# Development of GS-441524 Derivatives as Potent SARS-CoV-2 Mac1 Inhibitors via a Direct-to-Biology Approach

**DOI:** 10.64898/2026.06.24.734322

**Authors:** Kewen Peng, Suryadeep Chakraborty, Shamar D. Wallace, Jessica Caroline Gomes Noll, Jialin Shang, Xuan Lu, Annette Choi, Gary Whittaker, J. Christopher Fromme, Hening Lin

**Affiliations:** Department of Medicine, The University of Chicago, Chicago, IL 60637, USA; Center for Chemical Biology and Therapeutics, The University of Chicago, Chicago, IL 60637, USA; Department of Chemistry, The University of Chicago, Chicago, IL 60637, USA; Department of Chemistry and Chemical Biology, Cornell University, Ithaca, NY 14853, USA; Department of Molecular Biology and Genetics, Weill Institute for Cell and Molecular Biology, Cornell University, Ithaca, NY 14853, USA; Departments of Microbiology & Immunology, Cornell University College of Veterinary Medicine, Ithaca NY, USA; Howard Hughes Medical Institute, Department of Medicine, Department of Chemistry, The University of Chicago, Chicago, IL 60637, USA

## Abstract

Targeting viral macrodomains (Mac) has emerged as a promising strategy for antiviral drug development, especially after the outbreak of COVID-19 that claimed millions of lives worldwide. Several severe acute respiratory syndrome coronavirus 2 (SARS-CoV-2) Mac1 inhibitors have been reported in the past few years. In the present work, we converted GS-441524 (IC_50_ of ∼10 μM for SARS-CoV-2 Mac1) to KP-S54 (**18c**), a potent inhibitor of both SARS-CoV-2 Mac1 (IC_50_: 44 nM) and Middle East respiratory syndrome coronavirus (MERS-CoV) Mac1 (IC_50_: 91 nM) through an iterative direct-to-biology approach. This approach leverages efficient amide-coupling reaction and the mix-and-read fluorescence polarization (FP) assays where reaction mixtures could be screened directly without purification. Cocrystal structure of a selected derivative (**12p**) binding to SARS-CoV-2 Mac1 revealed the binding mode, which will guide future drug development against viral macrodomains.

## Introduction

A central defense mechanism of host cells against viral infection is the interferon (IFN) signaling pathway, which relies on several IFN-inducible adenosine diphosphate (ADP)-ribosyltransferases (ARTs or PARPs) and their ability to ADP-ribosylate protein substrates.^1–4^ Coronaviruses (CoVs) possess a family of highly conserved domains, known as the macrodomains (Mac), that can hydrolyze the mono-ADP-ribosylation (MAR) modification on proteins and hence counter the antiviral IFN signaling.^5^ The deMARylating activity of viral macrodomains has been shown to be indispensable for virulence as mutating or knocking out viral macrodomains significantly attenuates viral infections in cell and mouse models.^6–8^

Research from different labs has validated that macrodomain 1 (Mac1) of SARS-CoV-2, located on the non-structural protein 3 (nsp3), the largest protein in the coronavirus genome, can reverse IFN-induced ADP-ribosylation and promote viral replication.^9, 10^ Loss-of-function mutation or knocking out SARS-CoV-2 Mac1 significantly attenuated viral infection in cell cultures and in mouse models. ^6, 11^ The effects of Mac1 on SARS-CoV-2 infection in cells have been found to be highly dependent on the IFN-stimulating conditions, which is consistent with the induction of ADP-ribosylation by IFN and the functions of viral macrodomains as discussed above. Without IFN pretreatment, SARS-CoV-2 with Mac1 deletion or mutation replicates normally in cells, hinting that Mac1 inhibitors need to be evaluated in cellular infection assays under IFN-stimulation or directly in mice models.

Multiple SARS-CoV-2 Mac1 inhibitors have been reported in the literature with varying binding affinities. Wazir et. al.^12^ discovered a series of 2 amide-3-methylester thiophene derivatives as SARS-CoV-2 Mac1 inhibitors. Among them, MDOLL-0229 (Figure 1) is the strongest binder with binding affinity similar to ADPr but only showed limited activity in cellular assays. No mouse data was reported for this compound due to its unfavorable pharmacological properties like hydrolysis of the carboxylic acid ester moiety and other potential metabolic liabilities. A recent report disclosed a very potent SARS-CoV-2 Mac1 inhibitor, AVI-4206 (Figure 1), discovered through an extensive fragment-linking campaign and has an IC_50_ value of 20 nM.^13^ AVI-4206 shows moderate efficacy in both cellular assays and mouse models, suggesting that developing SARS-CoV-2 Mac1 inhibitors based on other scaffolds may be worthwhile.

**Figure 1.**
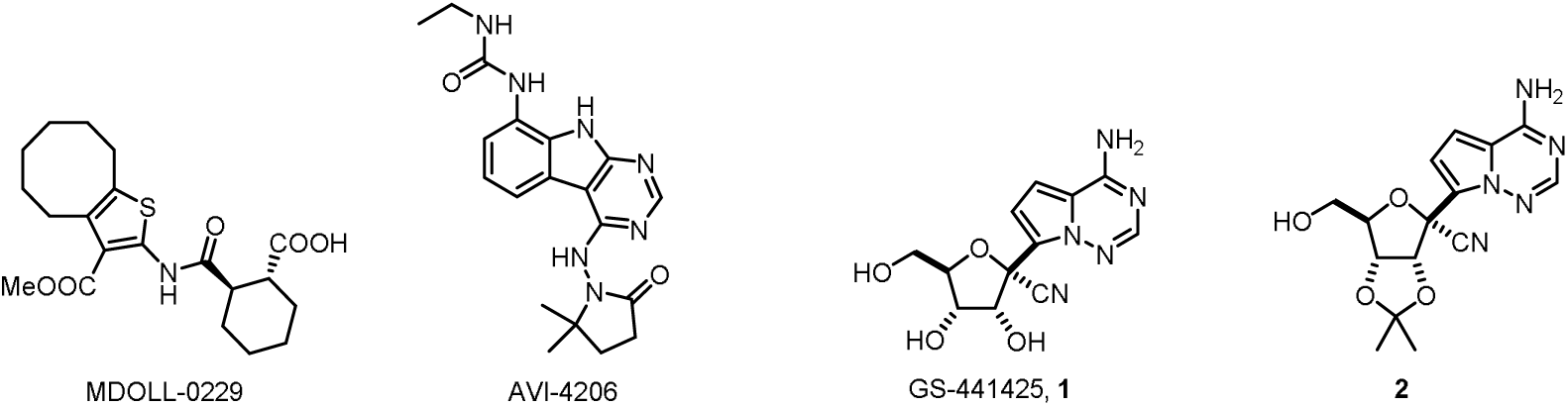
Chemical structures of MDOLL-0229, AVI-4206, GS-441524 (**1**) and 2’,3’-acetal protected GS-441524 (**2**).

A compound that we have been following is GS-441524 (Figure 1, **1**), which is an active metabolite of Remdesivir, an FDA approved anti-SARS-CoV-2 drug targeting RNA-dependent RNA polymerase (RdRp). GS-441524 was found to be a moderate binder of SARS-CoV-2 Mac1 with dissociation constant (*K_d_*) of about 10 μM, which is comparable to ADPr in isothermal titration calorimetry (ITC) experiments.^14^ Co-crystal structure shows this compound mimics the adenosine part of ADPr while its 1’-CN group forms extra hydrogen bonding interactions with the protein. Later studies confirmed that GS-441524 can inhibit the enzymatic activities of SARS-CoV-2 Mac1 as well as macrodomains from some other viruses albeit with lower potency.^15^ In our previous work,^16^ we synthesized some ADPr analogues of GS-441524 and obtained extremely potent binders of SARS-CoV-2 Mac1 with in vitro IC_50_ values down to single-digit nanomolar range. Interestingly, these compounds can also bind to macrodomains of Middle East respiratory syndrome coronavirus (MERS-CoV), Venezuelan equine encephalitis virus (VEEV), and Chikungunya virus (CHIKV). This result suggests GS-441524 has the potential of growing into tight-binding and possibly broad-spectrum antiviral drugs targeting viral macrodomains. However, the pyrophosphate moiety of these ADPr analogues makes them metabolic unstable and cell impermeable. Thus, new analogs with better metabolic stability and cellular permeability are needed.

Here we describe our effort to discover some potent and druglike GS-441524 derivatives using a direct-to-biology^17^ approach. Initial library construction of a GS-441524 5-NH_2_ analogue leveraging amide-coupling chemistry with hundreds of different carboxylic acids led to hit structures that share a glycine linker mimicking the pyrophosphate in ADPr. A second round of amide coupling-based compound library synthesis that introduced structural variability at the new amide bond yielded hits with better binding affinities. Further structural-activity relationship (SAR) studies led to compounds with nanomolar binding affinity for both SARS-CoV-2 Mac1 and MERS-CoV Mac1. X-ray crystal structure of a potent binder in complex with SARS-CoV-2 Mac1 protein shed light on the binding mode of the compound series developed. Further medicinal chemistry efforts led to KP-S54 (**18c**), a potent inhibitor for SARS-CoV-2 Mac1 (IC_50_: 44 nM) and MERS-CoV Mac1 (IC_50_: 91 nM) with improved liver microsomal stability and passive permeability.

## Results and Discussion

At the start of this project, the only reported binder with well-defined biophysical binding data for SARS-CoV-2 Mac1 was GS-441524 (Figure 1, **1**).^14^ We also confirmed its binding via fluorescence polarization (FP) assay using a TAMRA-labeled ADPr (ADPr-TAMRA) tracer, with data suggesting GS-441524 binds to SARS-CoV-2 Mac1 with IC_50_ of 15.2 μM, which is very close to that of ADPr (IC_50_: 15.5 μM) and consistent with literature. We later incorporated the GS-441524 nucleoside structure into the FP tracer and synthesized NDPr-TAMRA and Ph-NDPr-TAMRA (here N represents the GS-441524 nucleoside) which demonstrated over 100-fold affinity improvement over ADPr-TAMRA for not only SARS-CoV-2 Mac1 but also MERS-CoV, CHIKV, and VEEV macrodomains.^16^ Using the new FP tracers, we only need nanomolar concentrations of the viral macrodomain proteins for inhibitor screening, which bypassed the requirement for large amounts of proteins and enabled us to perform high-throughput screening assays against different viral macrodomains in a fast and mix-and-read fashion.

We decided to use GS-441524 as the starting point for SARS-CoV-2 Mac1 inhibitor design. GS-441524 is a very polar compound with multiple hydrogen bonding donors that potentially compromise its passive permeability. Although active transport mechanisms^18, 19^ might help the permeation of nucleoside analogues like GS-441524, our modified GS-441524 derivatives are unlikely to be actively transported into cells. Therefore, compounds’ lipophilicity was one of the key parameters we wished to optimize. Judged by the X-ray crystal structure of SARS-CoV-2 Mac1 in complex with GS-441524,^14^ the 2’,3’-hydroxyl groups have no contribution to the binding and can potentially be blocked to confer higher lipophilicity. Indeed, acetal-protected GS-441524 (Figure 1, **2**), a reported intermediate for Remdesivir synthesis,^20^ maintained the binding affinity for SARS-CoV-2 (IC_50_: 12 μM) and has much improved solubility in organic solvents, which allowed more efficient chemical reactions as well as easier compound purification. Thus, we decided to keep this acetal protection for all later derivatives.

**Figure 2.**
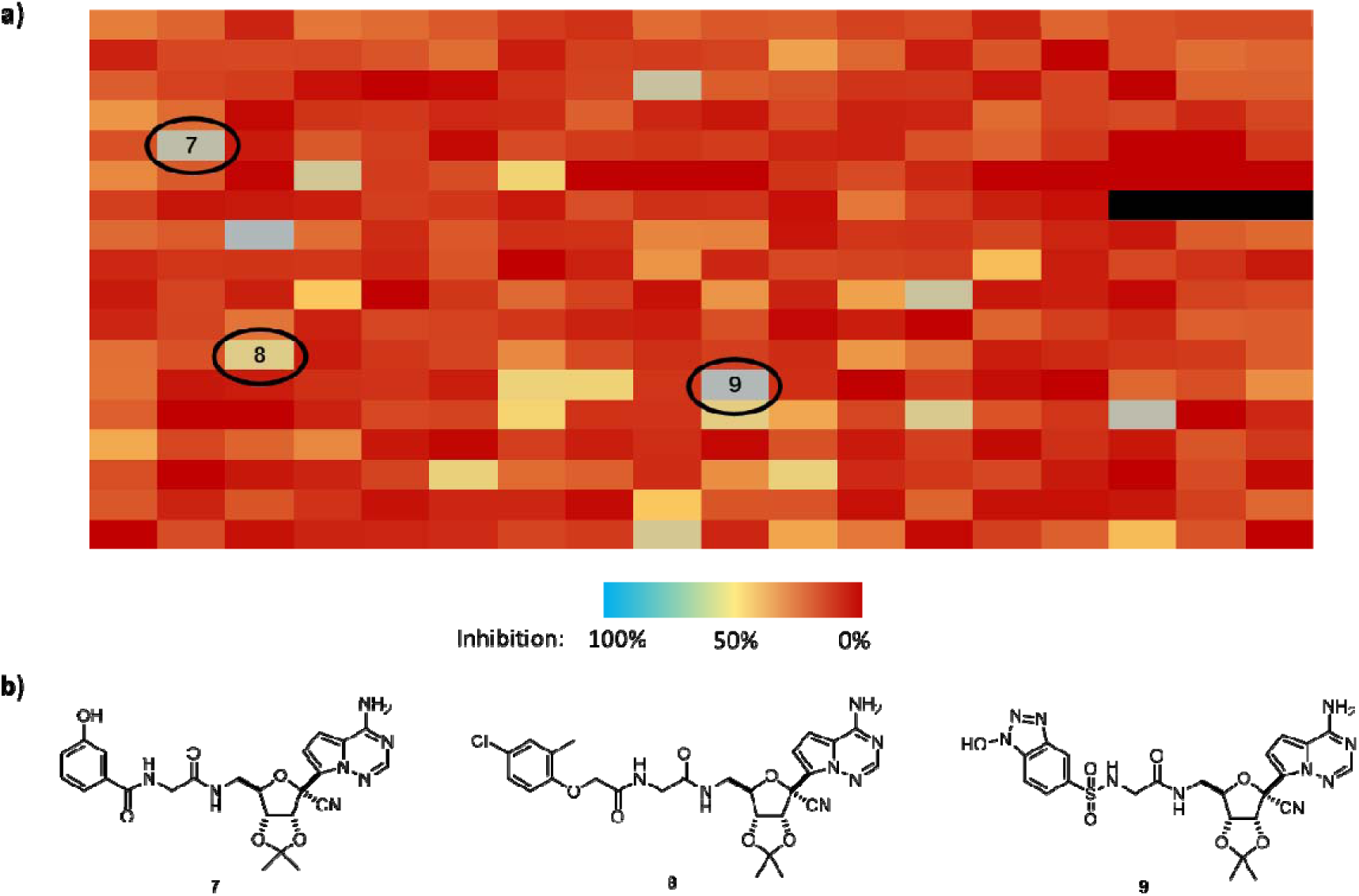
Results from the direct-to-biology FP screening against SARS-CoV-2 Mac1. **a**) Data from FP screening with selected hits (**7-9**) highlighted in ovals. **b**) Chemical structures of selected hits.

In order to quickly derivatize **2**, we converted its 5’-OH to an amine group leading to compound **6** that enabled library construction through simple and high-yielding amide coupling reactions with carboxylic acids. Additionally, we wished to mimic the phosphate linkage in NDPr with the amide moiety, since amides have been suggested to be excellent mimics of phosphate linkages in RNAs.^21^ Synthesis of the amine building block **8** is depicted in Scheme 1. We started with GS-441524, converting its 5-OH to iodide through an Appel reaction with iodine and triphenylphosphine, followed by azide substitution using sodium azide yielding intermediate **4**. The 2’,3’-dioxyl of **4** was protected with acetal using 2,2-dimethoxypropane in acetone under acid catalysis. The 5’-azide of **5** was reduced to amine via Staudinger reduction, furnishing amine **6** in good overall yield.

**Scheme 1.**
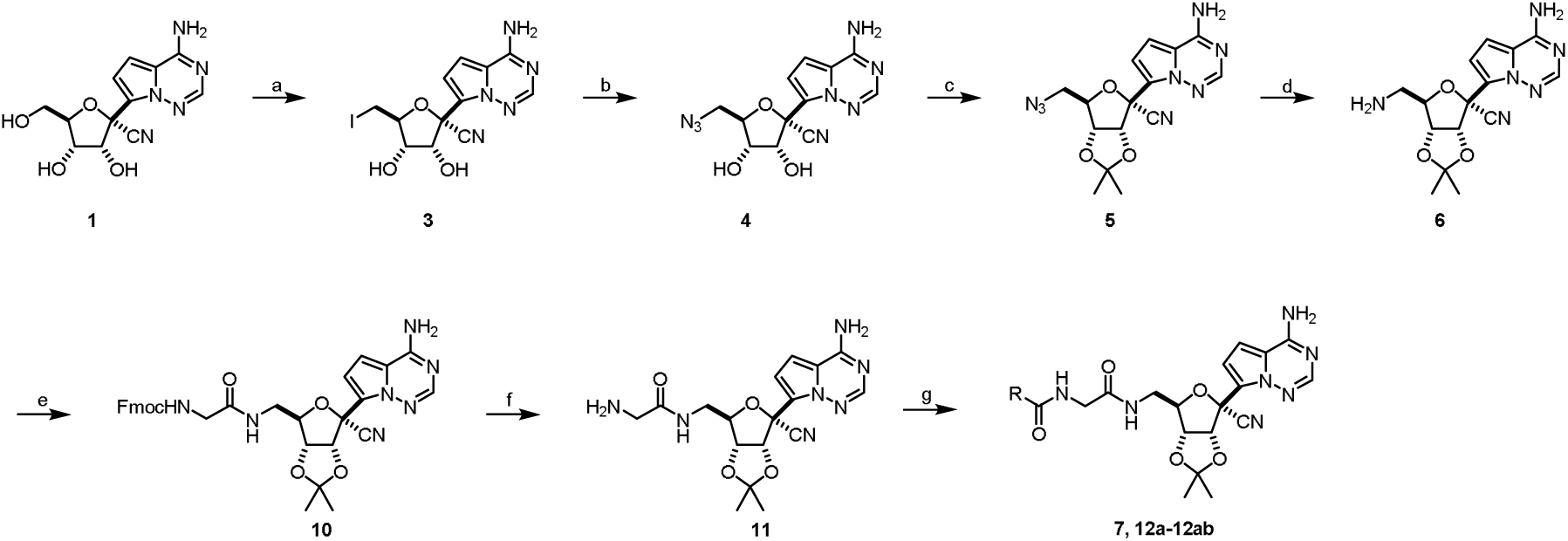
Synthesis of 5-NH_2_ GS-441524 (**6**) and its derivatives. Reagent and conditions: a) PPh_3_, I_2_, imidazole, DMF, 80□; b) NaN_3_, DMF, 50□; c) 2,2-dimethoxylpropane, acetone, conc. H_2_SO_4_, 40□; d) PPh_3_, THF, then water, reflux; e) Fmoc-Gly-OH, EDCI, HOBt, DIEA, DMF; f) 2N NHMe_2_ in THF, DCM; g) RCOOH, EDCI, HOBt, DIEA, DMF.

We performed amide coupling reactions of amine **6** with a panel of 318 carboxylic acids with high structural diversity in 96-well plate format using hexafluorophosphate azabenzotriazole tetramethyl uronium (HATU) as the condensing reagent, N,N-diisopropylethylamine (DIPEA) as the base and dimethyl sulfide (DMSO) as the solvent at a final concentration of 20 mM (see Materials and Methods for details). Among the 318 carboxylic acids used, 123 of them were from Enamine Essential Fragment Library, while the rest were chosen from Enamine in-stock building blocks based on structural similarity clustering to maximize the chemical space to be explored. LC-MS analysis of several randomly selected reactions all demonstrated high mass intensity of the amide products despite differential apparent conversion rates. The IC_50_ of amine **6** was tested to be >100 μM against SARS-CoV-2 Mac1 and hence unlikely to interfere with the screening readout if not fully converted. The crude reaction mixtures were directly screened in the FP assay at a theoretical concentration of 50 μM (with regard to the amide product assuming 100% conversion). We found the library generated is highly compatible with the high-throughput FP binding assay, as all the reagents used in the reactions were well tolerated by the assay and did not cause any interference. Moreover, the simple mix-and-read protocol enabled us to screen hundreds of reaction mixtures against four different viral macrodomains in a single day. The results from SARS-CoV-2 Mac1 screening are summarized in Figure 2. Some hits showed better apparent inhibitory activities, but later were determined to be false positives due to fluorescence quenching and were not further considered. Interestingly, three hits (Figure 2b, **7**-**9**) showed >50% inhibition and share a simple glycine linker and they all have another amide or sulfonamide linkage attached to the α-NH_2_ group of glycine.

We then synthesized compound **7** though the route depicted in Scheme 2. Briefly, **6** was coupled with Fmoc-Gly-OH, followed by Fmoc deprotection to yield key intermediate **11**. Compound **7** was prepared by coupling **11** with 3-hydroxybenzoic acid and later derivatives were prepared through the same method using different carboxylic acids. Testing the purified compound **7** confirmed that it is a genuine binder of SARS-CoV-2 Mac1 with IC_50_ of 4.2 μM. Interestingly, although GS-441524 does not bind MERS-CoV-2 Mac1, CHIKV Mac or VEEV Mac even at 100 μM,^22^ we found **7** is a strong binder for MERS-CoV-2 Mac1 with IC_50_ of only 1.6 μM but did not show any binding to CHIKV or VEEV macrodomains. This result can be rationalized by the fact that MERS-CoV is also a β-coronavirus and may share higher Mac1 structural similarity with SARS-CoV-2 than CHIKV or VEEV, both of which are α-coronaviruses. We reasoned that the two amide linkages can mimic the pyrophosphate linkage in NDPr and can be accommodated in the narrow pyrophosphate binding groove of SARS-CoV-2 or MERS-CoV Mac1, but may not be compatible with CHIKV or VEEV macrodomains. Nonetheless, compound **7** represents an excellent starting point for further derivatization towards more potent SARS-CoV-2 Mac1 inhibitors.

**Scheme 2.**
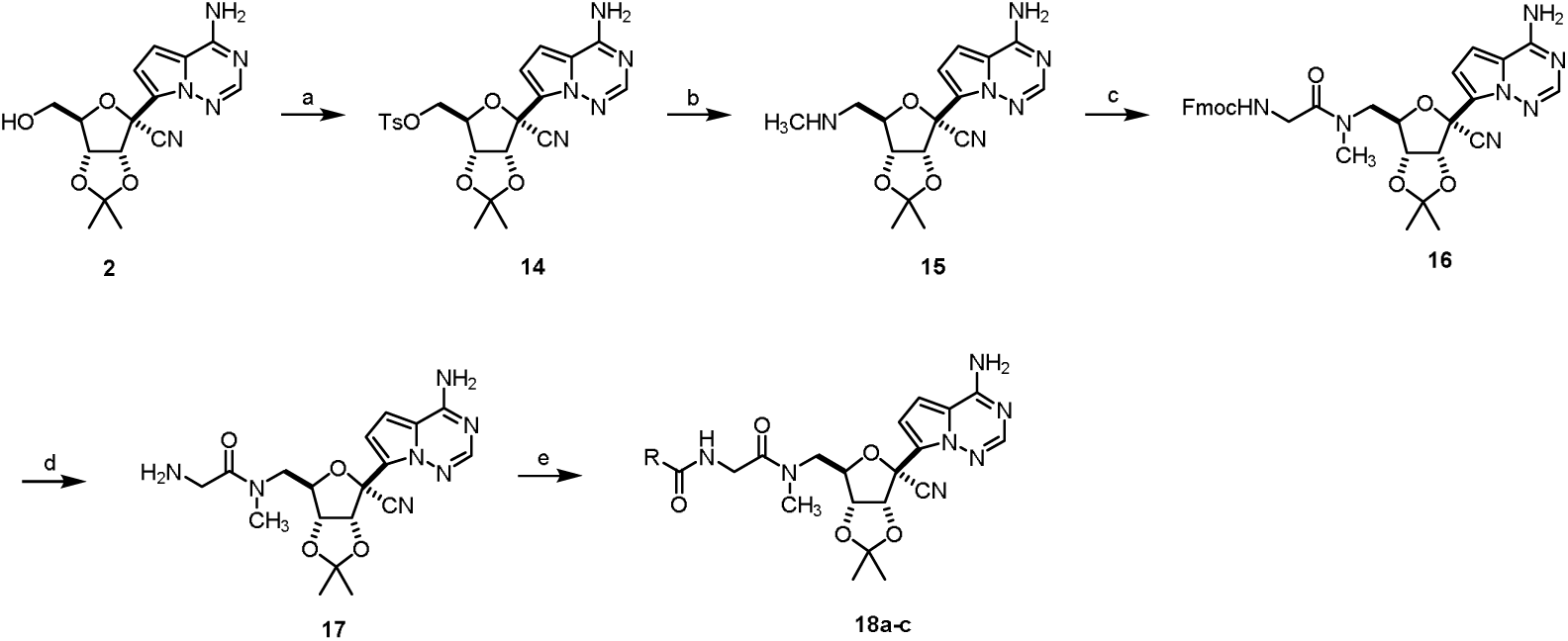
Synthesis of 5-NHCH_3_ GS-441524 derivatives **18a-c**. Reagent and conditions: a) TsCl, DMAP, DCM; b) MeNH_2_, THF, 100□/microwave irradiation; c) Fmoc-Gly-OH, EDCI, HOBt, DIEA, DMF; d) 2N NHMe_2_ in THF, DCM; e) RCOOH, EDCI, HOBt, DIEA, DMF.

The second amide linkage introduced in **7** provided a convenient handle for further derivatization. We set out to derivatize **7** first by conducting another direct-to-biology campaign with **11** and the same carboxylic acid building blocks used previously, also in 96-well plates. To our delight, we found that **12a** (Table 1), which was formed by reacting **11** with 5-hydroxynicotinic acid, showed 3-fold increase in binding affinity to both SARS-CoV-2 Mac1 and MERS-CoV Mac1 compared with **7**, suggesting the pyridine nitrogen is beneficial for binding.

**Table 1.**
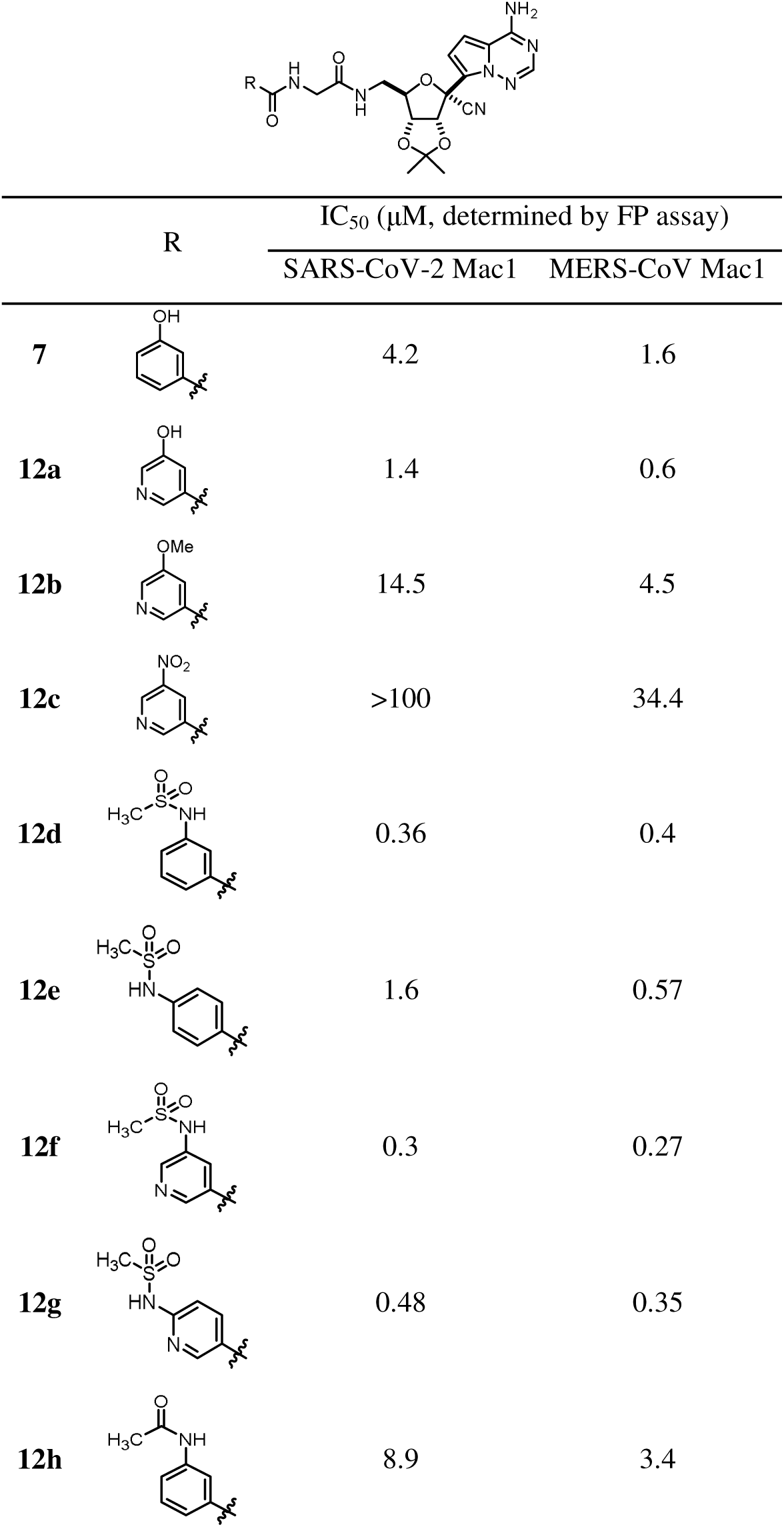
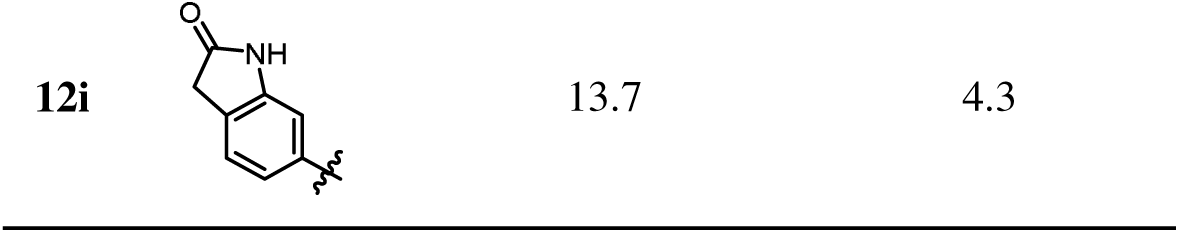
SAR of compounds 12a-i.

|  | R | IC<sub>50</sub> (μM, determined by FP assay) |  |
| --- | --- | --- | --- |
|  |  | SARS-CoV-2 Mac1 | MERS-CoV Mac1 |
| **7** |  | 4.2 | 1.6 |
| **12a** |  | 1.4 | 0.6 |
| **12b** |  | 14.5 | 4.5 |
| **12c** |  | >100 | 34.4 |
| **12d** |  | 0.36 | 0.4 |
| **12e** |  | 1.6 | 0.57 |
| **12f** |  | 0.3 | 0.27 |
| **12g** |  | 0.48 | 0.35 |
| **12h** |  | 8.9 | 3.4 |

The fact that the second round of direct-to-biology campaign did not yield much stronger inhibitors suggested that our carboxylic acid building blocks might still be limited in structural diversity. Therefore, we started to optimize **12a** through rational SAR studies. We first determined that the meta hydroxyl group is critical as compounds without it showed deteriorated binding activity. Blocking the hydroxyl group with a methyl group (**12b**) or replacing it with a nitro group (**12c**) led to much weaker activity, suggesting a hydrogen bonding donor is required at this position. Gratifyingly, attempt to mimic -OH in **7** with methyl sulfonamide (**12d**) led to >10-fold increase in activity for SARS-CoV-2 Mac1 and ∼4-fold increase in activity for MERS-CoV Mac1. Moving the -NHSO_2_Me from meta to para position (**12e**) is detrimental for SARS-CoV-2 Mac1 binding while MERS-CoV Mac1 binding is less affected. Switching the benzamide to nicotinamide led to slight increase in binding affinity (**12f** and **12g**). Interestingly, changing the sulfonamide group to carboxamide (**12h** and **12i**) led to a substantial decrease in activity, suggesting the sulfonamide not only mimics -OH as a hydrogen bonding donor but also impart additional interactions with the protein. Therefore, 3-sulfonamide appeared to be the optimal substituent on the aromatic ring.

Previously,^16, 22^ we found that attaching a simple phenyl ring at the C1’’-OH position can boost SARS-CoV-2 Mac1 binding by over ∼50 folds by forming hydrophobic interactions with an isoleucine sidechain from the distal ribose binding pocket. Judging from the overall geometry overlay of **12d** and ADPr, we hypothesized that the sulfonamide of **12d** may be located in or near the distal ribose binding pocket. Therefore, we replaced the terminal methyl group of **12d** with a benzene ring and synthesized **12j** (Table 2). Encouragingly, this compound showed 3-fold increase in binding affinity for SARS-CoV-2 Mac1. Adding a methylene between the benzene ring and the sulfonamide (**12k**) led to five-fold higher IC_50_ against SARS-CoV-2 Mac1 but binding for MERS-CoV Mac1 was retained. Replacing the benzamide moiety with nicotinamide (**12l**) led to a slight increase in binding activity as expected (IC_50_: 93 nM). Additionally, we found that **12l** has significantly improved water solubility (higher than 500 μM) over **12j** (less than 50 μM).

**Table 2.** SAR of compounds 12j-v.

|  | R | X | FP IC<sub>50</sub> (μM) |  | PAMPA P<sub>e</sub><br>(10<sup>-6</sup> cm/s) | MLM stability<br>(%remaining) |
| --- | --- | --- | --- | --- | --- | --- |
|  |  |  | SARS-CoV-2 Mac1 | MERS-CoV Mac1 |  |  |
| **12j** |  | CH | 0.12 | 0.23 | 4.10 | 3.9 |
| **12k** |  | CH | 0.59 | 0.32 | 3.68 | 0 |
| **12l** |  | N | 0.093 | 0.3 | 1.01 | 0 |
| **12m** |  | N | 0.12 | 0.23 | 1.16 | 0 |
| **12n** |  | N | 0.21 | 0.35 | 2.53 | ND |
| **12o** |  | N | 0.043 | 0.067 | 2.33 | 0 |
| **12p** |  | N | 0.019 | 0.078 | 3.11 | 0 |
| **12q** |  | N | 0.058 | 0.084 | 2.09 | 0 |
| **12r** |  | N | 0.048 | 0.105 | 3.90 | 30.1 |
| **12s** |  | N | 0.093 | 0.193 | 2.29 | 6.6 |
| **12t** |  | N | 0.18 | 0.29 | 0.82 | 72.0 |
| **12u** |  | N | 0.134 | 0.256 | 1.70 | 27.3 |
| **12v** |  | N | 0.141 | 0.352 | 0.82 | 19.3 |

**Table 3.**
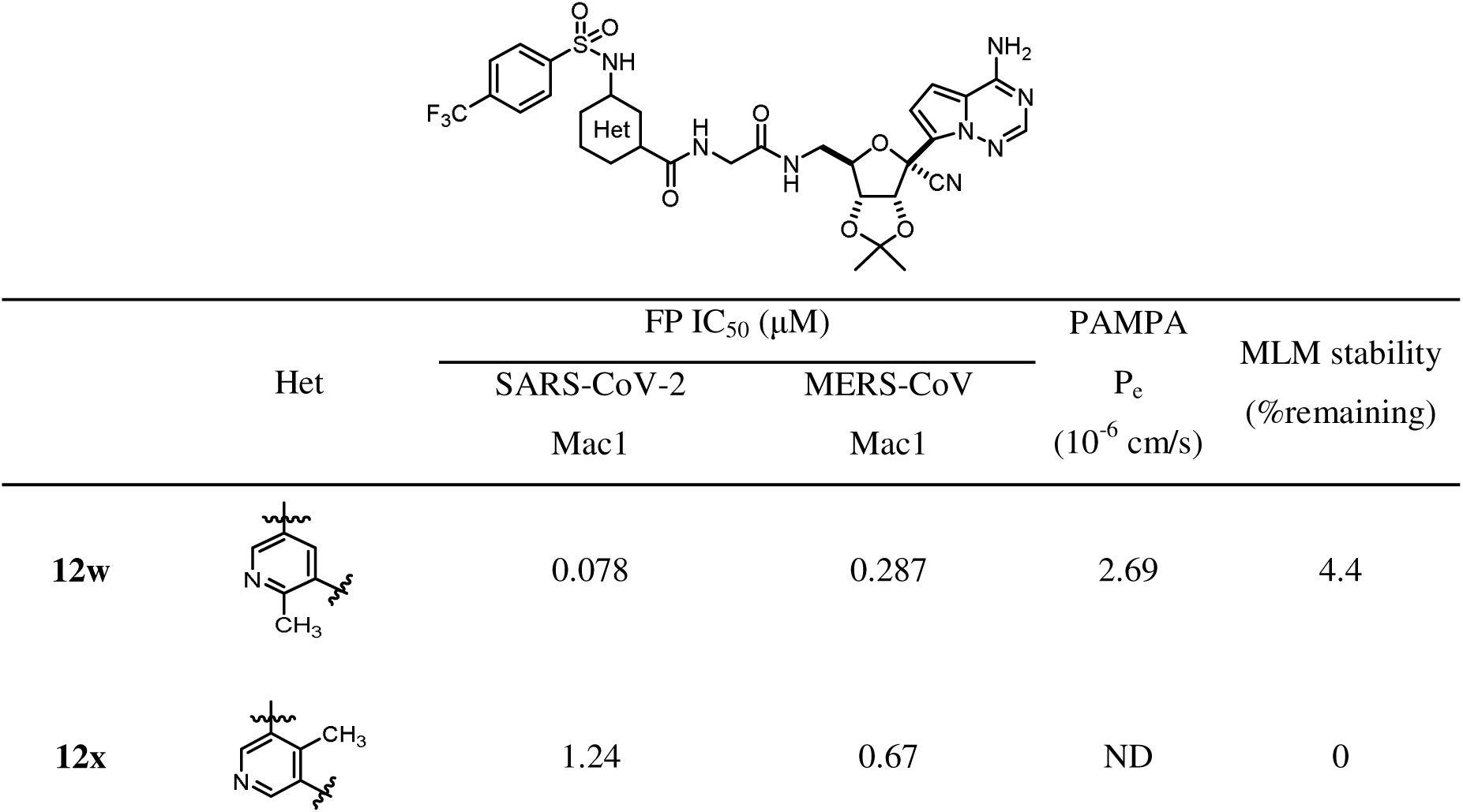
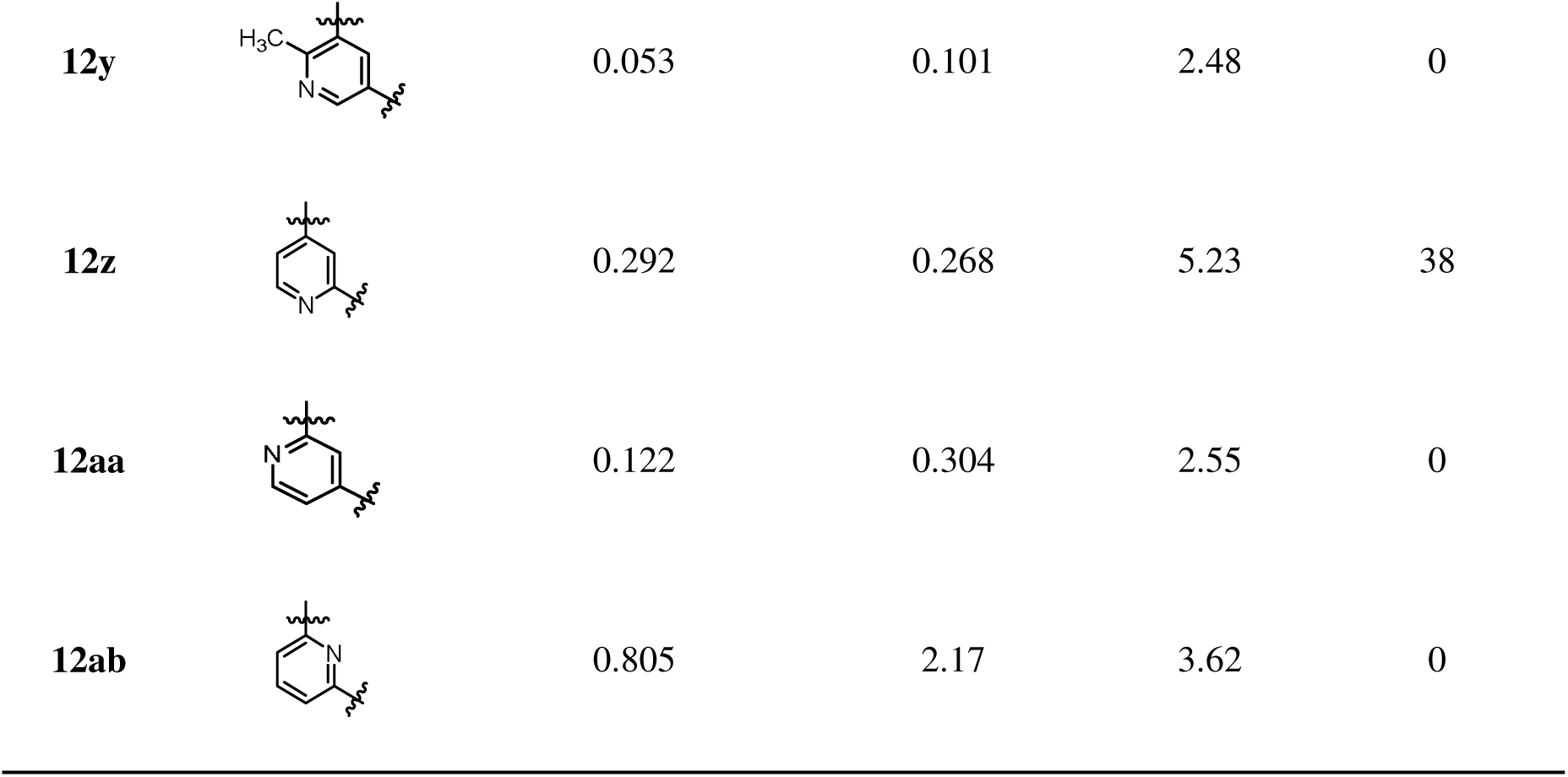
SAR of compounds 12w-ab.

|  | Het | FP IC<sub>50</sub> (μM) |  | PAMPA<br>P<sub>e</sub><br>(10<sup>-6</sup> cm/s) | MLM stability<br>(%remaining) |
| --- | --- | --- | --- | --- | --- |
|  |  | SARS-CoV-2<br>Mac1 | MERS-CoV<br>Mac1 |  |  |
| **12w** |  | 0.078 | 0.287 | 2.69 | 4.4 |
| **12x** |  | 1.24 | 0.67 | ND | 0 |
| **12y** |  | 0.053 | 0.101 | 2.48 | 0 |
| **12z** |  | 0.292 | 0.268 | 5.23 | 38 |
| **12aa** |  | 0.122 | 0.304 | 2.55 | 0 |
| **12ab** |  | 0.805 | 2.17 | 3.62 | 0 |

Now that inhibitors’ activity has already hit low nanomolar range, their pharmacological properties became more important than binding affinity in view of drug development. Hence, we started examining compounds’ passive permeability through parallel artificial membrane permeability assay (PAMPA) as well as their metabolic stability to mouse liver microsomes (MLM). We followed the lipid-oil-lipid trilayer PAMPA model developed by Chen et. al.,^23^ which has superior stability and better correlation to human oral drug absorption data. Although **12j** showed moderate effective permeability (P_e_) of 4.10 *10 ^-6^ cm/s, which is comparable to Remdesivir (see below), it was almost completely degraded in MLM after 15-min incubation. Similarly, **12l** was also unstable in MLM and showed poor permeability. We reasoned that the microsomal liability might come from the terminal benzene ring. Therefore, multiple different substituents were installed on the benzene ring to improve microsomal stability and also in the hope to optimize binding affinity. Incorporating a methoxyl (**12m**), nitro (**12n**) or fluorine (**12o**) at the meta position did not confer better microsomal stability, but it was interesting to see that fluorine (**12o**) boosted binding for SARS-CoV-2 Mac1 and MERS-CoV Mac1. Notably, moving the fluorine from meta to para position (**12p**) further boosted binding, yielding KP-S43 with IC_50_ of only 19 nM against SARS-CoV-2 Mac1 and 78 nM against MERS-CoV Mac1, albeit it was still unstable to MLM. The 3,4,5-trifluoro derivative (**12q**) was synthesized and tested but still showed no improvement in microsomal stability. Encouragingly, **12r** with a para trifluoromethyl group increased stability to MLM, having 30% remaining after 15-min incubation. Moreover, **12r** maintained moderate PAMPA permeability and high affinity to both SARS-CoV-2 Mac1 (IC_50_: 48 nM) and MERS-CoV Mac1 (IC_50_: 105 nM). Switching the benzene ring to a pyridine ring led to a microsomally stable compound **12t** but its permeability became poor. Adding an additional -CF_3_ on the pyridine ring (**12u**) helped with permeability but its microsomal stability was compromised.

We then further derivatized the nicotinamide core of **12r** (Table 2). Adding a methyl group at the 2 (**12w**) or 6 position (**12y**) maintained binding affinity but microsomal stability and PAMPA permeability deteriorated. A methyl group at the 4 position (**12x**), on the other hand, was not well tolerated, indicative of a steric clash with the binding pocket. This was later confirmed by cocrystal studies with SARS-CoV-2 Mac1 (see Figure 3). Moving the pyridine nitrogen atom to different positions did not yield more active compounds. Notably, **12z** with a picolinamide core demonstrated better microsomal stability and permeability compared with **12r** but its binding affinity for SARS-CoV-2 Mac1 decreased by 5-fold.

**Figure 3.**
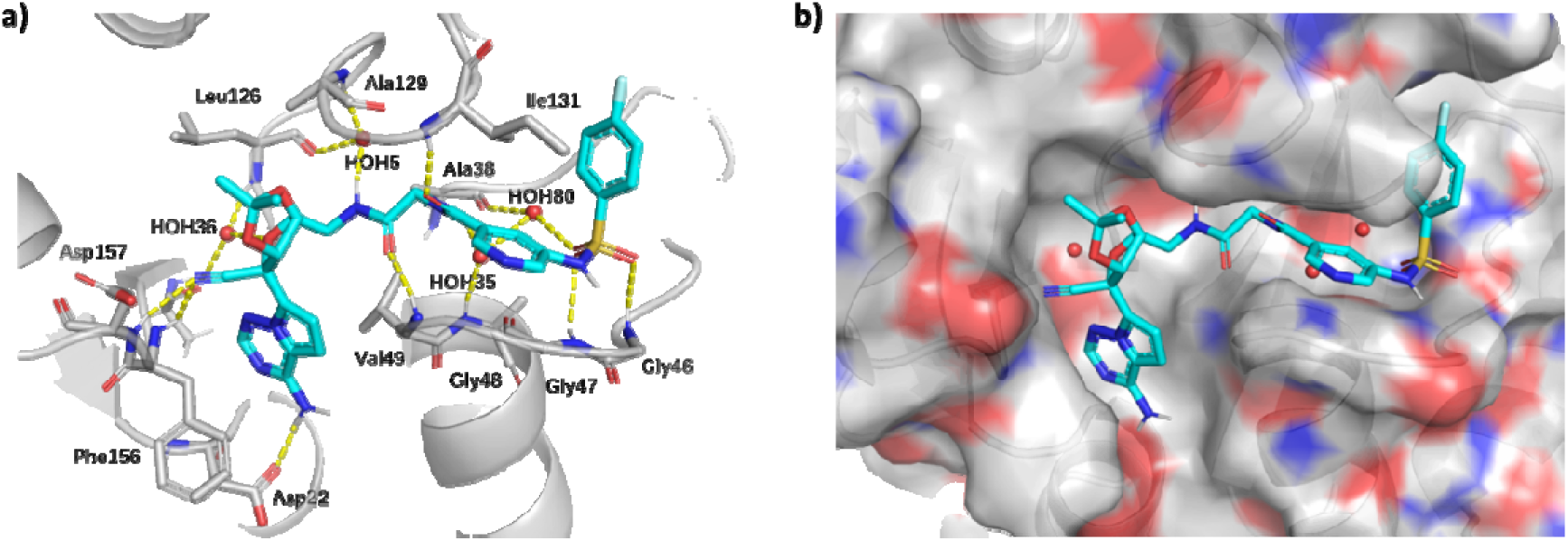
X-ray crystal structure of SARS-CoV-2 Mac1 in complex with **12p**. a) Binding interactions between **12p** and SARS-CoV-2 Mac1. Carbon atoms of **12p** and SARS-CoV-2 Mac1 are colored in cyan and gray, respectively. Structural water molecules are shown as red spheres. Hydrogen bonding interactions are shown as yellow dashes. b) Surface representation of a).

In order to increase compounds’ lipophilicity and hence permeability, we blocked either of the two carboxamide NHs in **12j** with a methyl group (**13** and **18a**, Table 4). Synthesis of **13** was analogous to previous 5-NH_2_ GS-441524 derivatives (Scheme 1) by reacting **6** with Fmoc-Sar-OH instead of Fmoc-Gly-OH, while synthesis of **18a** and its derivatives required slight modifications to the route as depicted in Scheme 2. Briefly, acetal-protected GS-441524 (**2**) was converted to 5-tosylate followed by substitution with methyl amine and condensation with Fmoc-Gly-OH. Fmoc deprotection of **16** and coupling to corresponding carboxylic acids yielded **18a-c**. Interestingly, blocking the NH near the ribose side (**18a**) led to 2-fold increased binding affinity to both macrodomains but putting a methyl group near the nicotinamide side (**13**) is detrimental for binding. Moreover, **18a** demonstrated a much higher P_e_ of 12.49 × 10^-6^ cm/s. We then incorporated the favorable methyl group into **12p** and **12r** with a nicotinamide core, yielding compounds **18b** and KP-S54 (**18c**), respectively. As expected, both **18b** and **18c** showed improved binding. **18b** has IC_50_ values of only 7.8 nM against SARS-CoV-2 Mac1 and 19.8 nM against MERS-CoV Mac1, which represents the strongest binder in the compound series. We also tried to replace the methyl group in **18b** with a slightly larger cyclopropyl group, but the IC_50_ increased by ∼50-fold. Although **18b** is the strongest binder, its P_e_ and MLM stability is less than ideal. On the other hand, KP-S54 (**18c**) with a trifluoromethyl group has significantly better P_e_ and MLM stability, albeit with slightly weaker binding affinity (44 nM for SARS-CoV-2 Mac1 and 91 nM against MERS-CoV Mac1).

**Scheme 3.**
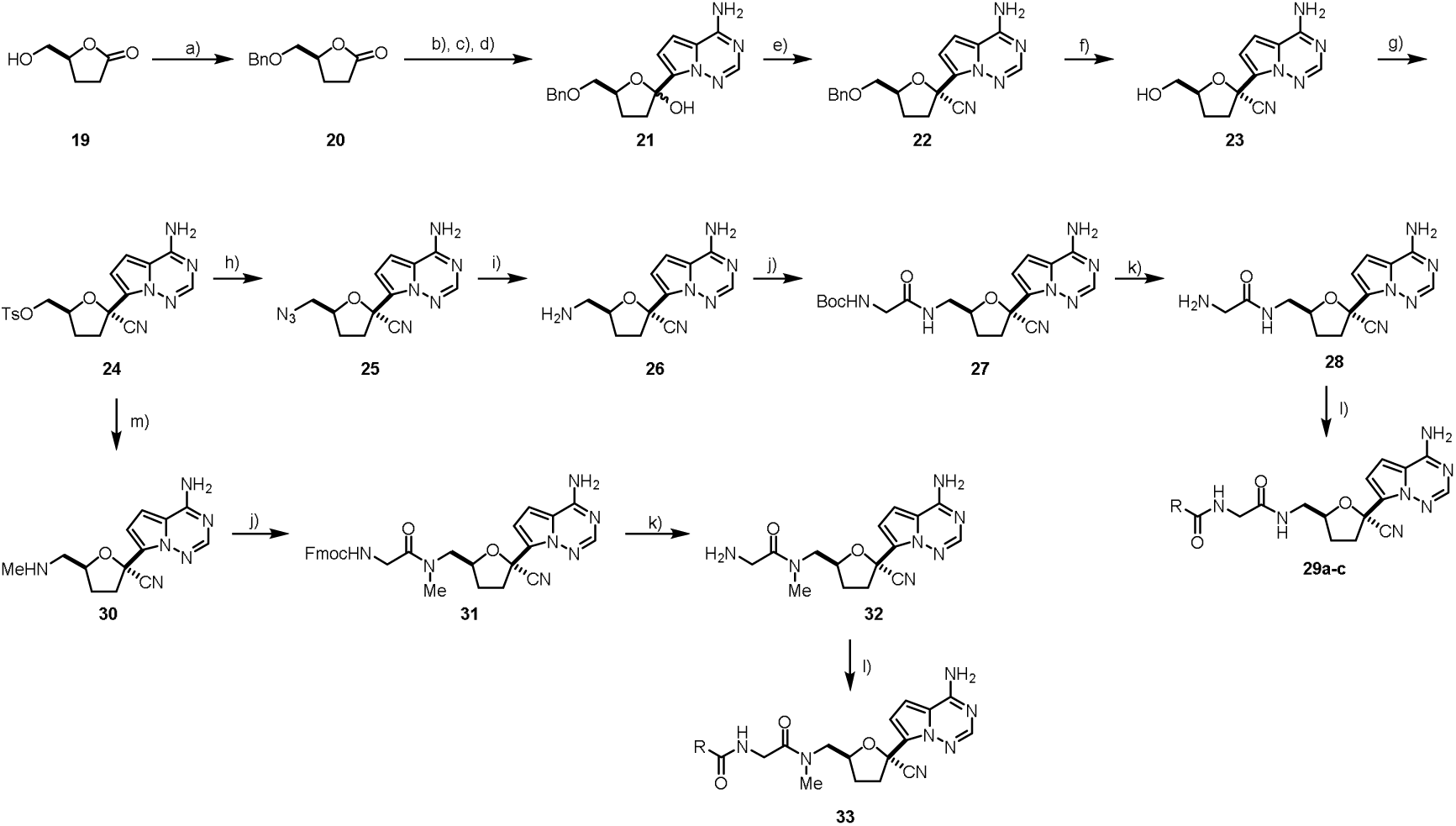
Synthesis of 2’,3’-dideoxy GS-441524 derivatives. Reagents and conditions: a) BnBr, NaH, TBAI, DCM; b) N,O-dimethylhydroxylamine hydrochloride, iPrMgCl, THF, 0□; c) TMSCl, imidazole, DCM, -15□; d) 7-iodopyrrolo[2,1-f][1,2,4]triazin-4-amine, TMSCl, MeMgBr, iPrMgCl·LiCl, THF, -15□ to 0□, then aq. HCl; e) TfOH, TMSOTf, TMSCN, DCM, -78□; f) BCl3, DCM, -78□; g) TsCl, DMAP, DCM; h) NaN3, DMF, 80□; i) PPh3, THF, then water, reflux; j) Fmoc-Gly-OH, EDCI, HOBt, DIEA, DMF; k) NHMe_2_ in THF, DCM; l) RCOOH, EDCI, HOBt, DIEA, DMF; m) MeNH_2_, THF, 100□/microwave irradiation.

**Table 4.**
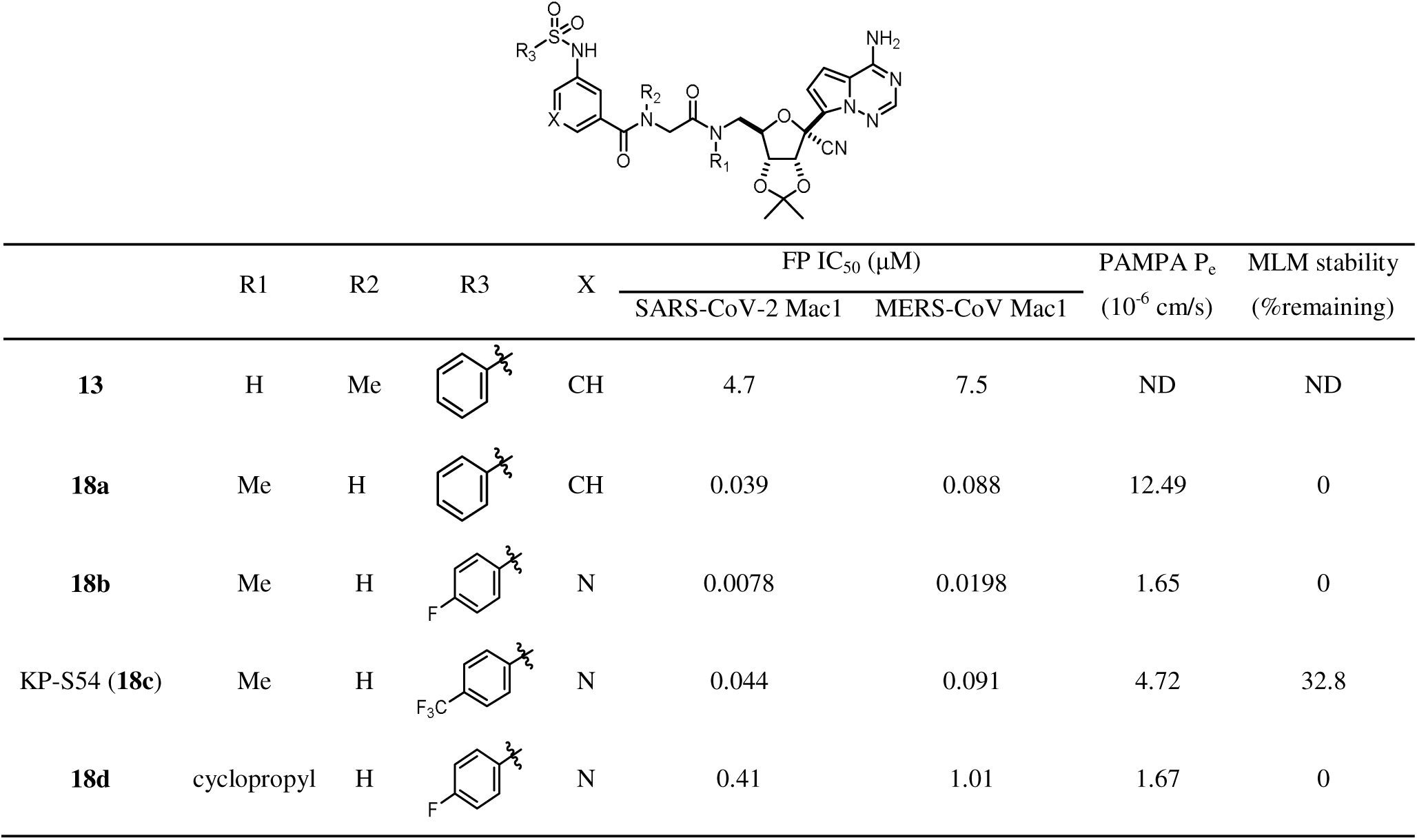
SAR of compounds 13 and 18a-c.

|  | R1 | R2 | R3 | X | FP IC<sub>50</sub> (μM) |  | PAMPA P<sub>e</sub><br>(10<sup>-6</sup> cm/s) | MLM stability<br>(%remaining) |
| --- | --- | --- | --- | --- | --- | --- | --- | --- |
|  |  |  |  |  | SARS-CoV-2 Mac1 | MERS-CoV Mac1 |  |  |
| **13** | H | Me |  | CH | 4.7 | 7.5 | ND | ND |
| **18a** | Me | H |  | CH | 0.039 | 0.088 | 12.49 | 0 |
| **18b** | Me | H |  | N | 0.0078 | 0.0198 | 1.65 | 0 |
| KP-S54 (**18c**) | Me | H |  | N | 0.044 | 0.091 | 4.72 | 32.8 |
| **18d** | cyclopropyl | H |  | N | 0.41 | 1.01 | 1.67 | 0 |

**Table 5.**
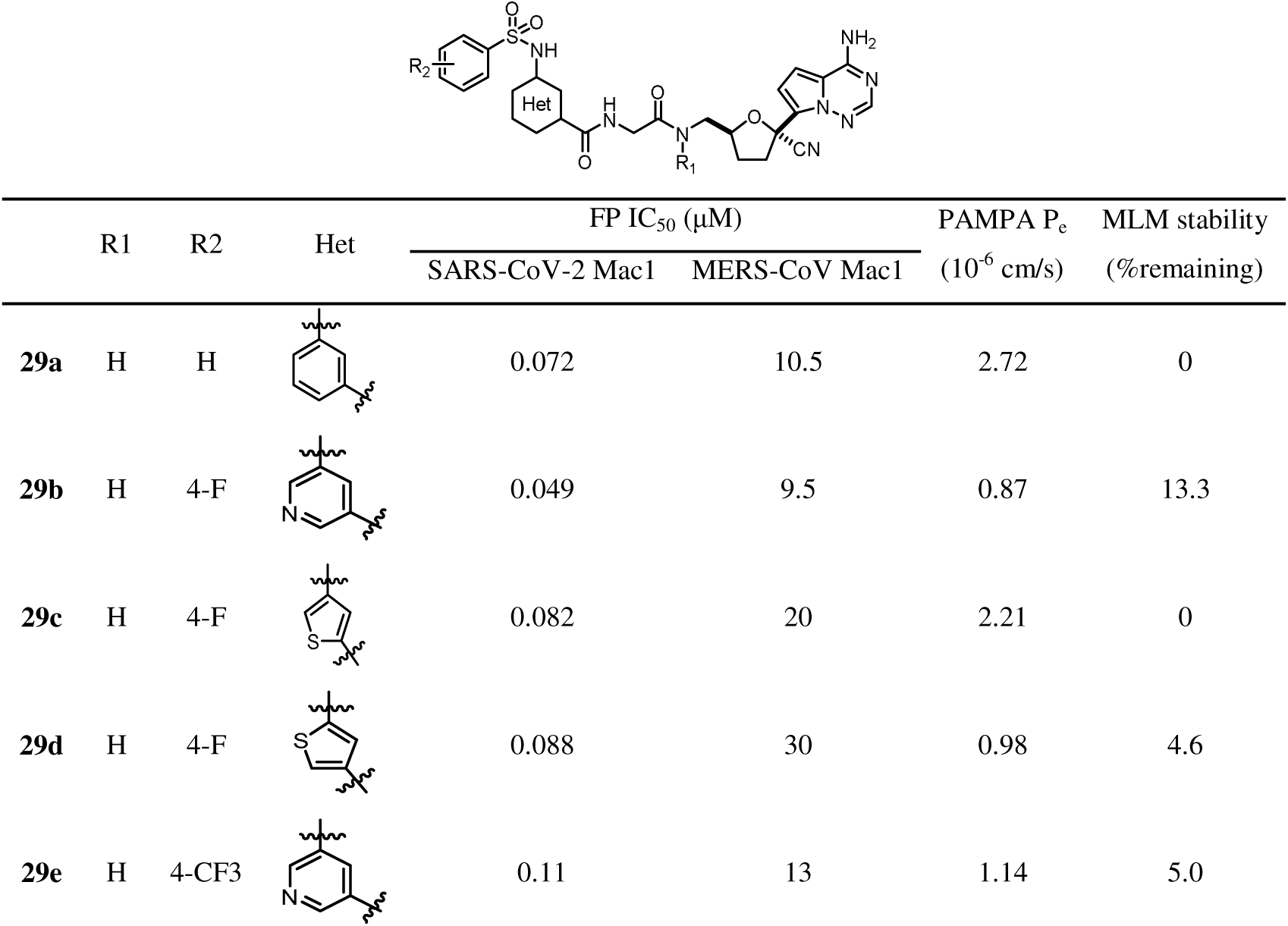
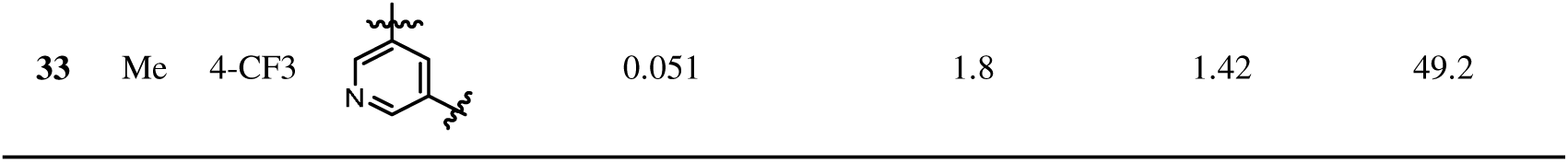
SAR of compounds 29a-e and 33.

| | R1 | R2 | Het | FP $IC_{50}$ (μM) | | PAMPA $P_e$<br>($10^{-6}$ cm/s) | MLM stability<br>(%remaining) |
| --- | --- | --- | --- | --- | --- | --- | --- |
|  |  |  |  | SARS-CoV-2 Mac1 | MERS-CoV Mac1 |  |  |
| **29a** | H | H |  | 0.072 | 10.5 | 2.72 | 0 |
| **29b** | H | 4-F |  | 0.049 | 9.5 | 0.87 | 13.3 |
| **29c** | H | 4-F |  | 0.082 | 20 | 2.21 | 0 |
| **29d** | H | 4-F |  | 0.088 | 30 | 0.98 | 4.6 |
| **29e** | H | 4-CF3 |  | 0.11 | 13 | 1.14 | 5.0 |

Considering the potential liability in gastric acid associated with the acetal moiety of the developed compounds if taken orally and the observation that 2’,3’-dioxy acetal moiety may have no contribution to macrodomain binding, we synthesized and tested several 2’,3’-dideoxy GS-441524 derivatives. Synthesis of these compounds necessitated a very different route as depicted in Scheme 3. The hydroxyl group of lactone **19** was first protected with a benzyl group, and the nucleophilic addition to the carbonyl group was achieved through a Weinreb amide approach reported by Xie et al. for the synthesis of a key remdesivir intermediate.^24^ Briefly, lactone **20** was converted into a Weinreb amide first and the resulting free -OH was tentatively protected with TMSCl before reacting with the heteroaryl Grinard reagent prepared from 7-iodopyrrolo[2,1-f][1,2,4]triazin-4-amine, followed by deprotection of the TMS group and ring closure in acid to yield intermediate **21**. For this conversion, we found this approach gave higher and more consistent yields than the condition used in the original route for Remdesivir synthesis.^20^ The dehydroxylative cyanation reaction of **21** was achieved with TfOH, TMSOTf and TMSCN at -78, but the stereoselectivity observed for GS-441524 derivatives was lost due to the absence of a 2’-oxygen atom that has been suggested to guide the stereoselective attack of TMSCN.^25^ Nonetheless, the α and β diastereomers (roughly in 1:1 ratio) could be separated by column chromatography. The -OBn of the desired β isomer was deprotected with boron trichloride at -78 to yield key intermediate 2’,3’-dideoxy GS-441524 (**23**), which then underwent tosylation and azide substitution to yield **25**. Staudinger reduction of azide **25** gave amine **26**, which was coupled with Boc-Gly-OH to yield **27** that was deprotected and subsequently coupled with different carboxylic acids to furnish target compounds **29a-e**. Similarly, **33** was prepared first by reacting tosylate **24** with methyl amine under microwave irradiation, followed by coupling with Boc-Gly-OH, deprotection and coupling to the corresponding carboxylic acid.

Interestingly, although the binding affinities of the 2’,3’-dideoxy derivatives for SARS-CoV-2 Mac1 are maintained, their affinity for MERS-CoV Mac1 is lost. Moreover, compared with their acetal counterparts, the 2’,3’-dideoxy compounds showed diminished PAMPA P_e_ and did not improve MLM stability much (compare **12j** with **29a**, **12p** with **29b**, **12r** with **29e** and **18c** with **33**). Additionally, we also attempted to replace the nicotinamide core with thiophene rings (**29c** and **29d**), but did not observe any improvement on binding affinity or liver microsomal stability. Taken together, we reasoned that the acetal-protected derivatives might be superior drug candidates for mouse studies compared with their 2’,3’-dideoxy counterparts considering their wider antiviral spectrum and better permeability, and the liability of the acetal moiety may be evaded by intraperitoneal (IP) injection or increasing dosage. Nonetheless, **33** might be a good oral drug candidate to treat SARS-CoV-2 infection, although further structural optimization might be needed to improve its permeability.

To shed light on how the GS-441524-glycine derivatives bind to SARS-CoV-2 Mac1 protein, we attempted to crystalize Mac1 with several derivatives with strong binding affinities and reasonable water solubility. We were able to obtain crystals and solve the structure of **12p** with SARS-CoV-2 Mac1. The binding pose is shown in Figure 3. Expectedly, the hydrogen bonding interactions with the Asp22 side chain and the backbone NH of Phe156 and Asp157 are maintained for the GS-441524 nucleoside part. The two amide carbonyl groups interact with the backbone NH of Val49 and Ala38, mimicking the interactions made by the phosphate moiety close to the adenosine portion of ADPr. The -SO_2_- moiety in **12p** interacts with the backbone NH of Gly46 and Gly47, which explains why changing the -OH in **7** with -NHSO_2_Me (**12d**) led to 10-fold increase in binding affinity. The terminal phenyl group can form hydrophobic interactions with Ile131 as expected. Notably, we found there are four structural water molecules (HOH36, HOH5, HOH35 and HOH80) that play important roles in the binding of **12p**. Among these, only HOH36 is maintained in ADPr binding to SARS-CoV-2 Mac1 while the other three are unique to **12p**. HOH5 interacts with Leu126, Ala129, and an -NH- from **12p**, but it is interesting that replacing the -NH- with a -NMe- group (**18b**) led to an increase in binding affinity, suggesting that displacing this water might contribute to stronger interactions. HOH35 and HOH80 form a hydrogen bonding network with Ala38, Gly48 and **12p**, which may contribute to the binding affinity.

Because compound KP-S54 has the best balance between Mac1 inhibition potency and desired pharmacological properties (permeability and liver microsomal stability), we next conducted a more thorough investigation on the drug-like properties of this compound by testing its liver microsomal stabilities in different species and comparing its microsomal stability and PAMPA permeability with selected reference drugs including Remdesivir, GS-441524 and Propranolol (Table 6). Although KP-S54 showed less-than-ideal mouse liver microsomal stability with half-life of only 17 min, its stability in rat and human liver microsomes are modestly better (half-lives of 41 min and 35 min, respectively). In line with literature report, Remdesivir was immediately and completely degraded upon incubation with different liver microsomes, while GS-441524 showed high cross-species liver microsomal stability. In terms of PAMPA permeability, KP-S54 (P_e_: 4.7 * 10^-6^ cm/s) is only slightly less permeable than Remdesivir (P_e_: 6.0 * 10^-6^ cm/s) but significantly better than GS-441524 (P_e_: 1.5 * 10^-6^ cm/s). The metabolic stability and permeability of KP-S54 (**18c**) prompted us to do some pilot tests in mouse models.

**Table 6.** Liver microsomal stability and PAMPA permeability of KP-S54, Remdesivir, GS-441524 and Propranolol.

|  | Liver microsomal half life (min) |  |  | PAMPA P<sub>e</sub><br>(10<sup>-6</sup> cm/s) |
| --- | --- | --- | --- | --- |
|  | Mouse | Rat | Human |  |
| KP-S54 (**18c**) | 17 | 41 | 35 | 4.7 |
| Remdesivir | <5 | <5 | <5 | 6.0 |
| GS-441524 | >60 | >60 | >60 | 1.5 |
| Propranolol | 17 | 31 | 46 | 21 |

Five mice (3 males, 2 females) were subjected to KP-S54 (100 mg/kg) or vehicle treatment intraperitoneally once daily starting from 1 day before SARS-CoV-2 infection. To our disappointment, mice from both the control and treated groups consistently lost weight starting from 2 days post-infection (Figure S1). All surviving animals reached the 20% weight loss cut-off point for euthanasia by day 6 post-infection. The PCR results showed similar viral loads in the lungs and the brains between the control and treated groups (Figure S2). The inactivity of KP-S54 in mice may be largely attributed to its poor pharmacokinetic profile. The drug concentration only reached ∼0.3 μg/mL (∼0.4 μM) one hour after IP injection (50 mg/kg) and was rapidly eliminated within 6 hours (Figure S3). This result is consistent with the data from mouse microsomal stability experiments where the compound’s half-life was less than 20 minutes. Therefore, further structural modification of KP-S54 to improve its PK profile is warranted.

The inhibitors we discovered here represent potent tool compounds that can be used to study SARS-CoV-2 and MERS-CoV *in vitro*. While the in vivo efficacy of KP-S54 is not as good as the previously reported AVI-4206, its ability to potently inhibit both SARS-CoV-2 and MERS-CoV macrodomains justifies its future development as it may lead to inhibitors with broad specificity. The structural information disclosed herein will aid in the design of better viral macrodomain inhibitors in the future. Moreover, our work provides an example that the direct-to-biology approach, leveraging efficient chemical reactions and robust screening assays, can greatly accelerate lead discovery.

## Materials and Methods

### Liver microsomal stability assay

To a microcentrifuge tube was sequentially added 513 μL phosphate buffer (100 mM, pH 7.4), 12 μL compound to be tested in water (50 μM, 1 μM final), and 60 μL NADPH in PB buffer (10 mM, 1 mM final). The tube was incubated at 37 for 15 minutes before adding 15 μL liver microsome in PB buffer (20 mg/mL, final 0.5 mg/mL) to initiate reaction. The final reaction volume is 600 μL and the tube was kept at 37 . At 0, 5, 10, 15, 30 min, 60 μL of reaction mixture was taken out and quenched with 60 μL acetonitrile and vortexed. The collected samples were centrifuged at 17,000 g for 5 minutes and the compound remaining at each time point was quantified by LC-MS. Half-life was obtained by plotting the graph of log (compound peak area) vs. incubation time. For stability screening, only 0 min and 15 min were tested and the reaction volume was scaled down to 150 μL accordingly.

### Parallel artificial membrane permeability assay

The tri-layer PAMPA assay was adapted from the method developed by Chen et. al.^23^, which uses a lipid-oil-lipid tri-layer assembled on a 96-well polycarbonate filter plate (Sigma, catalogue number: MPC4NTR10). Firstly, 20 μL of 5% hexadecane in hexanes was added to each well. The plate was put in a fume hood for 1 hour to evaporate hexanes. Then 10 μL of 5 mg/mL egg yolk lecithin (50 μg) was added and hexanes was evaporated again. The plate was flipped and 5 mg/mL egg yolk lecithin (50 μg) was added to the other side of each well and the plate was ready for use after. To each well of the filter plate (donor plate) was added 150 μL 500 μM compound in assay buffer (100 mM PB pH 7.4 + 2% DMSO). For less soluble compound, saturated solution in assay buffer was used instead. Separately, to each well of an acceptor plate was added 300 μL of assay buffer. The donor plate was carefully put on top of the acceptor plate, ensuring no gap or bubbles between the bottom of filter plate wells and the liquid surface of accepter wells. The sandwich was incubated in a humid chamber for 5 hours. For each assay compound, 100 μL solution from both top and bottom plates was pipetted to a UV-transparent 96-well plate and UV absorbance at 280 nm was measured as estimation of drug concentration. The apparent permeability was calculated using the following equation:

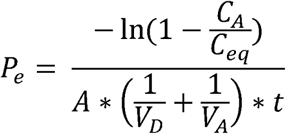

where P_e_ is permeability in the unit of cm/s; A = effective filter area; V_D_ = donor well volume; V_A_ = acceptor well volume; t = incubation time (s); C_A_ = compound concentration in acceptor well after incubation, and C_eq_ is the equilibrium concentration as if there was no membrane, calculated with following equation:

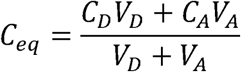

where C_D_ = compound concentration in donor well after incubation.

### Amide-coupling library construction

The amide library was constructed in 96-well plate format. Briefly, 1 μL of 100 mM carboxylic acid building blocks in DMSO was transferred to 96-well PCR plates, followed by the addiction of 3 μL of DMSO solution containing 33.3 mM amine and 66.6 mM DIPEA. The plates were centrifuged, and the reaction was initiated by adding 1 μL of 100 mM HATU in DMSO to each well. The final concentration of the carboxylic acid, amine, HATU and DIPEA are 20 mM (1 eq.), 20 mM (1 eq.), 20 mM (1. eq.) and 40 mM (2 eq.), respectively, and the final reaction volume was 5 μL. The plates were sealed and allowed to stand overnight at room temperature. LC-MS analysis of selected wells was conducted to examine reaction conversion.

### Expression and purification of macrodomains

The macrodomain proteins were expressed and purified as previously described.^16^

### Fluorescence polarization-based binding assay

The procedure for FP assay was as previously reported.^16^ For high-throughput screening of amide coupling-library, the reaction mixture was diluted with the FP buffer and directly assayed in 384-well plates. We used 30 nM protein and 20 nM **TAMRA-NDPr** for SARS-CoV-2 Mac1 screening, and 20 nM protein and 20 nM **TAMRA-Ph-NDPr** for MERS-CoV Mac1 screening.

### Co-crystallization of SARS-CoV-2 Mac1 bound to 12p

SARS-CoV-2 Mac1 was mixed with compound **12p** in buffer (5 mM HEPES-NaOH pH 7.5, 150 mM NaCl) to final concentrations of 0.056 mM and 1 mM respectively. The protein-inhibitor mixture was incubated at room temperature for 1 hr. and centrifuged at ∼ 16,000 x g for 5 minutes at 4 °C. The supernatant was concentrated until the Mac1 concentration was ∼ 1.75 mM. The Mac1-compound 12p complex was crystallized by the hanging-drop method at 20 °C by mixing 1 μL of the concentrated Mac1-**12p** solution with 1 μL of well solution (200 mM sodium acetate, 100 mM Tris−HCl pH 8, and 31% (w/v) PEG-4000). Needle-like and cube-shaped crystals were observed after 1-2 days. Before freezing with liquid nitrogen, crystals were cryo-protected in well solution containing 10% ethylene glycol and 1 mM compound **12p**. Only cube-shaped crystals were used for diffraction.

### Diffraction Data Collection, Structure Solution, Model Building, and Refinement

Diffraction data was collected on beamline ID7B-2 at the Center for High-Energy X-ray Sciences (CHEXS) at the Cornell High Energy Synchrotron Source (CHESS). Initial data processing was performed using Fast DP.^26–29^ The structure was solved by molecular replacement using Phaser^30^ in Phenix^31^ with a previously published structure of SARS-CoV-2 Mac1 (PDB: 8GIA).^32^ Coot^33^ was used for model building, and refinement and validation were performed in Phenix.^33^ For this structure, there are 2 copies of Mac1 bound to compound **12p** in the asymmetric unit. The final structures of both copies are nearly identical except for the position of the fluorophenyl group, which adopts slightly different conformations in each copy.

### Mouse anti-viral experiments

K18 female mice (The Jackson Laboratories) were randomly assigned to an experimental group and a negative control group. Mice in the experimental group were intraperitoneal administered **12p** at 100mg/kg daily starting from 1 day before SARS-CoV-2 intranasal infection (5x10^2^ PFU). The negative control mice were treated with only the vehicle (80% PBS, 10% DMSO and 10% Kolliphor EL). Mice were assessed daily for clinical signs and weight measurement. On day 6 post-infection, all mice were euthanized. Mouse brain and lung tissues were harvested for viral load quantification.

### Viral load quantification

Lung and brain tissues from the mice used in this study were harvested and stored at -80 °C until processing. RNA was extracted from the tissues in the Cornell BSL-3 facility using trizol and chloroform. The viral load was assessed using the Luna SARS-CoV-2-RT-qPCR Multiplex Assay Kit (New England Biolabs) according to the manufacturer’s instructions.

### Pharmacokinetic Studies

The pharmacokinetic study of KP-S54 was conducted with IP (50 mg/kg) dosing C57BL/6 mice (n = 2) using a formulation of 10% DMSO, 10% Kolliphor EL and 80% PBS. To quantify drug concentration in the plasma, ∼200 μL of blood was drawn from the eyeballs of each mouse at 1, 3 and 6 hours after injection. Blood samples were centrifuged at 21,000 g for 15 min at 4 °C and the supernatant was treated with an equal volume of methanol. The tubes were vortexed thoroughly and centrifuged at 21,000 g for 15 min at 4 °C and the supernatant was again centrifuged at 21,000 g for 15 min at 4 °C. Finally, 100 μL of the supernatant was injected into the LC-MS for compound quantification.

### General synthetic methods

Unless otherwise stated, all reactions were carried out under ambient atmosphere. Reagents and solvents were purchased from Sigma Aldrich, TCI, Alfa-aesar, Combi-blocks, Chem-impex international, Ambeed, VWR, and Fisher scientific and were used as received. Thin layer chromatography (TLC) was performed on precoated glass-backed silica gel 60 F254 plates (EMD Millipore), visualized by UV or staining with 10% H_2_SO_4_ in ethanol. SiliaFlash^®^ P60 (SiliCycle) silica gel was used for column chromatography purification. Alternatively, pre-packed C18 or silica gel columns were used in combination with a CombiFlash flash chromatography system. All unreported compounds were characterized by ^1^H-NMR, ^13^C-NMR and/or ^31^P-NMR spectra recorded on Bruker spectrometers running at 400, 500 or 600 MHz. Mass spectra (MS) were obtained on a LCQ Fleet Mass Spectrometer (Thermo Scientific) equipped with an electrospray ionization (ESI) source connected to a Shimadzu HPLC LC20-AD.

**General Procedure A** for the amide coupling reaction using EDCI and HOBt Amine (0.1 mmol, 1 eq.) and carboxylic acid (0.11 mmol, 1.1 eq.) were combined and dissolved in DMF (0.5 mL). DIPEA (0.22 mmol, 2.2 eq.) and HOBt (0.11 mmol, 1.1 eq.) were added, followed by the addition of EDCI (0.11 mmol, 1.1 eq.). The reaction mixture was allowed to stir at room temperature overnight. Water (2 mL) was added to the mixture, followed by extraction with ethyl acetate (EA, 2 mL, three times). The combined organic phase was washed with brine, dried over anhydrous Na_2_SO_4_ and evaporated to dryness. Silica gel column chromatography purification of the crude afforded the desired amide product. For target compounds with high water solubility and EA extraction was inefficient, the reaction mixture in DMF was directly subjected to reverse-phase purification on a C18 column using Combiflash.

### (2R,3R,4S,5S)-2-(4-aminopyrrolo[2,1-f][1,2,4]triazin-7-yl)-3,4-dihydroxy-5-(iodomethyl)tetrahydrofuran-2-carbonitrile *(3)*

GS-441524 (50 mg, 0.172 mmol, 1 eq.), PPh_3_ (67 mg, 0.257 mmol, 1.5 eq.) and imidazole (23 mg, 0.343 mmol, 2 eq.) were dissolved with THF (1 mL) in a sealed tube and the reaction mixture was heated to 80 with stirring. At this temperature, iodine (65 mg, 0.257 mmol, 1.5 eq) dissolved in THF (1 mL) was slowly added to the reaction mixture. The reaction mixture was stirred at 80 overnight and was directly evaporated under vacuum. The crude mixture was purified by flash column chromatography (DCM: MeOH 80:1 to 20:1) to yield product as a white solid (56 mg, yield: 81%)

**LC-MS (ESI)**: m/z calcd for C_12_H_13_IN O ^+^ [M+H]^+^ 402.0, found 402.1.

**^1^H NMR** (500 MHz, DMSO-*d*_6_) δ 7.94 (s, 1H), 7.89 (br, 2H), 6.92 (d, *J* = 4.5 Hz, 1H), 6.87 (d, *J* = 4.5 Hz, 1H), 6.30 (d, *J* = 6.2 Hz, 1H), 5.45 (d, *J* = 5.5 Hz, 1H), 4.80 (t, *J* = 5.6 Hz, 1H), 4.08 – 4.02 (m, 1H), 3.88 (q, *J* = 5.5 Hz, 1H), 3.57 (dd, *J* = 10.8, 4.6 Hz, 1H), 3.41 (dd, *J* = 10.8, 6.3 Hz, 1H).

**^13^C NMR** (126 MHz, DMSO) δ 156.1, 148.4, 123.7, 117.4, 117.1, 111.2, 101.3, 83.4, 79.5, 74.5, 74.2, 7.9.

### (2R,3R,4S,5R)-2-(4-aminopyrrolo[2,1-f][1,2,4]triazin-7-yl)-5-(azidomethyl)-3,4-dihydroxytetrahydrof uran-2-carbonitrile *(4)*

Compound **3** (55 mg, 0.137 mmol, 1 eq.) and sodium azide (18 mg, 0.274 mmol, 2 eq.) were combined and dissolved with 2 mL DMF. The reaction was stirred at 50 for 4 hours. Water (6 mL) was added, and the mixture was extracted with ethyl acetate (5 mL, three times). The combined organic phase was washed with brine, dried over anhydrous Na_2_SO_4_ and evaporated to yield product as a white solid (42 mg, yield: 97%)

**LC-MS (ESI)**: m/z calcd for C H N O ^+^ [M+H]^+^ 317.1, found 317.1.

**^1^H NMR** (500 MHz, DMSO-*d*_6_) δ 7.94 (s, 1H), 7.89 (br, 2H), 6.92 (d, *J* = 4.6 Hz, 1H), 6.84 (d, *J* = 4.5 Hz, 1H), 6.30 (d, *J* = 6.1 Hz, 1H), 5.40 (d, *J* = 5.6 Hz, 1H), 4.77 (t, *J* = 5.5 Hz, 1H), 4.18 (td, *J* = 6.1, 3.4 Hz, 1H), 3.94 (q, *J* = 5.6 Hz, 1H), 3.68 – 3.53 (m, 2H).

**^13^C NMR** (126 MHz, DMSO) δ 156.1, 148.4, 123.7, 117.5, 117.2, 110.9, 101.3, 83.0, 79.6, 74.3, 71.5, 51.8.

### (3aR,4R,6R,6aR)-4-(4-aminopyrrolo[2,1-f][1,2,4]triazin-7-yl)-6-(azidomethyl)-2,2-dimethyltetrahydrofuro[3,4-d][1,3]dioxole-4-carbonitrile *(5)*

Compound **4** (2.13 g, 6.73 mmol, 1 eq.) was dissolved in acetone (20 mL), followed by the sequential addition of 2,2-dimethoxypropane (3.32 mL, 26.92 mmol, 4 eq.) and concentrated H_2_SO_4_ (0.73 mL, 13.46 mmol, 2 eq.). The reaction was stirred at 45 for 1 hour and was allowed to cool to room temperature. NaHCO_3_ (2.26 g, 26.92 mmol, 4 eq.) was added to the reaction mixture which was then evaporated under vacuum to ∼1/5 of the original volume. Water (50 mL) and ethyl acetate (50 mL) were added, and the phases were separated. The aqueous phase was extracted with ethyl acetate (25 mL) for two additional times. The combine organic phase was washed with brine, dried and evaporated. Column chromatography purification (DCM: MeOH 60:1) afforded the product as a colorless syrup (2.05 g, yield: 96%).

**LC-MS (ESI)**: m/z calcd for C H N O ^+^ [M+H]^+^ 357.1, found 357.1.

**^1^H NMR** (400 MHz, Chloroform-*d*) δ 8.01 (s, 1H), 7.02 (d, *J* = 4.6 Hz, 1H), 6.63 (d, *J* = 4.7 Hz, 1H), 5.68 (s, 2H), 5.52 (d, *J* = 6.9 Hz, 1H), 4.96 (dd, *J* = 6.9, 4.4 Hz, 1H), 4.52 (q, *J* = 4.6 Hz, 1H), 3.68 (dd, *J* = 13.2, 4.4 Hz, 1H), 3.58 (dd, *J* = 13.2, 5.0 Hz, 1H), 1.76 (s, 3H), 1.41 (s, 3H).

### (3aR,4R,6R,6aR)-6-(aminomethyl)-4-(4-aminopyrrolo[2,1-f][1,2,4]triazin-7-yl)-2,2-dimethyltetrahydrofuro[3,4-d][1,3]dioxole-4-carbonitrile *(6)*

Azide **5** (1.7 g, 4.77 mmol, 1 eq.) and PPh_3_ (2.5 g, 9.54 mmol, 2 eq.) were combined and dissolved with THF (10 mL). The mixture was stirred at room temperature for 1 hour followed by the addition of water (10 mL). The resulting suspension was stirred overnight under reflux. The reaction mixture was evaporated under vacuum and the crude was purified on a silica gel column (DCM: MeOH 50:1 to 10:1 with 1% triethylamine) to afford the product as a white powder (1.21 g, yield: 77%)

**LC-MS (ESI)**: m/z calcd for C H N O ^+^ [M+H]^+^ 331.2, found 331.2.

**^1^H NMR** (500 MHz, Chloroform-*d*) δ 7.98 (s, 1H), 7.07 (d, *J* = 4.6 Hz, 1H), 6.62 (d, *J* = 4.7 Hz, 1H), 5.57 (d, *J* = 7.0 Hz, 1H), 5.42 (s, 2H), 5.05 (dd, *J* = 7.0, 4.0 Hz, 1H), 4.47 (q, *J* = 4.1 Hz, 1H), 3.18 (dd, *J* = 13.5, 3.8 Hz, 1H), 3.03 (dd, *J* = 13.5, 4.7 Hz, 1H), 1.77 (s, 3H), 1.41 (s, 3H).

### (9H-fluoren-9-yl)methyl

#### (2-((((3aR,4R,6R,6aR)-6-(4-aminopyrrolo[2,1-f][1,2,4]triazin-7-yl)-6-cyano-2,2-dimethyltetrahydrofur o[3,4-d][1,3]dioxol-4-yl)methyl)amino)-2-oxoethyl)carbamate *(10)*

**General procedure A** was followed. The product was obtained as a white solid (yield: 85%).

**LC-MS (ESI)**: m/z calcd for C H N O ^+^ [M+H]^+^ 610.2, found 610.4.

### 2-amino-N-(((3aR,4R,6R,6aR)-6-(4-aminopyrrolo[2,1-f][1,2,4]triazin-7-yl)-6-cyano-2,2-dimethyltetrahydrofuro[3,4-d][1,3]dioxol-4-yl)methyl)acetamide *(11)*

Compound **10** (1.37 g, 2.25 mmol, 1 eq.) was dissolved with DCM (20 mL). A 2 M solution of dimethylamine in THF (5.6 mL, 5 eq.) was added and the reaction mixture was allowed to stir at room temperature overnight. The solvent was evaporated and the mixture was purified by column chromatography (DCM:MeOH 50:1 to 10:1 with 1% triethylamine) to afford the product as a white solid (0.72 g, yield: 82%)

**LC-MS (ESI)**: m/z calcd for C H N O ^+^ [M+H]^+^ 388.2, found 388.2.

**^1^H NMR** (400 MHz, DMSO-*d*_6_) δ 8.02 (t, *J* = 5.8 Hz, 1H), 7.96 (s, 1H), 7.91 (br, 2H), 6.91 (d, *J* = 4.6 Hz, 1H), 6.89 (d, *J* = 4.6 Hz, 1H), 5.42 (d, *J* = 6.7 Hz, 1H), 4.84 (dd, *J* = 6.7, 3.4 Hz, 1H), 4.34 (td, *J* = 5.8, 3.3 Hz, 1H), 3.36 (td, *J* = 8.5, 5.1 Hz, 2H), 3.09 (s, 2H), 1.60 (s, 3H), 1.33 (s, 3H).

For target compounds **7** and **12a-12ab**, **11** and appropriate carboxylic acids were reacted following General Procedure A. The yields generally ranged from 70% to 90%.

### N-(2-((((3aR,4R,6R,6aR)-6-(4-aminopyrrolo[2,1-f][1,2,4]triazin-7-yl)-6-cyano-2,2-dimethyltetrahydrofuro[3,4-d][1,3]dioxol-4-yl)methyl)amino)-2-oxoethyl)-3-hydroxybenzamide *(7)*

**LC-MS (ESI)**: m/z calcd for C H N O ^+^ [M+H]^+^ 508.2, found 508.3.

**^1^H NMR** (400 MHz, DMSO-*d*_6_ + D_2_O) δ 7.97 (s, 1H), 7.32 – 7.21 (m, 3H), 6.95 – 6.87 (m, 3H), 5.41 (d, *J* = 6.6 Hz, 1H), 4.85 (dd, *J* = 6.6, 3.3 Hz, 1H), 4.37 (td, *J* = 5.8, 3.3 Hz, 1H), 3.83 (d, *J* = 2.1 Hz, 2H), 3.43 – 3.39 (m, 1H), 3.30 (dd, *J* = 14.0, 5.6 Hz, 1H), 1.62 (s, 3H), 1.36 (s, 3H).

**^13^C NMR** (101 MHz, DMSO) δ 170.0, 167.0, 157.6, 155.9, 148.6, 135.8, 129.8, 122.5, 118.7, 118.3, 117.4, 116.6, 116.1, 114.7, 111.3, 101.5, 84.2, 83.9, 82.5, 80.4, 43.0, 26.3, 25.6.

### N-(2-((((3aR,4R,6R,6aR)-6-(4-aminopyrrolo[2,1-f][1,2,4]triazin-7-yl)-6-cyano-2,2-dimethyltetrahydrofuro[3,4-d][1,3]dioxol-4-yl)methyl)amino)-2-oxoethyl)-5-hydroxynicotinamide *(12a)*

**LC-MS (ESI)**: m/z calcd for C H N O ^+^ [M+H]^+^ 509.2, found 509.2.

**^1^H NMR** (400 MHz, DMSO-*d*_6_ + D_2_O) δ 8.57 – 8.44 (m, 1H), 8.26 (d, *J* = 2.9 Hz, 1H), 7.97 (s, 1H), 7.64 – 7.49 (m, 1H), 6.90 (q, *J* = 4.6 Hz, 2H), 5.41 (d, *J* = 6.6 Hz, 1H), 4.85 (dd, *J* = 6.6, 3.3 Hz, 1H), 4.37 (td, *J* = 5.6, 3.1 Hz, 1H), 3.92 – 3.80 (m, 2H), 3.41 (d, *J* = 5.4 Hz, 1H), 3.31 (dd, *J* = 14.0, 5.7 Hz, 1H), 1.62 (s, 3H), 1.36 (s, 3H).

**^13^C NMR** (101 MHz, DMSO) δ 169.8, 165.7, 155.9, 153.7, 148.6, 140.8, 139.6, 130.7, 122.5, 121.3, 117.4, 116.6, 116.2, 111.3, 101.5, 84.2, 83.8, 82.5, 80.4, 43.0, 26.3, 25.6.

### N-(2-((((3aR,4R,6R,6aR)-6-(4-aminopyrrolo[2,1-f][1,2,4]triazin-7-yl)-6-cyano-2,2-dimethyltetrahydrofuro[3,4-d][1,3]dioxol-4-yl)methyl)amino)-2-oxoethyl)-5-methoxynicotinamide *(12b)*

**LC-MS (ESI)**: m/z calcd for C H N O ^+^ [M+H]^+^ 523.2, found 523.2.

**^1^H NMR** (500 MHz, DMSO-*d*_6_ + D_2_O) δ 8.63 (d, *J* = 1.7 Hz, 1H), 8.43 (d, *J* = 2.9 Hz, 1H), 7.97 (s, 1H), 7.75 (dd, *J* = 2.9, 1.8 Hz, 1H), 6.90 (q, *J* = 4.6 Hz, 2H), 5.40 (d, *J* = 6.6 Hz, 1H), 4.85 (dd, *J* = 6.6, 3.3 Hz, 1H), 4.37 (td, *J* = 5.6, 3.3 Hz, 1H), 3.89 (d, *J* = 2.2 Hz, 2H), 3.87 (s, 3H), 3.42 (d, *J* = 5.7 Hz, 1H), 3.31 (dd, *J* = 13.9, 5.6 Hz, 1H), 1.62 (s, 3H), 1.36 (s, 3H).

**^13^C NMR** (126 MHz, DMSO) δ 169.7, 165.5, 155.9, 155.6, 148.7, 141.0, 140.7, 130.4, 122.5, 119.1, 117.4, 116.6, 116.2, 111.3, 101.4, 84.2, 83.8, 82.5, 80.4, 56.2, 43.0, 26.3, 25.5.

#### N-(2-((((3aR,4R,6R,6aR)-6-(4-aminopyrrolo[2,1-f][1,2,4]triazin-7-yl)-6-cyano-2,2-dimethyltetrahydrofuro[3,4-d][1,3]dioxol-4-yl)methyl)amino)-2-oxoethyl)-5-nitronicotinamide *(12c)*

**LC-MS (ESI)**: m/z calcd for C H N O ^+^ [M+H]^+^ 538.2, found 538.2.

**^1^H NMR** (500 MHz, DMSO-*d*_6_ + D_2_O) δ 9.51 (d, *J* = 2.5 Hz, 1H), 9.36 (d, *J* = 1.9 Hz, 1H), 8.99 – 8.94 (m, 1H), 7.96 (s, 1H), 6.91 – 6.86 (m, 2H), 5.41 (d, *J* = 6.7 Hz, 1H), 4.86 (dd, *J* = 6.7, 3.4 Hz, 1H), 4.37 (td, *J* = 5.6, 3.4 Hz, 1H), 3.94 (s, 2H), 3.42 (d, *J* = 5.8 Hz, 1H), 3.34 (dd, *J* = 14.1, 5.6 Hz, 1H), 1.62 (s, 3H), 1.36 (s, 3H).

**^13^C NMR** (126 MHz, DMSO) δ 169.4, 163.7, 155.9, 154.1, 148.6, 147.2, 144.5, 130.6, 130.3, 122.5, 117.4, 116.2, 111.2, 101.4, 84.2, 83.8, 82.5, 80.4, 43.1, 25.6.

#### N-(2-((((3aR,4R,6R,6aR)-6-(4-aminopyrrolo[2,1-f][1,2,4]triazin-7-yl)-6-cyano-2,2-dimethyltetrahydrofuro[3,4-d][1,3]dioxol-4-yl)methyl)amino)-2-oxoethyl)-3-(methylsulfonamido)benzamide *(12d)*

**LC-MS (ESI)**: m/z calcd for C_25_H_29_N_8_O_7_S^+^ [M+H]^+^ 585.2, found 585.2.

**^1^H NMR** (400 MHz, DMSO-*d*_6_ + D_2_O) δ 7.97 (s, 1H), 7.69 (t, *J* = 2.0 Hz, 1H), 7.60 (dt, *J* = 7.7, 1.4 Hz, 1H), 7.43 (t, *J* = 7.9 Hz, 1H), 7.36 (ddd, *J* = 8.1, 2.3, 1.1 Hz, 1H), 6.91 (q, *J* = 4.6 Hz, 2H), 5.41 (d, *J* = Hz, 1H), 4.85 (dd, *J* = 6.6, 3.3 Hz, 1H), 4.37 (td, *J* = 5.6, 3.2 Hz, 1H), 3.94 – 3.79 (m, 2H), 3.46 – 3.39 (m, 1H), 3.31 (ddd, *J* = 12.9, 5.9, 4.3 Hz, 1H), 3.00 (s, 3H), 1.62 (s, 3H), 1.36 (s, 3H).

**^13^C NMR** (101 MHz, DMSO) δ 169.9, 166.7, 155.9, 148.7, 138.9, 135.6, 129.7, 123.2, 123.0, 122.5, 119.4, 117.4, 116.6, 116.2, 111.3, 101.5, 84.2, 83.9, 82.5, 80.4, 43.1, 39.8, 26.3, 25.6.

#### N-(2-((((3aR,4R,6R,6aR)-6-(4-aminopyrrolo[2,1-f][1,2,4]triazin-7-yl)-6-cyano-2,2-dimethyltetrahydrofuro[3,4-d][1,3]dioxol-4-yl)methyl)amino)-2-oxoethyl)-4-(methylsulfonamido)benzamide *(12e)*

**LC-MS (ESI)**: m/z calcd for C_25_H_29_N_8_O_7_S^+^ [M+H]^+^ 585.2, found 585.2.

**^1^H NMR** (500 MHz, DMSO-*d*_6_ + D_2_O) δ 7.97 (s, 1H), 7.84 (d, *J* = 8.7 Hz, 2H), 7.25 (d, *J* = 8.7 Hz, 2H), 6.92 (d, *J* = 4.6 Hz, 1H), 6.90 (d, *J* = 4.6 Hz, 1H), 5.41 (d, *J* = 6.6 Hz, 1H), 4.88 – 4.82 (m, 1H), 4.36 (qd, *J* = 6.0, 3.3 Hz, 1H), 3.90 – 3.80 (m, 2H), 3.41 (d, *J* = 5.6 Hz, 1H), 3.30 (ddd, *J* = 13.8, 5.8, 4.4 Hz, 1H), 3.05 (s, 3H), 1.62 (s, 3H), 1.35 (s, 3H).

**^13^C NMR** (126 MHz, DMSO) δ 170.1, 166.4, 155.9, 148.7, 141.7, 129.2, 129.1, 122.5, 118.4, 117.4, 116.6, 116.1, 111.3, 101.4, 84.2, 83.8, 82.5, 80.4, 40.3, 26.3, 25.6.

#### N-(2-((((3aR,4R,6R,6aR)-6-(4-aminopyrrolo[2,1-f][1,2,4]triazin-7-yl)-6-cyano-2,2-dimethyltetrahydrofuro[3,4-d][1,3]dioxol-4-yl)methyl)amino)-2-oxoethyl)-5-(methylsulfonamido)nicotinamide *(12f)*

**LC-MS (ESI)**: m/z calcd for C_24_H_28_N_9_O_7_S^+^ [M+H]^+^ 586.2, found 586.2.

**^1^H NMR** (500 MHz, DMSO-*d*_6_ + D_2_O) δ 8.76 (d, *J* = 1.9 Hz, 1H), 8.53 (d, *J* = 2.6 Hz, 1H), 8.00 (t, *J* = 2.3 Hz, 1H), 7.97 (s, 1H), 6.91 (d, *J* = 4.6 Hz, 1H), 6.90 (d, *J* = 4.6 Hz, 1H), 5.42 (d, *J* = 6.6 Hz, 1H), 4.85 (dd, *J* = 6.7, 3.3 Hz, 1H), 4.37 (td, *J* = 5.7, 3.2 Hz, 1H), 3.95 – 3.81 (m, 2H), 3.42 (d, *J* = 5.8 Hz, 1H), 3.31 (dd, *J* = 14.1, 5.7 Hz, 1H), 3.08 (s, 3H), 1.62 (s, 3H), 1.35 (s, 3H).

**^13^C NMR** (126 MHz, DMSO) δ 169.7, 168.5, 165.2, 155.9, 148.7, 144.1, 143.7, 130.4, 126.0, 122.5, 117.4, 116.2, 111.3, 101.4, 88.3, 84.2, 83.8, 82.5, 80.4, 43.0, 26.3, 25.6.

#### N-(2-((((3aR,4R,6R,6aR)-6-(4-aminopyrrolo[2,1-f][1,2,4]triazin-7-yl)-6-cyano-2,2-dimethyltetrahydrofuro[3,4-d][1,3]dioxol-4-yl)methyl)amino)-2-oxoethyl)-6-(methylsulfonamido)nicotinamide *(12g)*

**LC-MS (ESI)**: m/z calcd for C_24_H_28_N_9_O_7_S^+^ [M+H]^+^ 586.2, found 586.3.

**^1^H NMR** (400 MHz, DMSO-*d*_6_ + D_2_O) δ 8.66 (d, *J* = 2.4 Hz, 1H), 8.10 (dd, *J* = 8.8, 2.4 Hz, 1H), 7.95 (s, 1H), 6.99 (d, *J* = 8.7 Hz, 1H), 6.89 (q, *J* = 4.6 Hz, 2H), 5.39 (d, *J* = 6.6 Hz, 1H), 4.84 (dd, *J* = 6.7, 3.3 Hz, 1H), 4.35 (td, *J* = 5.7, 3.3 Hz, 1H), 3.85 (d, *J* = 1.7 Hz, 2H), 3.41 – 3.36 (m, 1H), 3.29 (dd, *J* = 14.0, 5.7 Hz, 1H), 3.25 (s, 3H), 1.60 (s, 3H), 1.34 (s, 3H).

**^13^C NMR** (101 MHz, DMSO) δ 169.9, 165.1, 155.9, 155.3, 148.6, 138.2, 122.5, 117.4, 116.6, 116.1, 112.0, 111.3, 101.5, 84.2, 83.8, 82.5, 80.4, 42.9, 42.0, 26.3, 25.6.

### 3-acetamido-N-(2-((((3aR,4R,6R,6aR)-6-(4-aminopyrrolo[2,1-f][1,2,4]triazin-7-yl)-6-cyano-2,2-dimethyltetrahydrofuro[3,4-d][1,3]dioxol-4-yl)methyl)amino)-2-oxoethyl)benzamide *(12h)*

**LC-MS (ESI)**: m/z calcd for C_26_H_29_N_8_O_6_S^+^ [M+H]^+^ 549.2, found 549.3.

**^1^H NMR** (500 MHz, DMSO-*d*_6_ + D_2_O) δ 8.64 (t, *J* = 5.9 Hz, 1H), 8.09 (t, *J* = 5.9 Hz, 1H), 7.96 (q, *J* = 2.0 Hz, 1H), 7.91 (s, 1H), 7.72 – 7.65 (m, 1H), 7.46 (dt, *J* = 7.9, 1.3 Hz, 1H), 7.32 (t, *J* = 7.9 Hz, 1H), 6.84 (q, *J* = 4.6 Hz, 2H), 5.35 (d, *J* = 6.7 Hz, 1H), 4.79 (dd, *J* = 6.6, 3.3 Hz, 1H), 4.30 (td, *J* = 5.7, 3.2 Hz, 1H), 3.89 – 3.73 (m, 2H), 3.39 – 3.32 (m, 1H), 3.24 (dt, *J* = 13.8, 5.5 Hz, 1H), 1.99 (s, 3H), 1.56 (s, 3H), 1.29 (s, 3H).

**^13^C NMR** (126 MHz, DMSO) δ 170.1, 169.0, 167.0, 155.9, 148.7, 139.7, 135.0, 129.1, 122.5, 122.3, 122.1, 118.8, 117.4, 116.6, 116.2, 111.3, 101.5, 84.2, 83.9, 82.5, 80.4, 43.2, 26.3, 25.6, 24.4.

#### N-(2-((((3aR,4R,6R,6aR)-6-(4-aminopyrrolo[2,1-f][1,2,4]triazin-7-yl)-6-cyano-2,2-dimethyltetrahydrofuro[3,4-d][1,3]dioxol-4-yl)methyl)amino)-2-oxoethyl)-2-oxoindoline-6-carboxamide *(12i)*

**LC-MS (ESI)**: m/z calcd for C_26_H_27_N_8_O_6_S^+^ [M+H]^+^ 547.2, found 547.3.

**^1^H NMR** (500 MHz, DMSO-*d*_6_ + D_2_O) δ 8.73 (t, *J* = 5.8 Hz, 1H), 8.13 (t, *J* = 5.9 Hz, 1H), 7.97 (s, 1H), 7.46 (dd, *J* = 7.6, 1.7 Hz, 1H), 7.31 – 7.25 (m, 2H), 6.92 – 6.86 (m, 2H), 5.40 (d, *J* = 6.7 Hz, 1H), 4.84 (dd, *J* = 6.6, 3.3 Hz, 1H), 4.37 (td, *J* = 5.5, 3.2 Hz, 1H), 3.84 (dd, *J* = 5.9, 1.7 Hz, 2H), 3.55 – 3.49 (m, 2H), 3.46 – 3.41 (m, 1H), 3.29 (dt, *J* = 13.8, 5.5 Hz, 1H), 1.62 (s, 3H), 1.35 (s, 3H).

**^13^C NMR** (126 MHz, DMSO) δ 176.9, 170.1, 169.6, 167.0, 160.5, 155.9, 148.7, 144.2, 129.8, 124.6, 122.5, 121.0, 117.4, 116.2, 111.3, 108.4, 101.4, 93.1, 84.2, 83.8, 82.4, 80.4, 43.2, 40.7, 26.3, 25.6.

#### N-(2-((((3aR,4R,6R,6aR)-6-(4-aminopyrrolo[2,1-f][1,2,4]triazin-7-yl)-6-cyano-2,2-dimethyltetrahydrofuro[3,4-d][1,3]dioxol-4-yl)methyl)amino)-2-oxoethyl)-3-(phenylsulfonamido)benzamide *(12j)*

**LC-MS (ESI)**: m/z calcd for C_30_H_31_N_8_O_7_S^+^ [M+H]^+^ 647.2, found 647.3.

**^1^H NMR** (500 MHz, DMSO-*d*_6_ + D_2_O) δ 7.97 (s, 1H), 7.79 – 7.73 (m, 2H), 7.63 – 7.57 (m, 2H), 7.56 –7.48 (m, 3H), 7.31 (t, *J* = 7.9 Hz, 1H), 7.24 (ddd, *J* = 8.0, 2.3, 1.1 Hz, 1H), 6.92 (d, *J* = 4.6 Hz, 1H), 6.89 (d, *J* = 4.6 Hz, 1H), 5.41 (d, *J* = 6.7 Hz, 1H), 4.84 (dd, *J* = 6.7, 3.3 Hz, 1H), 4.35 (td, *J* = 5.7, 3.3 Hz, 1H), 3.89 – 3.72 (m, 2H), 3.41 (dd, *J* = 14.0, 5.7 Hz, 1H), 3.29 (dd, *J* = 14.0, 5.7 Hz, 1H), 1.62 (s, 3H), 1.34 (s, 3H).

**^13^C NMR** (126 MHz, DMSO) δ 169.9, 166.4, 155.9, 148.7, 135.4, 133.4, 129.8, 129.6, 127.1, 123.1, 123.0, 122.5, 119.9, 117.4, 116.6, 116.1, 111.3, 101.5, 84.2, 83.8, 82.5, 80.4, 43.0, 26.3, 25.6.

#### N-(2-((((3aR,4R,6R,6aR)-6-(4-aminopyrrolo[2,1-f][1,2,4]triazin-7-yl)-6-cyano-2,2-dimethyltetrahydrofuro[3,4-d][1,3]dioxol-4-yl)methyl)amino)-2-oxoethyl)-3-((phenylmethyl)sulfonamido)benzamide *(12k)*

**LC-MS (ESI)**: m/z calcd for C_31_H_33_N_8_O_7_S^+^ [M+H]^+^ 661.2, found 661.3.

**^1^H NMR** (500 MHz, DMSO-*d*_6_ + D_2_O) δ 7.97 (s, 1H), 7.69 (t, *J* = 2.0 Hz, 1H), 7.58 – 7.53 (m, 1H), 7.40 (t, *J* = 7.9 Hz, 1H), 7.37 – 7.32 (m, 3H), 7.30 (dd, *J* = 8.8, 2.3 Hz, 1H), 7.25 (dd, *J* = 6.6, 3.0 Hz, 2H), 6.92 (d, *J* = 4.6 Hz, 1H), 6.90 (d, *J* = 4.6 Hz, 1H), 5.42 (d, *J* = 6.6 Hz, 1H), 4.86 (dd, *J* = 6.7, 3.3 Hz, 1H), 4.48 (s, 2H), 4.38 (td, *J* = 5.6, 3.3 Hz, 1H), 3.95 – 3.80 (m, 2H), 3.31 (dd, *J* = 14.0, 5.7 Hz, 1H), 1.62 (s, 3H), 1.36 (s, 3H).

**^13^C NMR** (126 MHz, DMSO) δ 169.9, 166.8, 155.9, 148.7, 135.6, 131.4, 129.7, 128.9, 128.7, 122.5, 122.3, 118.5, 117.4, 116.6, 116.2, 111.3, 101.5, 84.2, 83.9, 82.5, 80.4, 57.4, 43.0, 26.3, 25.6.

#### N-(2-((((3aR,4R,6R,6aR)-6-(4-aminopyrrolo[2,1-f][1,2,4]triazin-7-yl)-6-cyano-2,2-dimethyltetrahydrofuro[3,4-d][1,3]dioxol-4-yl)methyl)amino)-2-oxoethyl)-5-(phenylsulfonamido)nicotinamide *(12l)*

**LC-MS (ESI)**: m/z calcd for C_29_H_30_N_9_O_7_S^+^ [M+H]^+^ 648.2, found 648.3.

**^1^H NMR** (500 MHz, DMSO-*d*_6_ + D_2_O) δ 8.65 (d, *J* = 2.0 Hz, 1H), 8.34 (d, *J* = 2.6 Hz, 1H), 7.91 (s, 1H), 7.86 (t, *J* = 2.3 Hz, 1H), 7.75 – 7.67 (m, 2H), 7.59 – 7.53 (m, 1H), 7.50 (dd, *J* = 8.3, 6.8 Hz, 2H), 6.86 (d, *J* = 4.6 Hz, 1H), 6.83 (d, *J* = 4.6 Hz, 1H), 5.35 (d, *J* = 6.7 Hz, 1H), 4.78 (dd, *J* = 6.6, 3.3 Hz, 1H), 4.29 (td, *J* = 5.8, 3.3 Hz, 1H), 3.85 – 3.72 (m, 2H), 3.34 (dd, *J* = 14.1, 5.8 Hz, 1H), 3.24 (dd, *J* = 13.9, 5.7 Hz, 1H), 1.55 (s, 3H), 1.28 (s, 3H).

**^13^C NMR** (126 MHz, DMSO) δ 169.6, 164.9, 155.9, 148.7, 144.2, 144.1, 139.4, 135.0, 133.8, 130.3, 130.0, 127.1, 126.8, 124.6, 122.5, 119.2, 117.4, 116.6, 116.2, 111.3, 110.6, 101.5, 84.1, 83.8, 82.5, 80.4, 53.9, 42.2, 26.3, 25.5.

#### N-(2-((((3aR,4R,6R,6aR)-6-(4-aminopyrrolo[2,1-f][1,2,4]triazin-7-yl)-6-cyano-2,2-dimethyltetrahydrofuro[3,4-d][1,3]dioxol-4-yl)methyl)amino)-2-oxoethyl)-5-((3-methoxyphenyl)sulfonamido)nicotinamide *(12m)*

**LC-MS (ESI)**: m/z calcd for C_30_H_32_N_9_O_8_S^+^ [M+H]^+^ 678.2, found 678.3.

**^1^H NMR** (500 MHz, DMSO-*d*_6_ + D_2_O) δ 8.72 (d, *J* = 2.0 Hz, 1H), 8.41 (d, *J* = 2.6 Hz, 1H), 7.97 (s, 1H), 7.94 (t, *J* = 2.3 Hz, 1H), 7.47 (t, *J* = 8.0 Hz, 1H), 7.32 (dd, *J* = 7.3, 1.8 Hz, 1H), 7.27 (t, *J* = 2.2 Hz, 1H), 7.19 (dd, *J* = 8.1, 2.6 Hz, 1H), 6.92 (d, *J* = 4.7 Hz, 1H), 6.89 (d, *J* = 4.7 Hz, 1H), 5.41 (d, *J* = 6.7 Hz, 1H), 4.84 (dd, *J* = 6.7, 3.3 Hz, 1H), 4.35 (td, *J* = 5.8, 3.2 Hz, 1H), 3.90 – 3.80 (m, 2H), 3.77 (s, 3H), 3.39 (d, *J* = 5.5 Hz, 1H), 3.30 (dd, *J* = 14.1, 5.5 Hz, 1H), 1.62 (s, 3H), 1.34 (s, 3H).

**^13^C NMR** (126 MHz, DMSO) δ 164.9, 159.9, 155.9, 148.6, 144.2, 131.2, 130.3, 126.8, 122.5, 119.6, 119.2, 117.4, 116.2, 112.0, 111.3, 101.4, 84.1, 82.5, 80.4, 56.1, 26.3, 25.5.

#### N-(2-((((3aR,4R,6R,6aR)-6-(4-aminopyrrolo[2,1-f][1,2,4]triazin-7-yl)-6-cyano-2,2-dimethyltetrahydrofuro[3,4-d][1,3]dioxol-4-yl)methyl)amino)-2-oxoethyl)-5-((3-nitrophenyl)sulfonamido)nicotinamide *(12n)*

**LC-MS (ESI)**: m/z calcd for C_29_H_29_N_10_O_9_S^+^ [M+H]^+^ 693.2, found 693.3.

**^1^H NMR** (500 MHz, DMSO-*d*_6_ + D_2_O) δ 8.77 – 8.68 (m, 1H), 8.51 (t, *J* = 2.0 Hz, 1H), 8.47 – 8.42 (m, 1H), 8.41 (d, *J* = 2.6 Hz, 1H), 8.14 (dt, *J* = 7.9, 1.4 Hz, 1H), 7.96 (s, 1H), 7.91 (t, *J* = 2.3 Hz, 1H), 7.84 (t, *J* = 8.1 Hz, 1H), 6.91 (d, *J* = 4.5 Hz, 1H), 6.88 (d, *J* = 4.6 Hz, 1H), 5.40 (d, *J* = 6.6 Hz, 1H), 4.84 (dd, *J* = 6.7, 3.3 Hz, 1H), 4.35 (td, *J* = 5.7, 3.3 Hz, 1H), 3.85 (t, *J* = 3.7 Hz, 2H), 3.44 – 3.38 (m, 1H), 3.30 (dt, *J* = 13.9, 5.5 Hz, 1H), 1.61 (s, 3H), 1.34 (s, 3H).

**^13^C NMR** (126 MHz, DMSO) δ 169.7, 155.9, 148.6, 148.4, 132.9, 132.0, 122.5, 121.8, 117.4, 116.6, 116.2, 111.3, 101.5, 84.1, 83.8, 82.5, 80.4, 54.0, 26.3, 25.5.

#### N-(2-((((3aR,4R,6R,6aR)-6-(4-aminopyrrolo[2,1-f][1,2,4]triazin-7-yl)-6-cyano-2,2-dimethyltetrahydrofuro[3,4-d][1,3]dioxol-4-yl)methyl)amino)-2-oxoethyl)-5-((3-fluorophenyl)sulfonamido)nicotinamide *(12o)*

**LC-MS (ESI)**: m/z calcd for C_29_H_29_FN_9_O_7_S^+^ [M+H]^+^ 666.2, found 666.3.

**^1^H NMR** (500 MHz, DMSO-*d*_6_ + D_2_O) δ 8.70 (d, *J* = 1.9 Hz, 1H), 8.39 (d, *J* = 2.6 Hz, 1H), 7.97 (s, 1H), 7.90 (t, *J* = 2.2 Hz, 1H), 7.67 – 7.54 (m, 3H), 7.49 (ddt, *J* = 10.7, 8.0, 2.0 Hz, 1H), 6.91 (d, *J* = 4.6 Hz, 1H), 6.89 (d, *J* = 4.6 Hz, 1H), 5.41 (d, *J* = 6.7 Hz, 1H), 4.84 (dd, *J* = 6.6, 3.3 Hz, 1H), 4.36 (td, *J* = 5.7, 3.3 Hz, 1H), 3.91 – 3.77 (m, 2H), 3.40 (d, *J* = 5.8 Hz, 1H), 3.30 (dd, *J* = 13.8, 5.7 Hz, 1H), 1.62 (s, 3H), 1.34 (s, 3H).

**^13^C NMR** (126 MHz, DMSO) δ 169.7, 169.6, 165.0, 163.1, 161.2, 155.9, 148.6, 144.6, 132.4, 132.3, 130.3, 127.0, 123.3, 122.5, 117.4, 116.6, 116.1, 114.2, 114.0, 111.3, 101.4, 84.1, 83.8, 82.5, 80.4, 46.2, 43.0, 40.3, 40.2, 40.0, 39.8, 39.7, 39.5, 39.3, 26.3, 25.5.

#### N-(2-((((3aR,4R,6R,6aR)-6-(4-aminopyrrolo[2,1-f][1,2,4]triazin-7-yl)-6-cyano-2,2-dimethyltetrahydrofuro[3,4-d][1,3]dioxol-4-yl)methyl)amino)-2-oxoethyl)-5-((4-fluorophenyl)sulfonamido)nicotinamide *(12p)*

**LC-MS (ESI)**: m/z calcd for C_29_H_29_FN_9_O_7_S^+^ [M+H]^+^ 666.2, found 666.3.

**^1^H NMR** (500 MHz, DMSO-*d*_6_ + D_2_O) δ 8.60 (d, *J* = 1.9 Hz, 1H), 8.33 (d, *J* = 2.6 Hz, 1H), 7.97 (s, 1H), 7.82 (ddt, *J* = 12.2, 5.4, 2.8 Hz, 3H), 7.39 – 7.31 (m, 2H), 6.92 (d, *J* = 4.6 Hz, 1H), 6.89 (d, *J* = 4.6 Hz, 1H), 5.41 (d, *J* = 6.7 Hz, 1H), 4.84 (dd, *J* = 6.6, 3.3 Hz, 1H), 4.36 (td, *J* = 5.7, 3.3 Hz, 1H), 3.89 – 3.80 (m, 2H), 3.42 – 3.38 (m, 1H), 3.30 (dd, *J* = 14.0, 5.8 Hz, 1H), 1.62 (s, 3H), 1.34 (s, 3H).

**^13^C NMR** (126 MHz, DMSO) δ 169.7, 165.3, 163.5, 155.9, 148.7, 144.7, 143.3, 130.1, 130.0, 129.9, 126.5, 123.7, 122.5, 118.8, 117.4, 116.9, 116.7, 116.6, 116.1, 111.3, 111.3, 101.4, 84.2, 83.8, 82.5, 80.4, 46.1, 40.3, 40.2, 40.0, 39.8, 39.7, 39.5, 39.3, 26.3, 25.6.

#### N-(2-((((3aR,4R,6R,6aR)-6-(4-aminopyrrolo[2,1-f][1,2,4]triazin-7-yl)-6-cyano-2,2-dimethyltetrahydrofuro[3,4-d][1,3]dioxol-4-yl)methyl)amino)-2-oxoethyl)-5-((3,4,5-trifluorophenyl)sulfonamido)nicotinamide *(12q)*

**LC-MS (ESI)**: m/z calcd for C_29_H_27_F_3_N_9_O_7_S^+^ [M+H]^+^ 702.2, found 702.4.

**^1^H NMR** (400 MHz, DMSO-*d*_6_ + D_2_O) δ 8.75 (d, *J* = 1.9 Hz, 1H), 8.43 (d, *J* = 2.6 Hz, 1H), 8.19 (t, *J* = 5.9 Hz, 1H), 7.97 (s, 1H), 7.92 (t, *J* = 2.2 Hz, 1H), 7.73 (t, *J* = 6.5 Hz, 2H), 6.91 (d, *J* = 4.5 Hz, 1H), 6.87 (d, *J* = 4.6 Hz, 1H), 5.41 (d, *J* = 6.6 Hz, 1H), 4.84 (dd, *J* = 6.7, 3.3 Hz, 1H), 4.36 (td, *J* = 5.7, 3.2 Hz, 1H), 3.86 (q, *J* = 2.8, 2.0 Hz, 2H), 3.42 – 3.25 (m, 2H), 1.62 (s, 3H), 1.34 (s, 3H).

**^13^C NMR** (101 MHz, DMSO) δ 169.7, 165.1, 165.0, 155.9, 149.4, 148.6, 145.0, 144.4, 130.3, 127.4, 122.5, 117.4, 116.6, 116.1, 112.9, 112.8, 112.6, 111.3, 101.4, 84.1, 83.8, 82.5, 80.4, 46.1, 43.0, 26.3, 25.5.

#### N-(2-((((3aR,4R,6R,6aR)-6-(4-aminopyrrolo[2,1-f][1,2,4]triazin-7-yl)-6-cyano-2,2-dimethyltetrahydrofuro[3,4-d][1,3]dioxol-4-yl)methyl)amino)-2-oxoethyl)-5-((4-(trifluoromethyl)phenyl)sulfonamido)nicotinamide *(12r)*

**LC-MS (ESI)**: m/z calcd for C_30_H_29_F_3_N_9_O_7_S^+^ [M+H]^+^ 716.2, found 716.4.

**^1^H NMR** (400 MHz, DMSO-*d*_6_ + D_2_O) δ 9.04 (t, *J* = 5.8 Hz, 1H), 8.76 (d, *J* = 1.9 Hz, 1H), 8.42 (d, *J* = 2.5 Hz, 1H), 8.19 (t, *J* = 5.9 Hz, 1H), 8.05 – 7.86 (m, 6H), 6.91 (d, *J* = 4.6 Hz, 1H), 6.88 (d, *J* = 4.6 Hz, 1H), 5.41 (d, *J* = 6.6 Hz, 1H), 4.84 (dd, *J* = 6.6, 3.3 Hz, 1H), 4.36 (td, *J* = 5.7, 3.2 Hz, 1H), 3.95 – 3.79 (m, 2H), 3.44 – 3.26 (m, 2H), 1.61 (s, 3H), 1.34 (s, 3H).

**^13^C NMR** (101 MHz, DMSO) δ 169.7, 169.6, 164.8, 155.9, 148.6, 144.6, 144.6, 143.4, 134.5, 133.5, 133.2, 130.4, 128.1, 127.8, 127.3, 127.2, 125.1, 122.5, 122.4, 117.4, 116.6, 116.2, 111.3, 101.4, 84.1, 83.8, 82.5, 80.4, 43.0, 26.3, 25.5.

#### N-(2-((((3aR,4R,6R,6aR)-6-(4-aminopyrrolo[2,1-f][1,2,4]triazin-7-yl)-6-cyano-2,2-dimethyltetrahydrofuro[3,4-d][1,3]dioxol-4-yl)methyl)amino)-2-oxoethyl)-5-((3-(trifluoromethyl)phenyl)sulfonamido)nicotinamide *(12s)*

**LC-MS (ESI)**: m/z calcd for C_30_H_29_F_3_N_9_O_7_S^+^ [M+H]^+^ 716.2, found 716.4.

**^1^H NMR** (400 MHz, DMSO-*d*_6_ + D_2_O) δ 9.01 (t, *J* = 5.9 Hz, 1H), 8.71 (d, *J* = 1.9 Hz, 1H), 8.38 (d, *J* = 2.6 Hz, 1H), 8.18 (t, *J* = 5.9 Hz, 1H), 8.10 – 7.99 (m, 3H), 7.97 (s, 1H), 7.90 (t, *J* = 2.2 Hz, 1H), 7.80 (t, *J* = 7.8 Hz, 1H), 6.92 (d, *J* = 4.6 Hz, 1H), 6.89 (d, *J* = 4.6 Hz, 1H), 5.41 (d, *J* = 6.7 Hz, 1H), 4.84 (dd, *J* = 6.7, 3.3 Hz, 1H), 4.36 (td, *J* = 5.7, 3.2 Hz, 1H), 3.92 – 3.79 (m, 2H), 3.41 – 3.25 (m, 2H), 1.61 (s, 3H), 1.34 (s, 3H).

**^13^C NMR** (101 MHz, DMSO) δ 169.7, 169.6, 165.0, 155.9, 148.6, 144.8, 144.0, 131.5, 131.1, 130.6, 130.3, 130.2, 127.3, 125.1, 124.9, 123.6, 122.5, 119.4, 117.4, 116.6, 116.1, 111.3, 110.3, 101.5, 84.1, 83.8, 82.5, 80.4, 70.2, 46.1, 43.0, 26.3, 25.5.

#### N-(2-((((3aR,4R,6R,6aR)-6-(4-aminopyrrolo[2,1-f][1,2,4]triazin-7-yl)-6-cyano-2,2-dimethyltetrahydrofuro[3,4-d][1,3]dioxol-4-yl)methyl)amino)-2-oxoethyl)-5-(pyridine-3-sulfonamido)nicotinamide *(12t)*

**LC-MS (ESI)**: m/z calcd for C_28_H_29_N_10_O_7_S^+^ [M+H]^+^ 649.2, found 649.3.

**^1^H NMR** (500 MHz, DMSO-*d*_6_ + D_2_O) δ 8.90 (d, *J* = 2.4 Hz, 1H), 8.77 (dd, *J* = 4.9, 1.6 Hz, 1H), 8.73 –8.66 (m, 1H), 8.39 (d, *J* = 2.6 Hz, 1H), 8.13 (dt, *J* = 8.1, 1.9 Hz, 1H), 7.97 (s, 1H), 7.90 (t, *J* = 2.3 Hz, 1H), 7.59 (dd, *J* = 8.1, 4.8 Hz, 1H), 6.92 (d, *J* = 4.6 Hz, 1H), 6.89 (d, *J* = 4.6 Hz, 1H), 5.41 (d, *J* = 6.6 Hz, 1H), 4.84 (dd, *J* = 6.7, 3.3 Hz, 1H), 4.36 (td, *J* = 5.7, 3.2 Hz, 1H), 3.90 – 3.80 (m, 2H), 3.39 (dd, *J* = 5.8, 1.7 Hz, 1H), 3.34 – 3.27 (m, 1H), 1.62 (s, 3H), 1.34 (s, 3H).

**^13^C NMR** (126 MHz, DMSO) δ 169.7, 168.1, 165.0, 155.9, 148.6, 147.4, 144.8, 135.1, 130.3, 127.1, 124.9, 122.5, 117.4, 116.1, 111.3, 101.5, 84.1, 83.8, 82.5, 80.4, 26.3, 25.5.

#### N-(2-((((3aR,4R,6R,6aR)-6-(4-aminopyrrolo[2,1-f][1,2,4]triazin-7-yl)-6-cyano-2,2-dimethyltetrahydrofuro[3,4-d][1,3]dioxol-4-yl)methyl)amino)-2-oxoethyl)-5-((6-(trifluoromethyl)pyridine)-3-sulfonamido)nicotinamide (12u)

**LC-MS (ESI)**: m/z calcd for C_29_H_28_F_3_N_10_O_7_S^+^ [M+H]^+^ 717.2, found 717.4.

**^1^H NMR** (400 MHz, DMSO-*d*_6_ + D_2_O) δ 9.08 (d, *J* = 2.2 Hz, 1H), 8.99 (t, *J* = 5.9 Hz, 1H), 8.68 (d, *J* = 1.9 Hz, 1H), 8.46 – 8.34 (m, 2H), 8.19 (t, *J* = 5.9 Hz, 1H), 8.08 (d, *J* = 8.3 Hz, 1H), 7.97 (s, 1H), 7.91 (t, *J* = 2.2 Hz, 1H), 6.92 (d, *J* = 4.6 Hz, 1H), 6.88 (d, *J* = 4.6 Hz, 1H), 5.41 (d, *J* = 6.6 Hz, 1H), 4.85 (dd, *J* = 6.6, 3.3 Hz, 1H), 4.36 (td, *J* = 5.7, 3.2 Hz, 1H), 3.86 (dd, *J* = 5.5, 2.1 Hz, 2H), 3.40 (t, *J* = 5.9 Hz, 1H), 3.30 (dt, *J* = 13.9, 5.6 Hz, 1H), 1.61 (s, 3H), 1.34 (s, 4H).

**^13^C NMR** (101 MHz, DMSO) δ 169.7, 169.7, 165.2, 155.9, 148.6, 148.0, 145.2, 137.5, 130.3, 127.8, 127.3, 125.1, 122.8, 122.5, 122.2, 120.0, 119.5, 117.4, 116.6, 116.1, 111.3, 110.2, 101.5, 84.2, 83.8, 82.5, 80.4, 43.0, 26.3, 25.5.

#### N-(2-((((3aR,4R,6R,6aR)-6-(4-aminopyrrolo[2,1-f][1,2,4]triazin-7-yl)-6-cyano-2,2-dimethyltetrahydrofuro[3,4-d][1,3]dioxol-4-yl)methyl)amino)-2-oxoethyl)-5-((4-cyanophenyl)sulfonamido)nicotinamide *(12v)*

**LC-MS (ESI)**: m/z calcd for C_30_H_29_N_10_O_7_S^+^ [M+H]^+^ 673.2, found 673.4.

**^1^H NMR** (400 MHz, DMSO-*d*_6_ + D_2_O) δ 9.01 (t, *J* = 5.8 Hz, 1H), 8.72 (d, *J* = 1.9 Hz, 1H), 8.39 (d, *J* = 2.5 Hz, 1H), 8.19 (t, *J* = 5.9 Hz, 1H), 8.03 (d, *J* = 8.4 Hz, 2H), 7.95 (d, *J* = 10.9 Hz, 2H), 7.93 – 7.88 (m, 2H), 6.92 (d, *J* = 4.6 Hz, 1H), 6.88 (d, *J* = 4.5 Hz, 1H), 5.41 (d, *J* = 6.6 Hz, 1H), 4.84 (dd, *J* = 6.7, 3.3 Hz, 1H), 4.36 (td, *J* = 5.6, 3.2 Hz, 1H), 3.88 – 3.84 (m, 2H), 3.44 – 3.37 (m, 1H), 3.30 (dt, *J* = 13.9, 5.6 Hz, 1H), 1.62 (s, 3H), 1.34 (s, 3H).

**^13^C NMR** (101 MHz, DMSO) δ 169.6, 165.0, 155.9, 148.6, 144.8, 134.0, 130.3, 127.8, 127.6, 127.2, 125.0, 122.5, 119.4, 118.0, 117.4, 116.6, 116.2, 115.9, 111.3, 110.2, 101.5, 84.1, 83.8, 82.5, 80.4, 43.0, 26.3, 25.5.

#### N-(2-((((3aR,4R,6R,6aR)-6-(4-aminopyrrolo[2,1-f][1,2,4]triazin-7-yl)-6-cyano-2,2-dimethyltetrahydrofuro[3,4-d][1,3]dioxol-4-yl)methyl)amino)-2-oxoethyl)-2-methyl-5-((4-(trifluoromethyl)phenyl)sulfonamido)nicotinamide *(12w)*

**LC-MS (ESI)**: m/z calcd for C_31_H_31_F_3_N_9_O_7_S^+^ [M+H]^+^ 730.2, found 730.4.

**^1^H NMR** (400 MHz, DMSO-*d*_6_ + D_2_O) δ 8.67 (t, *J* = 6.0 Hz, 1H), 8.19 (d, *J* = 3.0 Hz, 1H), 8.07 – 7.81 (m, 5H), 7.49 (d, *J* = 2.6 Hz, 1H), 7.05 – 6.77 (m, 2H), 5.43 (d, *J* = 6.6 Hz, 1H), 4.86 (dd, *J* = 6.7, 3.3 Hz, 1H), 4.36 (td, *J* = 5.7, 3.2 Hz, 1H), 3.84 (dd, *J* = 5.7, 2.5 Hz, 2H), 3.41 (s, 1H), 3.38 – 3.25 (m, 1H), 2.43 (s, 3H), 1.62 (s, 3H), 1.34 (s, 3H).

**^13^C NMR** (101 MHz, DMSO) δ 169.5, 168.0, 168.0, 155.9, 152.1, 148.6, 143.4, 143.4, 142.3, 132.4, 128.1, 127.8, 127.2, 127.2, 122.5, 117.4, 116.6, 116.2, 111.3, 101.5, 84.2, 83.9, 82.5, 80.5, 26.3, 25.6, 22.4.

#### N-(2-((((3aR,4R,6R,6aR)-6-(4-aminopyrrolo[2,1-f][1,2,4]triazin-7-yl)-6-cyano-2,2-dimethyltetrahydrofuro[3,4-d][1,3]dioxol-4-yl)methyl)amino)-2-oxoethyl)-4-methyl-5-((4-(trifluoromethyl)phenyl)sulfonamido)nicotinamide *(12x)*

**LC-MS (ESI)**: m/z calcd for C_31_H_31_F_3_N_9_O_7_S^+^ [M+H]^+^ 730.2, found 730.4.

**^1^H NMR** (500 MHz, DMSO-*d*_6_ + D_2_O) δ 8.35 (s, 1H), 8.18 (t, *J* = 5.9 Hz, 1H), 8.07 (s, 1H), 8.03 – 7.94 (m, 3H), 7.91 (d, *J* = 8.3 Hz, 2H), 6.92 (d, *J* = 4.6 Hz, 1H), 6.90 (d, *J* = 4.6 Hz, 1H), 5.41 (d, *J* = 6.7 Hz, 1H), 4.85 (dd, *J* = 6.7, 3.3 Hz, 1H), 4.35 (td, *J* = 5.7, 3.3 Hz, 1H), 3.83 (d, *J* = 3.6 Hz, 2H), 2.08 (s, 3H), 1.61 (s, 3H), 1.32 (s, 3H).

**^13^C NMR** (151 MHz, DMSO) δ 169.6, 169.6, 167.3, 156.0, 155.9, 148.6, 134.0, 128.1, 127.1, 124.8,123.0, 122.5, 117.5, 117.4, 116.6, 116.1, 111.3, 101.5, 84.2, 83.9, 82.5, 80.4, 42.6, 40.7, 26.3, 25.5, 14.6.

#### N-(2-((((3aR,4R,6R,6aR)-6-(4-aminopyrrolo[2,1-f][1,2,4]triazin-7-yl)-6-cyano-2,2-dimethyltetrahydrofuro[3,4-d][1,3]dioxol-4-yl)methyl)amino)-2-oxoethyl)-6-methyl-5-((4-(trifluoromethyl)phenyl)sulfonamido)nicotinamide *(12y)*

**LC-MS (ESI)**: m/z calcd for C_31_H_31_F_3_N_9_O_7_S^+^ [M+H]^+^ 730.2, found 730.4.

**^1^H NMR** (400 MHz, DMSO-*d*_6_ + D_2_O) δ 9.05 (t, *J* = 5.8 Hz, 1H), 8.83 (d, *J* = 2.0 Hz, 1H), 8.25 (t, *J* = 5.9 Hz, 1H), 8.03 (t, *J* = 4.2 Hz, 3H), 7.99 – 7.89 (m, 3H), 6.98 (d, *J* = 4.6 Hz, 1H), 6.95 (d, *J* = 4.6 Hz, 1H), 5.47 (d, *J* = 6.6 Hz, 1H), 4.91 (dd, *J* = 6.7, 3.3 Hz, 1H), 4.42 (td, *J* = 5.6, 3.2 Hz, 1H), 4.02 – 3.81 (m, 2H), 3.47 (t, *J* = 5.9 Hz, 1H), 3.37 (dt, *J* = 13.9, 5.6 Hz, 1H), 2.20 (s, 3H), 1.68 (s, 3H), 1.41 (s, 3H).

**^13^C NMR** (101 MHz, DMSO) δ 169.8, 164.8, 155.9, 148.6, 133.4, 128.4, 127.9, 127.2, 125.2, 122.5, 117.4, 116.6, 116.2, 111.3, 101.5, 84.2, 83.9, 82.5, 80.4, 43.1, 26.3, 25.5, 21.1.

#### N-(2-((((3aR,4R,6R,6aR)-6-(4-aminopyrrolo[2,1-f][1,2,4]triazin-7-yl)-6-cyano-2,2-dimethyltetrahydrofuro[3,4-d][1,3]dioxol-4-yl)methyl)amino)-2-oxoethyl)-4-((4-(trifluoromethyl)phenyl)sulfonamido)picolinamide *(12z)*

**LC-MS (ESI)**: m/z calcd for C_30_H_29_F_3_N_9_O_7_S^+^ [M+H]^+^ 716.2, found 716.4.

**^1^H NMR** (500 MHz, DMSO-*d*6 + D_2_O) δ 8.42 (s, 1H), 8.15 (s, 1H), 8.08 (d, *J* = 8.2 Hz, 2H), 7.99 (d, *J* = 8.3 Hz, 2H), 7.96 (s, 1H), 7.74 (s, 1H), 7.33 – 7.23 (m, 1H), 6.90 (d, *J* = 4.5 Hz, 1H), 6.87 (d, *J* = 4.6 Hz, 1H), 5.39 (d, *J* = 6.6 Hz, 1H), 4.82 (dd, *J* = 6.7, 3.3 Hz, 1H), 4.34 (td, *J* = 5.6, 3.2 Hz, 1H), 3.88 (s, 2H), 3.39 (t, *J* = 6.2 Hz, 1H), 3.30 (dt, *J* = 13.4, 5.6 Hz, 1H), 1.61 (s, 3H), 1.34 (s, 3H).

**^13^C NMR** (101 MHz, DMSO) δ 167.4, 158.9, 155.9, 148.6, 143.5, 140.3, 128.1, 127.4, 125.4, 122.5, 117.4, 116.6, 116.1, 114.9, 111.3, 101.4, 99.1, 91.3, 83.8, 82.4, 80.4, 26.3, 25.6.

#### N-(2-((((3aR,4R,6R,6aR)-6-(4-aminopyrrolo[2,1-f][1,2,4]triazin-7-yl)-6-cyano-2,2-dimethyltetrahydrofuro[3,4-d][1,3]dioxol-4-yl)methyl)amino)-2-oxoethyl)-2-((4-(trifluoromethyl)phenyl)sulfonamido)iso nicotinamide *(12aa)*

**LC-MS (ESI)**: m/z calcd for C_30_H_29_F_3_N_9_O_7_S^+^ [M+H]^+^ 716.2, found 716.4.

**^1^H NMR** (500 MHz, DMSO-*d*_6_ + D_2_O) δ 8.16 (t, *J* = 5.9 Hz, 1H), 8.09 – 8.00 (m, 3H), 7.97 (s, 1H), 7.85 (d, *J* = 8.2 Hz, 2H), 7.32 (s, 1H), 7.07 (d, *J* = 5.6 Hz, 1H), 6.92 (d, *J* = 4.6 Hz, 1H), 6.90 (d, *J* = 4.6 Hz, 1H), 5.41 (d, *J* = 6.6 Hz, 1H), 4.84 (dd, *J* = 6.6, 3.3 Hz, 1H), 4.36 (td, *J* = 5.7, 3.2 Hz, 1H), 3.83 (d, *J* = 3.0 Hz, 2H), 3.43 – 3.37 (m, 1H), 3.30 (dt, *J* = 13.7, 5.4 Hz, 1H), 1.62 (s, 3H), 1.34 (s, 3H).

**^13^C NMR** (101 MHz, DMSO) δ 169.5, 165.5, 155.9, 148.6, 127.9, 126.2, 125.5, 122.8, 122.5, 117.4, 116.6, 116.1, 111.3, 101.5, 84.2, 83.8, 82.5, 80.4, 43.0, 26.3, 25.6.

#### N-(2-((((3aR,4R,6R,6aR)-6-(4-aminopyrrolo[2,1-f][1,2,4]triazin-7-yl)-6-cyano-2,2-dimethyltetrahydrofuro[3,4-d][1,3]dioxol-4-yl)methyl)amino)-2-oxoethyl)-6-((4-(trifluoromethyl)phenyl)sulfonamido)picolinamide *(12ab)*

**LC-MS (ESI)**: m/z calcd for C_30_H_29_F_3_N_9_O_7_S^+^ [M+H]^+^ 716.2, found 716.4.

**^1^H NMR** (500 MHz, DMSO-*d*_6_ + D_2_O) δ 8.34 (t, *J* = 5.9 Hz, 1H), 8.28 (d, *J* = 8.1 Hz, 2H), 8.00 – 7.90 (m, 3H), 7.88 (t, *J* = 7.9 Hz, 1H), 7.60 (d, *J* = 7.4 Hz, 1H), 7.16 (d, *J* = 8.2 Hz, 1H), 6.92 (s, 2H), 5.44 (d, *J* = 6.7 Hz, 1H), 4.88 (dd, *J* = 6.7, 3.4 Hz, 1H), 4.39 (td, *J* = 5.7, 3.3 Hz, 1H), 3.98 (s, 2H), 3.46 – 3.37 (m, 2H), 1.62 (s, 3H), 1.34 (s, 3H).

**^13^C NMR** (101 MHz, DMSO) δ 169.3, 163.6, 155.9, 148.6, 147.9, 140.7, 128.7, 127.0, 122.5, 117.4, 116.5, 116.2, 115.7, 111.3, 101.5, 84.2, 83.9, 82.5, 80.5, 26.3, 25.6.

#### ((3aR,4R,6R,6aR)-6-(4-aminopyrrolo[2,1-f][1,2,4]triazin-7-yl)-6-cyano-2,2-dimethyltetrahydrofuro[3, 4-d][1,3]dioxol-4-yl)methyl 4-methylbenzenesulfonate *(14)*

To a stirring solution of **2** (200 mg, 0.603 mmol, 1 eq.) in DCM (4 mL) was sequentially added DMAP (81 mg, 0.663 mmol, 1.1 eq.) and TsCl (115 mg, 0.603 mmol, 1 eq.). The reaction mixture was allowed to stir at room temperature overnight. The solvent was evaporated and the crude mixture was purified by column chromatography (hexanes: EA 1:1 to 1:2) to afford the product as a white solid (170 mg, yield: 58%).

**LC-MS (ESI):** m/z calcd for C_22_H_24_N_5_O_6_S^+^ [M+H]^+^ 486.1, found 486.2.

**^1^H NMR** (500 MHz, Chloroform-*d*) δ 7.96 (d, *J* = 1.8 Hz, 1H), 7.72 (dd, *J* = 8.4, 1.9 Hz, 2H), 7.25 (d, *J* = 8.0 Hz, 2H), 6.99 (dd, *J* = 4.7, 1.9 Hz, 1H), 6.63 (dd, *J* = 4.7, 1.9 Hz, 1H), 5.54 (s, 2H), 5.45 (dd, *J* = 6.8, 1.9 Hz, 1H), 4.96 (dd, *J* = 6.7, 3.9 Hz, 1H), 4.55 (q, *J* = 4.4 Hz, 1H), 4.37 – 4.24 (m, 2H), 2.42 (s, 3H), 1.76 (s, 3H), 1.41 (s, 3H).

**^13^C NMR** (126 MHz, CDCl_3_) δ 155.2, 147.3, 145.2, 132.3, 129.8, 128.0, 123.1, 117.3, 116.7, 115.5, 112.6, 99.9, 83.8, 82.3, 81.6, 81.4, 68.1, 26.3, 25.5, 21.7.

### (3aR,4R,6R,6aR)-4-(4-aminopyrrolo[2,1-f][1,2,4]triazin-7-yl)-2,2-dimethyl-6-((methylamino)methyl)t etrahydrofuro[3,4-d][1,3]dioxole-4-carbonitrile *(15)*

In a microwave vial, **14** (50 mg, 0.103 mmol, 1 eq.) was dissolved with THF (1.6 mL), followed by the addition of 40% MeNH_2_ in water (0.39 mL, 5.03 mmol, 50 eq.). The reaction mixture was subjected to microwave irradiation for 90 minutes at 100 . The solvent was evaporated and the crude was purified on a silica gel column (DCM: MeOH 50:1 to 10:1 with 1% TEA) to afford the product as a white solid (22 mg, yield: 62%).

**LC-MS (ESI):** m/z calcd for C H N O ^+^ [M+H]^+^ 345.2, found 345.2.

**^1^H NMR** (400 MHz, DMSO-*d*_6_) δ 8.00 (br, 2H), 7.97 (s, 1H), 6.94 (s, 2H), 5.46 (d, *J* = 6.7 Hz, 1H), 4.99 (dd, *J* = 6.7, 3.5 Hz, 1H), 4.49 (ddd, *J* = 7.0, 4.6, 3.5 Hz, 1H), 3.10 – 2.94 (m, 2H), 2.44 (s, 3H), 1.65 (s, 3H), 1.37 (s, 3H).

### (9H-fluoren-9-yl)methyl

#### (2-((((3aR,4R,6R,6aR)-6-(4-aminopyrrolo[2,1-f][1,2,4]triazin-7-yl)-6-cyano-2,2-dimethyltetrahydrofur o[3,4-d][1,3]dioxol-4-yl)methyl)(methyl)amino)-2-oxoethyl)carbamate *(16)*

General Procedure A was followed. **15** was reacted with Fmoc-Gly-OH to afford the product as a white solid (yield: 84%).

**LC-MS (ESI):** m/z calcd for C H N O ^+^ [M+H]^+^ 624.3, found 624.3.

**^1^H NMR** (500 MHz, DMSO-*d*_6_) δ 8.36 (s, 1H), 8.07 – 7.86 (m, 5H), 7.73 (t, *J* = 8.2 Hz, 2H), 7.42 (t, *J* = 7.3 Hz, 2H), 7.33 (t, *J* = 7.4 Hz, 2H), 6.94 (d, *J* = 4.7 Hz, 1H), 6.93 – 6.90 (m, 1H), 5.49 (d, *J* = 6.8 Hz, 0.3H), 5.45 (d, *J* = 6.6 Hz, 0.7H), 4.93 (dd, *J* = 6.8, 4.0 Hz, 0.3H), 4.88 (dd, *J* = 6.7, 3.8 Hz, 0.7H), 4.53 – 4.46 (m, 0.3H), 4.42 (td, *J* = 5.9, 3.7 Hz, 0.7H), 4.31 – 4.21 (m, 3H), 3.86 (d, *J* = 6.0 Hz, 1.4H), 3.79 –3.73 (m, 0.6H), 3.71 – 3.65 (m, 1.4H), 3.42 (dd, *J* = 14.0, 6.4 Hz, 0.6H), 2.89 (s, 2H), 2.83 (s, 1H), 1.65 (s, 1H), 1.62 (s, 2H), 1.37 (s, 1H), 1.34 (s, 2H).

**^13^C NMR** (126 MHz, DMSO) δ 169.8, 169.3, 157.0, 156.1, 148.7, 144.3, 141.2, 128.1, 127.6, 125.8, 122.5, 120.6, 117.6, 116.2, 111.2, 101.4, 84.1, 83.8, 82.6, 80.3, 66.2, 54.0, 49.4, 47.1, 42.6, 35.8, 31.2, 26.5, 25.7.

### 2-amino-N-(((3aR,4R,6R,6aR)-6-(4-aminopyrrolo[2,1-f][1,2,4]triazin-7-yl)-6-cyano-2,2-dimethyltetrahydrofuro[3,4-d][1,3]dioxol-4-yl)methyl)-N-methylacetamide *(17)*

**16** (144 mg, 0.23 mmol, 1 eq.) was dissolved with DCM (10 mL). A 2 M solution of dimethylamine in THF (2.3 mL, 10 eq.) was added and the reaction mixture was allowed to stir at room temperature overnight. The solvent was evaporated and the crude was purified by column chromatography (DCM: MeOH 50:1 to 10:1 with 1% TEA) to afford the product as a white solid (78 mg, yield: 85%).

**LC-MS (ESI):** m/z calcd for C H N O ^+^ [M+H]^+^ 402.2, found 402.2.

**^1^H NMR** (400 MHz, DMSO-*d*_6_) δ 8.11 – 7.82 (m, 3H), 6.98 – 6.77 (m, 2H), 5.51 – 5.39 (m, 1H), 4.92 –4.82 (m, 1H), 4.47 – 4.37 (m, 1H), 3.74 – 3.56 (m, 1H), 3.45 – 3.12 (m, 3H), 1.63 (s, 1H), 1.60 (s, 2H), 1.35 (s, 1H), 1.33 (s, 2H).

For target compounds **13**, **18a-18d** and appropriate carboxylic acids were reacted following General Procedure A. The yields generally ranged from 70% to 90%.

#### N-(2-((((3aR,4R,6R,6aR)-6-(4-aminopyrrolo[2,1-f][1,2,4]triazin-7-yl)-6-cyano-2,2-dimethyltetrahydrofuro[3,4-d][1,3]dioxol-4-yl)methyl)amino)-2-oxoethyl)-N-methyl-3-(phenylsulfonamido)benzamide *(13)*

**LC-MS (ESI)**: m/z calcd for C_31_H_33_N_8_O_7_S^+^ [M+H]^+^ 661.2, found 661.3.

**^1^H NMR** (500 MHz, DMSO-*d*_6_) δ 8.02 – 7.88 (m, 1H), 7.74 (t, *J* = 7.6 Hz, 2H), 7.65 – 7.46 (m, 3H), 7.38 – 7.20 (m, 1H), 7.19 – 6.96 (m, 3H), 6.95 – 6.86 (m, 2H), 5.42 (t, *J* = 6.2 Hz, 1H), 4.84 (ddd, *J* = 17.3, 6.7, 3.4 Hz, 1H), 4.42 – 4.25 (m, 1H), 3.70 (s, 1H), 3.45 – 3.27 (m, 4H), 2.86 (s, 1.5H), 2.73 (s, 1.5H), 1.62 (s, 3H), 1.34 (s, 3H).

#### N-(2-((((3aR,4R,6R,6aR)-6-(4-aminopyrrolo[2,1-f][1,2,4]triazin-7-yl)-6-cyano-2,2-dimethyltetrahydrofuro[3,4-d][1,3]dioxol-4-yl)methyl)(methyl)amino)-2-oxoethyl)-3-(phenylsulfonamido)benzamide *(18a)*

**LC-MS (ESI)**: m/z calcd for C_31_H_33_N_8_O_7_S^+^ [M+H]^+^ 661.2, found 661.3.

**^1^H NMR** (500 MHz, DMSO-*d*_6_ + D_2_O) δ 7.98 (s, 0.7H), 7.95 (s, 0.3H), 7.82 – 7.70 (m, 2H), 7.66 – 7.51 (m, 5H), 7.38 – 7.30 (m, 1H), 7.28 – 7.21 (m, 1H), 6.99 – 6.88 (m, 2H), 5.50 (d, *J* = 6.8 Hz, 0.3H), 5.44 (d, *J* = 6.7 Hz, 0.7H), 4.98 (dd, *J* = 6.8, 4.0 Hz, 0.3H), 4.87 (dd, *J* = 6.7, 3.7 Hz, 0.7H), 4.52 (td, *J* = 6.8, 6.2, 4.0 Hz, 0.3H), 4.42 (td, *J* = 5.9, 3.7 Hz, 0.7H), 4.20 – 3.99 (m, 1.7H), 3.90 (dd, *J* = 16.4, 4.4 Hz, 0.3H), 3.76 – 3.71 (m, 0.3H), 3.67 (dd, *J* = 14.1, 5.5 Hz, 0.7H), 3.45 – 3.40 (overlapped with water peak, 1H), 2.94 (s, 2H), 2.83 (s, 1H), 1.65 (s, 1H), 1.61 (s, 2H), 1.38 (s, 1H), 1.34 (s, 2H).

**^13^C NMR** (126 MHz, DMSO) δ 169.5, 169.1, 166.4, 166.3, 155.9, 148.7, 148.6, 139.7, 138.3, 135.6, 135.6, 133.5, 129.8, 129.7, 127.1, 123.1, 123.1, 122.5, 122.0, 119.8, 117.6, 117.4, 116.8, 116.5, 116.2, 111.4, 111.2, 101.4, 84.1, 83.8, 83.1, 82.5, 82.3, 80.7, 80.2, 55.3, 50.2, 49.4, 41.6, 41.5, 36.0, 34.1, 26.5, 26.4, 25.7.

#### N-(2-((((3aR,4R,6R,6aR)-6-(4-aminopyrrolo[2,1-f][1,2,4]triazin-7-yl)-6-cyano-2,2-dimethyltetrahydrofuro[3,4-d][1,3]dioxol-4-yl)methyl)(methyl)amino)-2-oxoethyl)-5-((4-fluorophenyl)sulfonamido)nicotinamide *(18b)*

**LC-MS (ESI)**: m/z calcd for C_30_H_31_FN_9_O_7_S^+^ [M+H]^+^ 680.2, found 680.4

**^1^H NMR** (400 MHz, DMSO-*d*_6_ + D_2_O) δ 8.88 (t, *J* = 5.7 Hz, 0.7H), 8.79 (t, *J* = 5.7 Hz, 0.3H), 8.72 (d, *J* = 19.2 Hz, 1H), 8.46 – 8.35 (m, 1H), 8.02 – 7.77 (m, 4H), 7.41 (t, *J* = 8.8 Hz, 2H), 6.92 (q, *J* = 4.6 Hz, 2H), 5.51 (d, *J* = 6.8 Hz, 0.3H), 5.44 (d, *J* = 6.7 Hz, 0.7H), 4.98 (dd, *J* = 6.8, 4.0 Hz, 0.3H), 4.87 (dd, *J* =, 3.7 Hz, 0.7H), 4.52 (dt, *J* = 7.6, 4.6 Hz, 0.3H), 4.43 (td, *J* = 5.9, 3.7 Hz, 0.7H, 4.25 – 4.04 (m, 1.7H), 3.93 (dd, *J* = 16.4, 5.2 Hz, 0.3H), 3.80 – 3.61 (m, 1.3H), 3.44 – 3.39 (m, 0.7H), 2.95 (s, 2H), 2.84 (s, 1H), 1.65 (s, 1H), 1.61 (s, 2H), 1.38 (s, 1H), 1.34 (s, 2H).

**^13^C NMR** (101 MHz, DMSO) δ 169.3, 168.8, 166.3, 164.9, 164.8, 163.8, 155.9, 148.6, 144.4, 144.3, 135.6, 134.6, 130.5, 130.3, 130.2, 127.0, 122.5, 122.0, 117.5, 117.4, 117.1, 116.8, 116.5, 116.3, 111.5, 111.3, 101.5, 84.1, 83.8, 82.5, 82.3, 80.7, 80.2, 50.3, 49.4, 41.6, 40.5, 40.3, 40.1, 39.9, 39.6, 39.4, 39.2, 36.0, 34.1, 26.5, 26.4, 25.7.

#### N-(2-((((3aR,4R,6R,6aR)-6-(4-aminopyrrolo[2,1-f][1,2,4]triazin-7-yl)-6-cyano-2,2-dimethyltetrahydrofuro[3,4-d][1,3]dioxol-4-yl)methyl)(methyl)amino)-2-oxoethyl)-5-((4-(trifluoromethyl)phenyl)sulfonamido)nicotinamide *(18c)*

**LC-MS (ESI)**: m/z calcd for C_31_H_31_F_3_N_9_O_7_S^+^ [M+H]^+^ 730.2, found 730.4.

**^1^H NMR** (400 MHz, DMSO-*d*_6_ + D_2_O) δ 8.72 (d, *J* = 1.9 Hz, 0.7H), 8.67 (d, *J* = 1.9 Hz, 0.3H), 8.40 (d, *J* = 2.5 Hz, 0.7H), 8.38 (d, *J* = 2.5 Hz, 0.3H), 8.01 – 7.93 (m, 5H), 7.92 (t, *J* = 2.2 Hz, 0.7H), 7.88 (t, *J* = 2.2 Hz, 0.3H), 6.98 – 6.87 (m, 2H), 5.51 (d, *J* = 6.8 Hz, 0.3H), 5.44 (d, *J* = 6.7 Hz, 0.7H), 4.98 (dd, *J* =, 4.0 Hz, 0.3H), 4.87 (dd, *J* = 6.7, 3.7 Hz, 0.7H), 4.52 (dt, *J* = 7.5, 4.8 Hz, 0.3H), 4.43 (td, *J* = 5.8, 3.6 Hz, 0.7H), 4.25 – 4.04 (m, 1.7H), 3.97 – 3.89 (m, 0.3H), 3.76 – 3.65 (m, 1H), 3.42 (overlapped with water peak, 1H), 2.95 (s, 2H), 2.84 (s, 1H), 1.65 (s, 1H), 1.61 (s, 2H), 1.38 (s, 1H), 1.34 (s, 2H).

**^13^C NMR** (101 MHz, DMSO) δ 169.3, 169.3, 164.9, 155.9, 148.6, 144.7, 143.9, 133.3, 133.0, 130.5, 128.1, 127.2, 127.1, 125.2, 122.5, 122.4, 122.0, 117.5, 116.8, 116.5, 116.2, 111.2, 101.4, 84.1, 83.8, 82.5, 80.7, 80.2, 49.4, 36.0, 34.1, 26.5, 26.4, 25.6.

#### N-(2-((((3aR,4R,6R,6aR)-6-(4-aminopyrrolo[2,1-f][1,2,4]triazin-7-yl)-6-cyano-2,2-dimethyltetrahydrofuro[3,4-d][1,3]dioxol-4-yl)methyl)(cyclopropyl)amino)-2-oxoethyl)-5-((4-fluorophenyl)sulfonamido) nicotinamide *(18d)*

**LC-MS (ESI)**: m/z calcd for C_32_H_33_FN_9_O_7_S^+^ [M+H]^+^ 706.2, found 706.3.

**^1^H NMR** (500 MHz, DMSO-*d*_6_ + D_2_O) δ 8.71 (S, 1H), 8.39 (d, *J* = 2.6 Hz, 1H), 7.97 (s, 1H), 7.90 (t, *J* = 2.3 Hz, 1H), 7.86 – 7.80 (m, 2H), 7.44 – 7.33 (m, 2H), 6.93 (d, *J* = 4.6 Hz, 1H), 6.90 (d, *J* = 4.6 Hz, 1H), 5.45 (d, *J* = 6.6 Hz, 1H), 4.88 (dd, *J* = 6.7, 3.7 Hz, 1H), 4.46 (td, *J* = 6.1, 3.7 Hz, 1H), 4.26 (qd, *J* = 16.8, 4.9 Hz, 2H), 3.70 (dd, *J* = 14.1, 5.5 Hz, 1H), 3.39 (overlapped with water peak, 1H), 2.75 (h, *J* = 4.5, 3.5 Hz, 1H), 1.61 (s, 3H), 1.35 (s, 3H), 0.82 – 0.65 (m, 4H).

**^13^C NMR** (126 MHz, DMSO) δ 171.7, 165.9, 165.0, 163.9, 155.9, 155.9, 148.6, 144.5, 130.4, 130.2, 130.2, 126.9, 122.5, 117.2, 117.1, 116.3, 111.2, 101.4, 84.1, 83.3, 82.7, 80.2, 48.2, 42.3, 40.3, 30.0, 26.5, 25.6, 9.3.

#### (S)-5-((benzyloxy)methyl)dihydrofuran-2(3H)-one *(20)*

To a solution of (S)-5-(hydroxymethyl)dihydrofuran-2(3H)-one (**19**, 25g, 215 mmol, 1 eq.) in DCM (200 mL) was added BnBr (28.1 mL, 237 mml, 1.1 eq.) and a small spatula of tetra-*n*-butylammonium iodide (TBAI). The reaction was cooled in an ice bath, and NaH (60% in paraffin, 12.9 g, 323 mmol, 1.5 eq) was slowly added in potions so that gas evolution was not too vigorous. Upon complete addition of NaH, the ice bath was removed, and the reaction mixture was allowed to stir overnight. Saturated NH_4_Cl (5 mL) was added to quench unreacted NaH, followed by the addition of water (50 mL). Phases were separated and the aqueous phase was extracted twice more with DCM (50 mL). The combined organic phase was washed with brine and dried over Na_2_SO_4_ before being evaporated to dryness. Column chromatography (Hexanes: EA 3:1) afforded the product as a colorless oil (23.6 g, yield: 53%)

**^1^H NMR** (500 MHz, Chloroform-*d*) δ 7.39 – 7.27 (m, 5H), 4.68 (ddt, *J* = 7.9, 6.1, 3.8 Hz, 1H), 4.62 –4.53 (m, 2H), 3.69 (dd, *J* = 10.7, 3.4 Hz, 1H), 3.59 (dd, *J* = 10.7, 4.1 Hz, 1H), 2.63 (ddd, *J* = 16.7, 10.0, 6.6 Hz, 1H), 2.49 (ddd, *J* = 17.5, 10.1, 7.0 Hz, 1H), 2.30 (dddd, *J* = 13.0, 10.2, 7.9, 6.6 Hz, 1H), 2.22 –2.08 (m, 1H).

**^13^C NMR** (126 MHz, CDCl_3_) δ 177.3, 137.7, 128.5, 127.9, 127.7, 79.0, 71.6, 28.4, 24.2.

#### (S)-2-(4-aminopyrrolo[2,1-f][1,2,4]triazin-7-yl)-5-((benzyloxy)methyl)tetrahydrofuran-2-ol *(21)*

Lactone **20** (23.6 g, 115 mmol, 1 eq.) and 2-methoxyethoxymethyl chloride (16.8 g, 172 mmol, 1.5 eq.) was combined and dissolved with THF (100 mL). The reaction was cooled in an ice bath and a 2 M solution of *i*PrMgCl in THF (172 mL, 344 mmol, 3 eq.) was slowly added. After 2 hours, TLC indicated full conversion of **20**, and 200 mL of saturated NH_4_Cl was added. The resulting mixture was extracted with EA (200 mL, three times). The combined organic phase was washed with brine, dried over anhydrous Na_2_SO_4_ and evaporated to dryness to afford the crude Weinreb amide as a colorless oil (28.6 g) that was directly used in the next conversion.

The crude Weinreb amide (28.6 g, 101 mmol, 1 eq.) and imidazole (12.3 g, 181 mmol, 1.8 eq.) were combined and dissolved with DCM (300 mL). The solution was cooled to -15, and TMSCl (15.3 mL, 121 mmol, 1.2 eq.) was slowly added. Upon the complete addition of TMSCl, the reaction was allowed to warm to 0 and was kept at this temperature for 2 hours. The reaction mixture was sequentially washed with water (100 mL) and brine and dried over anhydrous Na_2_SO_4_. The organic phase was evaporated to dryness to afford the crude TMS-protected Weinreb amide.

To prepare the Grignard reagent, 7-iodopyrrolo[2,1-f][1,2,4]triazin-4-amine (26.2 g, 101 mmol, 1 eq.) was dissolved with THF (300 mL) and cooled to -15 . TMSCl (28.1 mL, 221 mmol, 2.2 eq.) was slowly added and the mixture was allowed to stir at -15 for 15 minutes. MeMgBr (3 M in Et_2_O, 67 mL, 201 mmol, 2 eq.) was slowly added through an addition funnel and the reaction mixture was kept at the same temperature for 20 minutes upon complete addition. Finally, *i*PrMgCl·LiCl (1.3 M in THF, 108 mL, 141 mmol, 1.4 eq.) was slowly added to the reaction mixture. Upon complete addition of *i*PrMgCl·LiCl, the reaction mixture was allowed to stir for another 30 minutes. The previously prepared TMS-protected Weinreb amide in THF (100 mL) was slowly added to the reaction mixture, after which the reaction was allowed to warm to 0 and was kept at this temperature for 1 hour. The pH of the reaction mixture was adjusted to 1∼2 through the addition of 1 N HCl. The solvent was largely evaporated under vacuum and the resulting mixture was diluted with EA (500 mL), washed with brine, dried over Na_2_SO_4_ and again evaporated. The crude was purified through column chromatography (EA: MeOH 60:1) to give the desired product as a white solid (26.2 g, three-step yield: 67%).

**LC-MS (ESI)**: m/z calcd for C H N O ^+^ [M+H]^+^ 341.2, found 341.2.

**^1^H NMR** (500 MHz, Chloroform-*d*) δ 8.12 (s, 1H), 7.38 (d, *J* = 4.8 Hz, 1H), 7.35 – 7.27 (m, 6H), 6.62 (d, *J* = 4.8 Hz, 1H), 5.73 (s, 2H), 5.55 (s, 1H), 4.57 (s, 2H), 3.95 (tt, *J* = 8.1, 3.6 Hz, 1H), 3.57 (dd, *J* = 9.6, 3.5 Hz, 1H), 3.44 (dd, *J* = 9.6, 7.2 Hz, 1H), 3.32 (t, *J* = 7.0 Hz, 2H).

**^13^C NMR** (126 MHz, CDCl_3_) δ 190.9, 155.4, 148.4, 138.0, 129.9, 128.5, 127.8, 117.9, 117.7, 100.9, 74.5, 73.4, 69.9, 60.4, 37.8, 27.5.

### (2R,5S)-2-(4-aminopyrrolo[2,1-f][1,2,4]triazin-7-yl)-5-((benzyloxy)methyl)tetrahydrofuran-2-carbonitrile *(22)*

At -78, to a solution of **21** (2.7 g, 7.93 mol, 1 eq.) in DCM (80 mL) was added TfOH (1.40 mL, 15.9 mmol, 2 eq.), followed by the addition of TMSOTf (3.07 mL, 16.9 mmol, 2.1 eq.). The mixture was allowed to stir for 30 minutes before TMSCN (3.97 mL, 31.7 mmol, 4 eq.) was added. The reaction mixture was kept -78 for 2 hours and then quenched with TEA (5 mL). The resulting mixture was stirred at room temperature for 1 hour and then evaporated to dryness under vacuum. Column chromatography (Hexanes: EA 1:1) afforded the desired β isomer as a syrup (0.86 g, yield: 31%).

**LC-MS (ESI)**: m/z calcd for C H N O ^+^ [M+H]^+^ 350.2, found 350.2.

**^1^H NMR** (500 MHz, Chloroform-*d*) δ 8.03 (s, 1H), 7.37 – 7.29 (m, 5H), 6.84 (d, *J* = 4.6 Hz, 1H), 6.60 (d, *J* = 4.6 Hz, 1H), 5.68 (s, 2H), 4.69 – 4.61 (m, 1H), 4.58 (s, 2H), 3.69 – 3.61 (m, 2H), 2.89 (ddd, *J* = 12.9, 8.0, 4.9 Hz, 1H), 2.77 (dt, *J* = 13.0, 8.5 Hz, 1H), 2.43 (dq, *J* = 12.7, 8.0 Hz, 1H), 2.18 – 2.08 (m, 1H).

**^13^C NMR** (126 MHz, CDCl_3_) δ 155.2, 147.3, 138.0, 128.4, 127.8, 127.7, 125.9, 119.0, 116.0, 110.6, 99.6, 80.1, 75.6, 73.4, 71.5, 36.8, 27.3.

### (2R,5S)-2-(4-aminopyrrolo[2,1-f][1,2,4]triazin-7-yl)-5-(hydroxymethyl)tetrahydrofuran-2-carbonitrile *(23)*

At -78, to a solution of **22** (0.53 g, 1.52 mol, 1 eq.) in DCM (10 mL) was added BCl_3_ (1 M in DCM, 4.60 mL, 4.60 mmol, 3 eq.). The reaction mixture was stirred at -78 for 4 hours and then quenched by MeOH (1 mL). The resulting mixture was directly evaporated to dryness and purified by column chromatography (DCM: MeOH 60:1 to 20:1) to afford the product as a white solid (0.14 g, yield: 47%)

**LC-MS (ESI)**: m/z calcd for C H N O ^+^ [M+H]^+^ 260.1, found 260.1.

**^1^H NMR** (500 MHz, Methanol-*d*_4_) δ 7.83 (s, 1H), 6.87 (d, *J* = 4.5 Hz, 1H), 6.86 (d, *J* = 4.5 Hz, 1H), 4.45 (ddt, *J* = 8.0, 5.5, 4.0 Hz, 1H), 3.71 (dd, *J* = 12.0, 3.8 Hz, 1H), 3.61 (dd, *J* = 12.0, 4.7 Hz, 1H), 2.77 – 2.72 (m, 2H), 2.36 (dq, *J* = 12.7, 8.1 Hz, 1H), 2.07 (ddt, *J* = 13.4, 7.9, 5.6 Hz, 1H).

**^13^C NMR** (126 MHz, MeOD) δ 155.8, 146.8, 125.3, 118.8, 116.3, 110.0, 101.2, 81.7, 76.2, 63.6, 36.6, 26.3.

### ((2S,5R)-5-(4-aminopyrrolo[2,1-f][1,2,4]triazin-7-yl)-5-cyanotetrahydrofuran-2-yl)methyl 4-methylbenzenesulfonate *(24)*

**23** (70 mg, 0.270 mmol, 1 eq.) was dissolved in pyridine (2 mL), followed by the addition of TsCl (62 mg, 0.324 mmol, 1.2 eq.). The reaction mixture was stirred at room temperature overnight and then evaporated to dryness. The crude was purified by column chromatography (DCM: MeOH 40:1) to afford the product as an off-white solid (65 mg, yield: 58%)

**LC-MS (ESI)**: m/z calcd for C_19_H_20_N_5_O_4_S^+^ [M+H]^+^ 414.1, found 414.2.

**^1^H NMR** (500 MHz, Chloroform-*d*) δ 7.94 (s, 1H), 7.69 (d, *J* = 8.1 Hz, 2H), 7.24 (d, *J* = 8.0 Hz, 2H), 6.74 (d, *J* = 4.6 Hz, 1H), 6.60 (d, *J* = 4.6 Hz, 1H), 5.98 (s, 2H), 4.59 (dq, *J* = 9.1, 4.6 Hz, 1H), 4.21 – 4.05 (m, 2H), 2.88 – 2.71 (m, 2H), 2.46 (dt, *J* = 13.0, 8.3 Hz, 1H), 2.40 (s, 3H), 2.16 – 2.08 (m, 1H).

**^13^C NMR** (126 MHz, CDCl_3_) δ 155.4, 147.3, 145.2, 132.4, 129.8, 127.9, 124.8, 118.6, 116.3, 110.7, 99.9, 77.9, 75.8, 70.3, 36.4, 27.0, 21.7.

### (2R,5S)-2-(4-aminopyrrolo[2,1-f][1,2,4]triazin-7-yl)-5-(azidomethyl)tetrahydrofuran-2-carbonitrile *(25)*

NaN_3_ (19 mg, 0.295 mmol, 2 eq.) was added to a stirring solution of **24** (61 mg, 0.148 mmol, 1 eq.) in DMF (1 mL). The reaction mixture was stirred at 80 for 2 hours, followed by the addition of water (4 mL). The resulting mixture was extracted three times with EA (3 mL). The combined EA phase was washed with brine, dried over Na_2_SO_4_ and evaporated to afford desired product (24 mg, yield: 82%).

**LC-MS (ESI)**: m/z calcd for C_12_H_13_N_8_O^+^ [M+H]^+^ 285.1, found 285.1.

**^1^H NMR** (500 MHz, Chloroform-*d*) δ 8.05 (s, 1H), 6.88 (d, *J* = 4.6 Hz, 1H), 6.62 (d, *J* = 4.6 Hz, 1H), 5.46 (s, 2H), 4.62 (dtd, *J* = 7.9, 5.4, 4.0 Hz, 1H), 3.56 (dd, *J* = 13.0, 4.0 Hz, 1H), 3.45 (dd, *J* = 13.1, 5.1 Hz, 1H), 2.94 (ddd, *J* = 12.8, 8.1, 4.5 Hz, 1H), 2.78 (dt, *J* = 13.1, 8.7 Hz, 1H), 2.46 (dq, *J* = 12.9, 8.2 Hz, 1H), 2.11 (dddd, *J* = 13.2, 9.0, 5.8, 4.6 Hz, 1H).

### (2R,5S)-5-(aminomethyl)-2-(4-aminopyrrolo[2,1-f][1,2,4]triazin-7-yl)tetrahydrofuran-2-carbonitrile *(26)*

Compound **25** (29 mg, 0.103 mmol, 1 eq.) and PPh_3_ (54 mg, 0.206 mmol, 2 eq.) were combined and dissolved in 1,4-dioxane (1 mL). The reaction mixture was stirred for 30 minutes before 1 mL of water was added, after which the reaction was brought to reflux and allowed to stir overnight. The solvent was evaporated and the crude mixture was purified by silica gel column chromatography (100% MeCN to MeCN: water 9:1 with 0.6% TEA) to afford the product as an off-white solid (19 mg, yield: 71%)

**LC-MS (ESI)**: m/z calcd for C_12_H_15_N_6_O^+^ [M+H]^+^ 259.1, found 259.1.

**^1^H NMR** (500 MHz, DMSO-*d*_6_) δ 7.94 (s, 1H), 7.89 (br, 2H), 6.90 (d, *J* = 4.5 Hz, 1H), 6.73 (d, *J* = 4.4 Hz, 1H), 4.52 (p, *J* = 6.2 Hz, 1H), 3.40 – 3.25 (m, 2H), 2.81 – 2.74 (m, 1H), 2.67 (ddd, *J* = 12.9, 8.5, 7.4 Hz, 1H), 2.38 – 2.23 (m, 2H), 2.01 – 1.88 (m, 2H).

### tert-butyl

#### (2-((((2S,5R)-5-(4-aminopyrrolo[2,1-f][1,2,4]triazin-7-yl)-5-cyanotetrahydrofuran-2-yl)methyl)amino)-2-oxoethyl)carbamate *(27)*

General Procedure A was followed. **26** was reacted with Boc-Gly-OH to yield the product as a white powder (yield: 78%).

**LC-MS (ESI)**: m/z calcd for C H N O ^+^ [M+H]^+^ 416.2, found 416.2.

**^1^H NMR** (500 MHz, Chloroform-*d*) δ 8.07 (s, 1H), 7.14 (s, 1H), 6.98 (d, *J* = 4.7 Hz, 1H), 6.65 (d, *J* = 4.6 Hz, 1H), 5.68 (s, 2H), 4.64 – 4.56 (m, 1H), 3.93 – 3.72 (m, 2H), 2.95 (dt, *J* = 13.1, 9.3 Hz, 1H), 2.59 (ddd, *J* = 12.7, 5.8, 2.1 Hz, 1H), 2.55 – 2.39 (m, 1H), 2.06 (dddd, *J* = 13.0, 9.2, 5.3, 3.5 Hz, 1H), 1.42 (s, 9H).

**^13^C NMR** (126 MHz, CDCl_3_) δ 169.6, 155.4, 147.6, 124.4, 118.8, 116.8, 112.4, 100.1, 79.4, 53.4, 44.1, 42.5, 36.3, 28.3, 27.3.

### 2-amino-N-(((2S,5R)-5-(4-aminopyrrolo[2,1-f][1,2,4]triazin-7-yl)-5-cyanotetrahydrofuran-2-yl)methyl)acetamide *(28)*

To a solution of **27** (184 mg, 0.443 mmol, 1 eq.) in DCM (10 mL) was added 4 M HCl in 1,4-dioxane (210 μL, 4.43 mmol, 10 eq.). The resulting solution was allowed to stir at room temperature overnight. The product was obtained as an HCl salt by evaporating the solvent under vacuum and was directly used in the next step without purification.

**LC-MS (ESI)**: m/z calcd for C H N O ^+^ [M+H]^+^ 316.2, found 316.2.

For target compounds **29a-29e**, **28** and appropriate carboxylic acids were reacted following General Procedure A. The yields generally ranged from 70% to 90%.

#### N-(2-((((2S,5R)-5-(4-aminopyrrolo[2,1-f][1,2,4]triazin-7-yl)-5-cyanotetrahydrofuran-2-yl)methyl)amino)-2-oxoethyl)-3-(phenylsulfonamido)benzamide *(29a)*

**LC-MS (ESI)**: m/z calcd for C_27_H_27_N_8_O_5_S^+^ [M+H]^+^ 575.2, found 575.3.

**^1^H NMR** (500 MHz, DMSO-*d*_6_ + D_2_O) δ 7.94 (s, 1H), 7.80 – 7.72 (m, 2H), 7.62 – 7.57 (m, 2H), 7.56 – 7.47 (m, 3H), 7.32 (t, *J* = 7.9 Hz, 1H), 7.25 (ddd, *J* = 8.1, 2.4, 1.0 Hz, 1H), 6.89 (d, *J* = 4.5 Hz, 1H), 6.82 (d, *J* = 4.5 Hz, 1H), 4.42 – 4.27 (m, 1H), 3.82 (d, *J* = 1.2 Hz, 2H), 3.35 – 3.24 (m, 2H), 2.74 (ddd, *J* = 13.4, 7.9, 5.9 Hz, 1H), 2.69 – 2.60 (m, 1H), 2.23 (dq, *J* = 12.7, 7.5 Hz, 1H), 1.83 (ddt, *J* = 12.6, 8.4, 6.1 Hz, 1H).

**^13^C NMR** (126 MHz, DMSO) δ 169.7, 166.4, 155.9, 148.4, 135.4, 133.5, 129.8, 129.6, 127.1, 125.0, 123.1, 123.0, 119.9, 119.7, 116.6, 110.0, 101.3, 80.1, 75.2, 42.9, 42.5, 36.8, 28.0.

#### N-(2-((((2S,5R)-5-(4-aminopyrrolo[2,1-f][1,2,4]triazin-7-yl)-5-cyanotetrahydrofuran-2-yl)methyl)amino)-2-oxoethyl)-5-((4-fluorophenyl)sulfonamido)nicotinamide *(29b)*

**LC-MS (ESI)**: m/z calcd for C_26_H_25_FN_9_O_5_S^+^ [M+H]^+^ 594.2, found 594.2.

**^1^H NMR** (500 MHz, DMSO-*d*_6_ + D_2_O) δ 8.73 (d, *J* = 1.9 Hz, 1H), 8.40 (d, *J* = 2.6 Hz, 1H), 7.94 (s, 1H), 7.91 (t, *J* = 2.3 Hz, 1H), 7.86 – 7.82 (m, 2H), 7.43 – 7.38 (m, 2H), 6.89 (d, *J* = 4.5 Hz, 1H), 6.82 (d, *J* = 4.5 Hz, 1H), 4.40 – 4.32 (m, 1H), 3.86 (d, *J* = 2.5 Hz, 2H), 3.33 – 3.29 (m, 2H), 2.74 (ddd, *J* = 13.5, 7.9, 5.9 Hz, 1H), 2.68 – 2.61 (m, 1H), 2.23 (dq, *J* = 12.6, 7.4 Hz, 1H), 1.83 (ddt, *J* = 12.6, 8.4, 6.1 Hz, 1H).

**^13^C NMR** (126 MHz, DMSO) δ 169.5, 169.4, 166.0, 164.8, 163.9, 155.9, 148.4, 144.4, 144.3, 135.8, 134.9, 130.4, 130.3, 130.2, 127.0, 125.0, 124.8, 119.7, 119.4, 117.3, 117.1, 116.6, 110.4, 110.0, 101.3, 80.1, 75.2, 42.9, 42.6, 42.5, 36.8, 28.0.

#### N-(2-((((2S,5R)-5-(4-aminopyrrolo[2,1-f][1,2,4]triazin-7-yl)-5-cyanotetrahydrofuran-2-yl)methyl)amino)-2-oxoethyl)-4-((4-fluorophenyl)sulfonamido)thiophene-2-carboxamide *(29c)*

**LC-MS (ESI)**: m/z calcd for C_25_H_24_FN O S ^+^ [M+H]^+^ 599.1, found 599.3.

**^1^H NMR** (500 MHz, DMSO-*d*_6_ + D_2_O) δ 7.94 (s, 1H), 7.85 – 7.79 (m, 2H), 7.56 (d, *J* = 1.6 Hz, 1H), 7.43 – 7.36 (m, 2H), 7.05 (d, *J* = 1.6 Hz, 1H), 6.90 (d, *J* = 4.5 Hz, 1H), 6.83 (d, *J* = 4.5 Hz, 1H), 4.41 –4.30 (m, 1H), 3.79 (d, *J* = 2.5 Hz, 2H), 3.30 (d, *J* = 5.4 Hz, 2H), 2.74 (ddd, *J* = 13.6, 7.8, 5.9 Hz, 1H), 2.65 (dt, *J* = 13.1, 8.0 Hz, 1H), 2.23 (dq, *J* = 12.7, 7.4 Hz, 1H), 1.83 (ddt, *J* = 12.6, 8.4, 6.1 Hz, 1H).

**^13^C NMR** (126 MHz, DMSO) δ 169.5, 165.8, 163.8, 161.4, 155.9, 148.5, 139.1, 130.2, 130.1, 125.0, 123.9, 119.7, 117.0, 116.8, 116.6, 110.0, 101.3, 80.1, 79.6, 79.4, 79.1, 75.3, 42.6, 42.5, 36.8, 28.0.

#### N-(2-((((2S,5R)-5-(4-aminopyrrolo[2,1-f][1,2,4]triazin-7-yl)-5-cyanotetrahydrofuran-2-yl)methyl)amino)-2-oxoethyl)-5-((4-fluorophenyl)sulfonamido)thiophene-3-carboxamide *(29d)*

**LC-MS (ESI)**: m/z calcd for C_25_H_24_FN O S ^+^ [M+H]^+^ 599.1, found 599.3.

**^1^H NMR** (500 MHz, DMSO-*d*_6_ + D_2_O) δ 8.07 (t, *J* = 6.0 Hz, 1H), 7.94 (s, 1H), 7.84 – 7.74 (m, 2H), 7.66 (s, 1H), 7.46 – 7.34 (m, 2H), 6.94 – 6.91 (m, 1H), 6.90 (d, *J* = 4.5 Hz, 1H), 6.82 (d, *J* = 4.5 Hz, 1H), 4.38 – 4.30 (m, 1H), 3.82 – 3.72 (m, 2H), 3.33 – 3.26 (m, 2H), 2.77 – 2.70 (m, 1H), 2.64 (dt, *J* = 13.0, 7.9 Hz, 1H), 2.22 (dq, *J* = 12.7, 7.4 Hz, 1H), 1.82 (ddt, *J* = 12.6, 8.4, 6.1 Hz, 1H).

**^13^C NMR** (126 MHz, DMSO) δ 169.8, 163.9, 162.4, 155.9, 148.4, 135.4, 130.3, 130.3, 129.6, 127.1, 125.1, 119.7, 117.0, 116.8, 116.6, 110.0, 101.3, 80.1, 75.2, 42.6, 36.8, 28.0.

#### N-(2-((((2S,5R)-5-(4-aminopyrrolo[2,1-f][1,2,4]triazin-7-yl)-5-cyanotetrahydrofuran-2-yl)methyl)amino)-2-oxoethyl)-5-((4-(trifluoromethyl)phenyl)sulfonamido)nicotinamide *(29e)*

**LC-MS (ESI)**: m/z calcd for C_27_H_25_F_3_N_9_O_5_S^+^ [M+H]^+^ 644.2, found 644.3.

**^1^H NMR** (600 MHz, DMSO-*d*_6_ + D_2_O) δ 8.74 (d, *J* = 2.0 Hz, 1H), 8.41 (d, *J* = 2.6 Hz, 1H), 8.14 (t, *J* = 6.0 Hz, 1H), 8.02 – 7.91 (m, 7H), 6.89 (d, *J* = 4.4 Hz, 1H), 6.81 (d, *J* = 4.5 Hz, 1H), 4.38 – 4.33 (m, 1H), 3.91 – 3.81 (m, 2H), 3.45 – 3.37 (m, 2H), 3.31 (overlapped with water peak, 1H), 2.77 – 2.71 (m, 1H), 2.64 (dt, *J* = 13.1, 8.0 Hz, 1H), 2.23 (dq, *J* = 12.7, 7.4 Hz, 1H), 1.84 (ddt, *J* = 12.6, 8.5, 6.2 Hz, 1H).

**^13^C NMR** (101 MHz, DMSO) δ 169.5, 169.4, 164.8, 155.9, 148.5, 144.6, 130.4, 128.1, 127.2, 127.2, 125.1, 125.0, 122.4, 119.7, 116.6, 111.5, 109.9, 102.2, 101.3, 86.7, 80.1, 75.2, 63.0, 36.8, 28.0.

### (2R,5S)-2-(4-aminopyrrolo[2,1-f][1,2,4]triazin-7-yl)-5-((methylamino)methyl)tetrahydrofuran-2-carbonitrile *(30)*

In a microwave vial, 24 (45 mg, 0.109 mmol) was dissolved with 2 M NH_2_Me in THF (2.74 mL, 5.48 mmol, 50 eq.). The reaction mixture was subjected to microwave irradiation at 100 for 4 hours. The reaction mixture was evaporated to dryness and the crude was purified by column chromatography (DCM: MeOH 40:1 to 10:1 with 1% TEA) to afford the product as a white solid (15 mg, yield: 51%).

**LC-MS (ESI)**: m/z calcd for C_13_H_17_N_6_O^+^ [M+H]^+^ 273.2, found 273.2.

### tert-butyl

## (2-((((2S,5R)-5-(4-aminopyrrolo[2,1-f][1,2,4]triazin-7-yl)-5-cyanotetrahydrofuran-2-yl)methyl)(methy l)amino)-2-oxoethyl)carbamate *(31)*

General Procedure A was followed. **30** was reacted with Fmoc-Gly-OH to yield the product as a white powder (yield: 87%).

**LC-MS (ESI)**: m/z calcd for C_30_H_30_N_7_O_4_^+^ [M+H]^+^ 552.2, found 552.4.

### 2-amino-N-(((2S,5R)-5-(4-aminopyrrolo[2,1-f][1,2,4]triazin-7-yl)-5-cyanotetrahydrofuran-2-yl)methyl)-N-methylacetamide *(32)*

**31** (60 mg, 0.11 mmol, 1 eq.) was dissolved with DCM (1 mL). A 2 M solution of dimethylamine in THF (0.55 mL, 10 eq.) was added and the reaction mixture was allowed to stir at room temperature overnight. The solvent was evaporated and the mixture was purified by column chromatography (DCM:MeOH 50:1 to 10:1 with 1% TEA) to afford the product as a white solid (26 mg, yield: 79%).

**LC-MS (ESI)**: m/z calcd for C H N O ^+^ [M+H]^+^ 330.2, found 330.2.

#### N-(2-((((2S,5R)-5-(4-aminopyrrolo[2,1-f][1,2,4]triazin-7-yl)-5-cyanotetrahydrofuran-2-yl)methyl)(met hyl)amino)-2-oxoethyl)-5-((4-(trifluoromethyl)phenyl)sulfonamido)nicotinamide *(33)*

32 and 5-((4-(trifluoromethyl)phenyl)sulfonamido)nicotinic acid were reacted following General Procedure A to yield the target compound.

**LC-MS (ESI)**: m/z calcd for C_28_H_27_F_3_N_9_O_5_S^+^ [M+H]^+^ 658.2, found 658.4.

**^1^H NMR** (400 MHz, DMSO-*d*_6_+ D_2_O) δ 8.86 – 8.78 (m, 1H), 8.72 – 8.78 (m, 1H), 8.42 (d, *J* = 2.5 Hz, 1H), 8.09 – 7.97 (m, 5H), 7.93 (q, *J* = 2.6 Hz, 1H), 6.97 (dd, *J* = 4.5, 2.2 Hz, 1H), 6.89 (t, *J* = 4.1 Hz, 1H), 4.68 – 4.58 (m, 0.35H), 4.53 (td, *J* = 6.9, 4.6 Hz, 0.65H), 4.32 – 4.04 (m, 2H), 3.67 – 3.63 (m, 1H), 3.49 (overlapped with water peak, 1H), 3.07 (s, 2H), 2.93 (s, 1H), 2.91 – 2.68 (m, 2H), 2.46 – 2.36 (m, 0.35H), 2.29 (dq, *J* = 14.1, 7.3 Hz, 0.65H), 1.91 (tdd, *J* = 15.0, 12.6, 6.0 Hz, 1H).

**^13^C NMR** (101 MHz, DMSO) δ 169.0, 168.8, 165.1, 155.9, 148.5, 144.8, 143.0, 130.5, 128.0, 127.0, 126.9, 125.2, 125.2, 124.8, 122.5, 119.7, 116.6, 109.8, 109.7, 101.3, 80.0, 79.7, 75.4, 75.1, 51.3, 41.4, 35.9, 34.6, 28.2.

## Acknowledgement

This work made use of the Cornell University NMR Facility, which is supported, in part, by the NSF through MRI award CHE-1531632. This work is based on research conducted at the Center for High-Energy X-ray Sciences (CHEXS), which is supported by the National Science Foundation (BIO, ENG and MPS Directorates) under award DMR-1829070, and the Macromolecular Diffraction at CHESS (MacCHESS) facility, which is supported by award 1-P30-GM124166-01A1 from the National Institute of General Medical Sciences, National Institutes of Health, and by New York State’s Empire State Development Corporation (NYSTAR).

