## Supplementary Tables and Figures for "Development of GS-441524 Derivatives as Potent SARS-CoV-2 Mac1 Inhibitors via a Direct-to-Biology Approach"

**Table 1. Data collection and refinement statistics**

(values for the highest-resolution shell are shown in parentheses)

| | |
|---|---|
| **Wavelength (Å)** | 0.9686 |
| **Resolution range (Å)** | 29.07 - 1.526 (1.56 - 1.53) |
| **Space group** | P 1 $2_1$ 1 |
| **Unit cell** | 42.91Å 88.90Å 43.04 Å<br>90.00° 94.88° 90.00° |
| **Total reflections** | 289424 (9284) |
| **Unique reflections** | 46501 (2249) |
| **Multiplicity** | 6.2 (4.1) |
| **Completeness (%)** | 95.21 (65.94) |
| **Mean I/sigma(I)** | 11.30 (1.77) |
| **Wilson B-factor** | 17.00 |
| **R-merge** | 0.08087 (0.4605) |
| **R-meas** | 0.08792 (0.5264) |
| **R-pim** | 0.03408 (0.2452) |
| **CC1/2** | 0.998 (0.715) |
| **CC*** | 0.999 (0.913) |
| **Reflections used in refinement** | 46498 (2284) |
| **Reflections used for R-free** | 2007 (86) |
| **R-work** | 0.1603 (0.2483) |
| **R-free** | 0.1887 (0.2431) |
| **Number of non-hydrogen atoms** | 2940 |
| **Protein residues** | 335 |
| **RMS(bonds)** | 0.005 |
| **RMS(angles)** | 0.75 |
| **Ramachandran favored (%)** | 99.40 |
| **Ramachandran allowed (%)** | 0.60 |
| **Ramachandran outliers (%)** | 0.00 |
| **Rotamer outliers (%)** | 0.00 |
| **Clashscore** | 1.70 |
| **Average B-factor** | 20.63 |

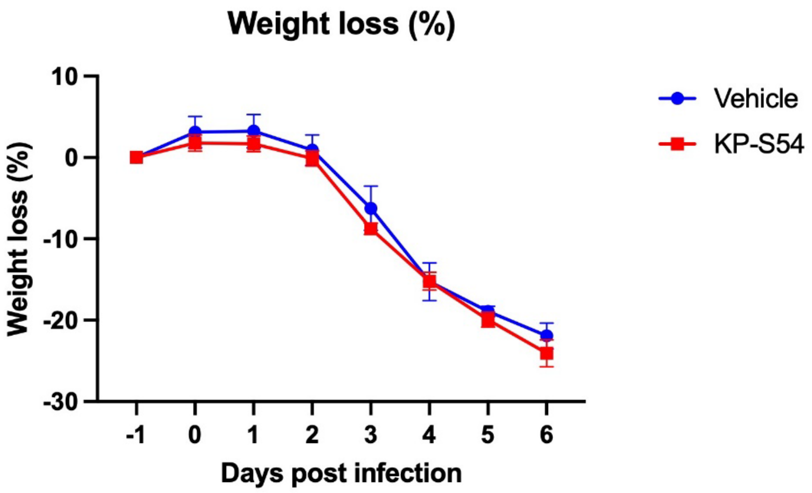


**Figure S1.** Effects of KP-S54 at 50 mg/kg and the vehicle on mouse body weight after SARS-CoV-2 infection (n = 5).

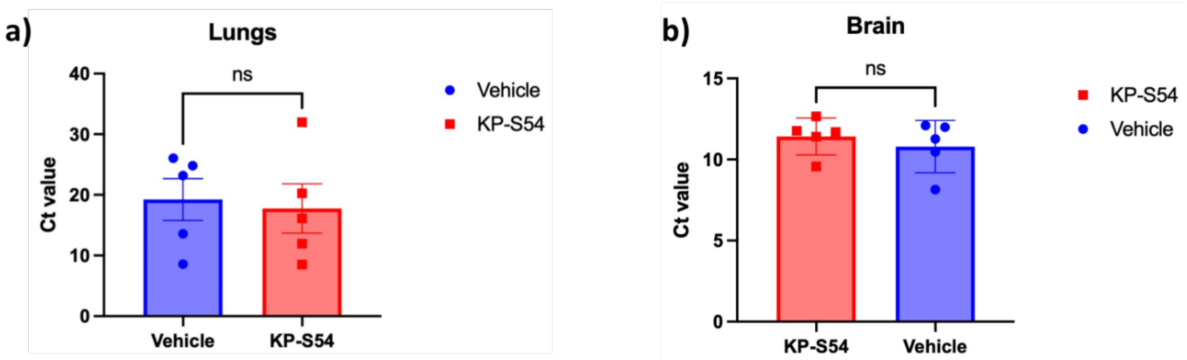


**Figure S2.** Effects of KP-S54 and the vehicle on SARS-CoV-2 viral loads in mouse **(a)** lungs and **(b)** brains (n = 5).

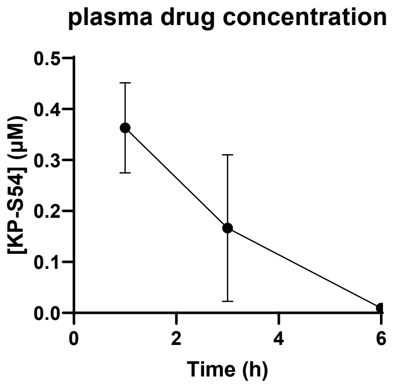


**Figure S3.** KP-S54 concentration in mouse plasma over time following intraperitoneal injection at 50 mg/kg (n = 2).

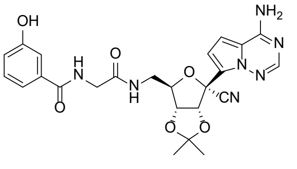


**7**

KP106CR74.10.fid

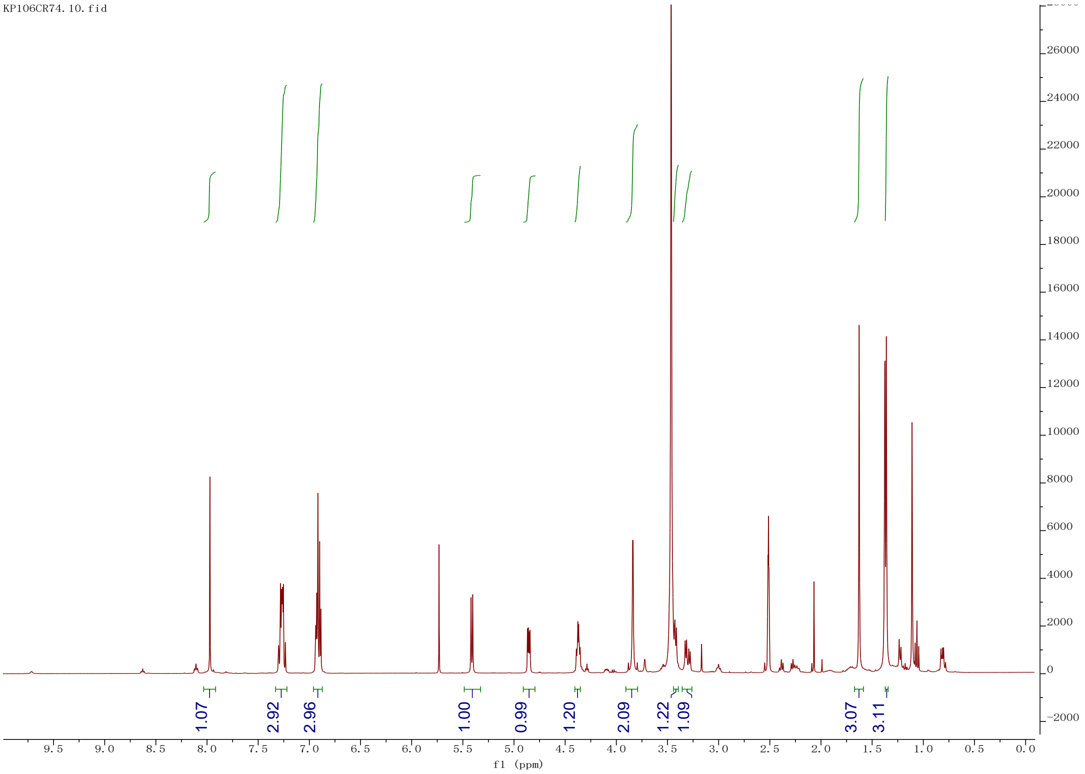


KP106CR74.12.fid

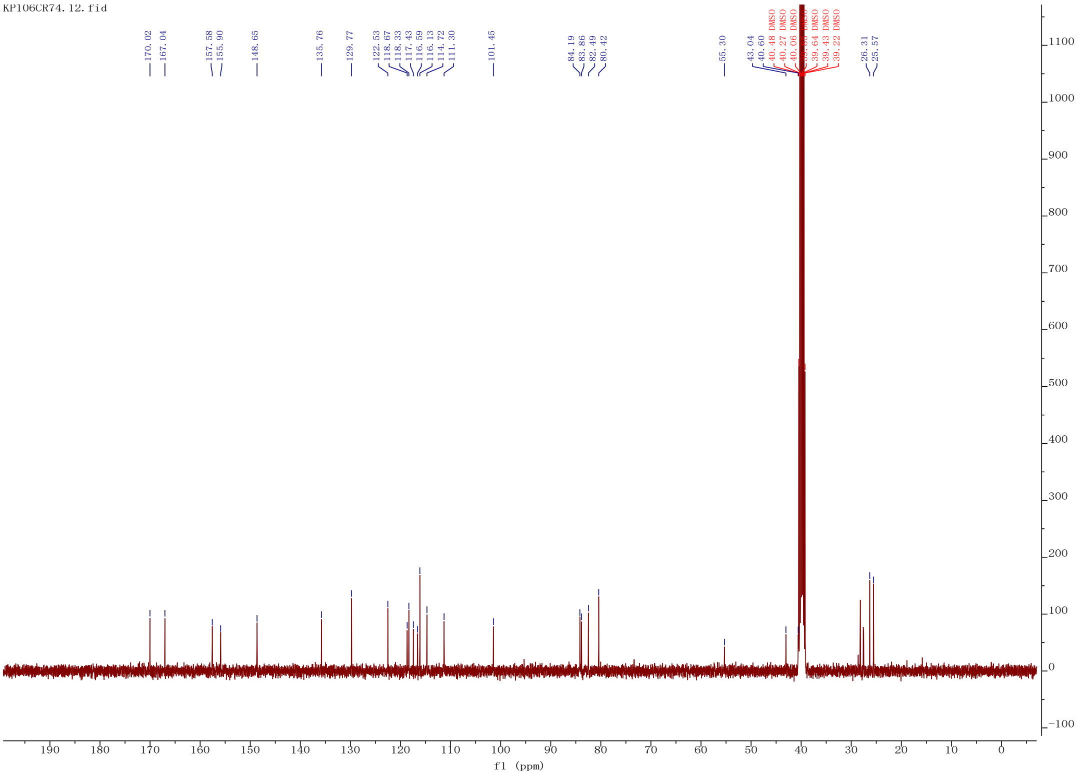

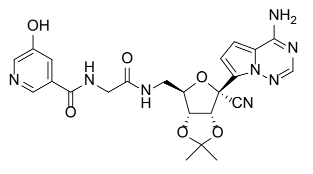
OH
N
H
N
O
N
H
O
O
N
N
NH2
N
CN
O
O

**12a**

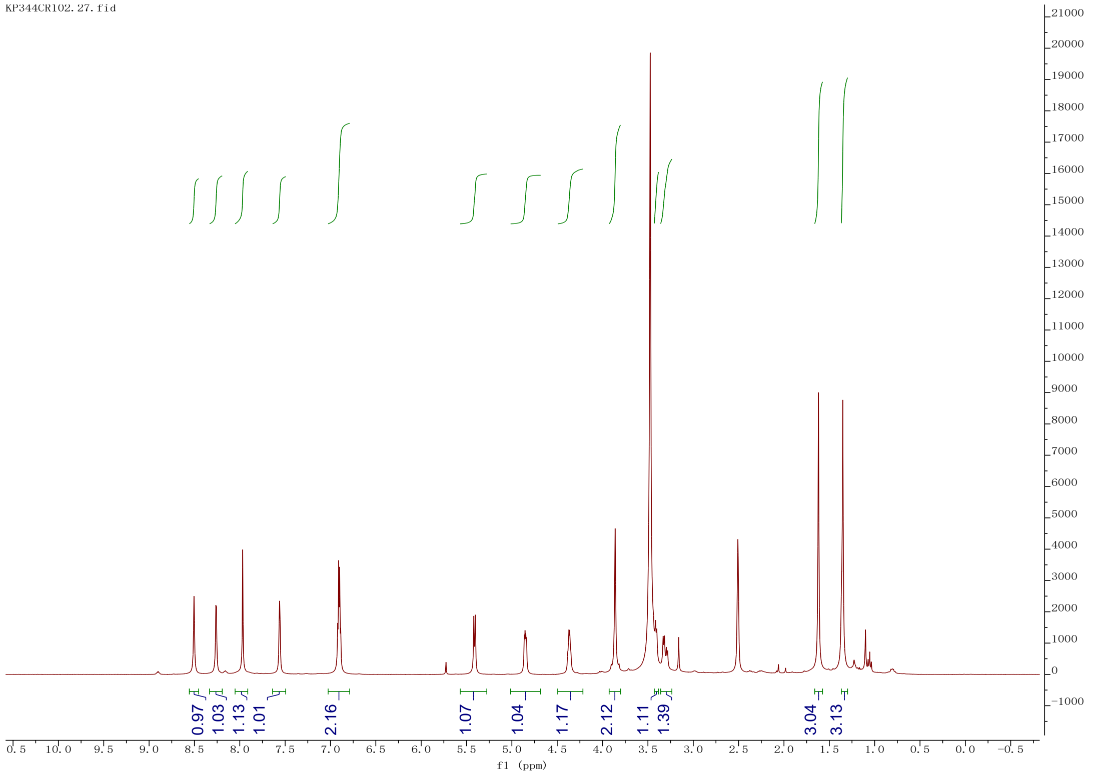
KP344CR102.27.fid
0.97
1.03
1.13
1.01
2.16
1.07
1.04
1.17
2.12
1.11
1.39
3.04
3.13
f1 (ppm)

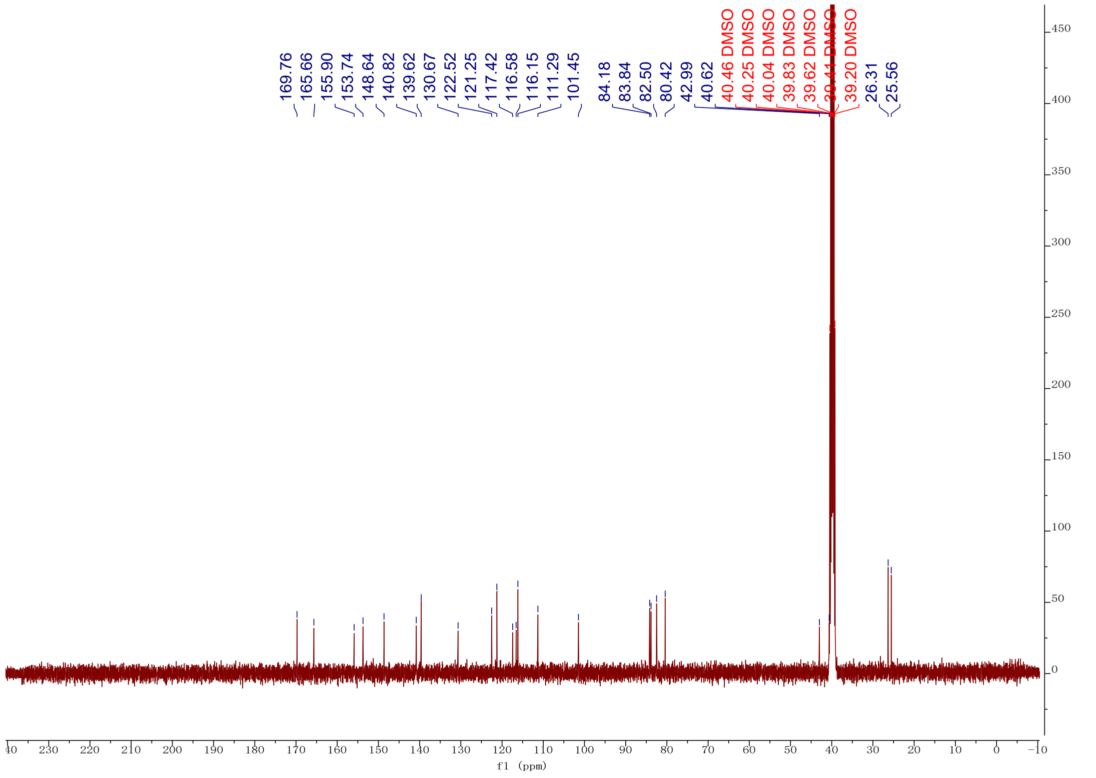
169.76
165.66
155.90
153.74
148.64
140.82
139.62
130.67
122.52
121.25
117.42
116.58
116.15
111.29
101.45
84.18
83.84
82.50
80.42
42.99
40.62
40.46 DMSO
40.25 DMSO
40.04 DMSO
39.83 DMSO
39.62 DMSO
39.41 DMSO
39.20 DMSO
26.31
25.56
f1 (ppm)

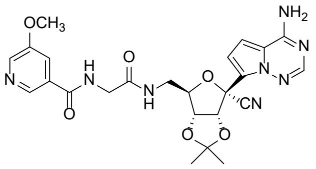
OCH3
NH2
N
H
O
N
O
N
H
O
O
O
N
N
N
CN

**12b**

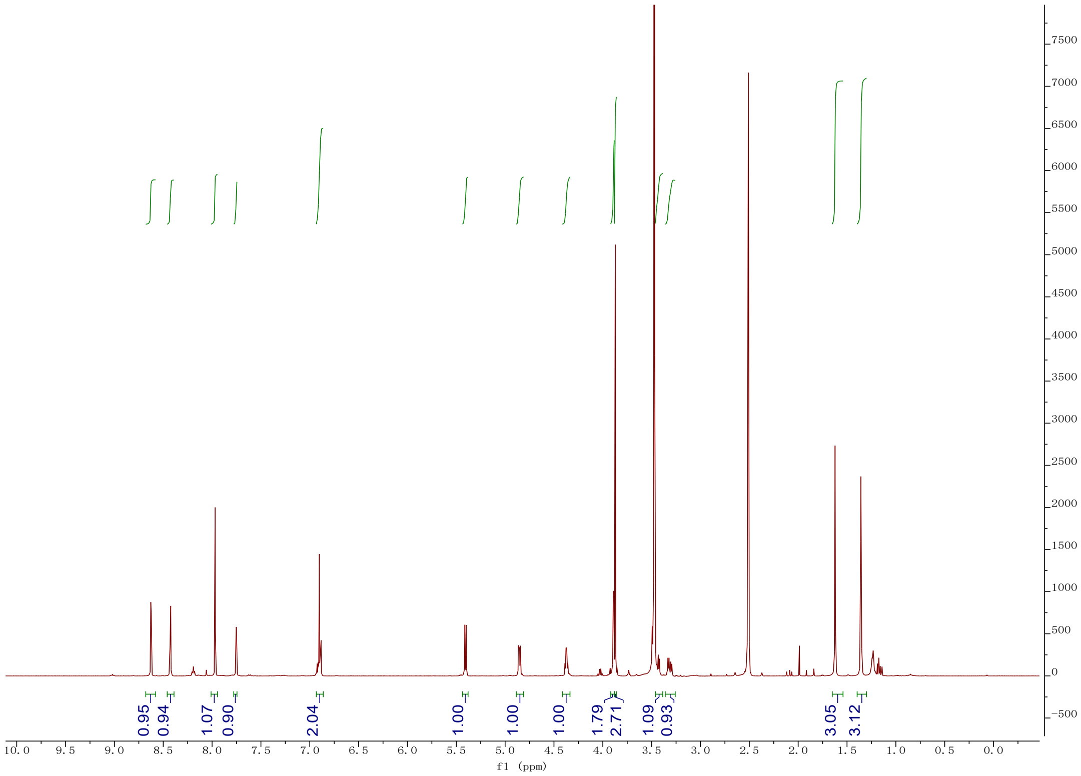
0.95
0.94
1.07
0.90
2.04
1.00
1.00
1.00
1.79
2.71
1.09
0.93
3.05
3.12
f1 (ppm)

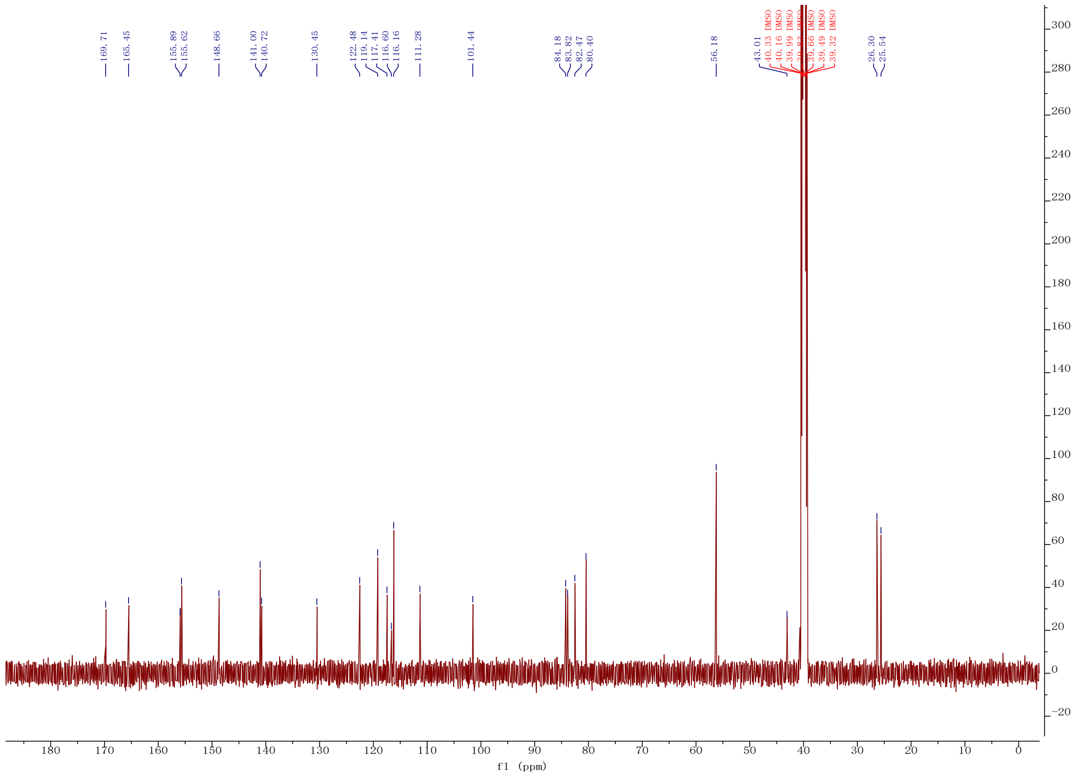
169.71
165.45
155.89
155.62
148.66
141.00
140.72
130.45
122.48
119.14
117.41
116.60
116.16
111.28
101.44
84.18
83.82
82.47
80.40
56.18
43.01
40.33 DMSO
40.16 DMSO
39.99 DMSO
39.66 DMSO
39.49 DMSO
39.32 DMSO
26.30
25.54
f1 (ppm)

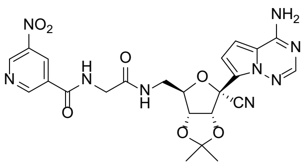

**12c**

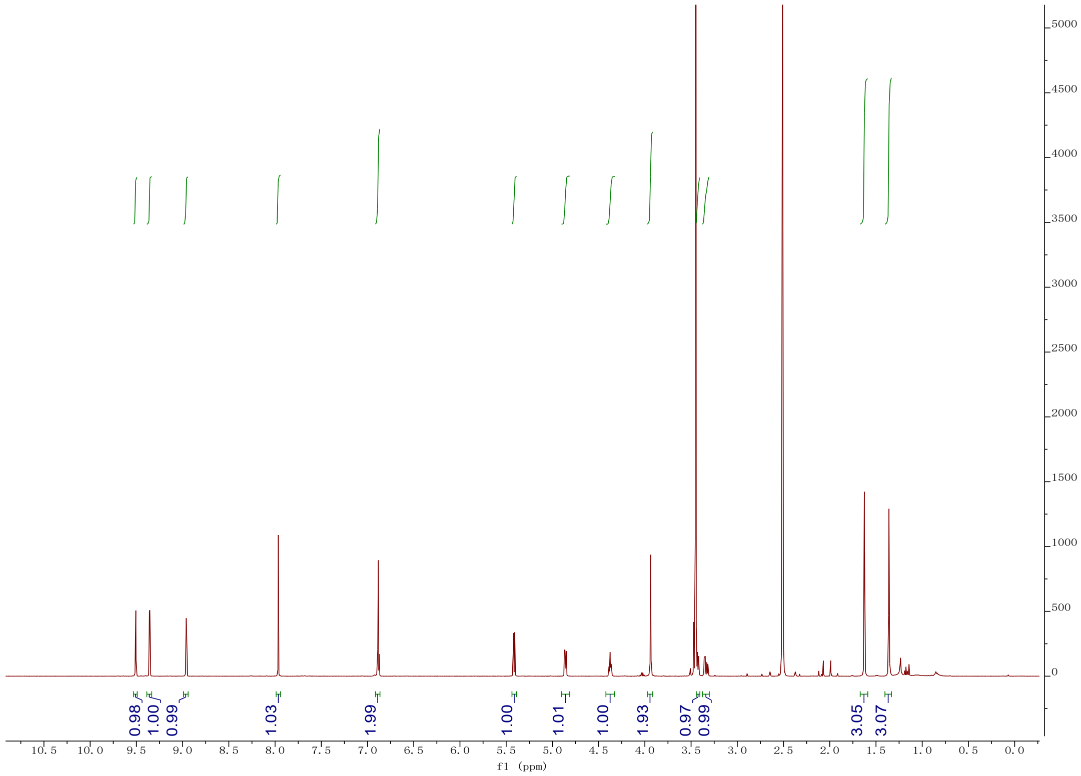

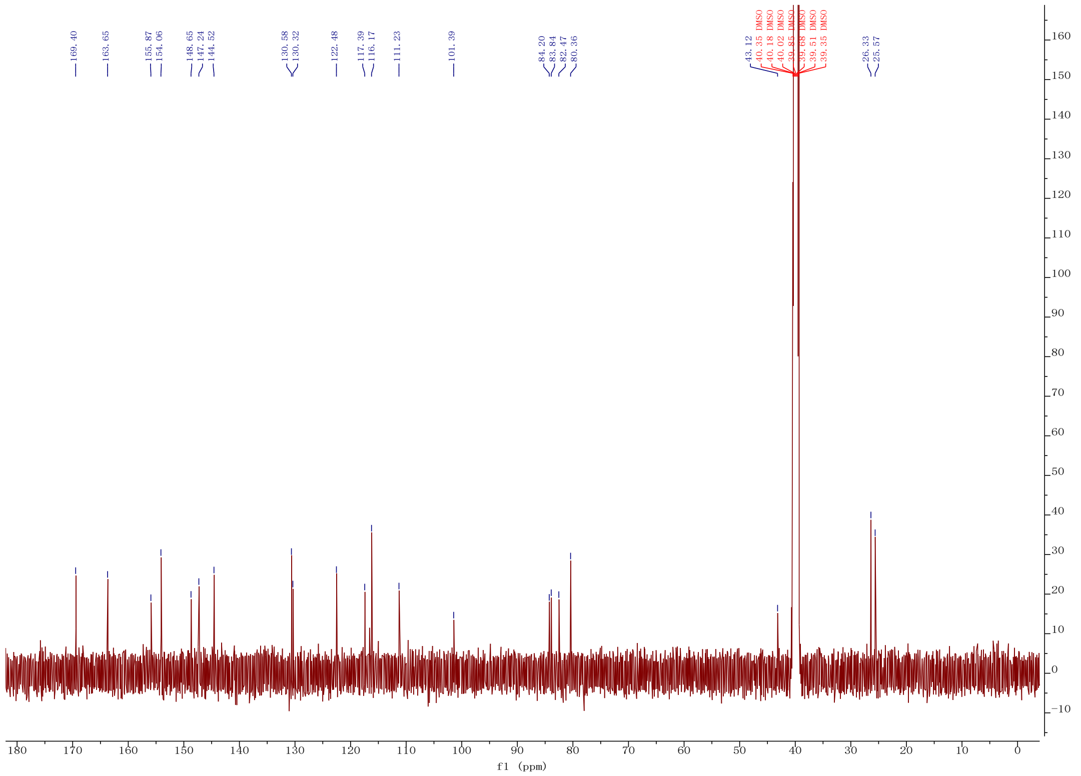

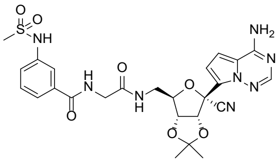

**12d**

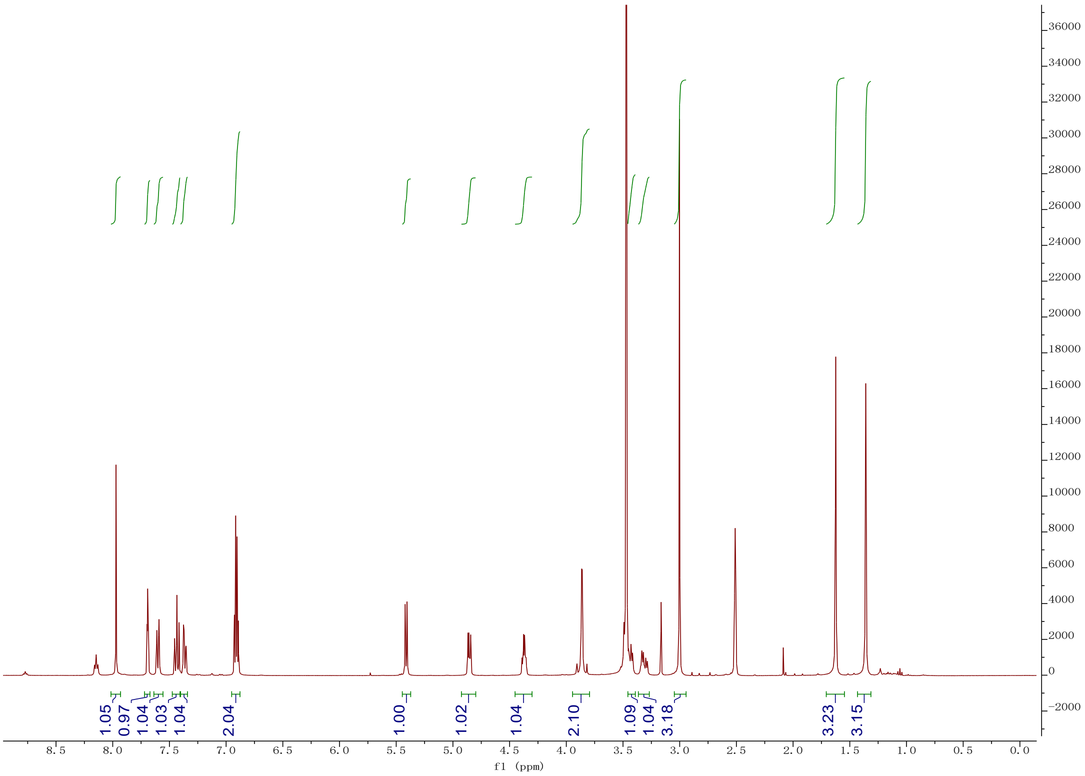

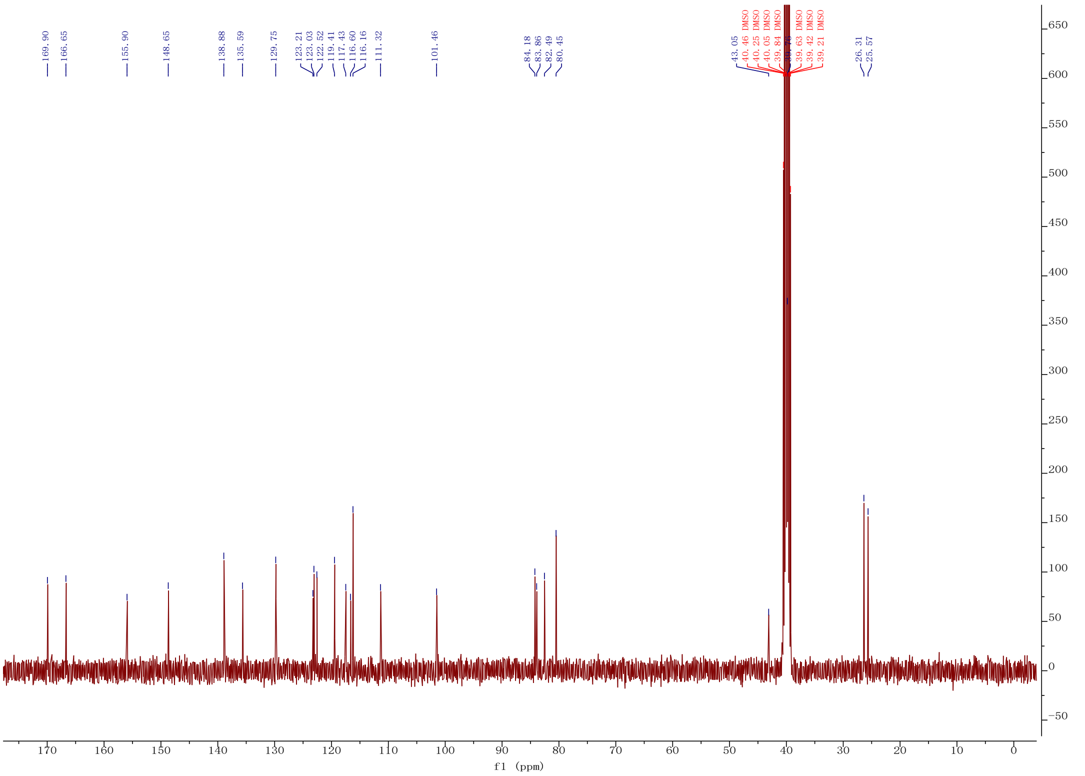

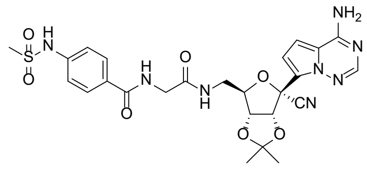

**12e**

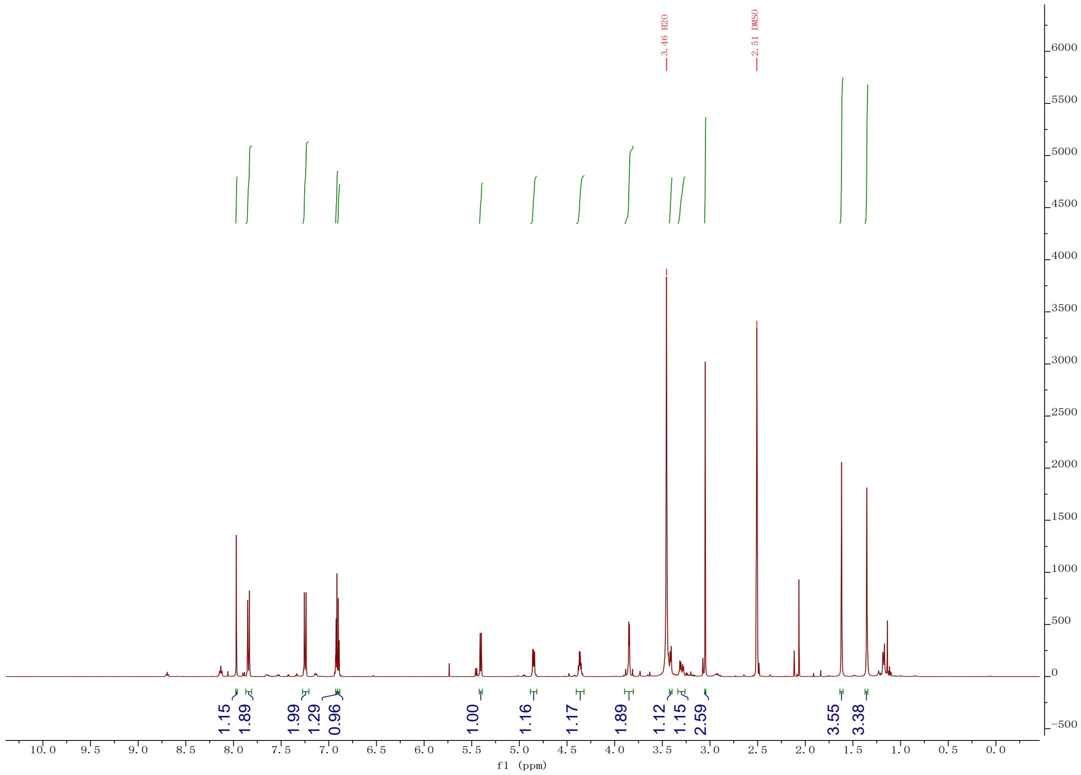
3.46 H2O
2.51 DMSO
1.15
1.89
1.99
1.29
0.96
1.00
1.16
1.17
1.89
1.12
1.15
2.59
3.55
3.38
f1 (ppm)

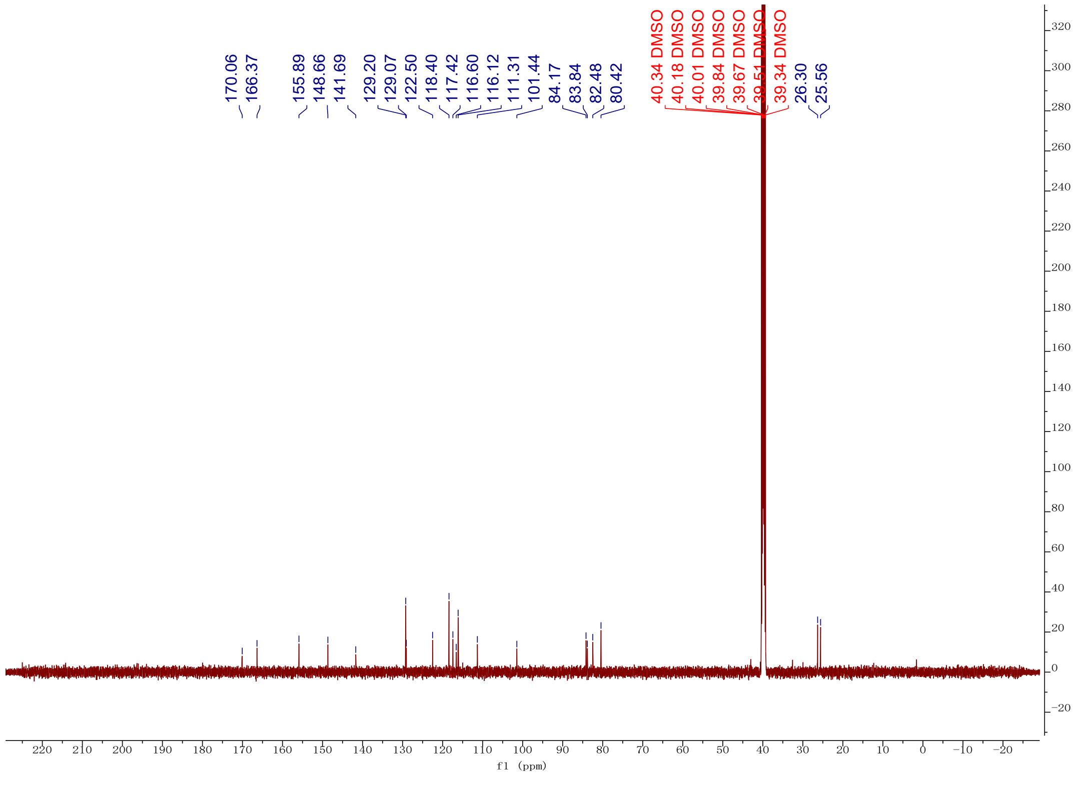
170.06
166.37
155.89
148.66
141.69
129.20
129.07
122.50
118.40
117.42
116.60
116.12
111.31
101.44
84.17
83.84
82.48
80.42
40.34 DMSO
40.18 DMSO
40.01 DMSO
39.84 DMSO
39.67 DMSO
39.51 DMSO
39.34 DMSO
26.30
25.56
f1 (ppm)

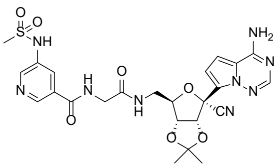

NH2
NH
N
O
H
N
O
N
H
O
N
N
N
CN
O
O


**12f**

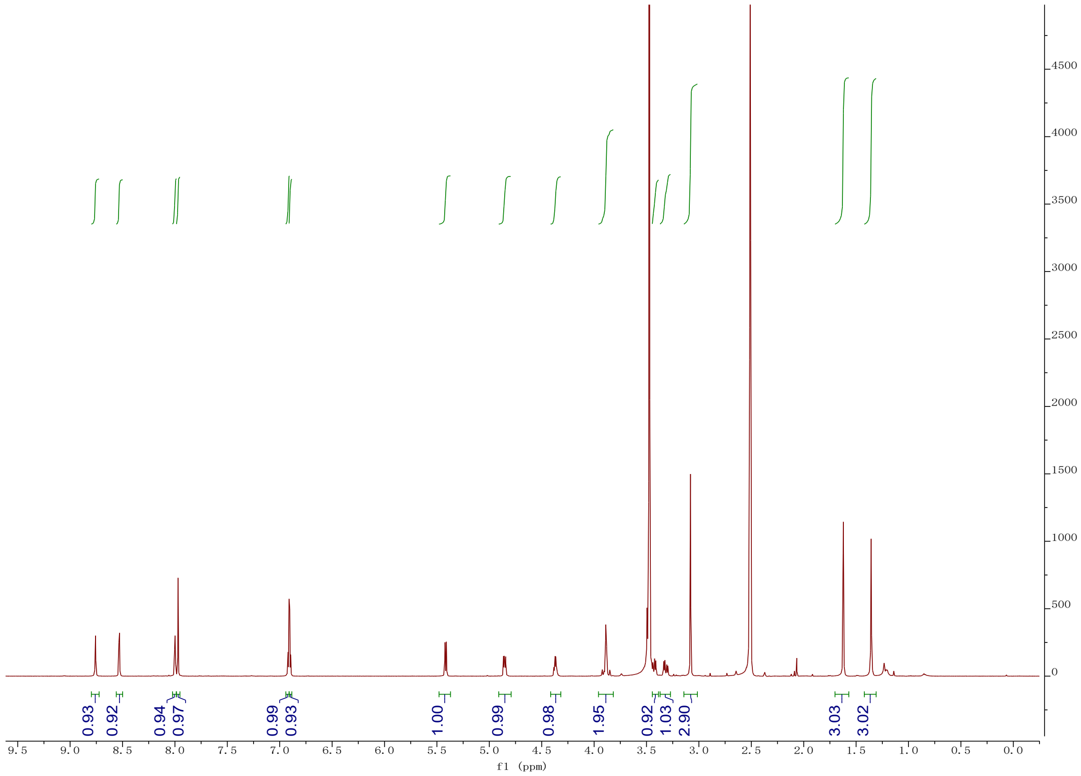

0.93
0.92
0.94
0.97
0.99
0.93
1.00
0.99
0.98
1.95
0.92
1.03
2.90
3.03
3.02
f1 (ppm)


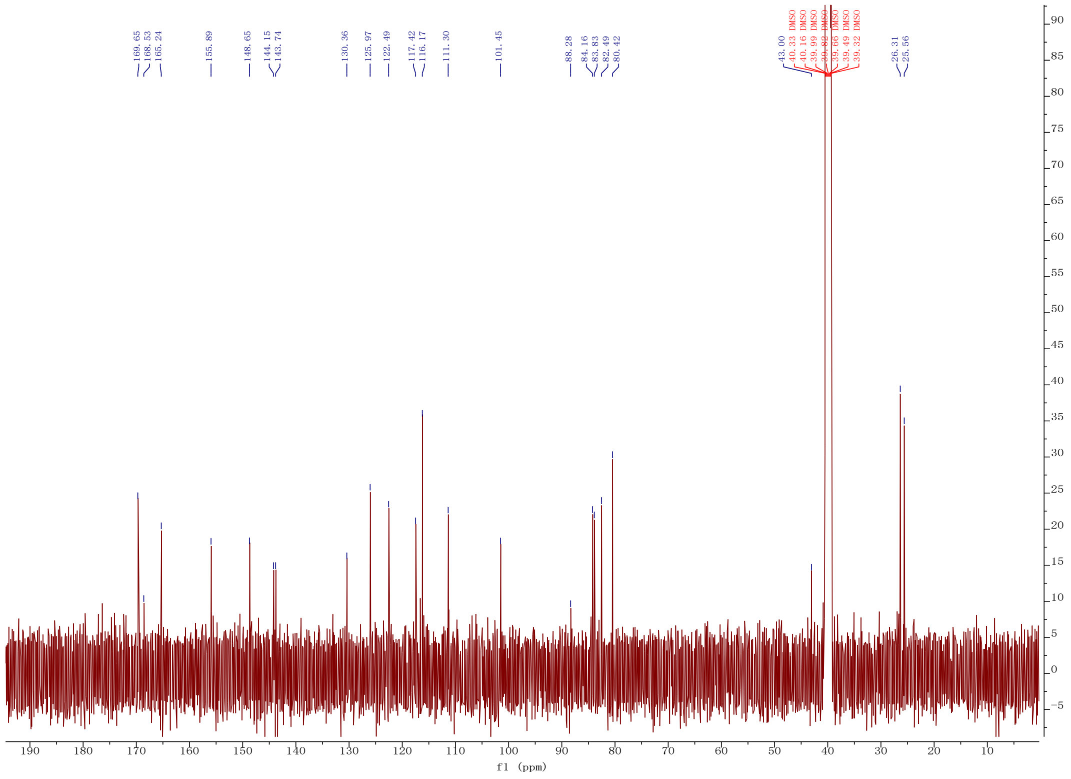

169.65
168.53
165.24
155.89
148.65
144.15
143.74
130.36
125.97
122.49
117.42
116.17
111.30
101.45
88.28
84.16
83.83
82.49
80.42
43.00
40.33 DMSO
40.16 DMSO
39.99 DMSO
39.82 DMSO
39.66 DMSO
39.49 DMSO
39.32 DMSO
26.31
25.56
f1 (ppm)

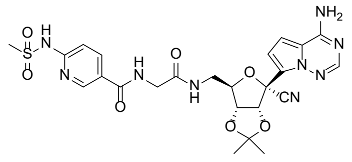

NH2
N
N
N
O
H
N
S
O
N
H
N
O
N
H
O
O
CN
O
O


**12g**

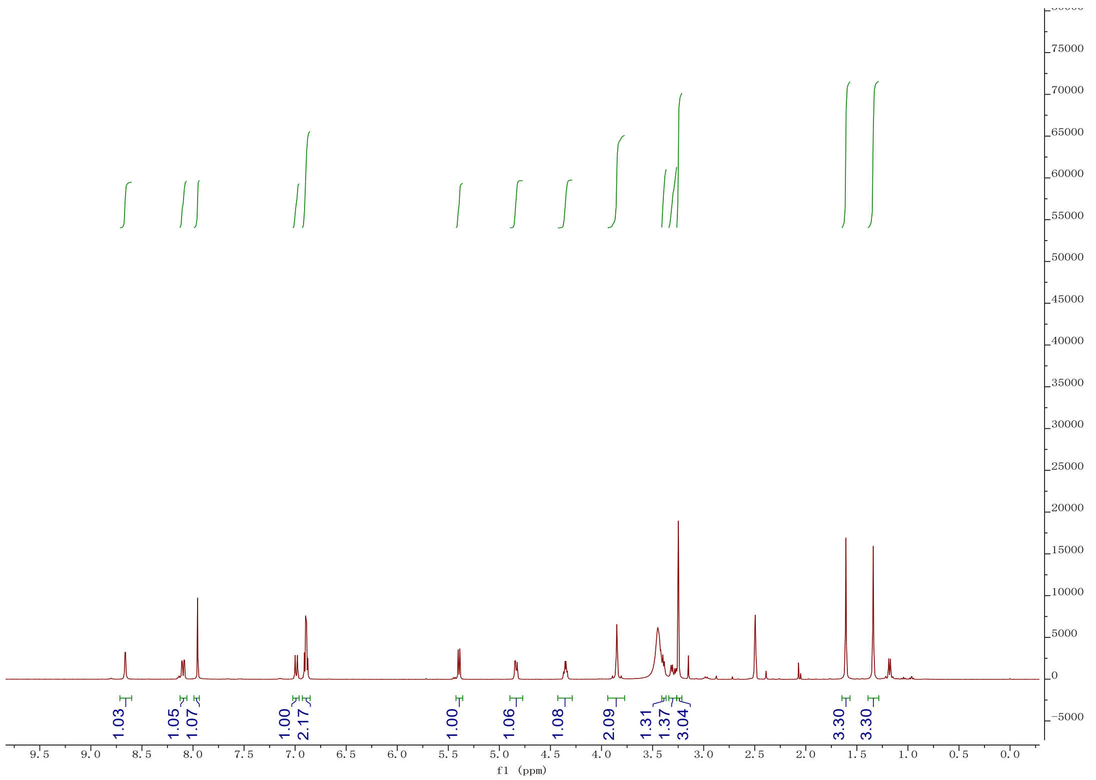

1.03
1.05
1.07
1.00
2.17
1.00
1.06
1.08
2.09
1.31
1.37
3.04
3.30
3.30
f1 (ppm)


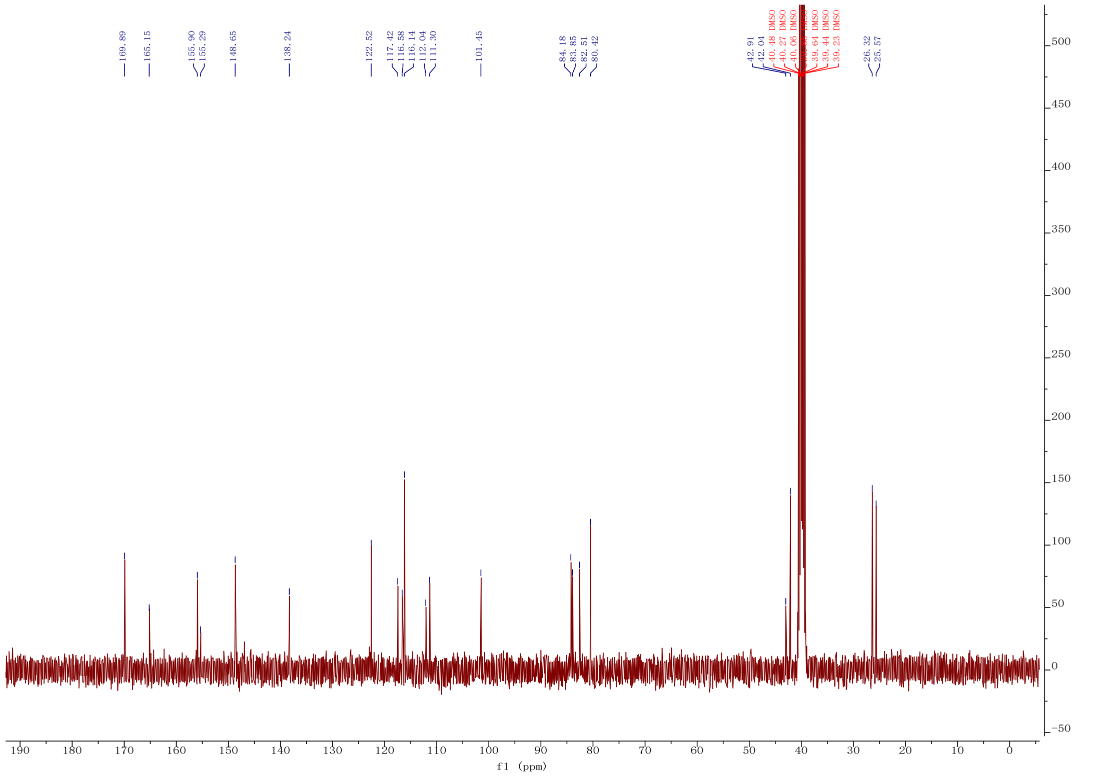

169.89
165.15
155.90
155.29
148.65
138.24
122.52
117.42
116.58
116.14
112.04
111.30
101.45
84.18
83.85
82.51
80.42
42.91
42.04
40.48 DMSO
40.27 DMSO
40.06 DMSO
39.64 DMSO
39.44 DMSO
39.23 DMSO
26.32
25.57
f1 (ppm)

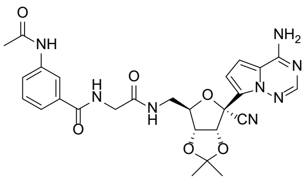

**12h**

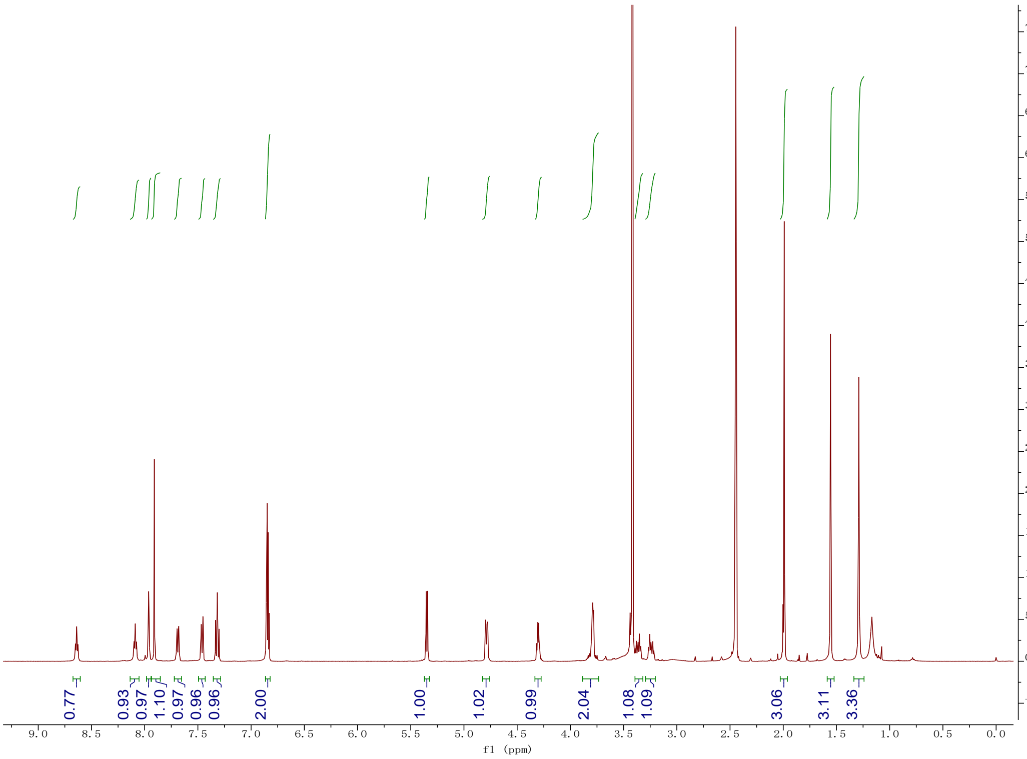
0.77
0.93
0.97
1.10
0.97
0.96
0.96
2.00
1.00
1.02
0.99
2.04
1.08
1.09
3.06
3.11
3.36
f1 (ppm)

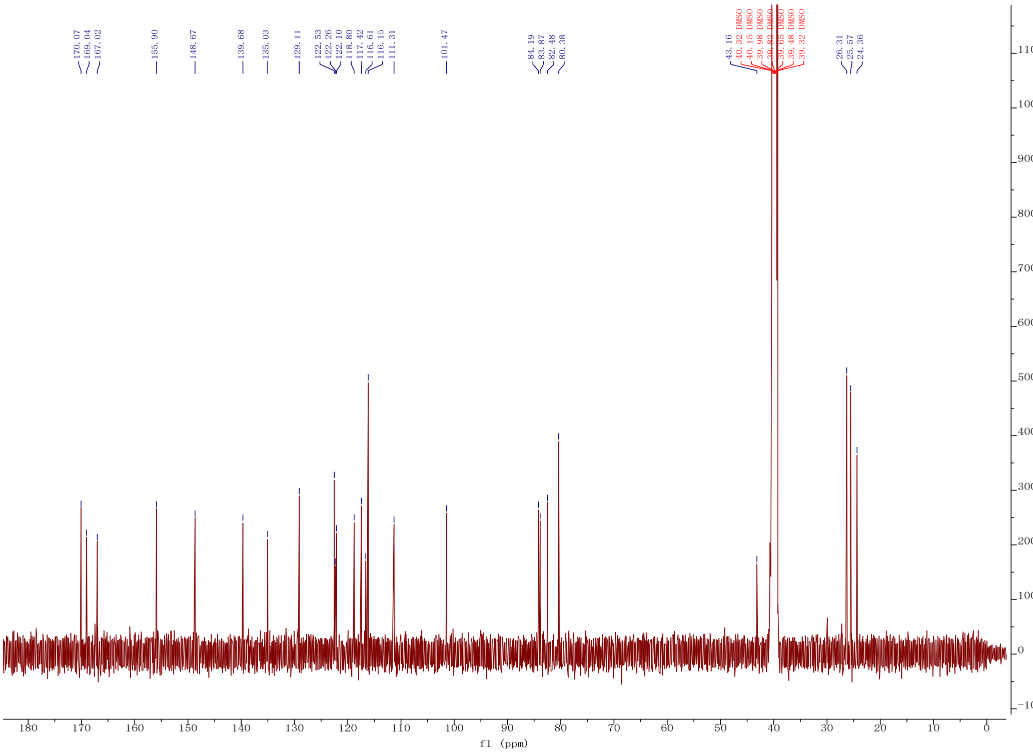
170.07
169.04
167.02
155.90
148.67
139.68
135.03
129.11
122.53
122.26
122.10
118.80
117.42
116.61
116.15
111.31
101.47
84.19
83.87
82.48
80.38
43.16
40.32 DMSO
40.15 DMSO
39.98 DMSO
39.82 DMSO
39.65 DMSO
39.48 DMSO
39.32 DMSO
26.31
25.57
24.36
f1 (ppm)

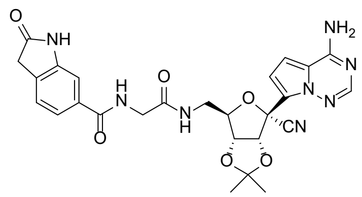

**12i**

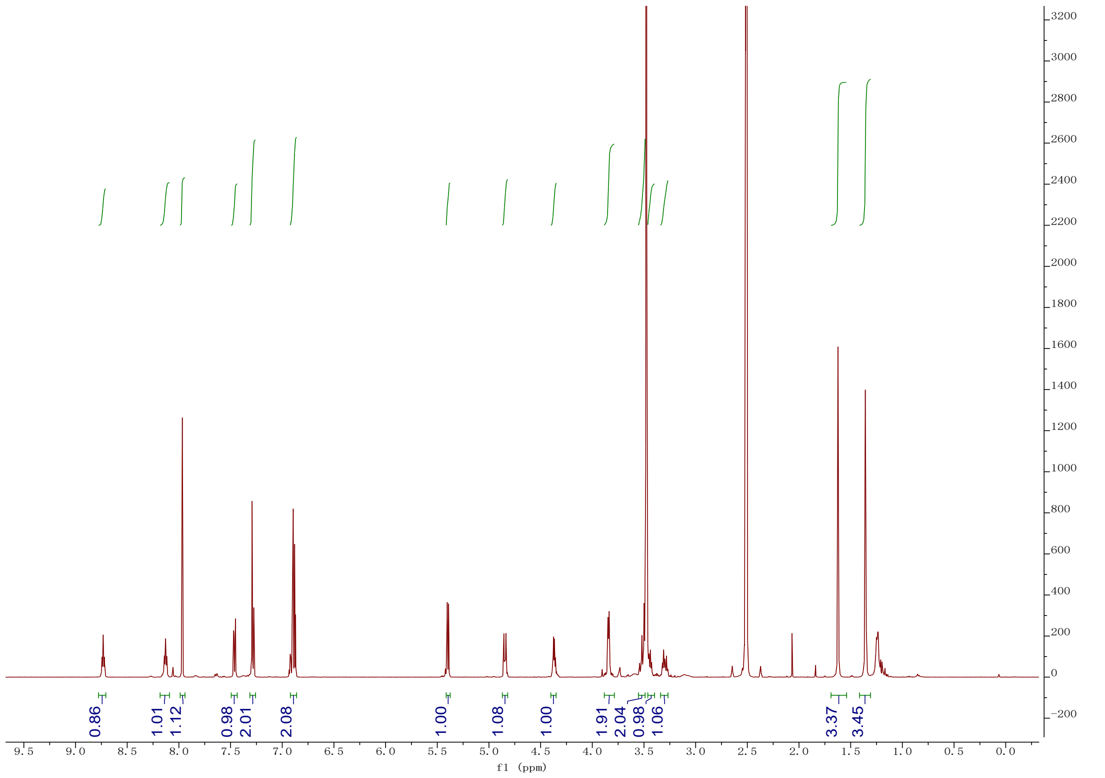

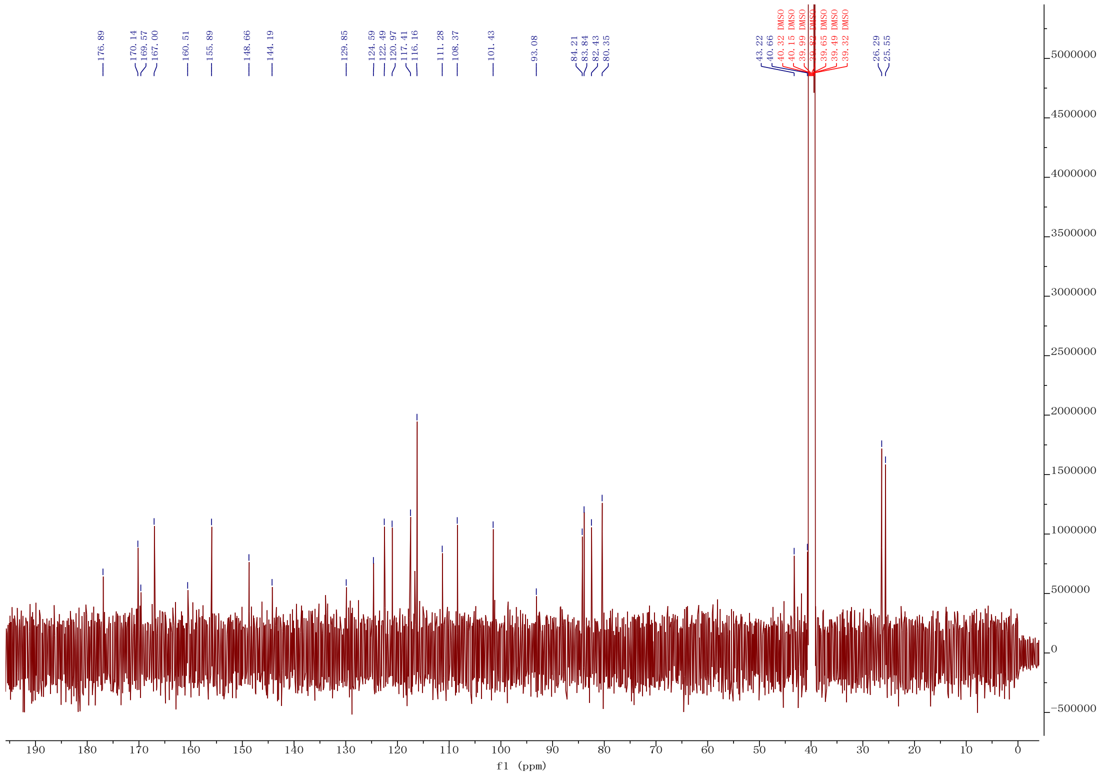

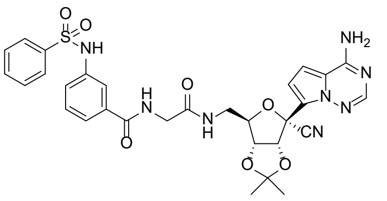
NH2
N
N
N
O
CN
O
O
NH
S
O
O
H
N
O
N
H
O

**12j**

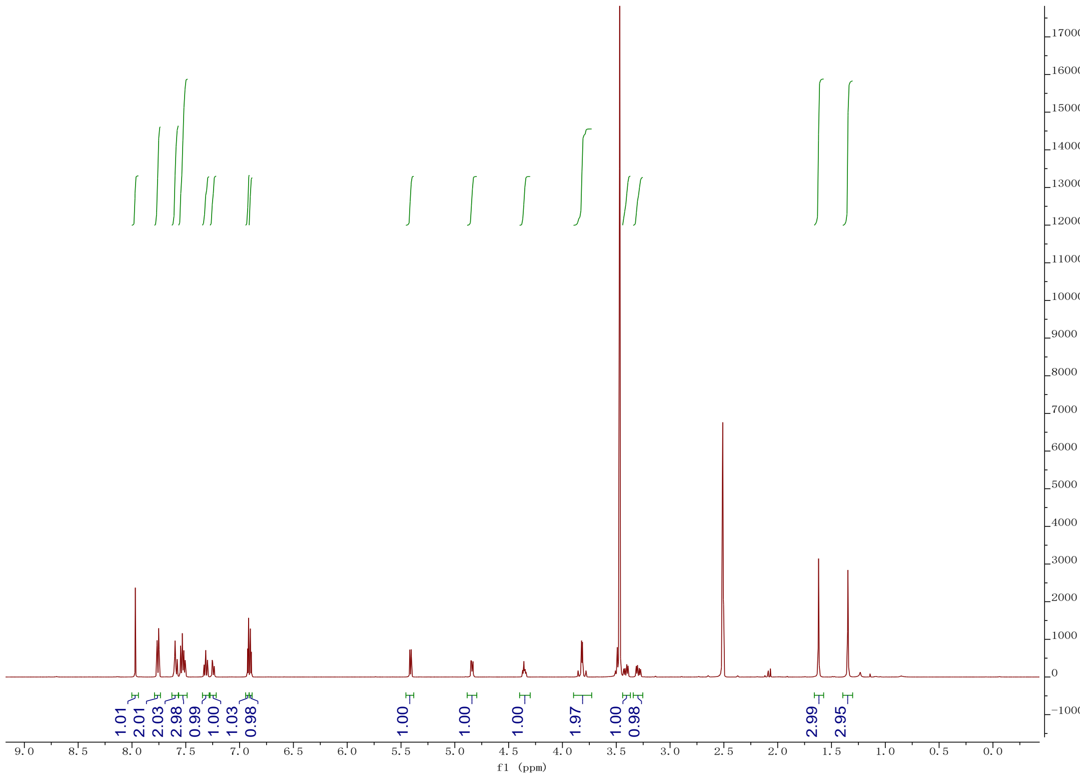
1.01
2.01
2.03
2.98
0.99
1.00
1.03
0.98
1.00
1.00
1.00
1.97
1.00
0.98
2.99
2.95
f1 (ppm)

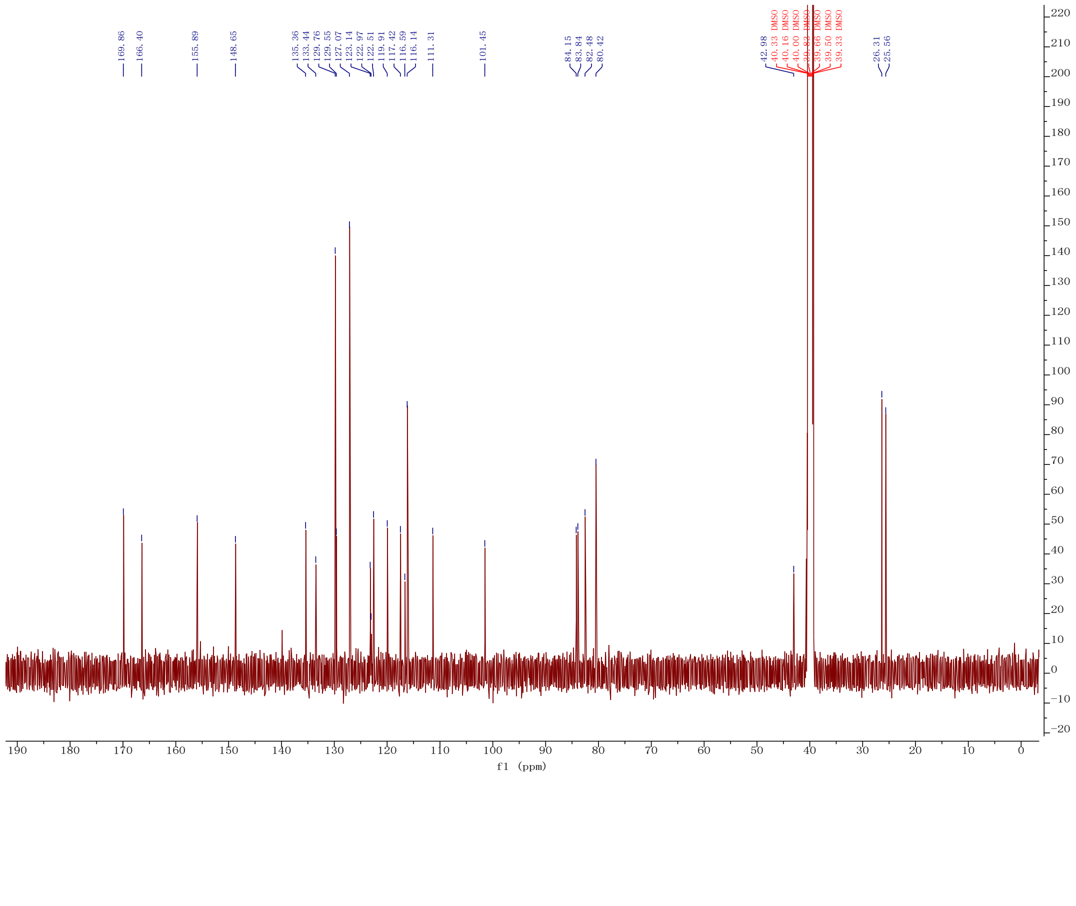
169.86
166.40
155.89
148.65
135.36
133.44
129.76
129.55
127.07
123.14
122.97
122.51
119.91
117.42
116.59
116.14
111.31
101.45
84.15
83.84
82.48
80.42
42.98
40.33 DMSO
40.16 DMSO
40.00 DMSO
39.50 DMSO
39.33 DMSO
26.31
25.56
f1 (ppm)

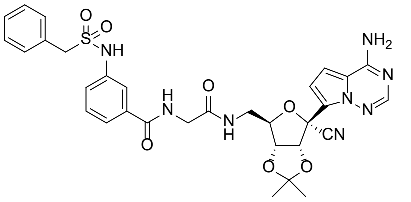
NH2
NH
O
O
S
H
N
O
N
H
O
O
N
N
N
CN
O
O

**12k**

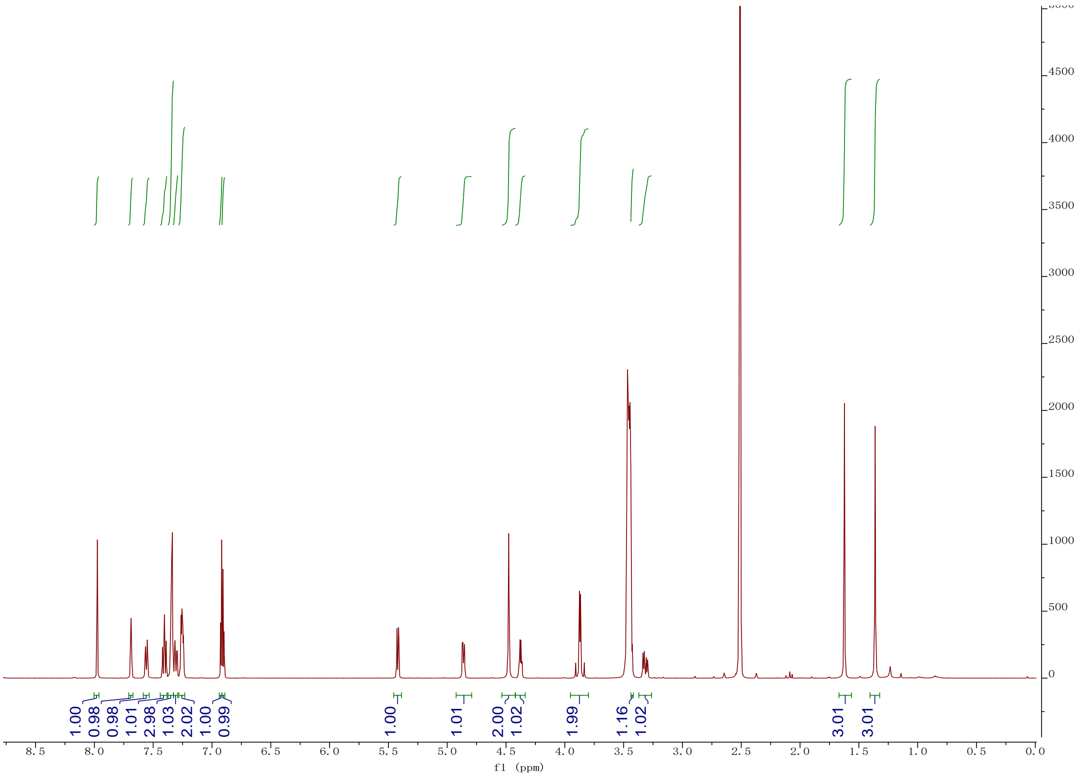
1.00
0.98
0.98
1.01
2.98
1.03
2.02
1.00
0.99
1.00
1.01
2.00
1.02
1.99
1.16
1.02
3.01
3.01
f1 (ppm)

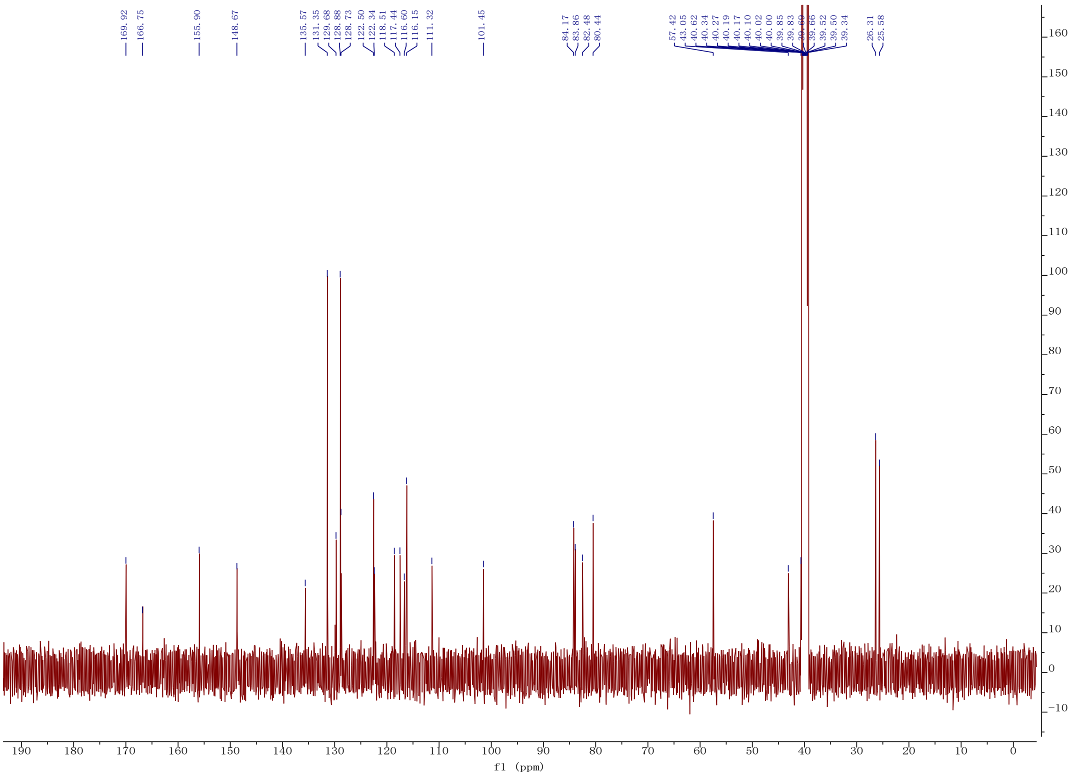
169.92
166.75
155.90
148.67
135.57
131.35
129.68
128.88
128.73
122.50
122.34
118.51
117.44
116.60
116.15
111.32
101.45
84.17
83.86
82.48
80.44
57.42
43.05
40.62
40.34
40.27
40.19
40.17
40.10
40.02
40.00
39.85
39.83
39.52
39.50
39.34
26.31
25.58
f1 (ppm)

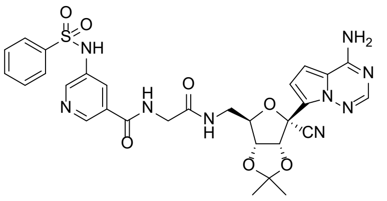
NH2
N
N
N
O
CN
O
O
H
N
O
N
H
O
N
NH
S
O
O

**12l**

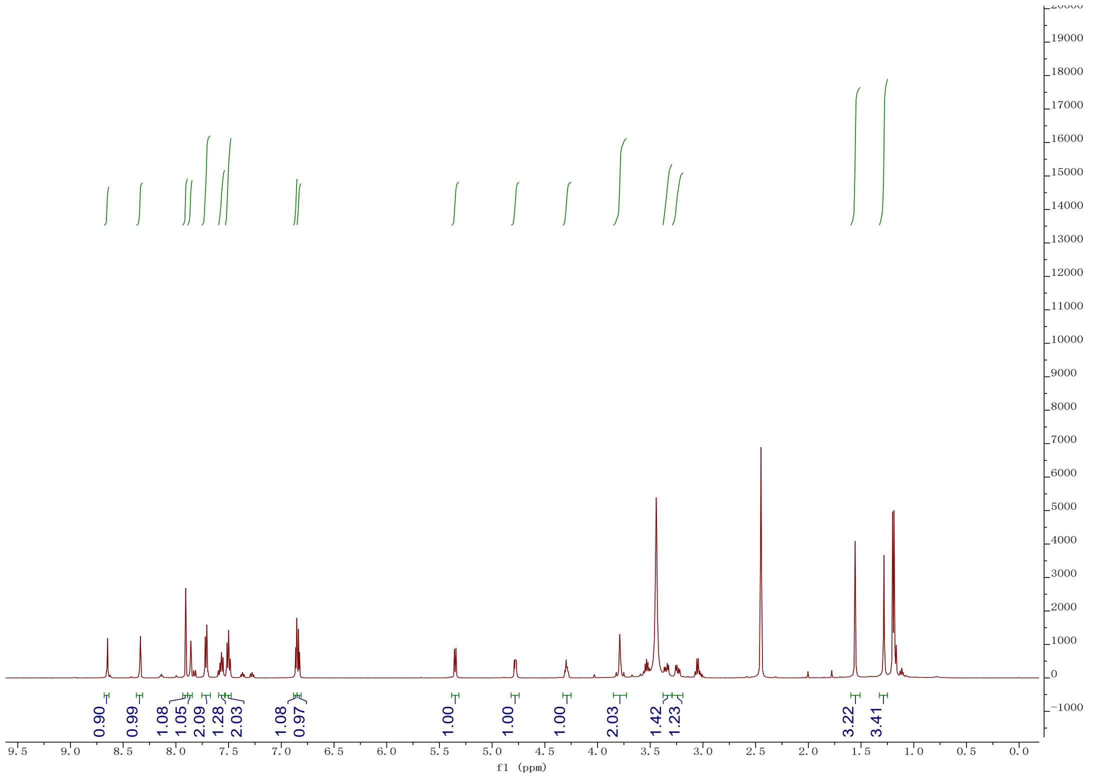
0.90
0.99
1.08
1.05
2.09
1.28
2.03
1.08
0.97
1.00
1.00
1.00
2.03
1.42
1.23
3.22
3.41
f1 (ppm)

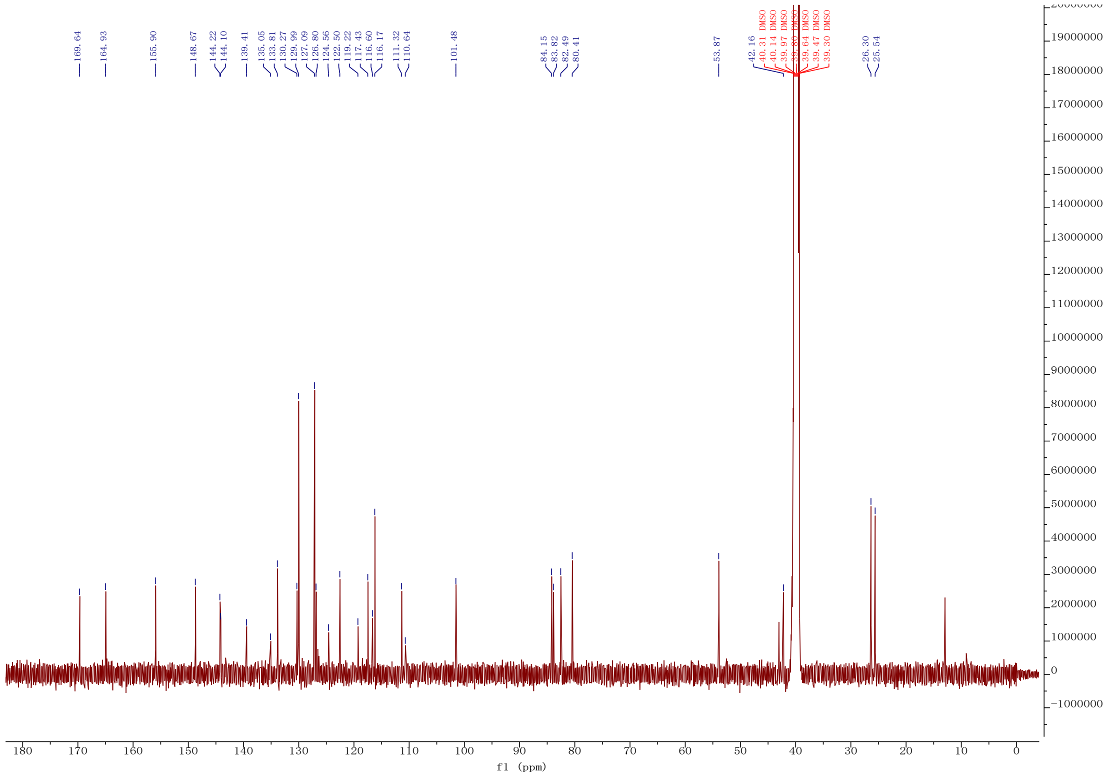
169.64
164.93
155.90
148.67
144.22
144.10
139.41
135.05
133.81
130.27
129.99
127.09
126.80
124.56
122.50
119.22
117.43
116.60
116.17
111.32
110.64
101.48
84.15
83.82
82.49
80.41
53.87
42.16
40.31 DMSO
40.14 DMSO
39.97 DMSO
39.80 DMSO
39.64 DMSO
39.47 DMSO
39.30 DMSO
26.30
25.54
f1 (ppm)

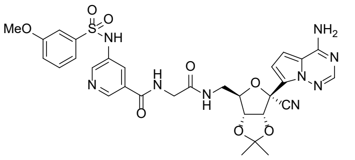
MeO
O
O
S
NH
N
H
N
O
N
H
O
O
O
NH2
N
N
N
CN

**12m**

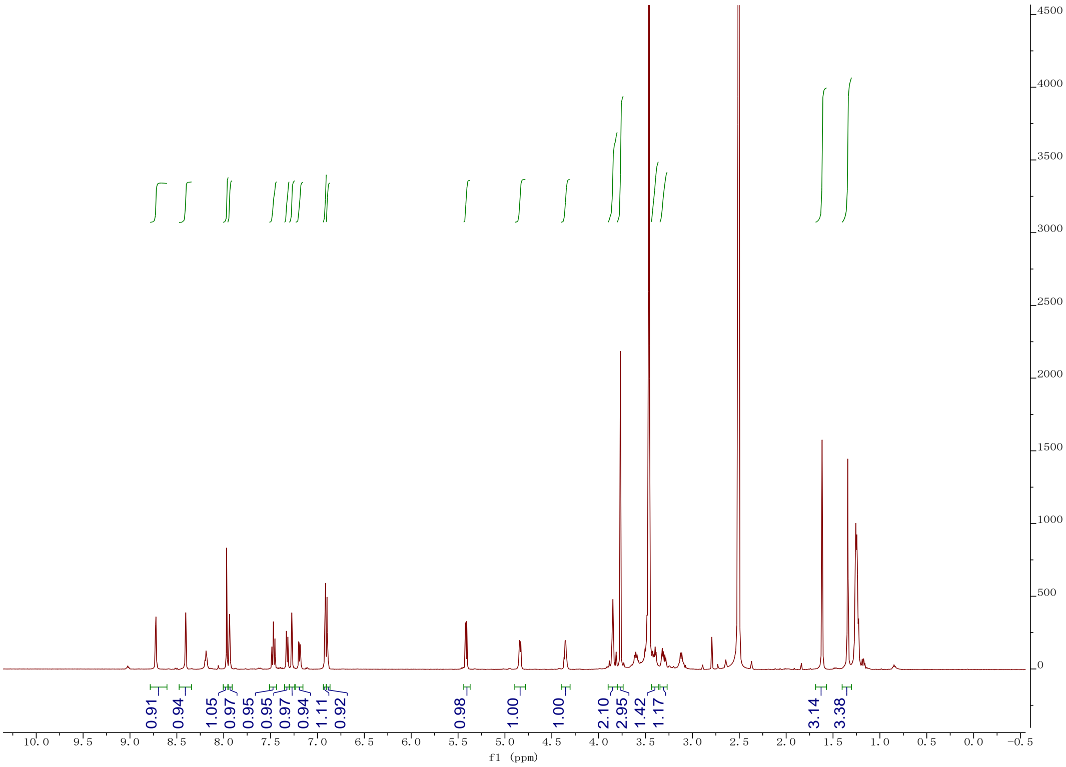
0.91
0.94
1.05
0.97
0.95
0.95
0.97
0.94
1.11
0.92
0.98
1.00
1.00
2.10
2.95
1.42
1.17
3.14
3.38
f1 (ppm)

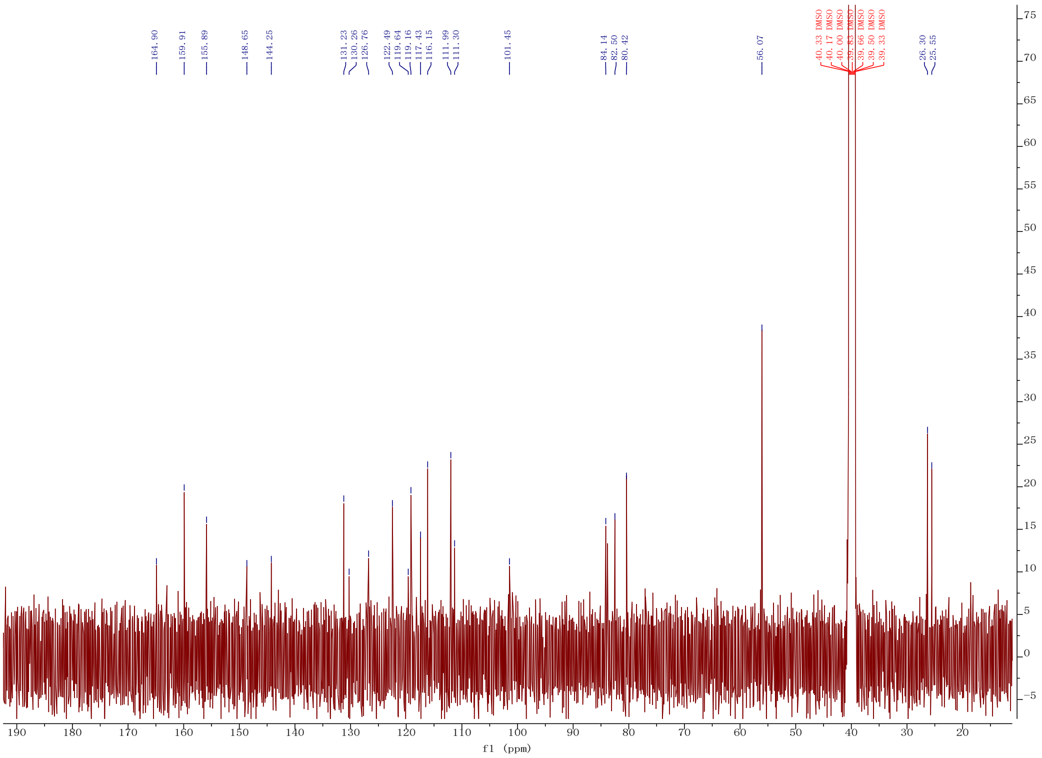
164.90
159.91
155.89
148.65
144.25
131.23
130.26
126.76
122.49
119.64
119.16
117.43
116.15
111.99
111.30
101.45
84.14
82.50
80.42
56.07
40.33 DMSO
40.17 DMSO
40.00 DMSO
39.83 DMSO
39.66 DMSO
39.50 DMSO
39.33 DMSO
26.30
25.55
f1 (ppm)

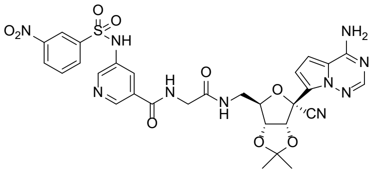

$O_2N$
O
O
S
NH
N
H
N
O
N
H
O
$NH_2$
N
N
N
CN
O
O
O


**12n**

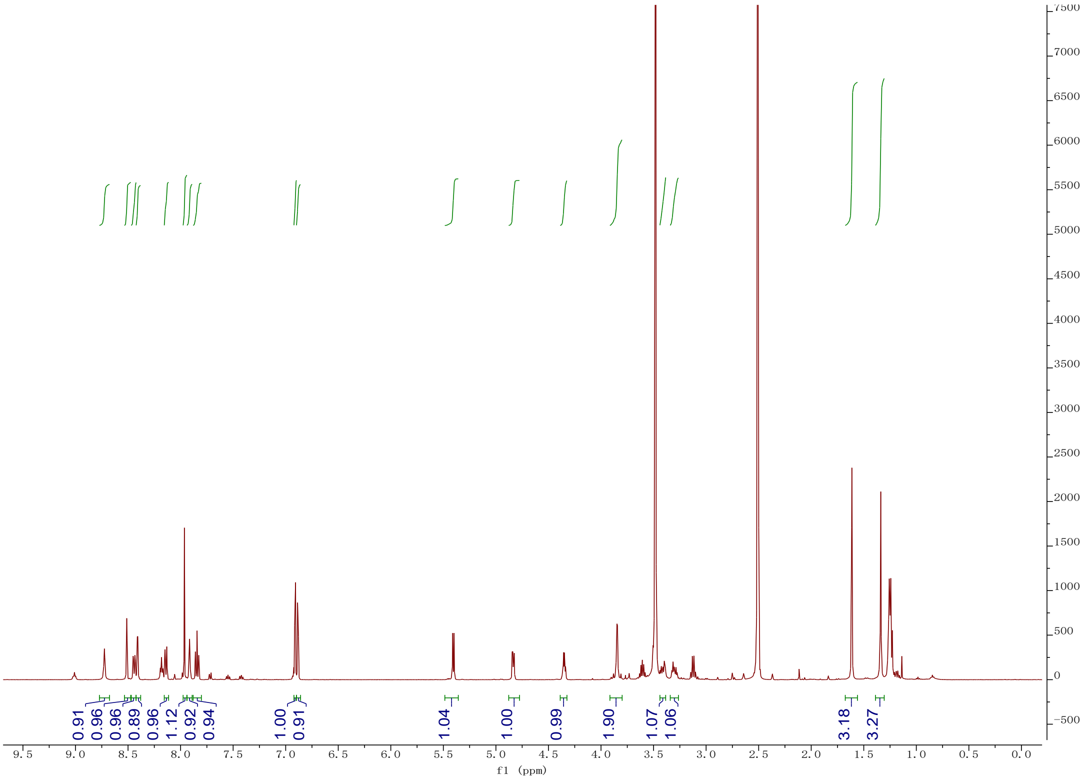

0.91
0.96
0.96
0.89
0.96
1.12
0.92
0.94
1.00
0.91
1.04
1.00
0.99
1.90
1.07
1.06
3.18
3.27
f1 (ppm)


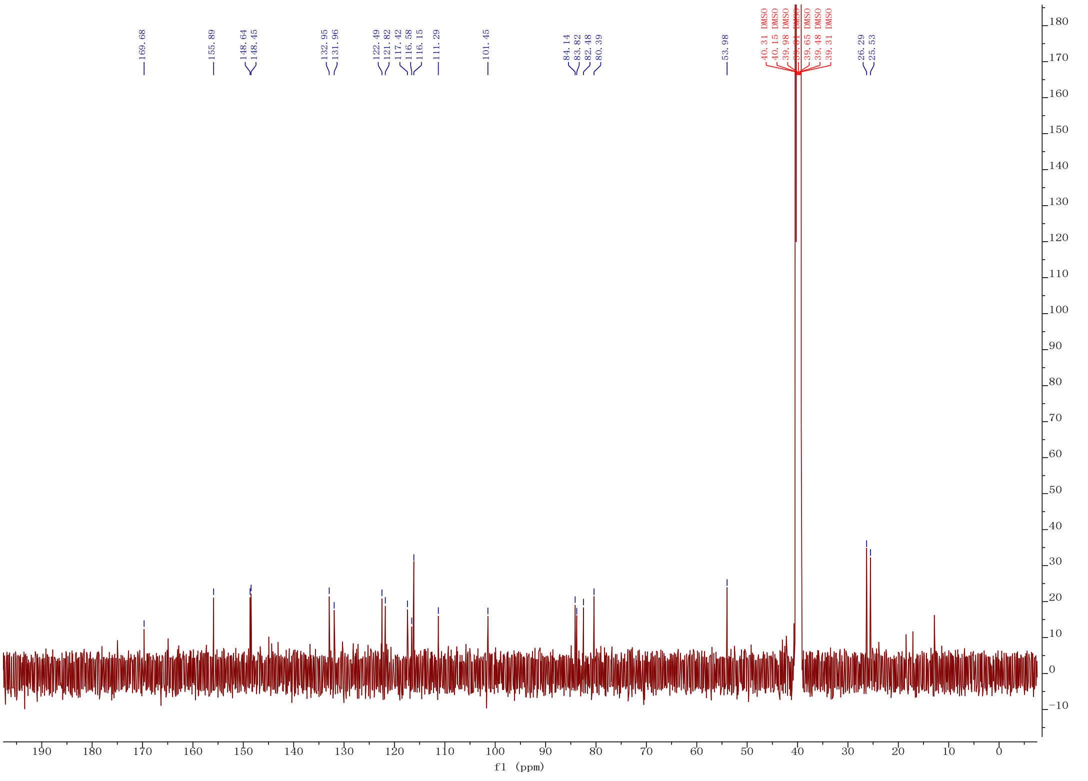

169.68
155.89
148.64
148.45
132.95
131.96
122.49
121.82
117.42
116.58
116.15
111.29
101.45
84.14
83.82
82.48
80.39
53.98
40.31 DMSO
40.15 DMSO
39.98 DMSO
39.81 DMSO
39.65 DMSO
39.48 DMSO
39.31 DMSO
26.29
25.53
f1 (ppm)

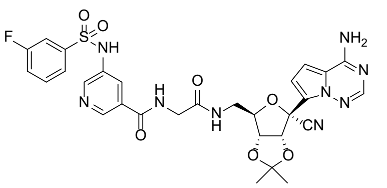

**12o**

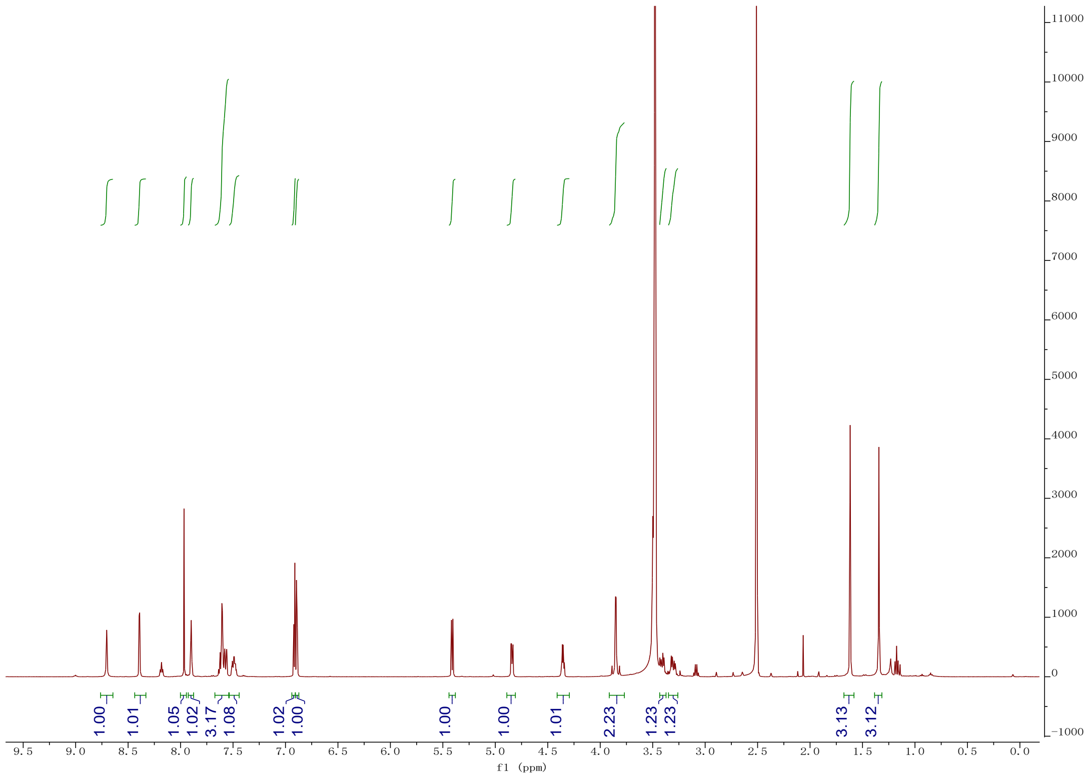

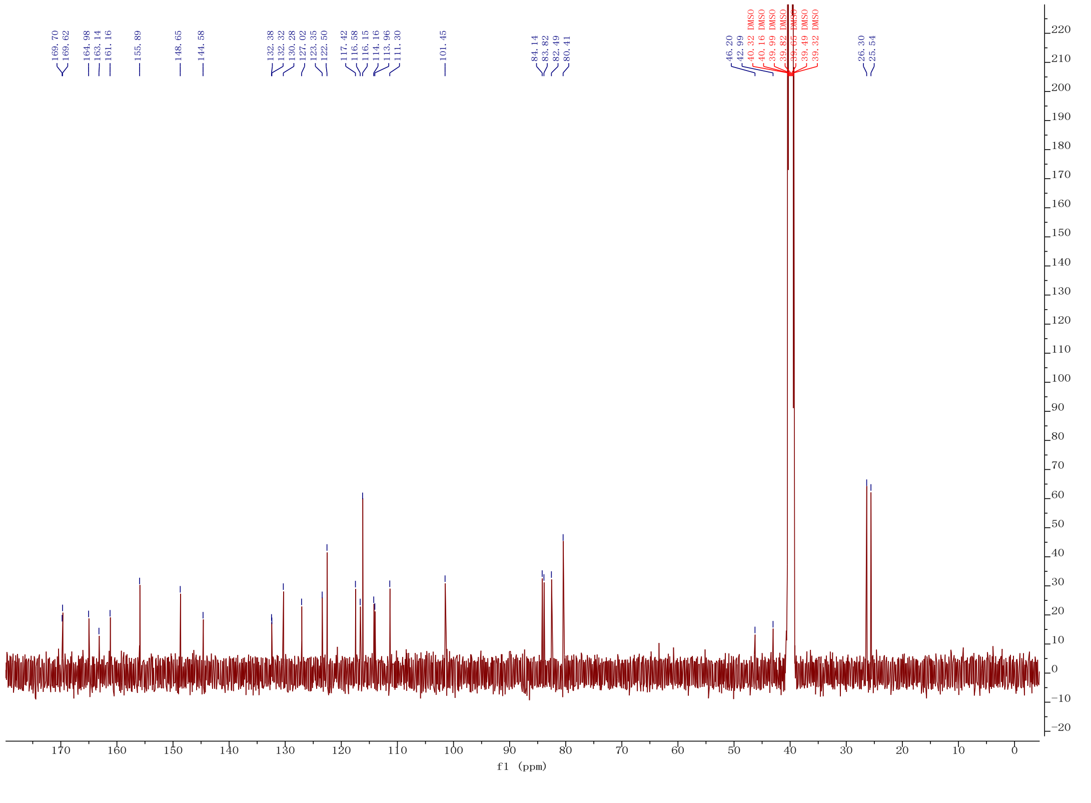

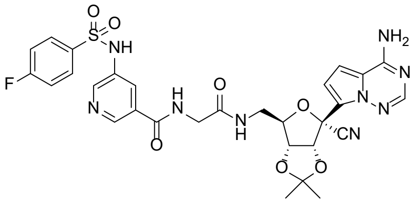
O
O
S
NH
F
N
H
N
O
O
N
H
O
NH2
N
N
N
CN
O
O

**12p**

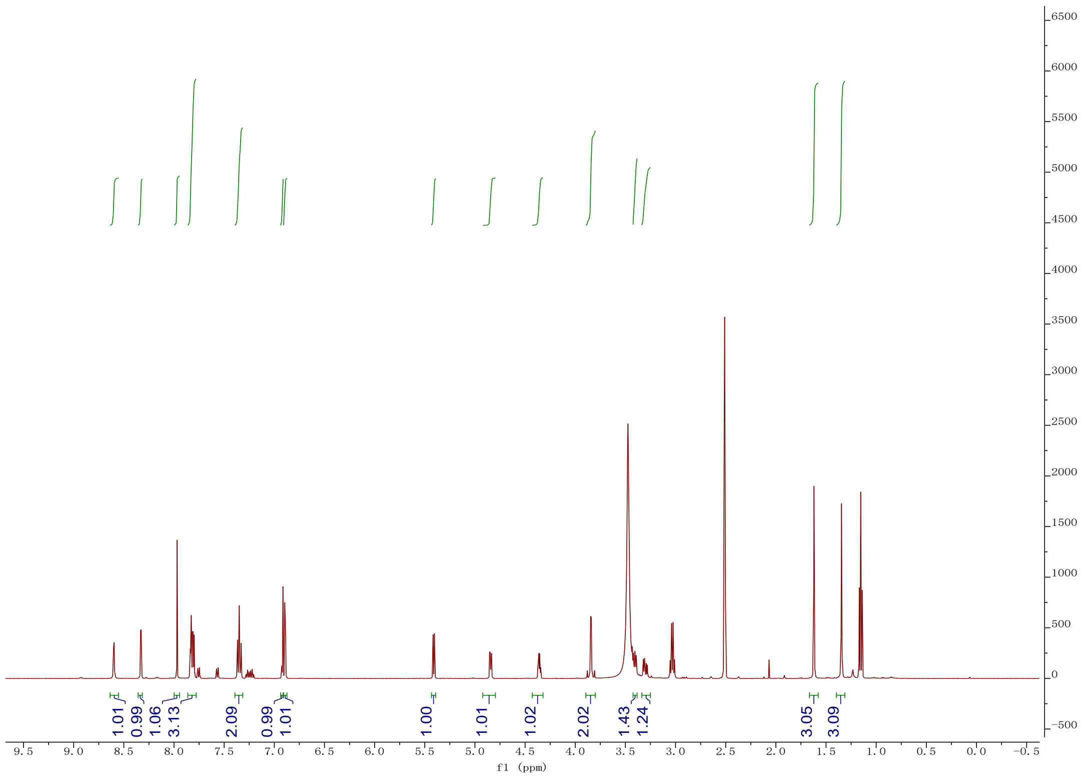
1.01
0.99
1.06
3.13
2.09
0.99
1.01
1.00
1.01
1.02
2.02
1.43
1.24
3.05
3.09
f1 (ppm)

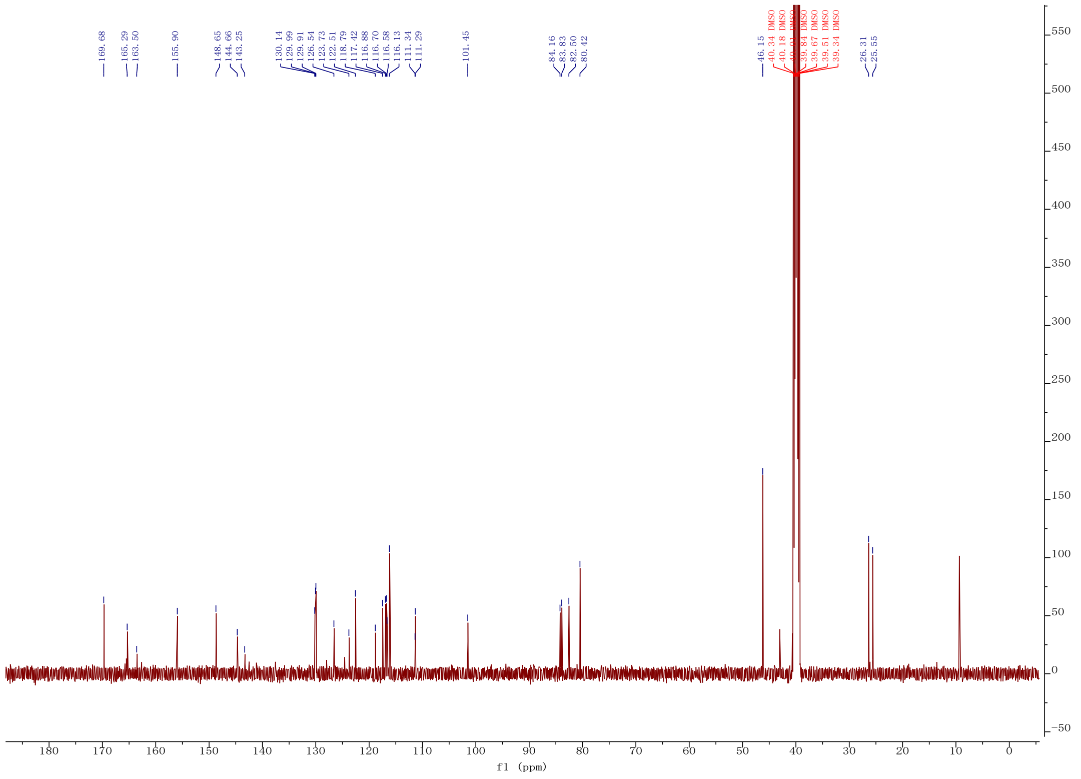
169.68
165.29
163.50
155.90
148.65
144.66
143.25
130.14
129.99
129.91
126.54
123.73
122.51
118.79
117.42
116.88
116.70
116.58
116.13
111.34
111.29
101.45
84.16
83.83
82.50
80.42
46.15
40.34 DMSO
40.18 DMSO
39.84 DMSO
39.67 DMSO
39.51 DMSO
39.34 DMSO
26.31
25.55
f1 (ppm)

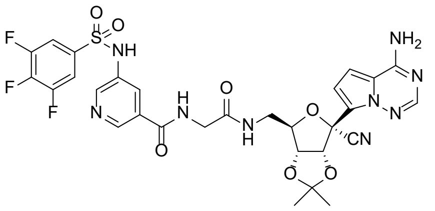
F
F
F
O
O
S
NH
N
H
N
O
O
N
H
O
NH2
N
N
N
CN
O
O

**12q**

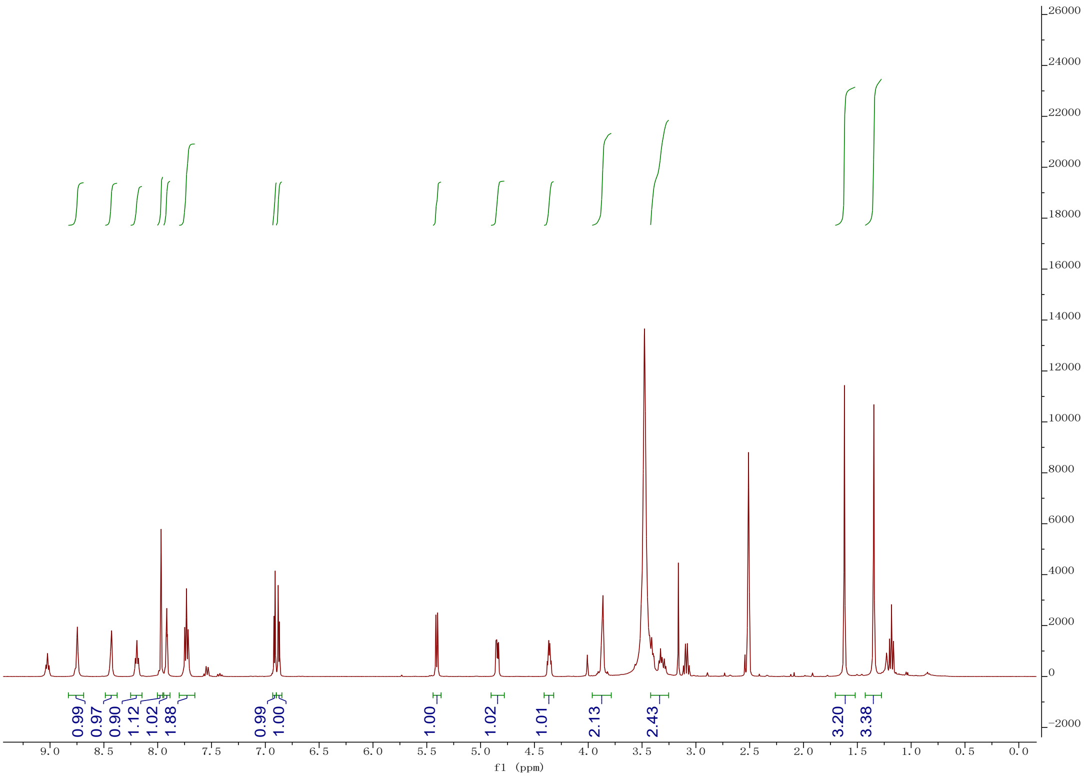
0.99
0.97
0.90
1.12
1.02
1.88
0.99
1.00
1.00
1.02
1.01
2.13
2.43
3.20
3.38
f1 (ppm)

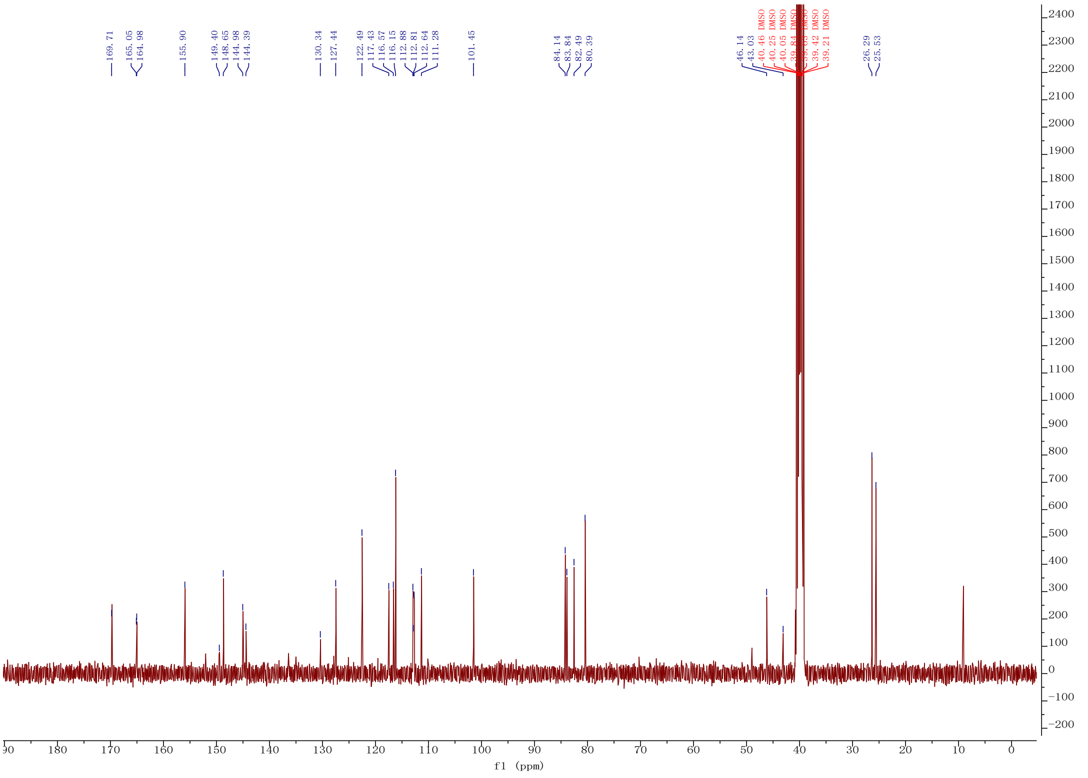
169.71
165.05
164.98
155.90
149.40
148.65
144.98
144.39
130.34
127.44
122.49
117.43
116.57
116.15
112.88
112.81
112.64
111.28
101.45
84.14
83.84
82.49
80.39
46.14
43.03
40.46 DMSO
40.25 DMSO
40.05 DMSO
39.84 DMSO
39.42 DMSO
39.21 DMSO
26.29
25.53
f1 (ppm)

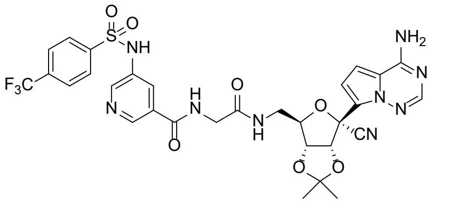
O
S
O
NH
F$_3$C
N
H
N
O
O
N
H
O
NH$_2$
N
N
N
CN
O
O

**12r**

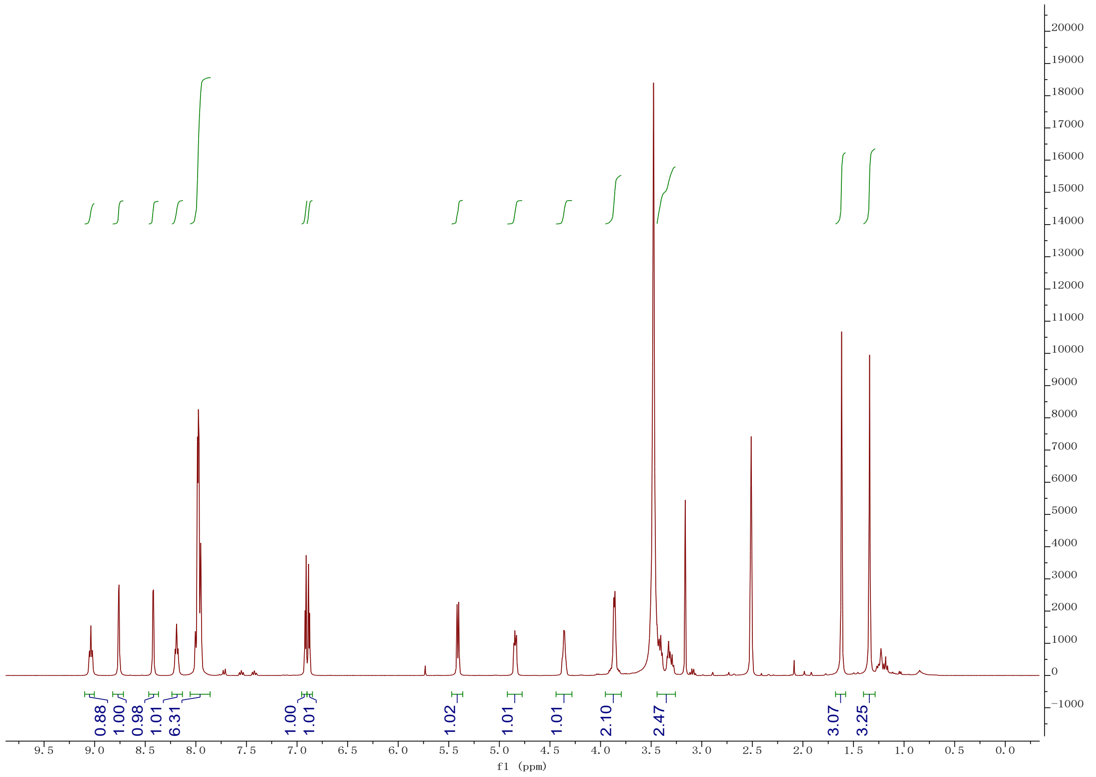
0.88
1.00
0.98
1.01
6.31
1.00
1.01
1.02
1.01
1.01
2.10
2.47
3.07
3.25
f1 (ppm)

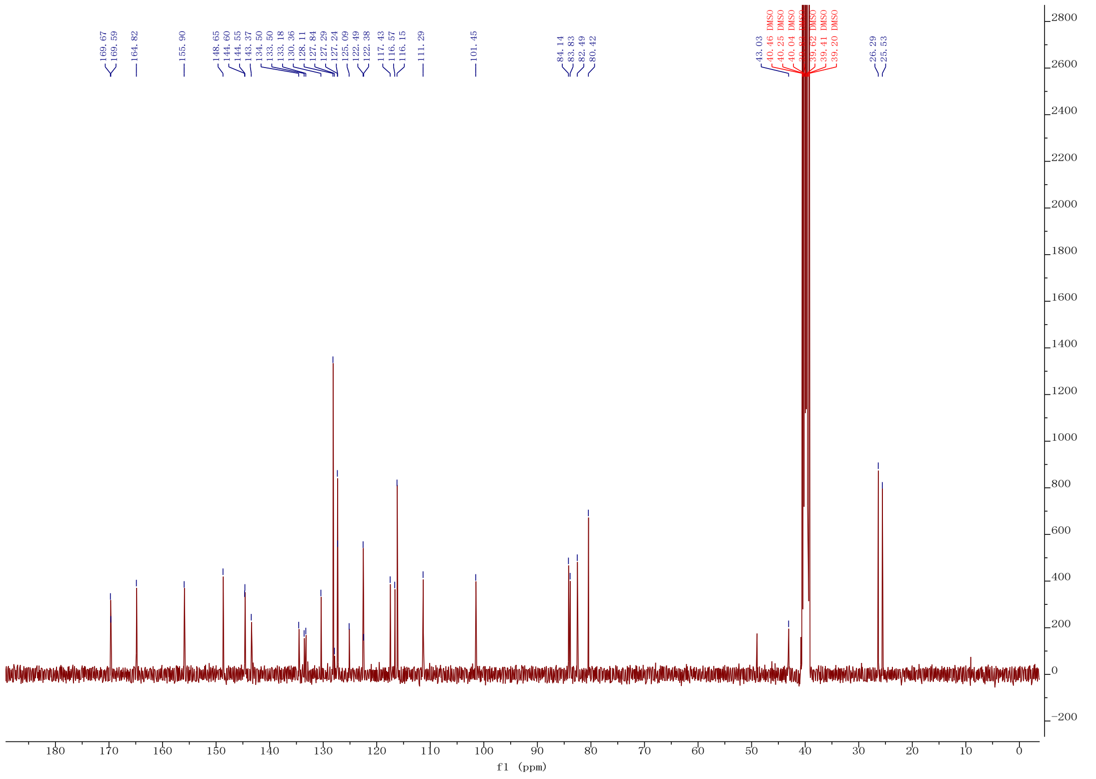
169.67
169.59
164.82
155.90
148.65
144.60
144.55
143.37
134.50
133.50
133.18
130.36
128.11
127.84
127.29
127.24
125.09
122.49
122.38
117.43
116.57
116.15
111.29
101.45
84.14
83.83
82.49
80.42
43.03
40.46 DMSO
40.25 DMSO
40.04 DMSO
39.62 DMSO
39.41 DMSO
39.20 DMSO
26.29
25.53
f1 (ppm)

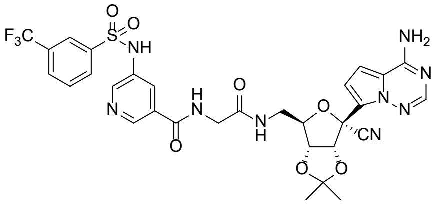
F3C
O
S
O
NH
N
H
N
O
O
N
H
O
NH2
N
N
N
CN
O
O

**12s**

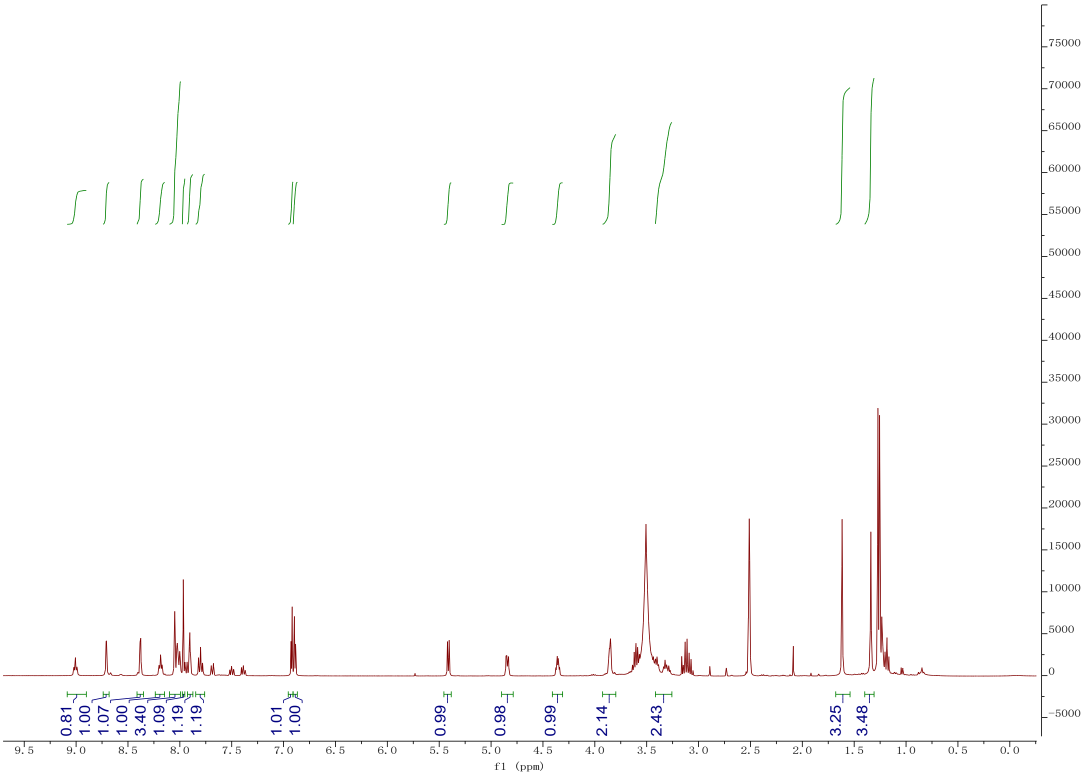
0.81
1.00
1.07
1.00
3.40
1.09
1.19
1.19
1.01
1.00
0.99
0.98
0.99
2.14
2.43
3.25
3.48
f1 (ppm)

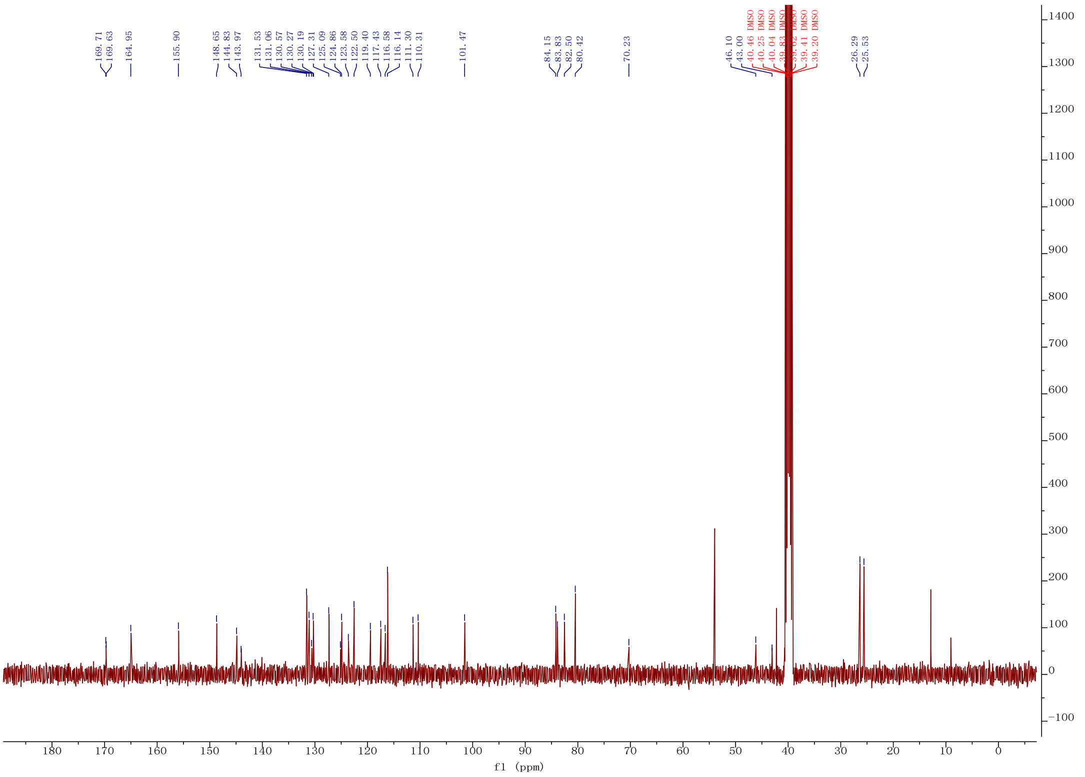
169.71
169.63
164.95
155.90
148.65
144.83
143.97
131.53
131.06
130.57
130.27
130.19
127.31
125.09
124.86
123.58
122.50
119.40
117.43
116.58
116.14
111.30
110.31
101.47
84.15
83.83
82.50
80.42
70.23
46.10
43.00
40.46 DMSO
40.25 DMSO
40.04 DMSO
39.83 DMSO
39.62 DMSO
39.41 DMSO
39.20 DMSO
26.29
25.53
f1 (ppm)

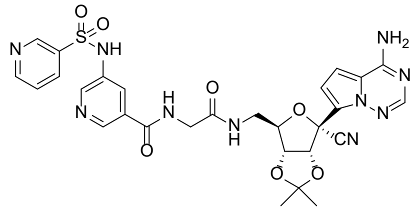
NH
N
NH2
N
N
N
H
O
O
O
S
O
CN
O
O
H
N
N

**12t**

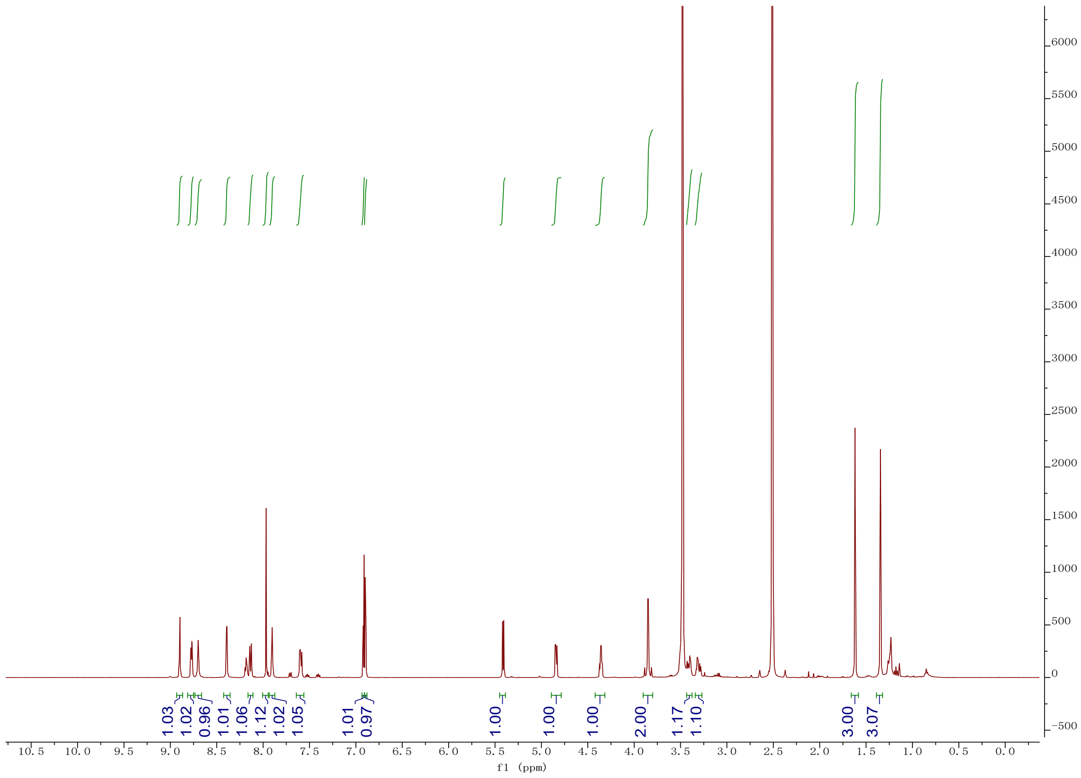
1.03
1.02
0.96
1.01
1.06
1.12
1.02
1.05
1.01
0.97
1.00
1.00
1.00
2.00
1.17
1.10
3.00
3.07
f1 (ppm)

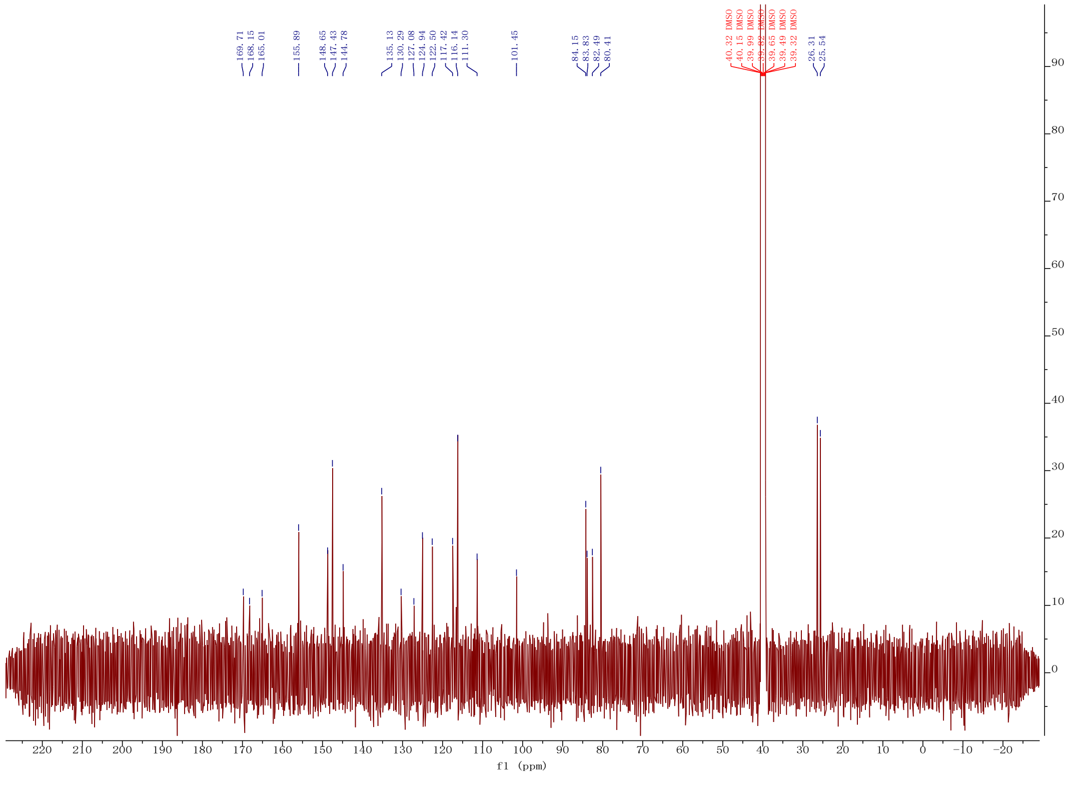
169.71
168.15
165.01
155.89
148.65
147.43
144.78
135.13
130.29
127.08
124.94
122.50
117.42
116.14
111.30
101.45
84.15
83.83
82.49
80.41
40.32 DMSO
40.15 DMSO
39.99 DMSO
39.82 DMSO
39.65 DMSO
39.49 DMSO
39.32 DMSO
26.31
25.54
f1 (ppm)

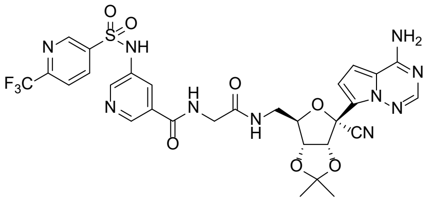

O
O
S
N
NH
F3C
N
H
N
O
N
O
H
N
O
NH2
N
N
N
CN
O
O


**12u**

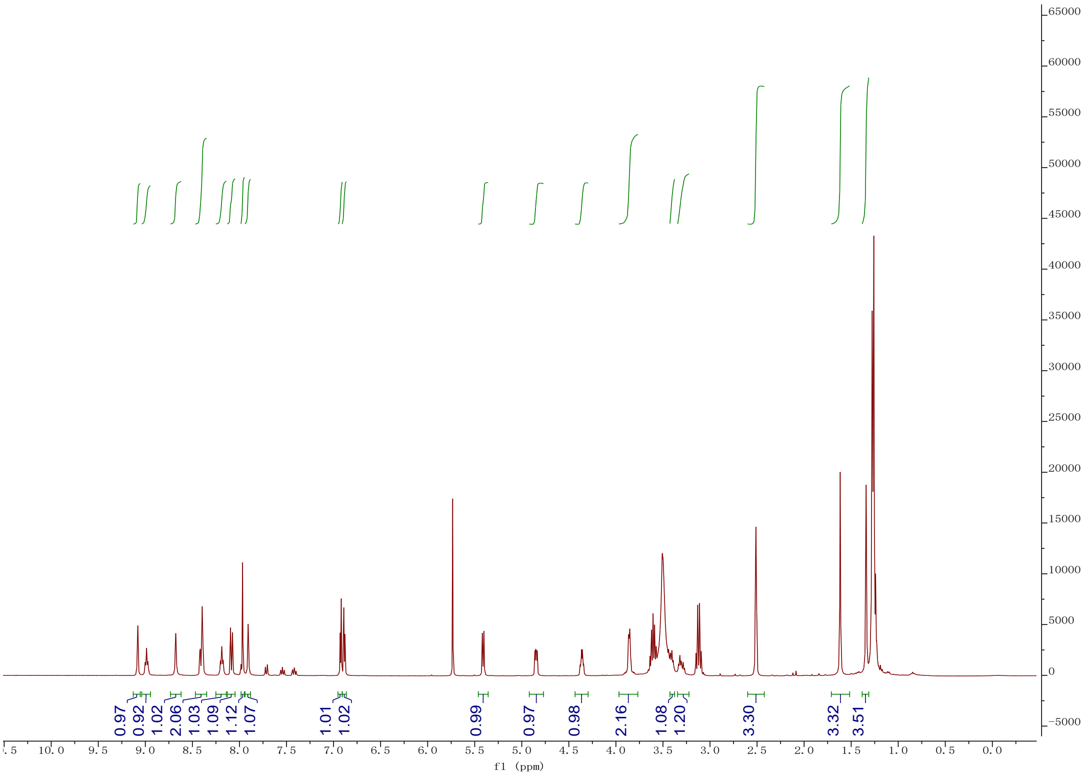

0.97
0.92
1.02
2.06
1.03
1.09
1.12
1.07
1.01
1.02
0.99
0.97
0.98
2.16
1.08
1.20
3.30
3.32
3.51
f1 (ppm)


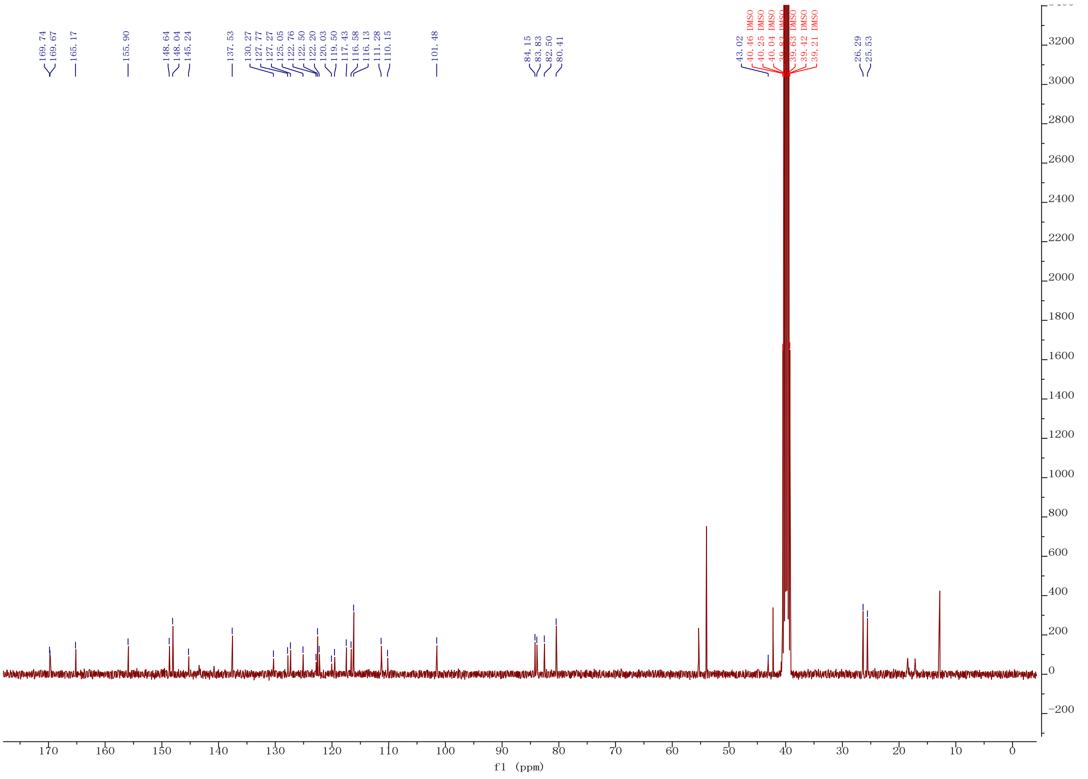

169.74
169.67
165.17
155.90
148.64
148.04
145.24
137.53
130.27
127.77
127.27
125.05
122.76
122.50
122.20
120.03
119.50
117.43
116.58
116.13
111.28
110.15
101.48
84.15
83.83
82.50
80.41
43.02
40.46 DMSO
40.25 DMSO
40.04 DMSO
39.63 DMSO
39.42 DMSO
39.21 DMSO
26.29
25.53
f1 (ppm)

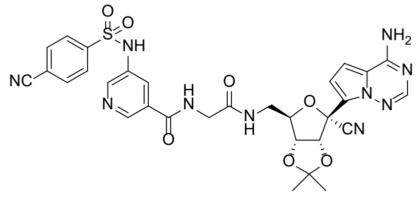
O
S
O
NH
NC
N
H
N
O
O
N
H
O
NH2
N
N
N
CN
O
O

**12v**

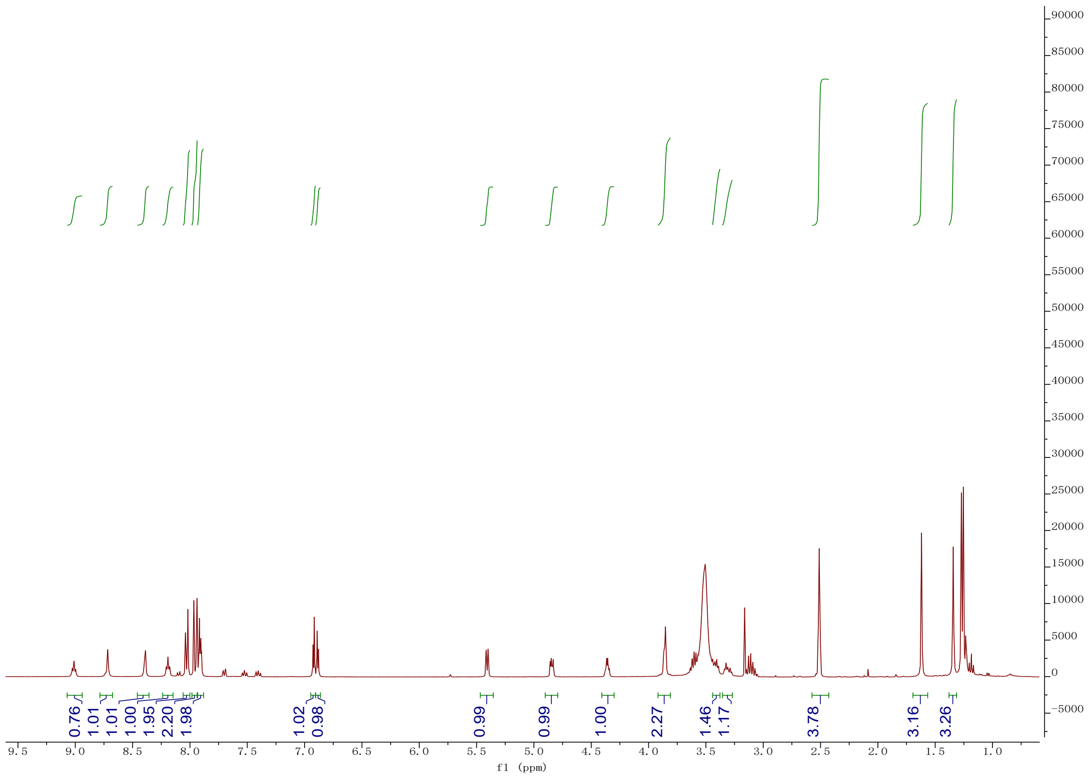
0.76
1.01
1.00
1.95
2.20
1.98
1.02
0.98
0.99
0.99
1.00
2.27
1.46
1.17
3.78
3.16
3.26
f1 (ppm)

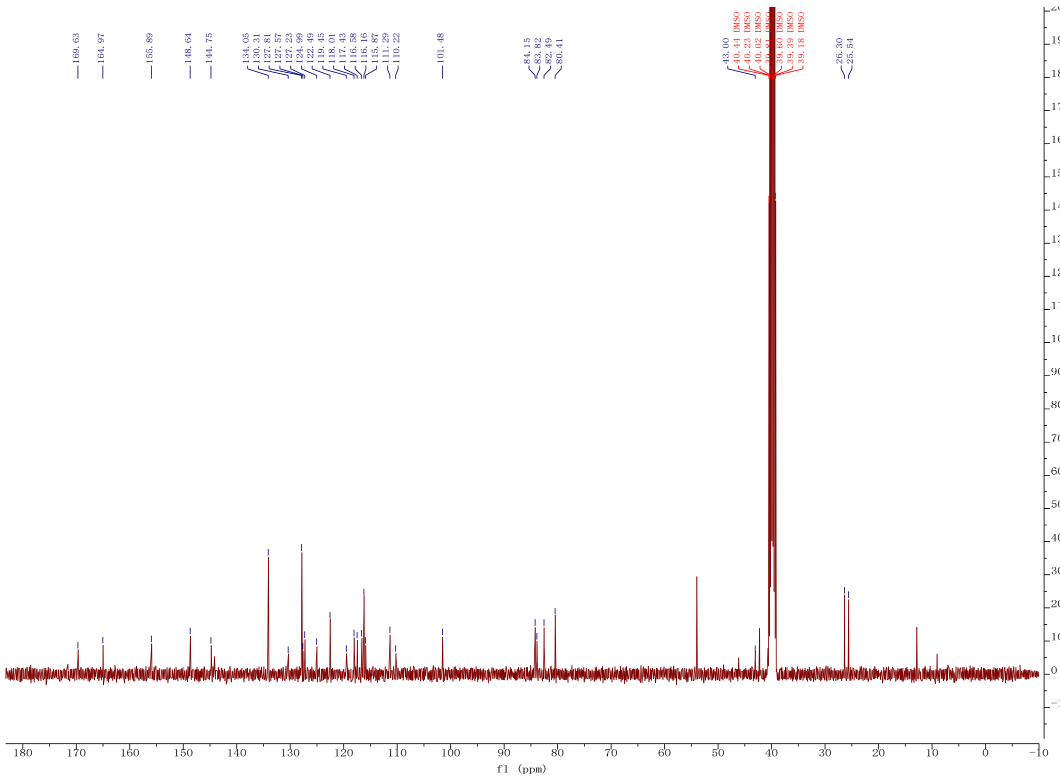
169.63
164.97
155.89
148.64
144.75
134.05
130.31
127.81
127.57
127.23
124.99
122.49
119.45
118.01
117.43
116.58
116.16
115.87
111.29
110.22
101.48
84.15
83.82
82.49
80.41
43.00
40.44 DMSO
40.23 DMSO
40.02 DMSO
39.60 DMSO
39.39 DMSO
39.18 DMSO
26.30
25.54
f1 (ppm)

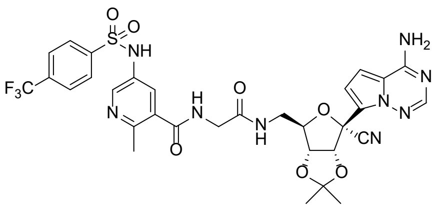

NH
NH2
F3C
N
CN


**12w**

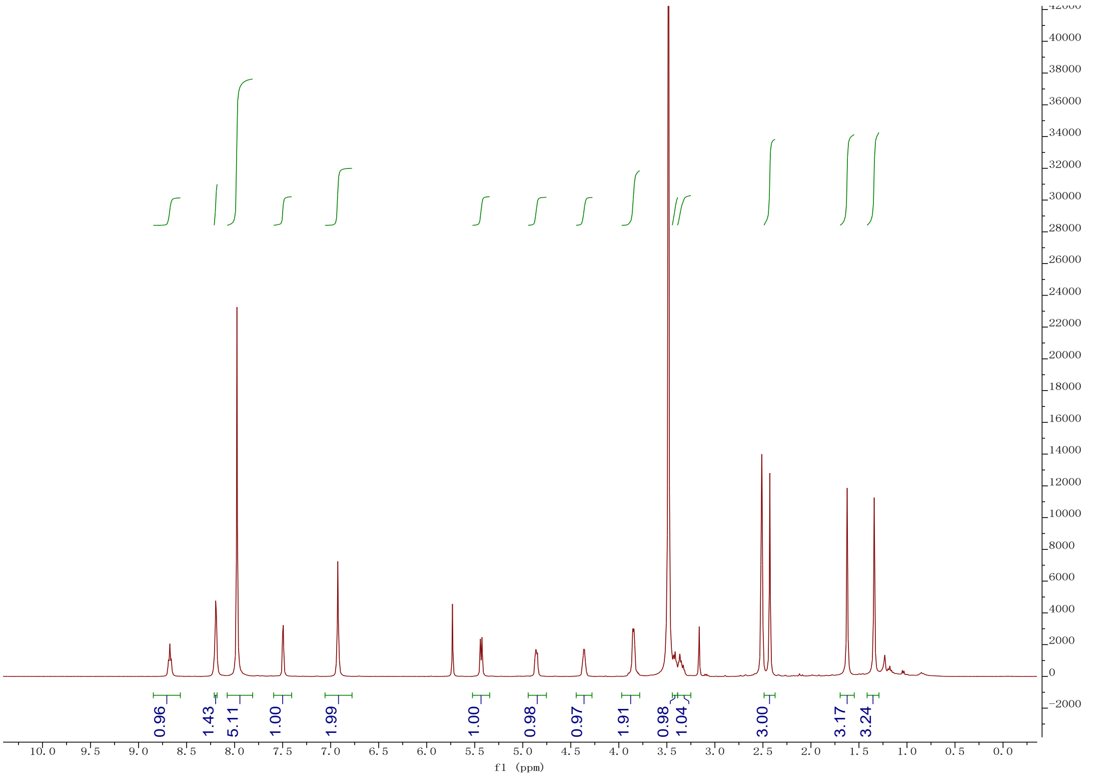

0.96
1.43
5.11
1.00
1.99
1.00
0.98
0.97
1.91
0.98
1.04
3.00
3.17
3.24
f1 (ppm)


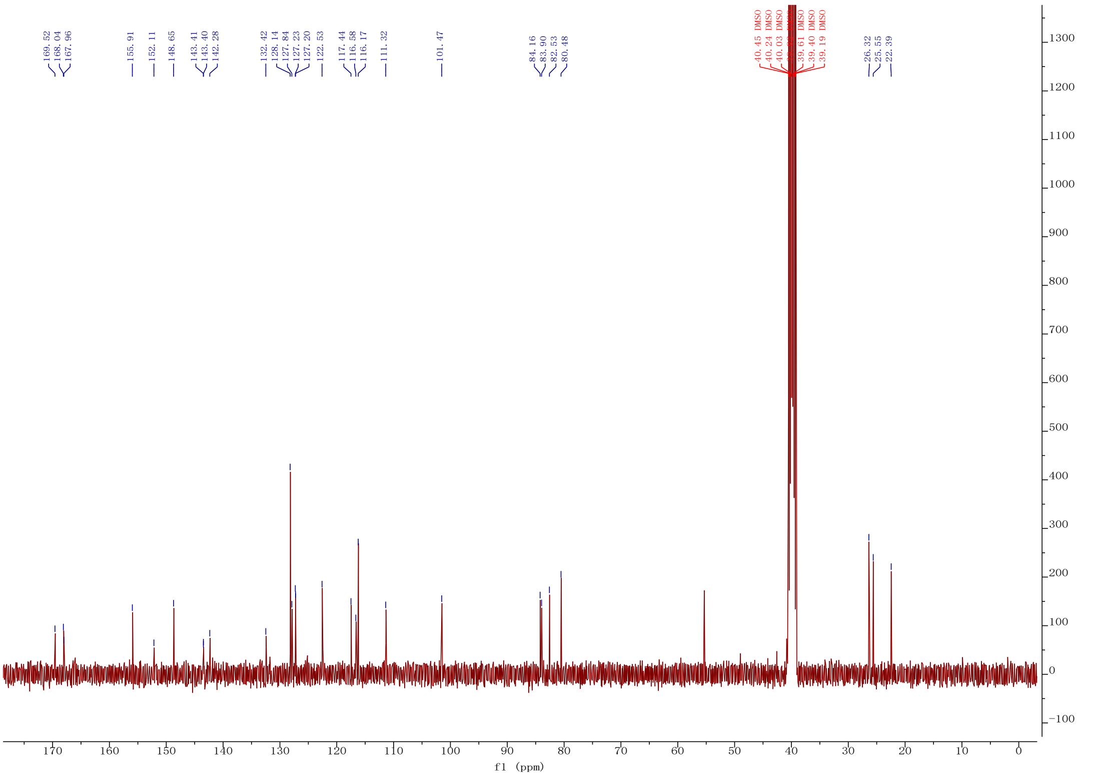

169.52
168.04
167.96
155.91
152.11
148.65
143.41
143.40
142.28
132.42
128.14
127.84
127.23
127.20
122.53
117.44
116.58
116.17
111.32
101.47
84.16
83.90
82.53
80.48
40.45 DMSO
40.24 DMSO
40.03 DMSO
39.61 DMSO
39.40 DMSO
39.19 DMSO
26.32
25.55
22.39
f1 (ppm)

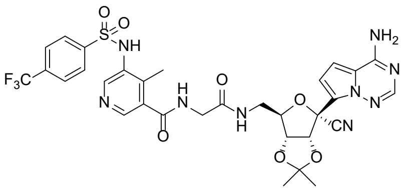

NH
NH2
F3C
N
H
O
N
H
O
O
O
CN


**12x**

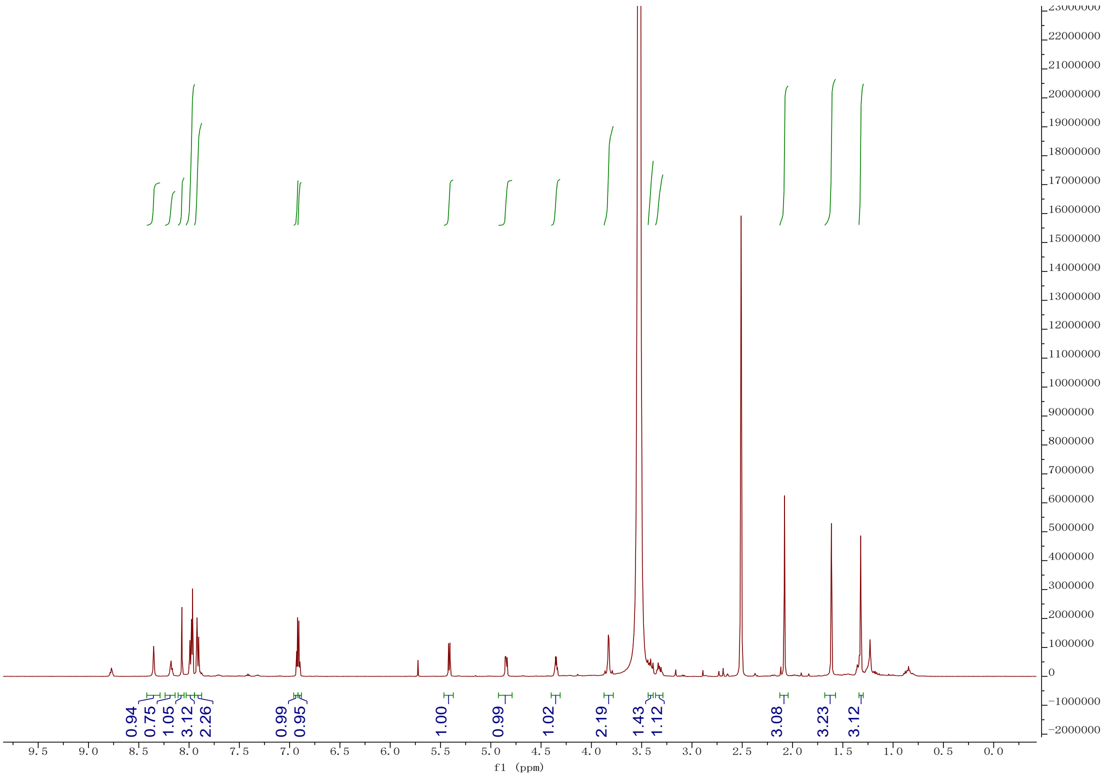

0.94
0.75
1.05
3.12
2.26
0.99
0.95
1.00
0.99
1.02
2.19
1.43
1.12
3.08
3.23
3.12
f1 (ppm)


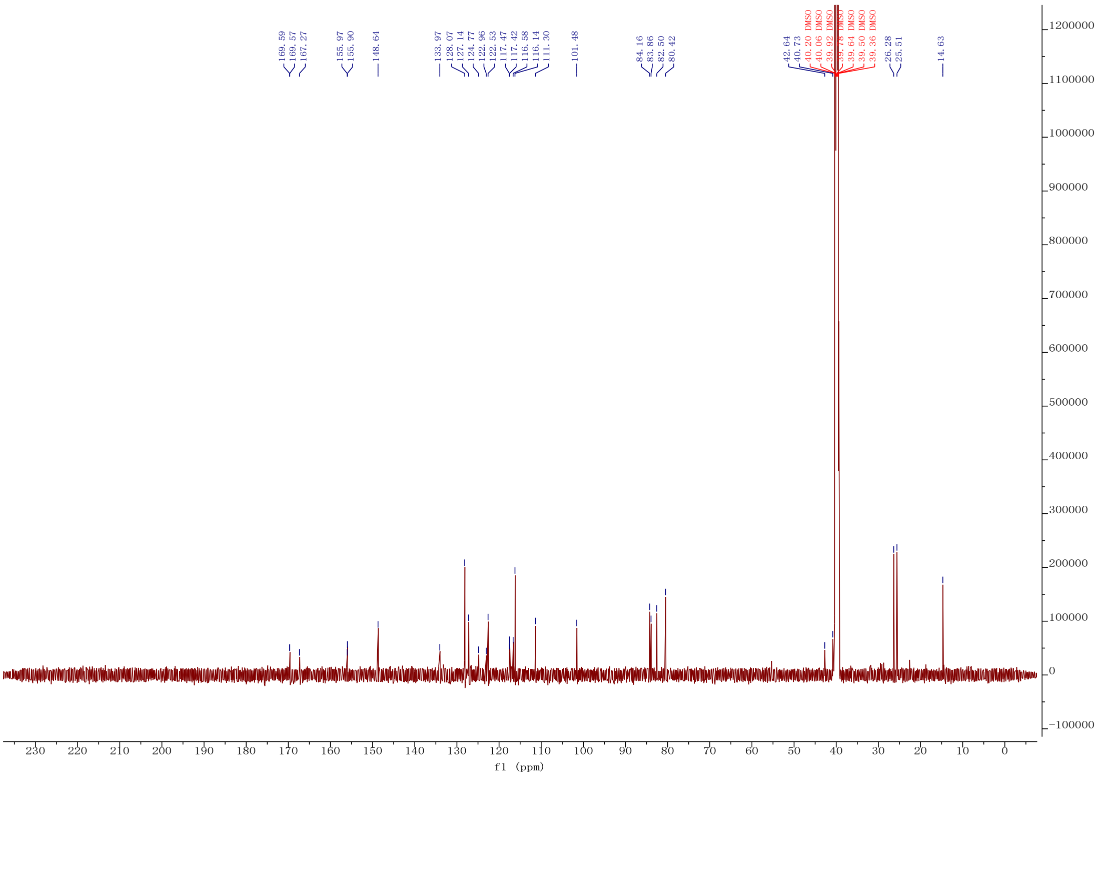

169.59
169.57
167.27
155.97
155.90
148.64
133.97
128.07
127.14
124.77
122.96
122.53
117.47
117.42
116.58
116.14
111.30
101.48
84.16
83.86
82.50
80.42
42.64
40.73
40.20 DMSO
40.06 DMSO
39.92 DMSO
39.78 DMSO
39.64 DMSO
39.50 DMSO
39.36 DMSO
26.28
25.51
14.63
f1 (ppm)

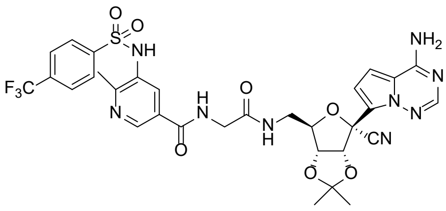
O
O
S
NH
F3C
N
H
N
O
O
N
H
O
O
O
CN
NH2
N
N
N

**12y**

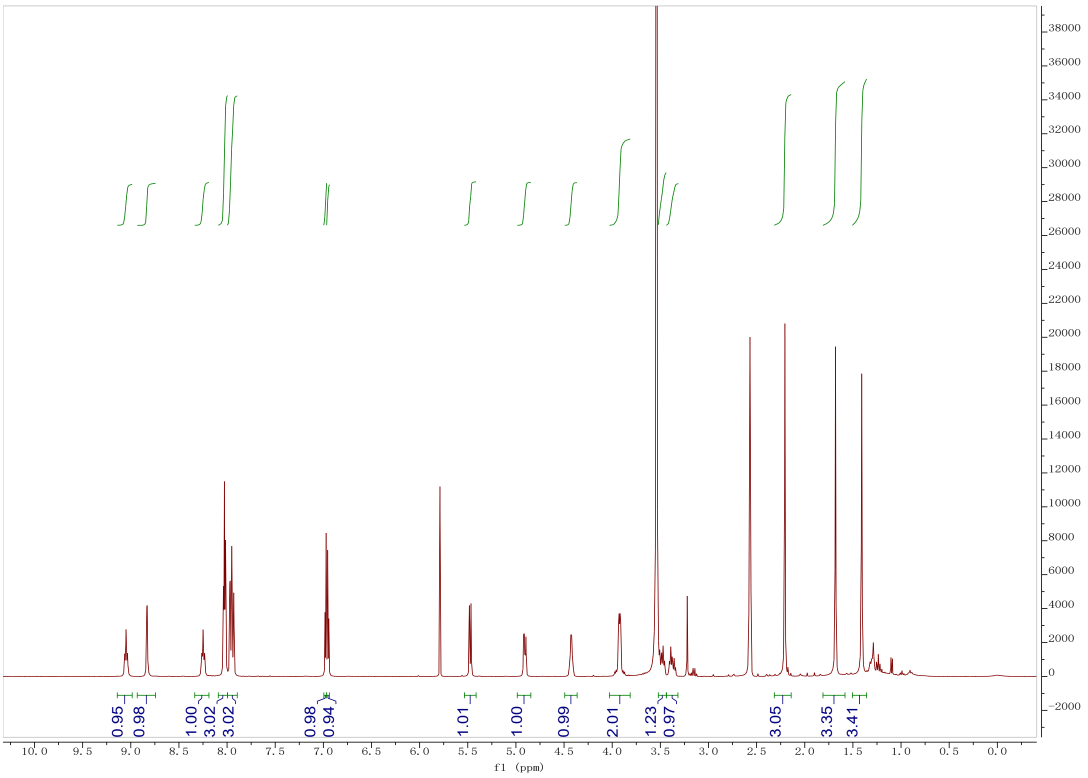
0.95
0.98
1.00
3.02
3.02
0.98
0.94
1.01
1.00
0.99
2.01
1.23
0.97
3.05
3.35
3.41
f1 (ppm)

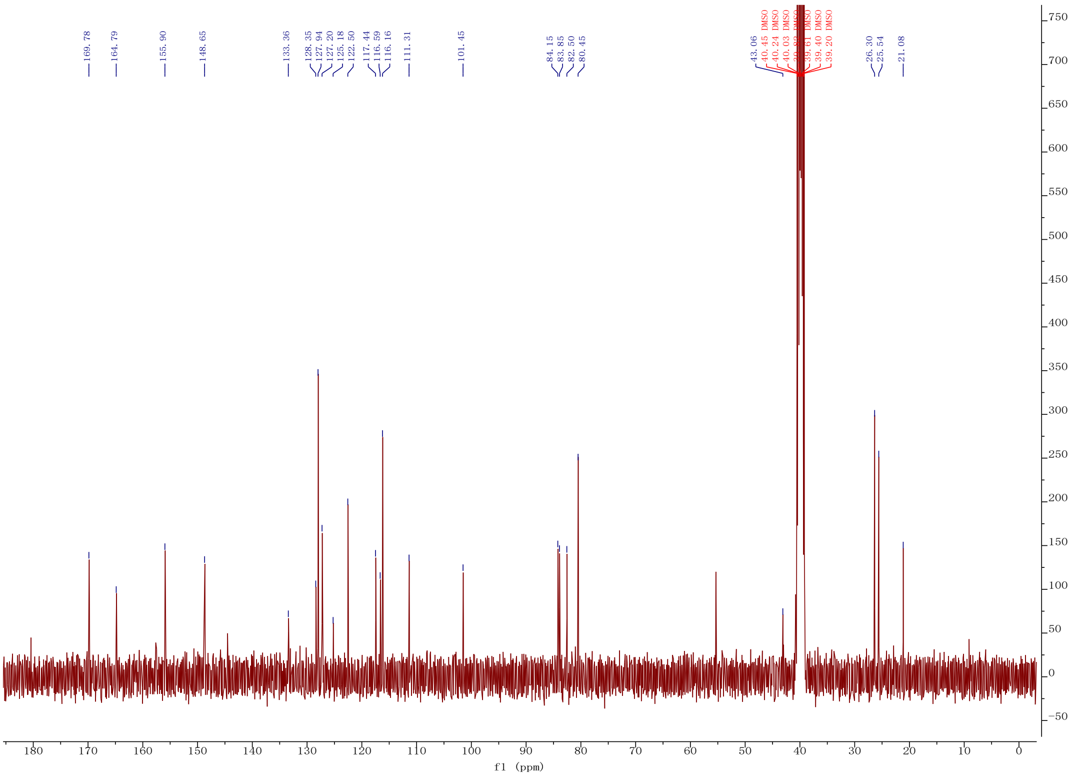
169.78
164.79
155.90
148.65
133.36
128.35
127.94
127.20
125.18
122.50
117.44
116.59
116.16
111.31
101.45
84.15
83.85
82.50
80.45
43.06
26.30
25.54
21.08
f1 (ppm)

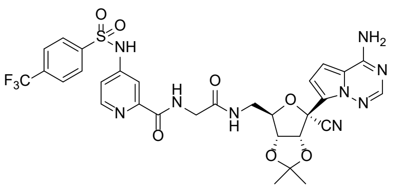
O
S
O
NH
F3C
H
N
N
O
O
N
H
O
NH2
N
N
N
CN
O
O

**12z**

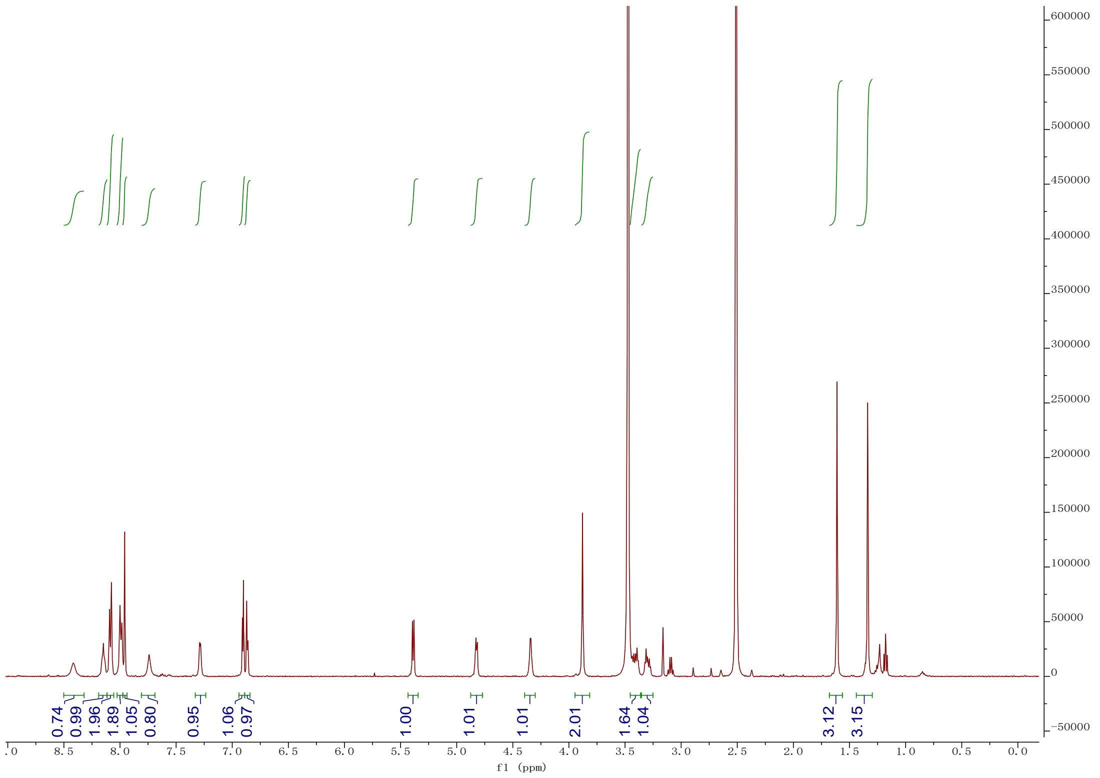
0.74
0.99
1.96
1.89
1.05
0.80
0.95
1.06
0.97
1.00
1.01
1.01
2.01
1.64
1.04
3.12
3.15
f1 (ppm)

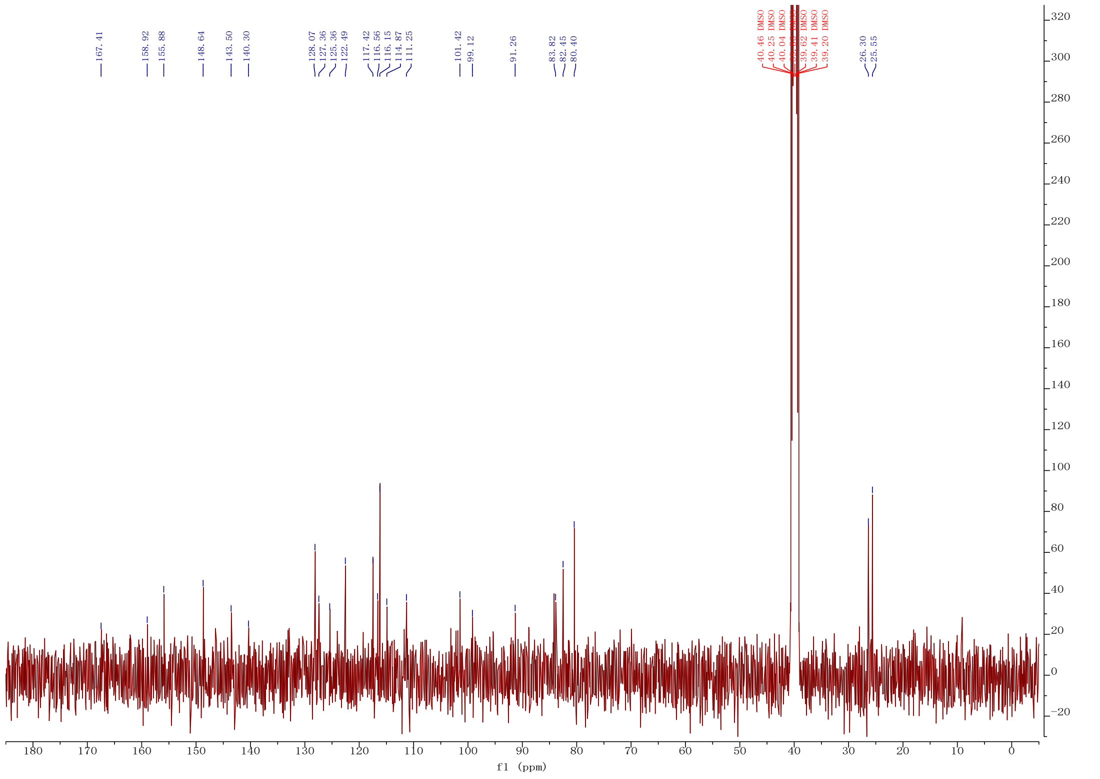
167.41
158.92
155.88
148.64
143.50
140.30
128.07
127.36
125.36
122.49
117.42
116.56
116.15
114.87
111.25
101.42
99.12
91.26
83.82
82.45
80.40
40.46 DMSO
40.25 DMSO
40.04 DMSO
39.41 DMSO
39.20 DMSO
26.30
25.55
f1 (ppm)

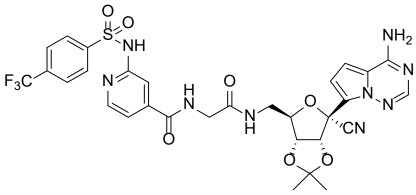

**12aa**

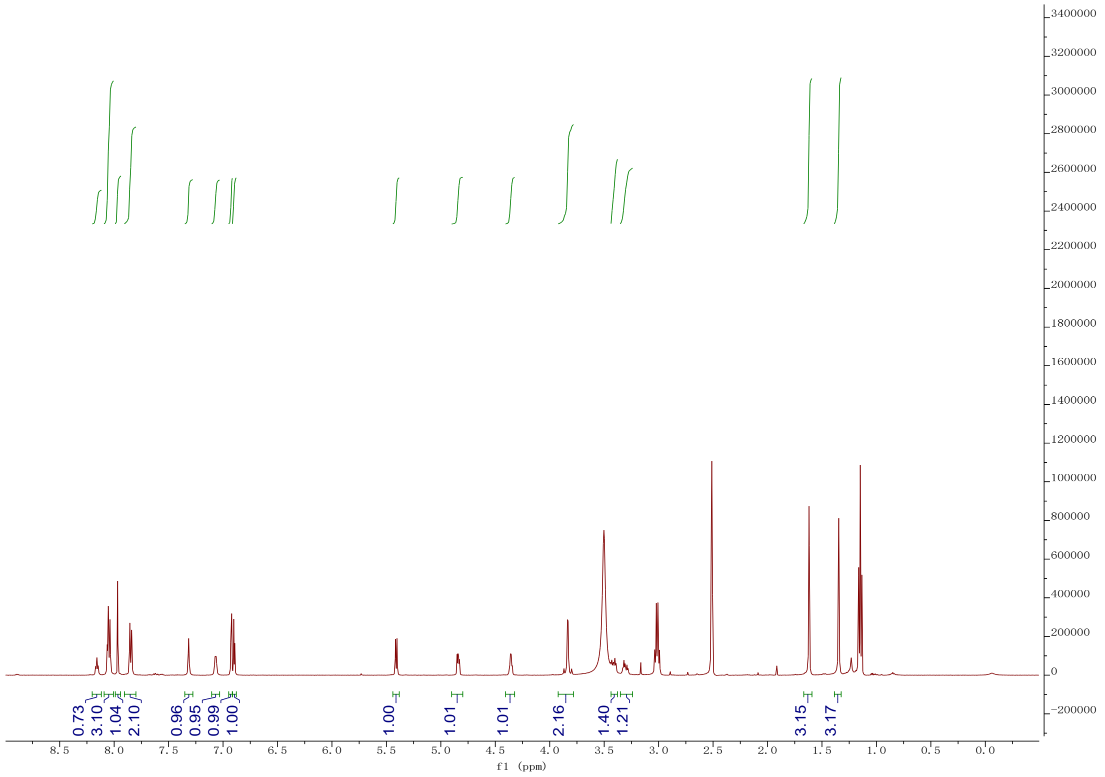

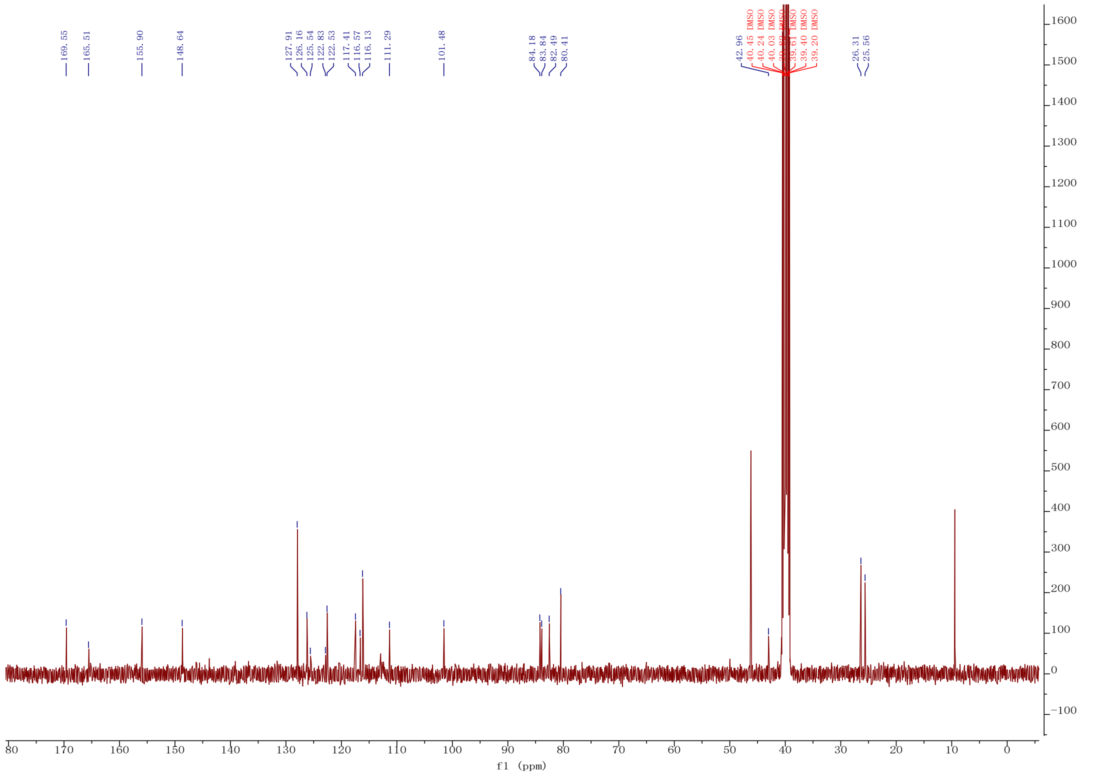

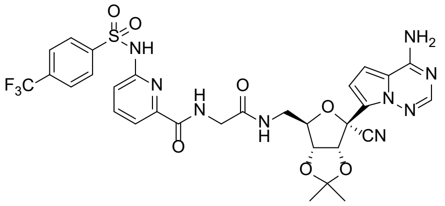

**12ab**

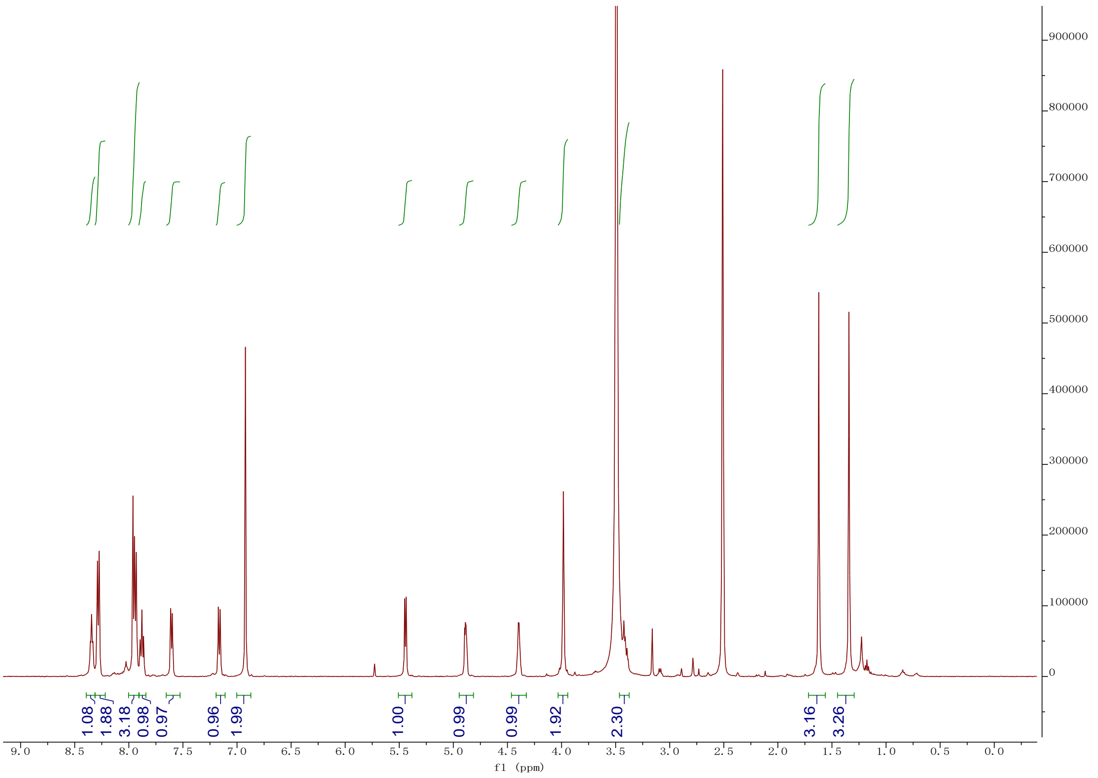
1.08
1.88
3.18
0.98
0.97
0.96
1.99
1.00
0.99
0.99
1.92
2.30
3.16
3.26
f1 (ppm)

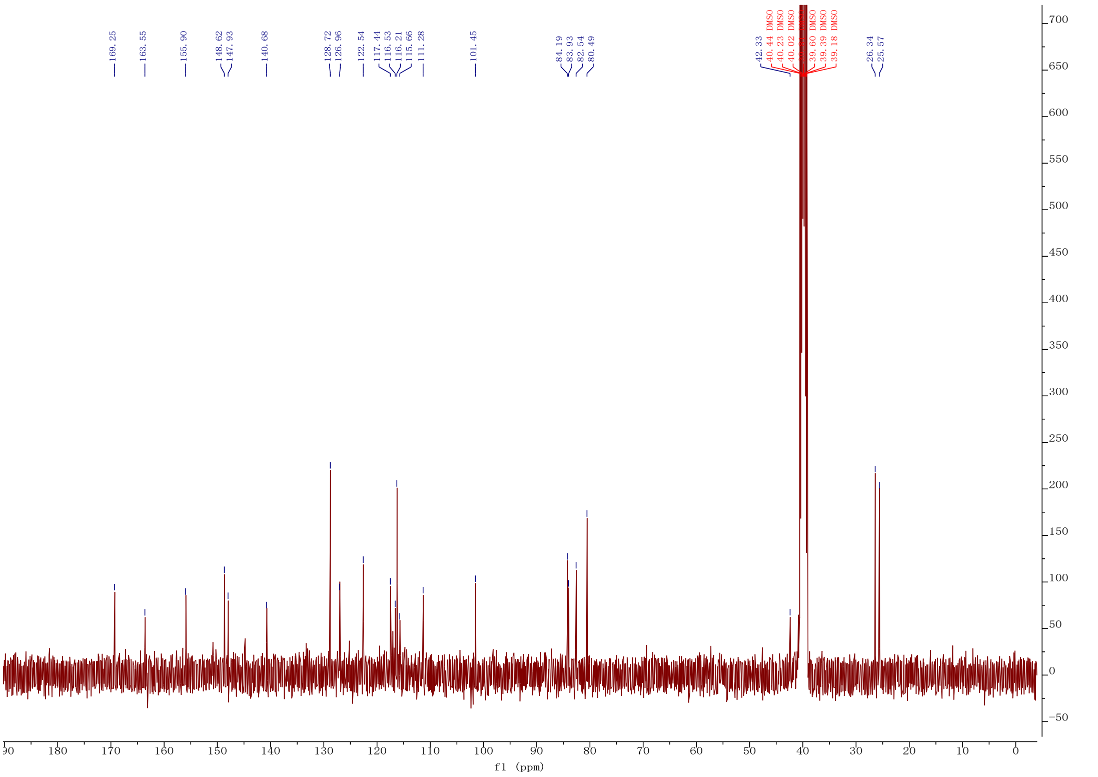
169.25
163.55
155.90
148.62
147.93
140.68
128.72
126.96
122.54
117.44
116.53
116.21
115.66
111.28
101.45
84.19
83.93
82.54
80.49
42.33
40.44 DMSO
40.23 DMSO
40.02 DMSO
39.60 DMSO
39.39 DMSO
39.18 DMSO
26.34
25.57
f1 (ppm)

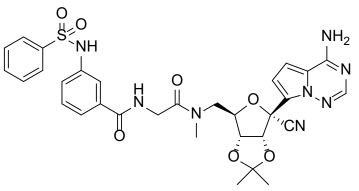
O
O
S
NH
NH2
N
N
N
H
N
O
N
O
CN
O
O
O

**18a**

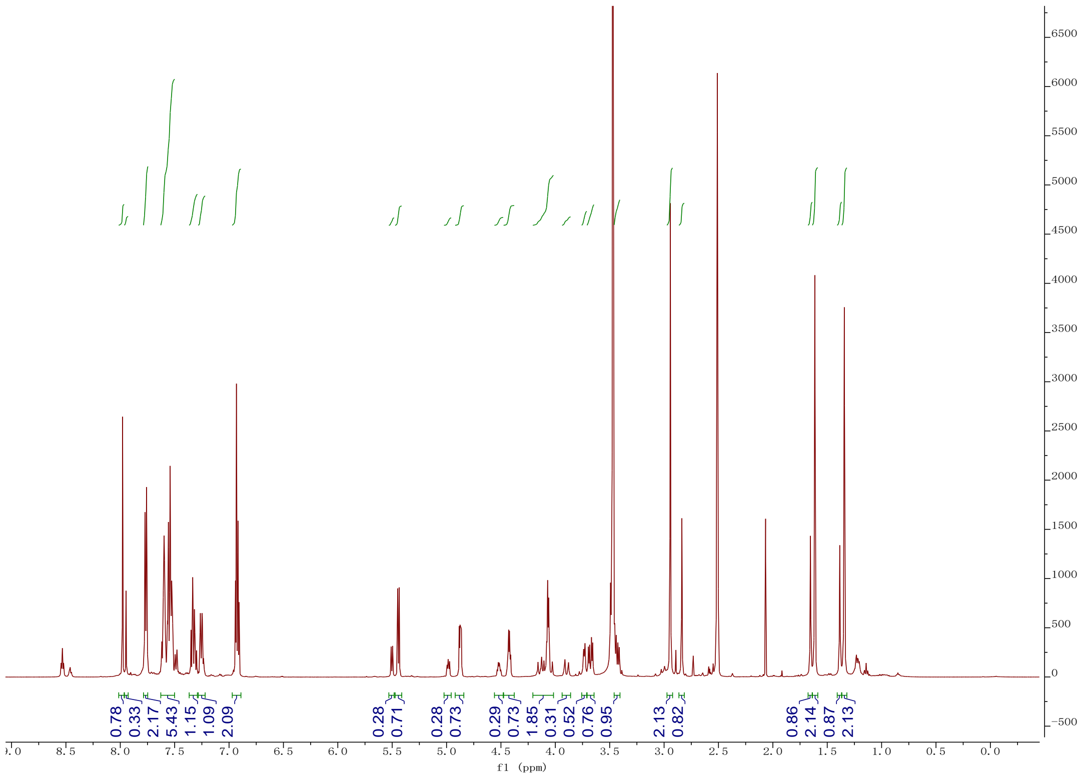
0.78
0.33
2.17
5.43
1.15
1.09
2.09
0.28
0.71
0.28
0.73
0.29
0.73
1.85
0.31
0.52
0.76
0.95
2.13
0.82
0.86
2.14
0.87
2.13
f1 (ppm)

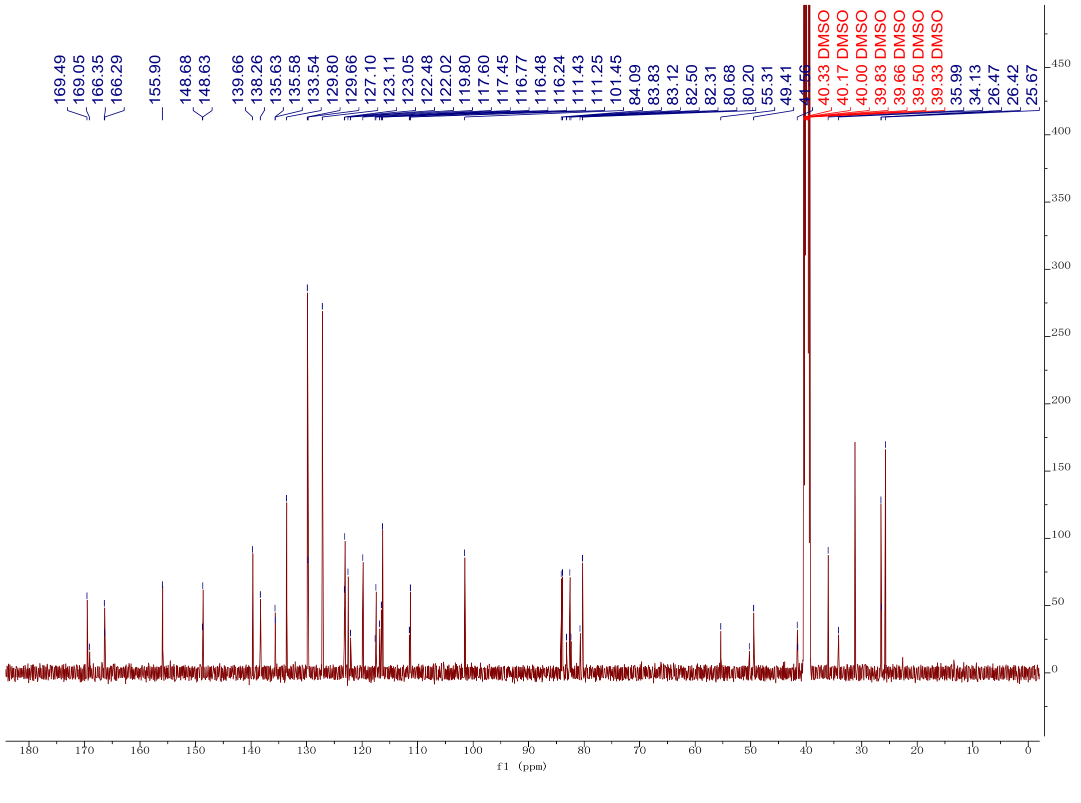
169.49
169.05
166.35
166.29
155.90
148.68
148.63
139.66
138.26
135.63
135.58
133.54
129.80
129.66
127.10
123.11
123.05
122.48
122.02
119.80
117.60
117.45
116.77
116.48
116.24
111.43
111.25
101.45
84.09
83.83
83.12
82.50
82.31
80.68
80.20
55.31
49.41
40.33 DMSO
40.17 DMSO
40.00 DMSO
39.83 DMSO
39.66 DMSO
39.50 DMSO
39.33 DMSO
35.99
34.13
26.47
26.42
25.67
f1 (ppm)

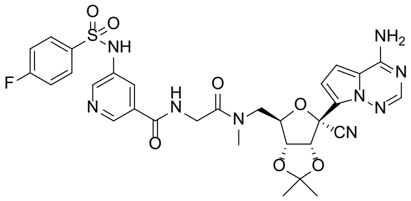

**18b**

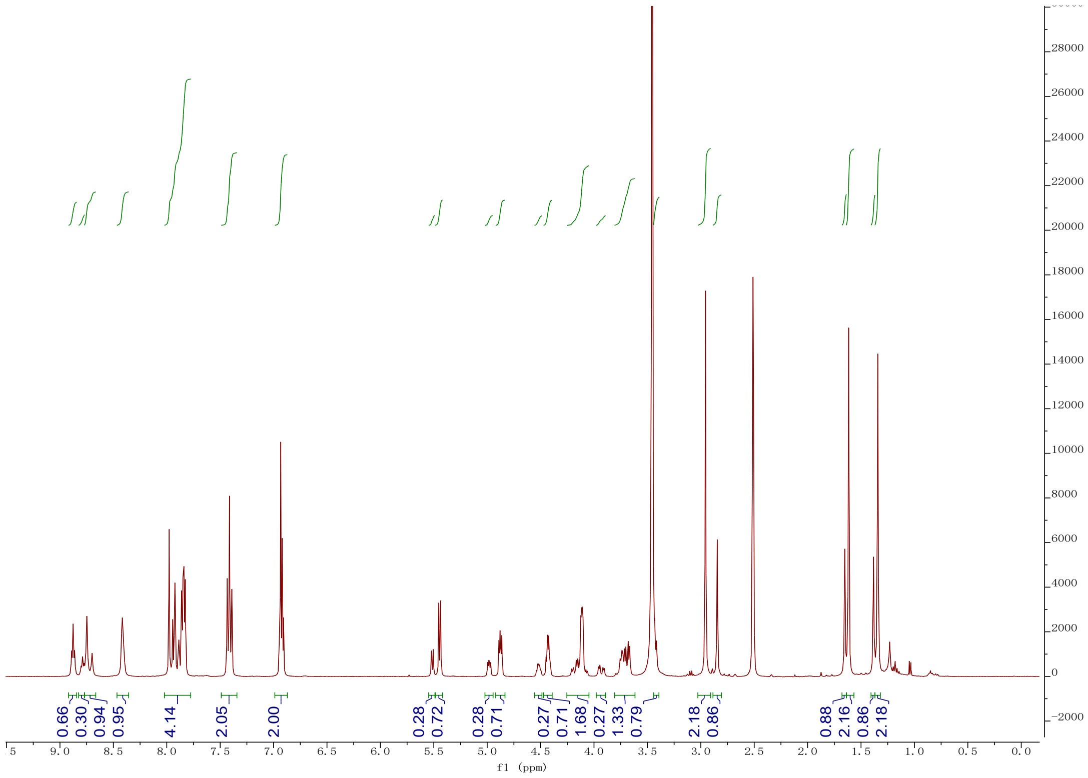

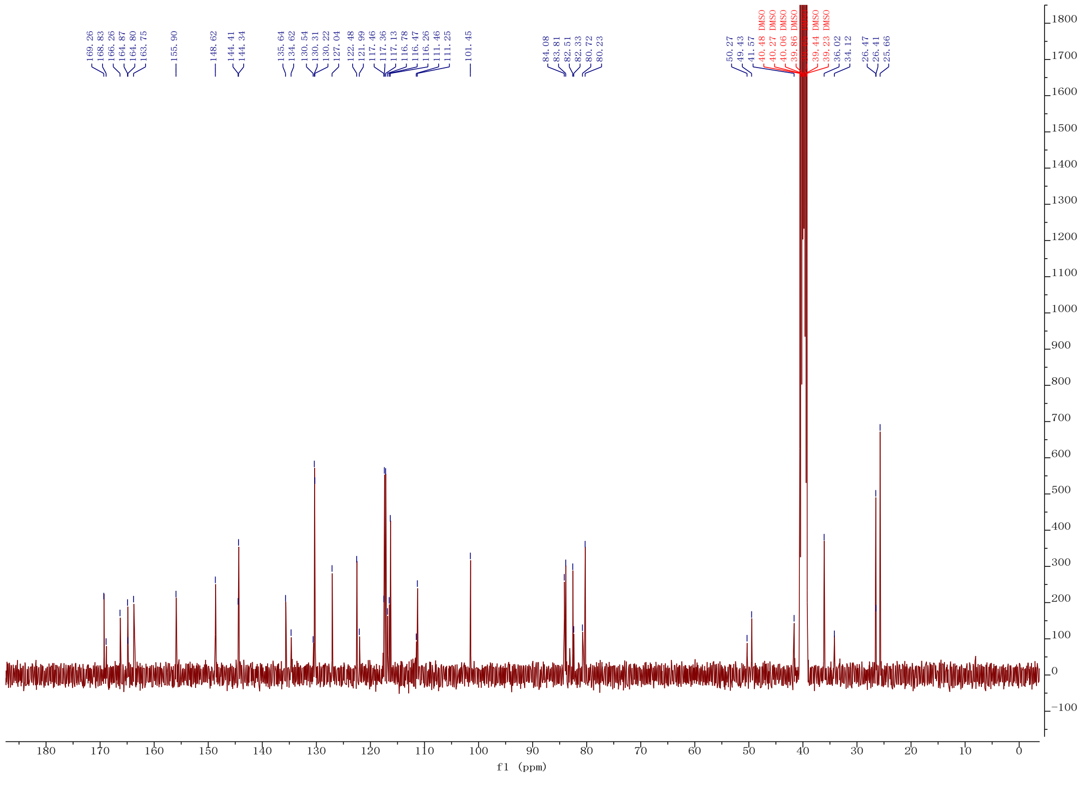

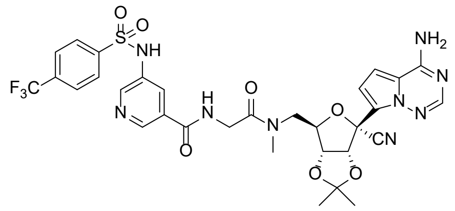

O
S
O
NH
F3C
N
H
N
O
N
O
NH2
N
N
N
O
CN
O
O


**18c**

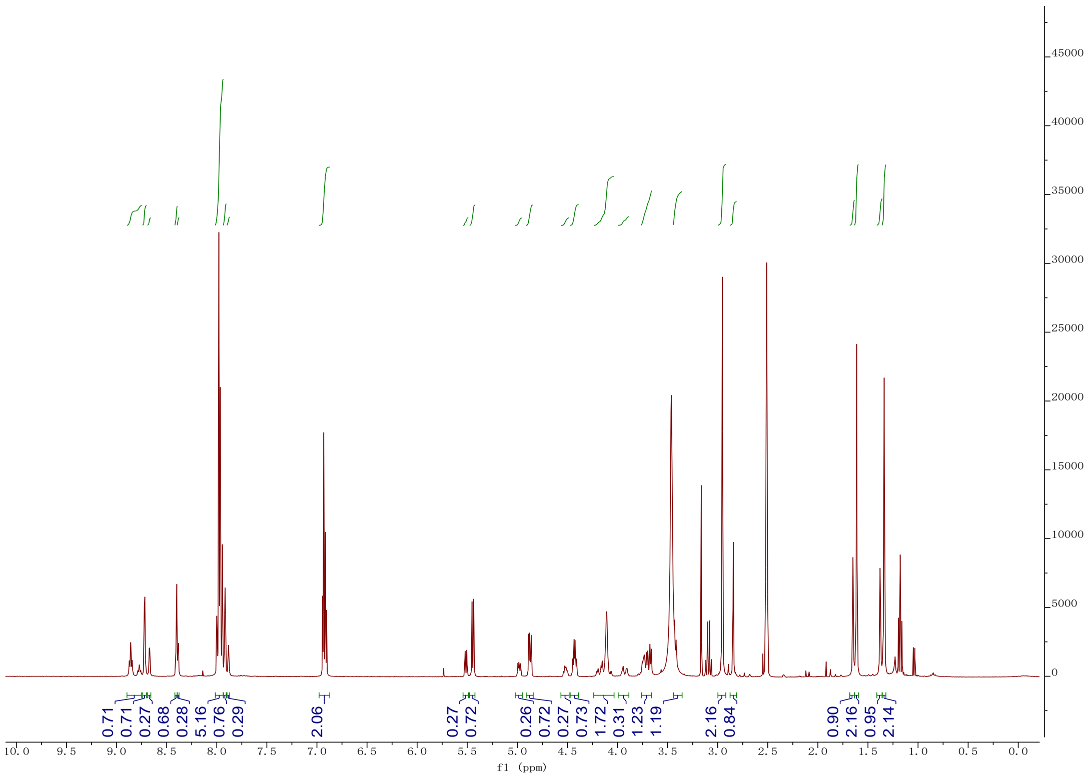

0.71
0.71
0.27
0.68
0.28
5.16
0.76
0.29
2.06
0.27
0.72
0.26
0.72
0.27
0.73
1.72
0.31
1.23
1.19
2.16
0.84
0.90
2.16
0.95
2.14
f1 (ppm)


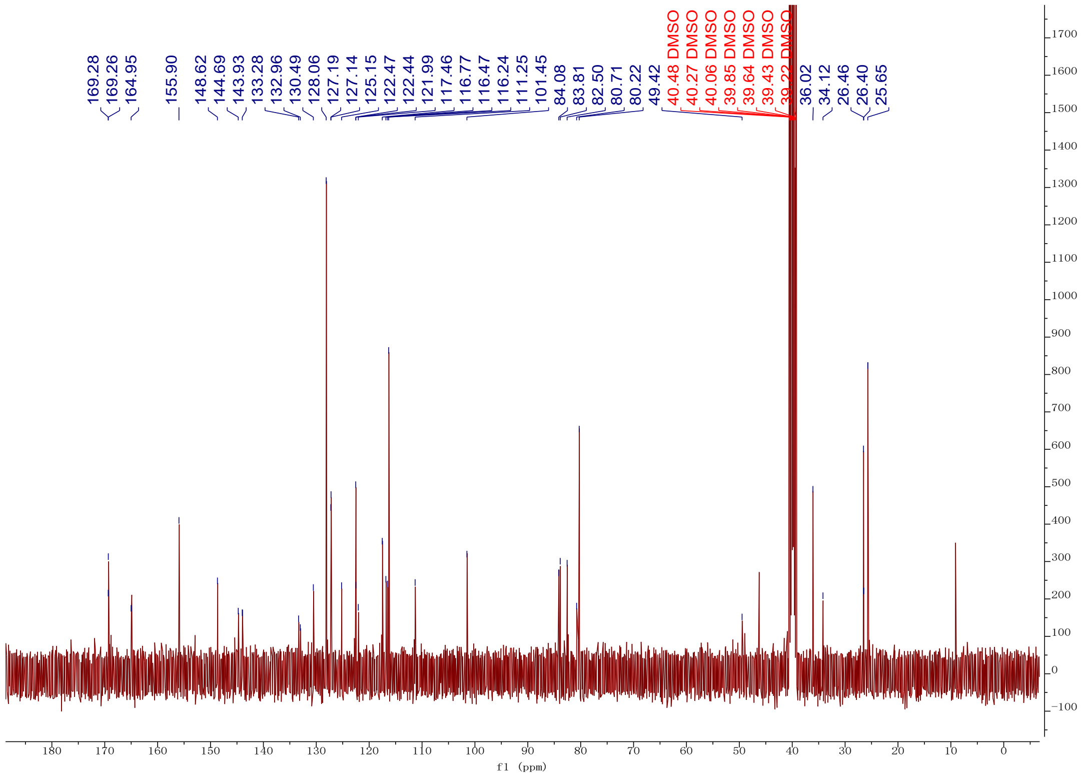

169.28
169.26
164.95
155.90
148.62
144.69
143.93
133.28
132.96
130.49
128.06
127.19
127.14
125.15
122.47
122.44
121.99
117.46
116.77
116.47
116.24
111.25
101.45
84.08
83.81
82.50
80.71
80.22
49.42
40.48 DMSO
40.27 DMSO
40.06 DMSO
39.85 DMSO
39.64 DMSO
39.43 DMSO
39.22 DMSO
36.02
34.12
26.46
26.40
25.65
f1 (ppm)

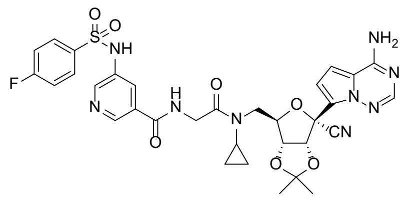

NH
F
N
H
O
N
O
N
O
CN
O
O
N
N
N
NH2


**18d**

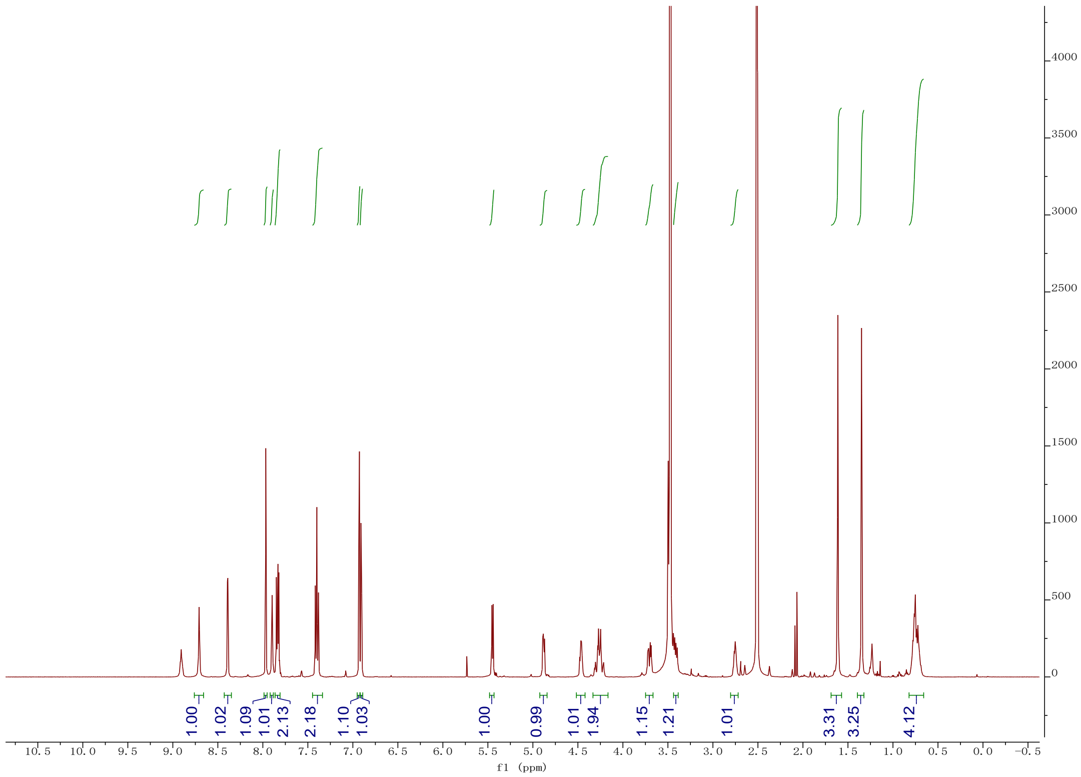

1.00
1.02
1.09
1.01
2.13
2.18
1.10
1.03
1.00
0.99
1.01
1.94
1.15
1.21
1.01
3.31
3.25
4.12
f1 (ppm)


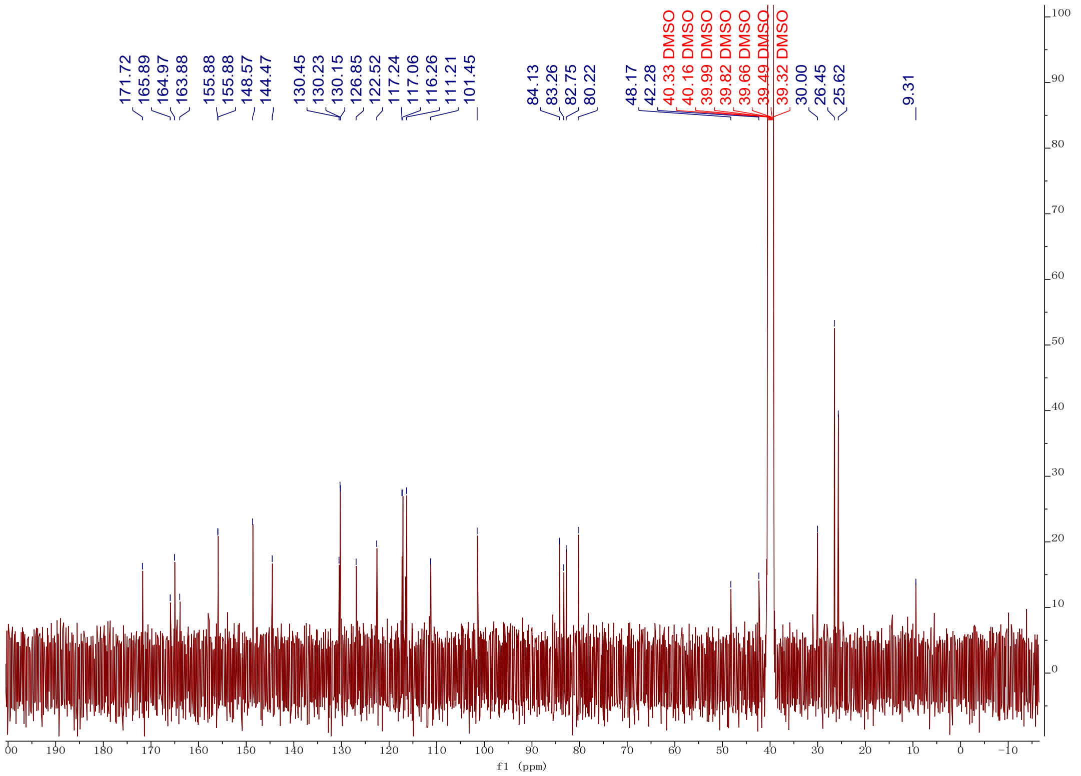

171.72
165.89
164.97
163.88
155.88
155.88
148.57
144.47
130.45
130.23
130.15
126.85
122.52
117.24
117.06
116.26
111.21
101.45
84.13
83.26
82.75
80.22
48.17
42.28
40.33 DMSO
40.16 DMSO
39.99 DMSO
39.82 DMSO
39.66 DMSO
39.49 DMSO
39.32 DMSO
30.00
26.45
25.62
9.31
f1 (ppm)

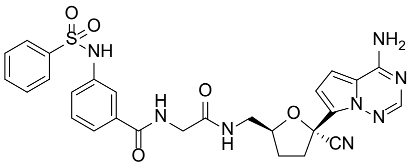
NH2
NH
O
O
S
N
N
N
H
N
O
H
N
O
O
CN

**29a**

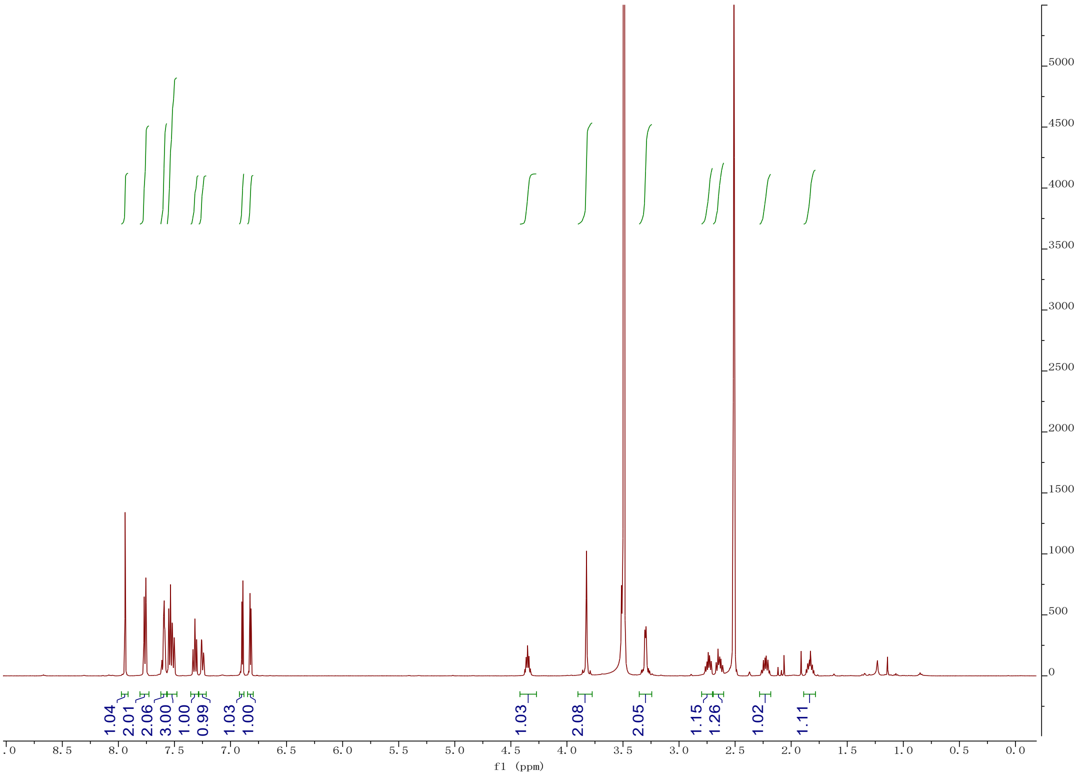
1.04
2.01
2.06
3.00
1.00
0.99
1.03
1.00
1.03
2.08
2.05
1.15
1.26
1.02
1.11
f1 (ppm)

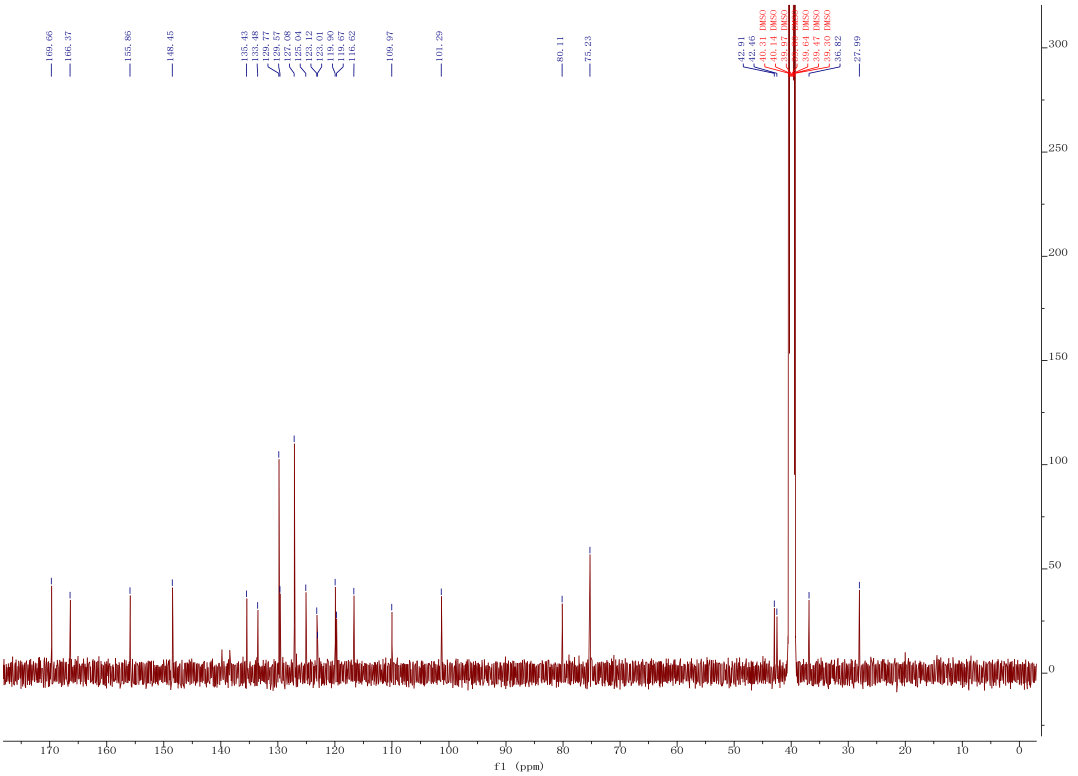
169.66
166.37
155.86
148.45
135.43
133.48
129.77
129.57
127.08
125.04
123.12
123.01
119.90
119.67
116.62
109.97
101.29
80.11
75.23
42.91
42.46
40.31 DMSO
40.14 DMSO
39.97 DMSO
39.64 DMSO
39.47 DMSO
39.30 DMSO
36.82
27.99
f1 (ppm)

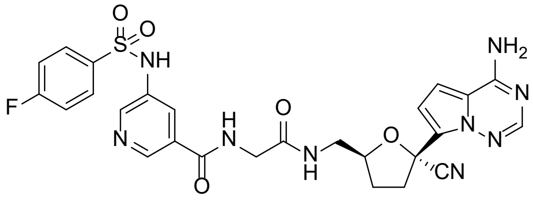

NH
F
N
H
N
O
O
N
H
O
NH2
N
N
N
CN


**29b**

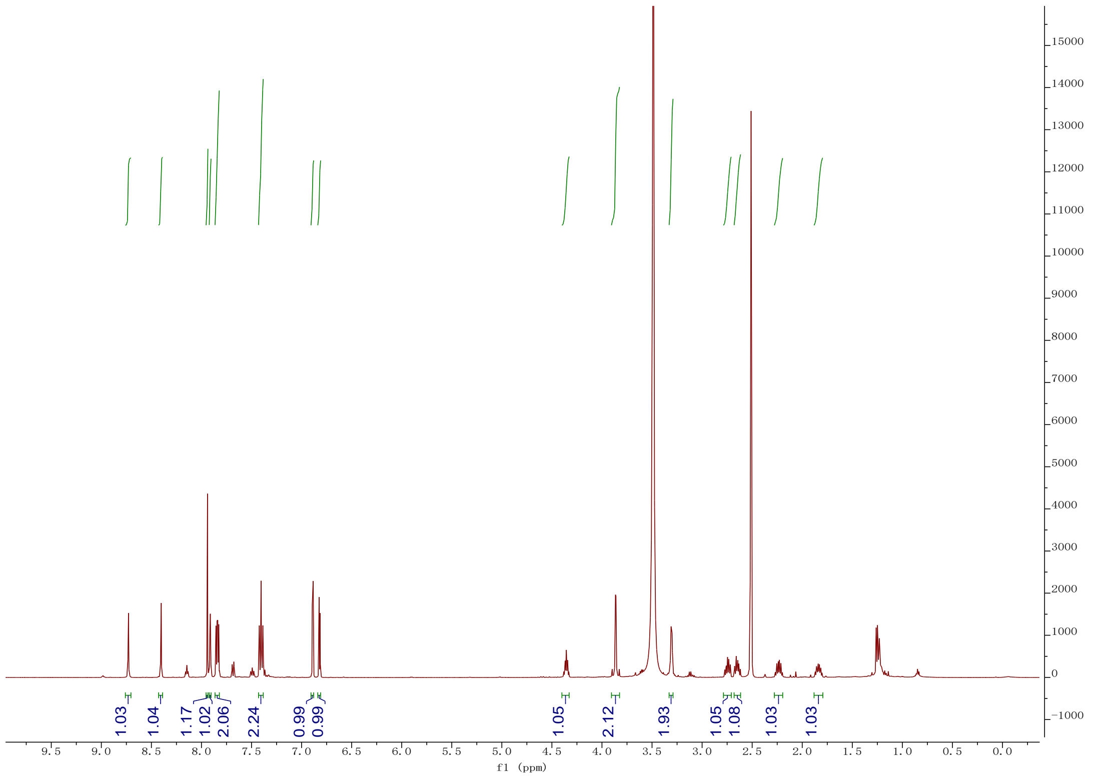

1.03
1.04
1.17
1.02
2.06
2.24
0.99
0.99
1.05
2.12
1.93
1.05
1.08
1.03
1.03
f1 (ppm)


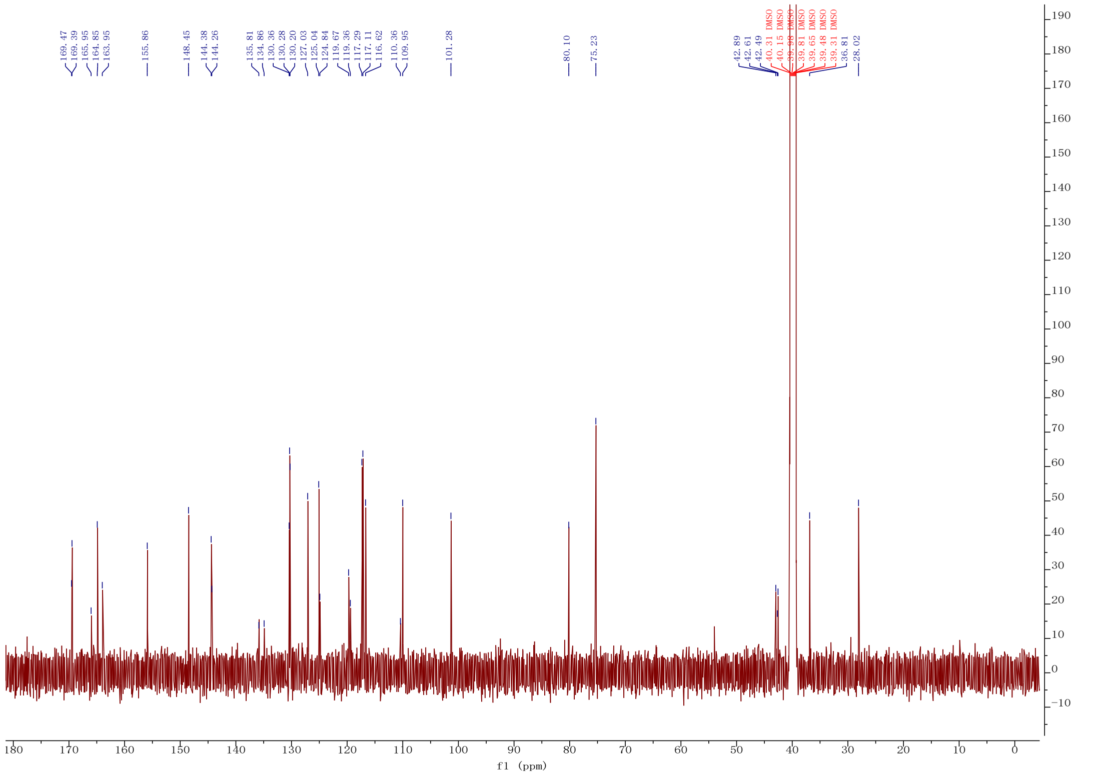

169.47
169.39
165.95
164.85
163.95
155.86
148.45
144.38
144.26
135.81
134.86
130.36
130.28
130.20
127.03
125.04
124.84
119.67
119.36
117.29
117.11
116.62
110.36
109.95
101.28
80.10
75.23
42.89
42.61
42.49
40.31 DMSO
40.15 DMSO
39.81 DMSO
39.65 DMSO
39.48 DMSO
39.31 DMSO
36.81
28.02
f1 (ppm)

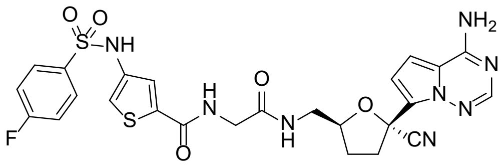
O O
S
NH
F
S
H
N
O
O
N
H
O
CN
NH2
N
N
N

**29c**

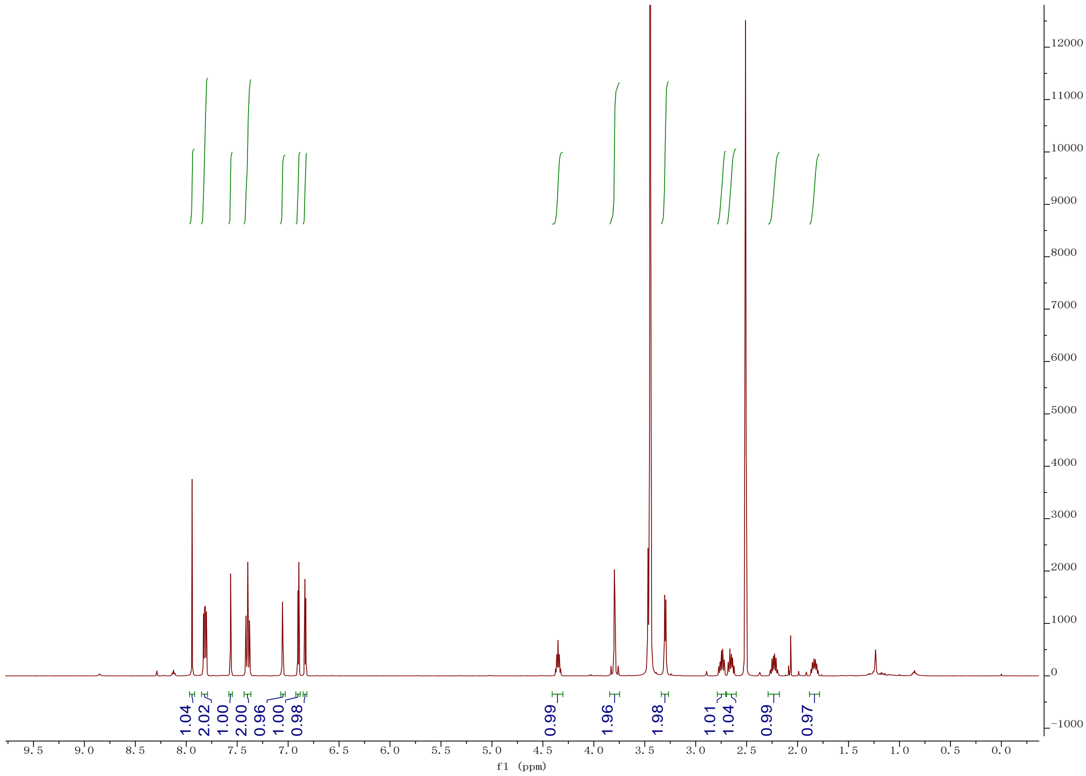
1.04
2.02
1.00
2.00
0.96
1.00
0.98
0.99
1.96
1.98
1.01
1.04
0.99
0.97
f1 (ppm)

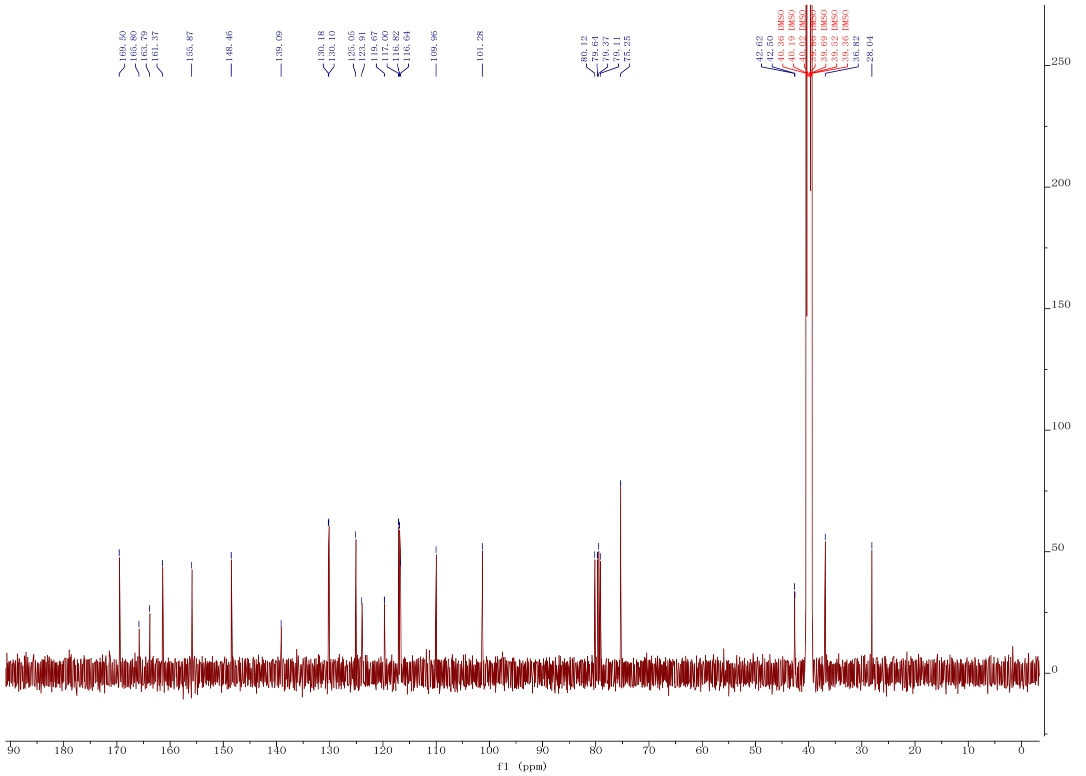
169.50
165.80
163.79
161.37
155.87
148.46
139.09
130.18
130.10
125.05
123.91
119.67
117.00
116.82
116.64
109.96
101.28
80.12
79.64
79.37
79.11
75.25
42.62
42.50
40.36 DMSO
40.19 DMSO
40.02 DMSO
39.86 DMSO
39.69 DMSO
39.52 DMSO
39.36 DMSO
36.82
28.04
f1 (ppm)

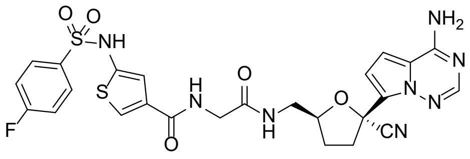
NH2
N
N
N
O
CN
O
H
N
O
N
H
O
S
NH
O
O
S
F

**29d**

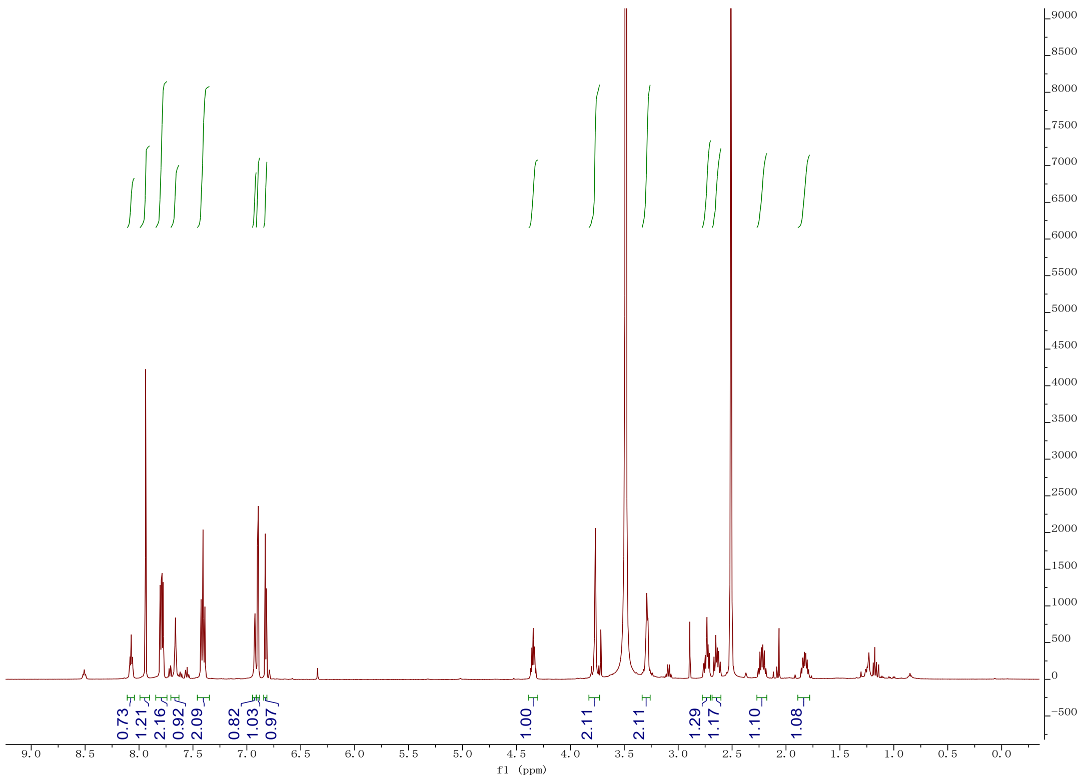
0.73
1.21
2.16
0.92
2.09
0.82
1.03
0.97
1.00
2.11
2.11
1.29
1.17
1.10
1.08
f1 (ppm)

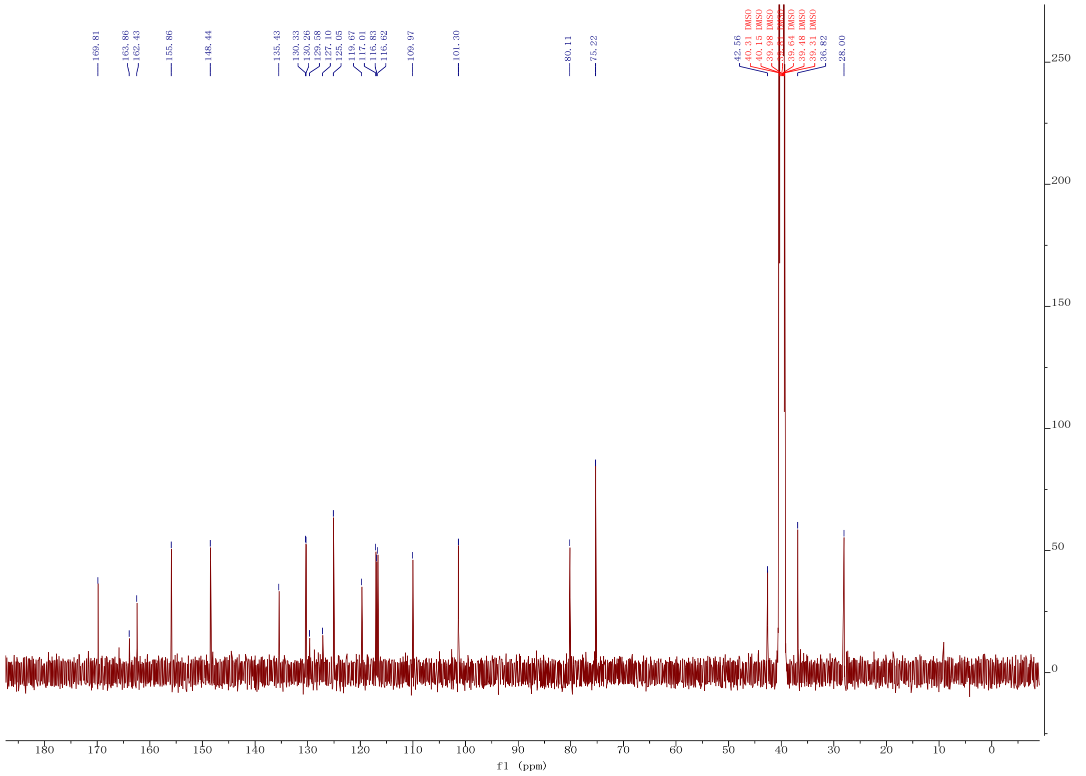
169.81
163.86
162.43
155.86
148.44
135.43
130.33
130.26
129.58
127.10
125.05
119.67
117.01
116.83
116.62
109.97
101.30
80.11
75.22
42.56
40.31 DMSO
40.15 DMSO
39.98 DMSO
39.64 DMSO
39.48 DMSO
39.31 DMSO
36.82
28.00
f1 (ppm)

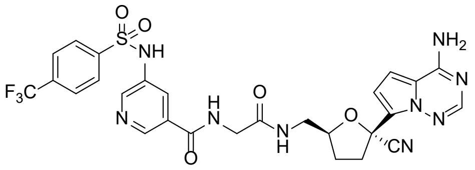
NH2
NH
F3C
N
H
O
O
N
H
O
CN
N
N
N
O
S
O

**29e**

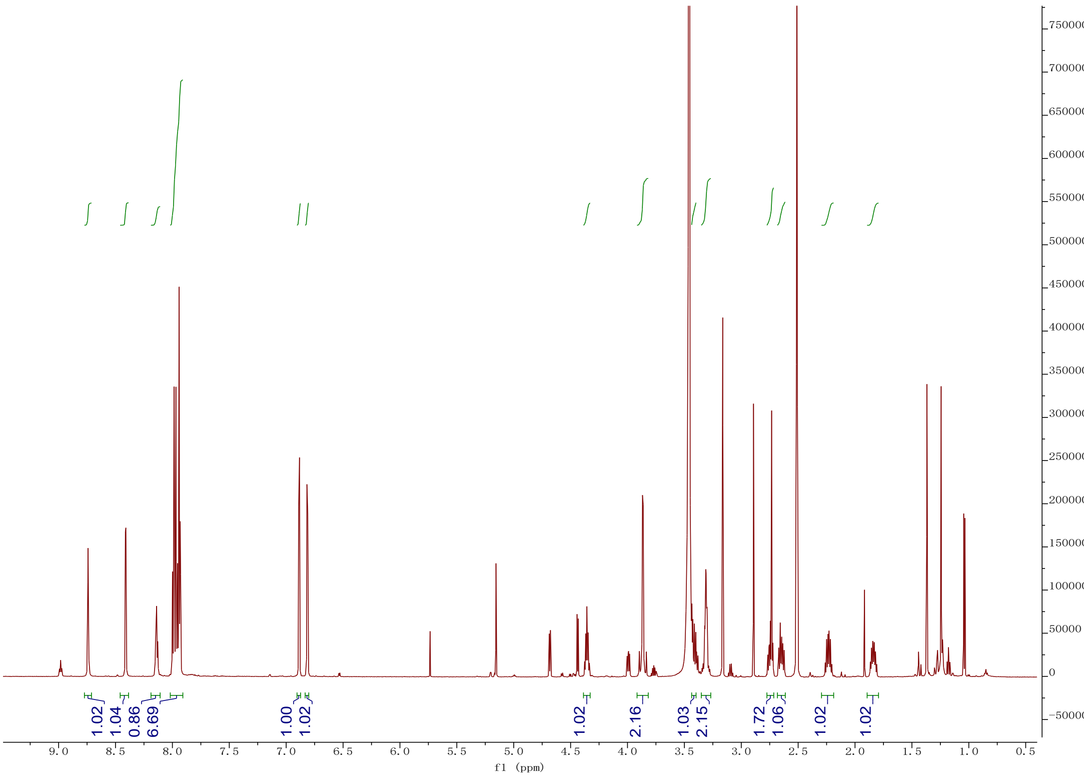
1.02
1.04
0.86
6.69
1.00
1.02
1.02
2.16
1.03
2.15
1.72
1.06
1.02
1.02
f1 (ppm)

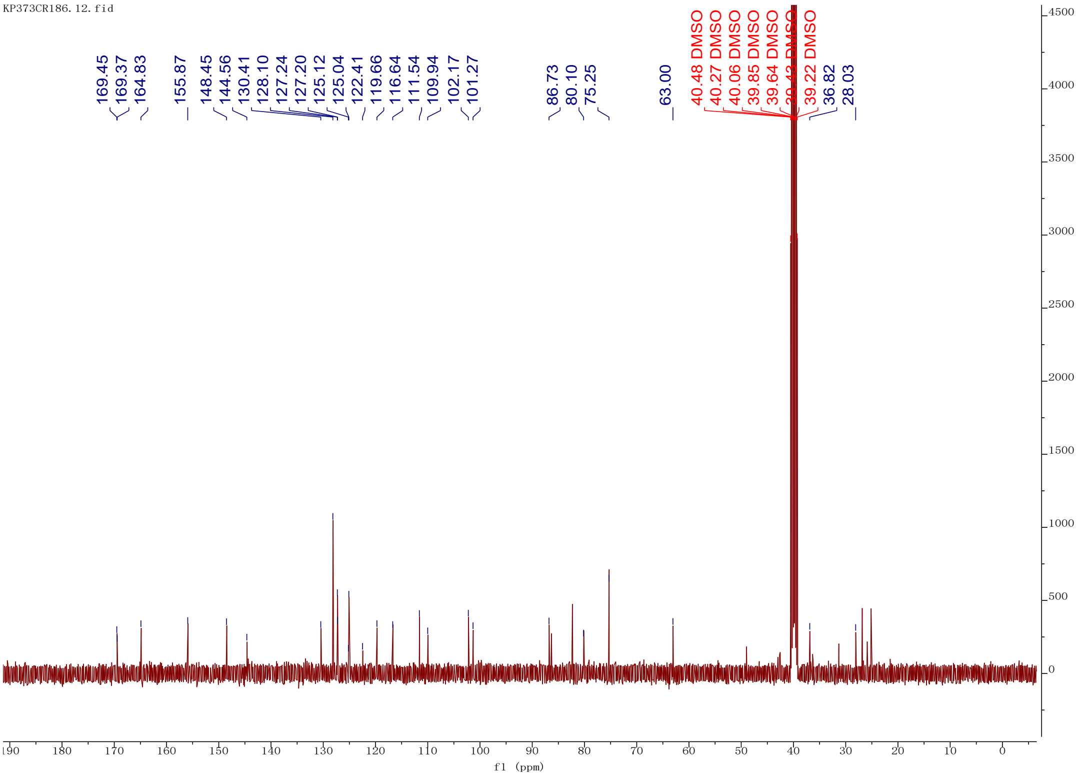
KP373CR186.12.fid
169.45
169.37
164.83
155.87
148.45
144.56
130.41
128.10
127.24
127.20
125.12
125.04
122.41
119.66
116.64
111.54
109.94
102.17
101.27
86.73
80.10
75.25
63.00
40.48 DMSO
40.27 DMSO
40.06 DMSO
39.85 DMSO
39.64 DMSO
39.22 DMSO
36.82
28.03
f1 (ppm)

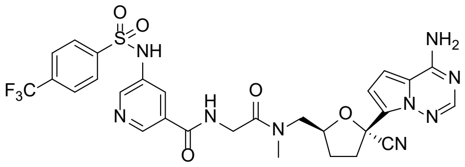


**33**

KP456CR186.10.fid

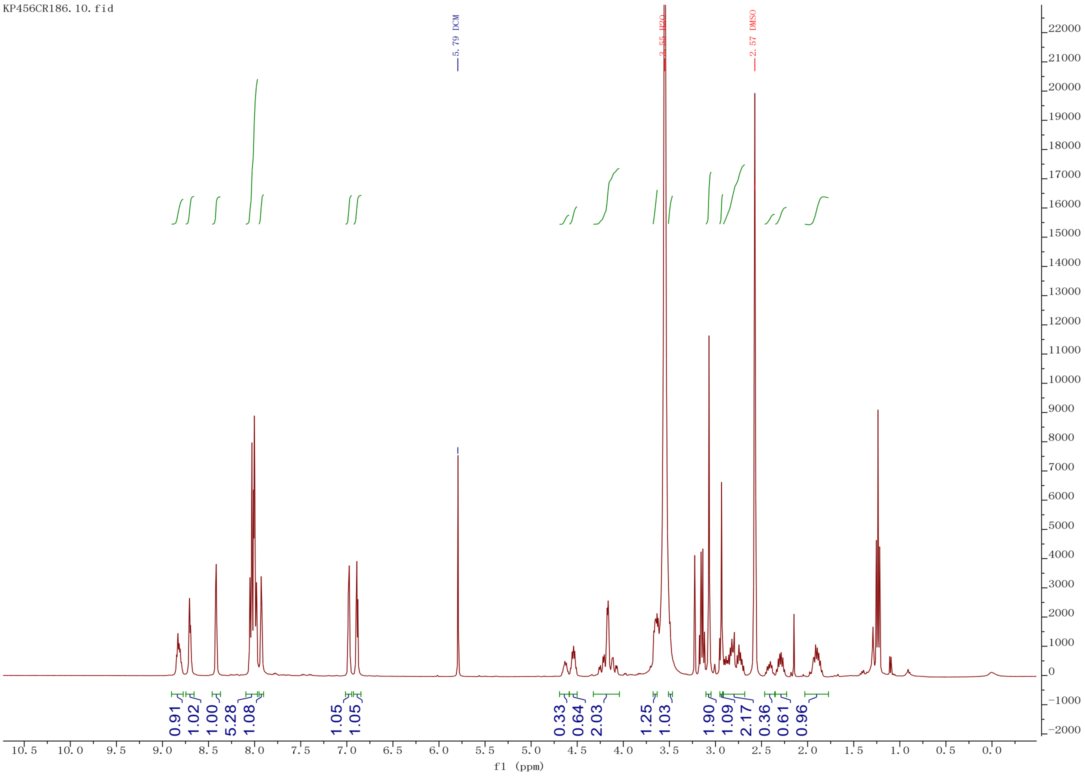


KP456CR186.12.fid

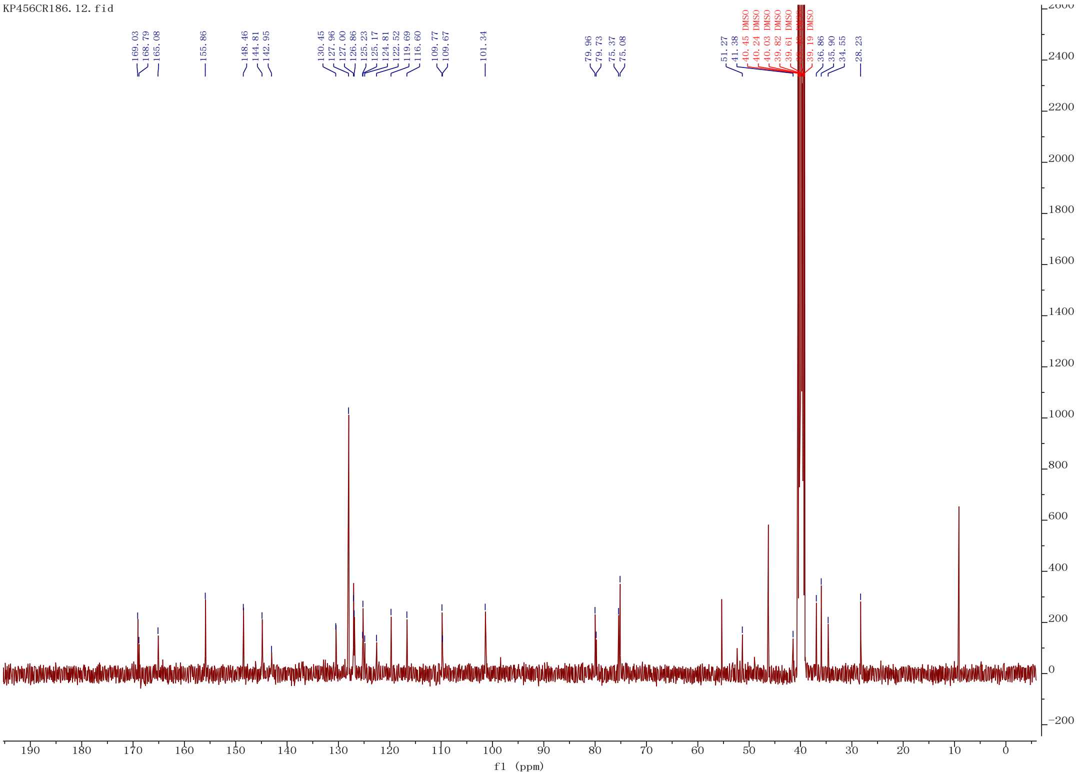
